# Causal inference clarifies the roles of background selection and mutation rate variation in shaping human genetic diversity

**DOI:** 10.64898/2026.06.02.727906

**Authors:** Gustavo V. Barroso, Nicholas W. Collier, Chenlu Di, Kirk E. Lohmueller, Aaron P. Ragsdale

## Abstract

Decades of theoretical and empirical work have struggled to reconcile competing views on how evolution shapes patterns of genetic variation. We now understand the genome as a mosaic molded by both neutral and selective forces, but quantifying their relative contributions remains an open challenge. A major obstacle has been the tendency to analyze each evolutionary process in isolation. But different processes may leave similar signatures on genetic variation, making it challenging to draw conclusions from correlations alone. To address this gap, we make predictions of the landscape of diversity based on background selection and mutation rate variation. We then develop structural equation models describing how mutation, recombination and selection jointly shape the genomic landscape of diversity. This approach offers a more realistic representation of biological interactions and enables rigorous evaluation of hypothesized causal structures, marking both conceptual and practical improvements over previous studies. Analyses of human data reveal large variation in the explanatory power of candidate models across chromosomes. We find that previous studies likely overestimated the predictive accuracy of background selection; although it emerges as the overall main driver of diversity, in some chromosomes mutation rate variation has a comparable impact. We show that recombination increases diversity more through its influence on background selection than its mutagenic effect, resolving a longstanding debate. This work demonstrates that modeling variation inherent in genome biology substantially improves our ability to explain human genetic diversity. At the same time, our results suggest inadequacies in available genomic maps or in background selection as the unique mode of linked selection in humans.

**Significance:** Understanding the mechanisms that create and maintain genetic diversity is fundamental to explaining how populations evolve. Early work emphasized the role of linked selection, but this focus has obscured the contributions of other processes. To overcome this limitation, we introduce structural equation models that encode the causal relationships among mutation, recombination, and selection. We find that background selection is the primary driver of genetic diversity, but its predictive power has likely been overstated in previous studies. Meanwhile, mutation rate variation substantially impacts diversity in some chromosomes. We also identify mismatches between model expectations and observed data that point to limitations in available genomic maps or to the insufficiency of background selection models to fully describe linked selection in humans.

## Introduction

> *In a sense, a description of the genetic variation in a population is the fundamental datum of evolutionary studies; and it is necessary to explain the origin and maintenance of this variation and to predict its evolutionary consequences.*
>
> — Hubby and Lewontin (1966)

In his classic treatment of genetic diversity, Lewontin (1974) steered modern population genetics toward explaining the causes and consequences of genetic variation. By understanding diversity, he argued, we gain a fuller appreciation of the mechanisms of evolution. While early work quantified allozyme variation across species (Lewontin and Hubby, 1966; Lewontin, 1991), subsequent studies turned to the fine-scale patterns of nucleotide diversity (*π*) along chromosomes (Nachman, 2001; Andolfatto and Przeworski, 2000, 2001; Hellmann et al., 2005; Reed et al., 2005; Pouyet and Gilbert, 2021). A central challenge in these efforts is to disentangle the similar signatures that disparate evolutionary processes leave on sequence data.

In an influential study, Begun and Aquadro (1992) showed that *π* increases with local recombination rate across a set of *Drosophila melanogaster* genes, which they interpreted as evidence of genetic hitchhiking (Smith and Haigh, 1974; Birky Jr and Walsh, 1988; Kaplan et al., 1989). This result motivated investigation of linked selection in humans (Payseur and Nachman, 2002), but clarifying the role of recombination would prove more challenging in our species. Hellmann et al. (2003) proposed that the correlation between human recombination and diversity reflects a mutagenic rather than a selective effect. Subsequent studies produced mixed results, anticipating the challenges of the genomic era in elucidating patterns of diversity along the human genome (Hellmann et al., 2005; Reed et al., 2005; Spencer et al., 2006).

More recent work has expanded the focus to the influence of the mutation landscape on *π* (Hodgkinson et al., 2009; Hodgkinson and Eyre-Walker, 2011; Aggarwala and Voight, 2016). Initial estimates suggested that mutation rate variation explains a substantial proportion of human genetic diversity (Harpak et al., 2016; Smith et al., 2018), muddling the picture even further. We now understand the genome as a mosaic shaped by both neutral and selective forces, but quantifying their relative contributions remains an open challenge (Pouyet and Gilbert, 2021; Galtier, 2024; Rodrigues et al., 2024). Each evolutionary process has traditionally been studied in isolation, but the recent availability of high-resolution maps of recombination (Spence and Song, 2019; Barroso et al., 2019; Halldorsson et al., 2019) and mutation (Carlson et al., 2018; Karczewski et al., 2020; Seplyarskiy et al., 2023), alongside mature models of linked selection (Santiago and Caballero, 2016; Barroso and Ragsdale, 2026), enables a framework that jointly considers the influence of multiple biological processes shaping human genetic variation.

Models of background selection (BGS) describe how local genetic diversity is reduced by the continual influx and removal of deleterious variants at constrained loci (Charlesworth et al., 1993; Hudson and Kaplan, 1995a,b; Nordborg et al., 1996; Nordborg, 1997; Santiago and Caballero, 1995, 1998, 2016; Barroso and Ragsdale, 2026). Early work provided convincing evidence that strong negative selection in the ancestral population of humans and chimpanzees shaped the genomic landscape of divergence between the two species (McVicker et al., 2009), presenting an alternative to neutral explanations that were prominent at the time, such as mutation rate variation (Lercher and Hurst, 2002; Hellmann et al., 2003) and reticulated demography (Osada and Wu, 2005; Patterson et al., 2006). Later refinements considered more realistic features of selection, offering quantitative improvements on this classic model. First, Murphy et al. (2022) showed that selective sweeps have negligible effect, and their best-fitting BGS model explained ≈ 60% of the genetic diversity (*R*^2^) at the 1 Mb scale in a cohort of Yoruba individuals. Soon after, by incorporating weak negative selection, Buffalo and Kern (2024) fit a model to explain ≈ 67% of diversity at this scale, arguing that this is close to the theoretical limit of *R*^2^, with the remaining fluctuations in diversity attributable to genealogical noise.

Thus, in the span of 15 years, linked selection went from contending alternative to most compelling component of the baseline model of human genome evolution (Lohmueller et al., 2011; Charlesworth, 2012, 2013; Cutter and Payseur, 2013; Johri et al., 2020, 2023). Yet the studies that have reinforced this view (McVicker et al., 2009; Murphy et al., 2022; Buffalo and Kern, 2024) share important caveats. First, inference of the distribution of fitness effects (DFE) from linked selective effects is likely underpowered. Consequently, fitting DFEs to the landscape of diversity resulted in parameter estimates that are incompatible with the rest of the literature (Williamson et al., 2005; Boyko et al., 2008; Keightley and Eyre-Walker, 2007; Kim et al., 2017) and inflated the correlation between *B* and *π*. Second, recent studies of BGS ignored mutation rate variation, instead normalizing their predictions by local average diversity (Murphy et al., 2022; Buffalo and Kern, 2024). And finally, the correlation between observed and predicted diversity (*R*^2^), used to assess model fit, does not disentangle how mutation, recombination, and selection influence diversity via direct and indirect causal paths, nor can it quantify their relative importance.

Here we address these points simultaneously. We use our recently developed BGS model that accommodates single-population demography, allows for flexible genome annotations and mutation rates, and covers the entire distribution of selection coefficients (Barroso and Ragsdale, 2026); we ground our *B*-value predictions on estimates of the DFE based on the site frequency spectrum (SFS) (Kim et al., 2017; Di et al., 2025) and on independently inferred mutation maps (Carlson et al., 2018; Karczewski et al., 2020; Seplyarskiy et al., 2023); and we use causal inference (Wright, 1920, 1921, 1923; Pearl, 2000; Shipley, 2016) to jointly model the effect of mutation, recombination and background selection on the landscape of diversity. We formalize causal models of genetic variation as Directed Acyclic Graphs (DAGs) and fit structural equation models that incorporate estimation error in the genomic maps and quantify direct and indirect effects along causal paths. We uncover between-chromosome heterogeneity in variance explained and in the relative importance of mutation, recombination and selection to diversity. Our models also raise questions about the accuracy of the available genomic maps and the adequacy of BGS as the unique mode of linked selection in humans.

## Results and Discussion

### New predictions of *B*-values incorporating multiple genomic features

The original background selection models assumed strong selection, independence among constrained sites, and a population at equilibrium (Charlesworth et al., 1993; Nordborg et al., 1996). Subsequent studies have revealed wide distributions of selection coefficients (Boyko et al., 2008; Kim et al., 2017), whereas both simulations (Torres et al., 2020) and numerical predictions (Barroso and Ragsdale, 2026) indicate that fluctuating population sizes can distort *B*-values. This suggests that classic BGS should fit poorly to human data, yet recent work suggests it accounts for at least 60% of the variation in diversity at the 1 Mb scale (Murphy et al., 2022; Buffalo and Kern, 2024). However, the inferred DFE and deleterious mutation rate are poorly constrained in these models, resulting in overfitting and possibly overestimating the effects of background selection.

To refine models of human genetic diversity, we predicted *B*-values using moments++ and bgshr (Barroso and Ragsdale, 2026) across multiple combinations of genomic maps. We considered two annotations of selectively constrained sites (or *elements*): one based on noncoding phastCons sites, which are putatively functional sites predicted from evolutionarily conservation in a 17-way primate phylogeny (Siepel et al., 2005), and another based on promoters and enhancers that were identified from methylation patterns (Ernst and Kellis, 2012; Vu and Ernst, 2022). We incorporated protein-coding exons into both sets, yielding the *CDS* +*phastCons* and *CDS* +*regulatory* groups, and modeled negative selection using DFEs independently estimated from the SFS (Methods). We further explored two models of selection in exons: one that combines all genes under the Kim et al. (2017) DFE and another where they are partitioned into deciles of constraint based on their tolerance to loss-of-function (LoF) mutations (Zeng et al., 2024). In the latter, we used the SFS to infer a DFE separately for nonsynonymous mutations within each set of genes (Table S4).

For each of these four configurations of selected elements, we predicted *B*-values using three mutation maps (Carlson et al., 2018; Karczewski et al., 2020; Seplyarskiy et al., 2023). We first used single-nucleotide mutation rates to estimate diversity in the absence of linked selection (E[*π*_0_]) at both neutral and constrained sites along the genome, thus incorporating the effect of direct selection on *π*_0_. We then used the location of constrained elements, together with their DFE and a fine-scale recombination map (Spence and Song, 2019), to predict equilibrium *B*-values across the autosomal genome, since BGS is well approximated by steady-state dynamics when population sizes have not undergone major fluctuations (Supplemental sections 4.2, 5.4). The equilibrium *N_e_* was estimated separately for each chromosome and model (Methods, Figure S57), enabling moments++ to accurately represent the effect of drift on *B*-values and bgshr to correct for selective interference (Figure 1). Expected diversity is then E[*π*] = *B* × E[*π*_0_]. When estimating *π̂* (using the high-coverage 1000 Genomes Project cohort of Yoruba individuals from Ibidan, Nigeria (YRI), Byrska-Bishop et al., 2022), we average over all sites within a window, deviating from previous work which focused exclusively on putatively neutral sites (Murphy et al., 2022; Buffalo and Kern, 2024). Predicting *B* in constrained sites allows our framework to better capture the weak-to-moderate selection regime (*N_e_s* ∼ 1), where diversity reductions are sharp but narrow, extending just over hundreds of base pairs (Barroso and Ragsdale, 2026).

**Figure 1:**
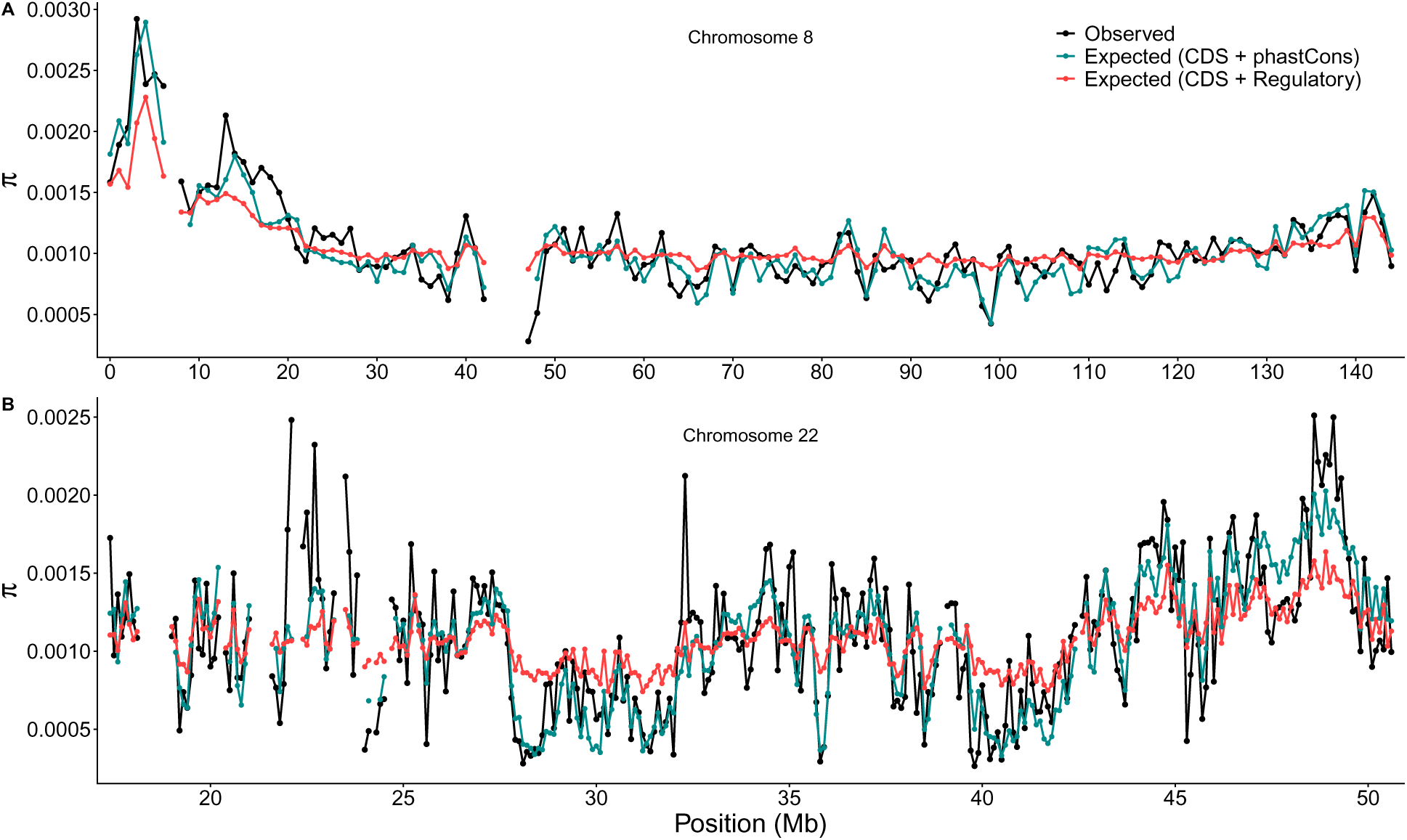
Observed and predicted landscapes of genetic diversity (E[*π*] = *B* × E[*π*_0_] under the Roulette mutation map). Cyan: E[*π*] predicted using *phastCons* elements. Red: E[*π*] predicted *regulatory* elements. In both cases, the exonic DFE is partitioned into deciles of constraint. **A**: chromosome 8 at the 1 Mb scale, where the Roulette map helps accommodating the peak in diversity at the left arm. **B**: in chromosome 22, our *phastCons* model explains ≈ 70% of diversity at the 100 kb scale. See Supplemental File for other landscapes.

We find that the *B*-value map changes little when the exonic DFE is partitioned by tolerance to LoF mutations (Figure S18). However, since they assign more weight to strongly selected sites, models that partition genes are able to capture larger reductions in diversity due to unlinked BGS (Table S6). Therefore, we showcase results from split exonic DFEs. The three mutation maps lead to *B*-values that are highly similar within annotation groups (pairwise correlations *>* 0.99). To succinctly examine their causal effects on diversity (Figure 2), we construct latent representations of *B* and *µ* (Methods). Finally, the two annotation sets predict *B*-values that diverge in several genomic regions (Figures 1, S17), as we discuss next.

**Figure 2:**
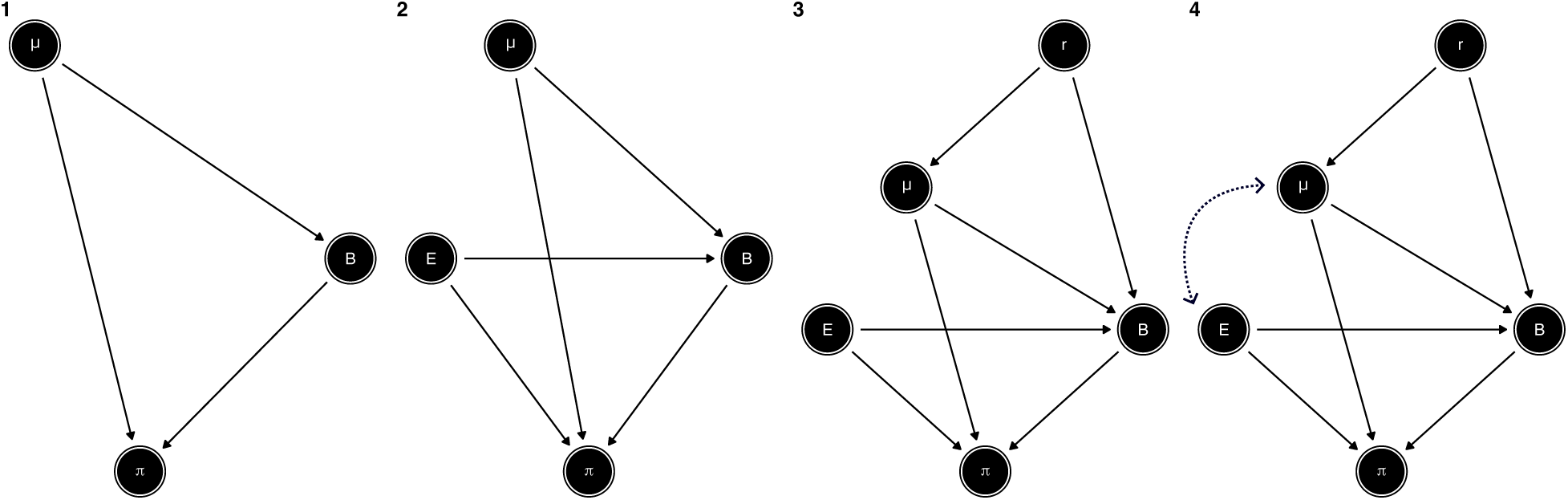
Candidate models describing causal relationships among human genomic landscapes. Straight arrows represent direct, linear effects of any sign. *π*: genetic diversity; *B*: *B*-values; *µ*: mutation rate; *E*: density of sites under direct selection; *r*: recombination rate. The curved arrow between *E* and *µ* in DAG 4 represents a free covariance parameter that is fixed to zero in DAG 3.

### Genome-wide correlations and goodness-of-fit of predicted landscapes

To assess how well linked selection predicts genetic diversity, pioneering work correlated levels of polymorphism with recombination rate and density of selected sites (Begun and Aquadro, 1992; Payseur and Nachman, 2002; Hellmann et al., 2003; Lohmueller et al., 2011). These proxies were later replaced with model expectations (*B*-values), which offered a better fit to the observed landscape of *π* (Murphy et al., 2022; Buffalo and Kern, 2024). An important finding has been that modeling background selection using phylogenetic conservation scores predicts local diversity better than models prioritizing functional annotations (such as exons, introns and UTRs) (Buffalo and Kern, 2024), highlighting a trade-off between predictability and interpretability.

We found higher genome-wide correlations between *π* and *B*-values for *CDS* +*phastCons* (*R*^2^ = 0.51) than for *CDS* +*regulatory* (*R*^2^ = 0.27) models at the 1 Mb scale (Figures S20, S21A). Although benchmarking shows that moments++ improves on earlier BGS models (Barroso and Ragsdale, 2026), our predictive power is lower than reported by Murphy et al. (2022) (*R*^2^ = 0.60) and Buffalo and Kern (2024) (*R*^2^ = 0.67). We attribute this discrepancy to our use of independent estimates of mutation rates and the DFEs, which mitigates overfitting. Notably, Murphy et al. (2022) restricted selection to the top 6% phastCons scores and observed declining correlations with other thresholds – a striking difference from our use of the top 60% noncoding scores (≈ 30% of the genome, Methods and Supplemental section 5.6), motivated by the inclusion of phastCons elements with reliably inferred DFEs (Di et al., 2025). In contrast, the best-fitting model of Buffalo and Kern (2024) combines classes of selection targets (e.g., phastCons, exons, introns) in a priority scheme to resolve overlapping annotations. Although their BGS model incorporates weak selection, their inferred selection coefficients for phastCons are roughly an order of magnitude higher than in Murphy et al. (2022), underscoring the difficulties of fitting selection models from the landscape of pairwise diversity alone.

Incorporating mutation rate variation into the prediction of diversity (E[*π*] = *B* × E[*π*_0_]) increases genome-wide correlations (Figure S21). For *CDS* +*phastCons* models, *R*^2^ = 0.62 (Roulette), *R*^2^ = 0.56 (gnomAD) and *R*^2^ = 0.50 (Carlson), whereas for *CDS* +*regulatory* models *R*^2^ = 0.57 (Roulette), *R*^2^ = 0.38 (gnomAD) and *R*^2^ = 0.37 (Carlson) at the 1 Mb scale (Figure S20). The larger *R*^2^ using *CDS* +*phastCons* models compared to *CDS* +*regulatory* models suggests that the regulatory annotations we use do not fully capture selection in the noncoding genome (Table S7, Supplemental section 5.7). Collectively, these results indicate that mutation rate variation accounts for a non-negligible fraction of human diversity, in agreement with previous work (Harpak et al., 2016; Aggarwala and Voight, 2016; Smith et al., 2018). However, such correlation measures conflate paths of influence and do not represent the proportion of variance in *π* explained by individual predictors. To obtain models that are not only predictive but also explanatory, we aim to tease apart causal effects.

### Proposing causal models of genetic diversity

Causal inference provides methods to identify cause-effect relationships from observational data (Denis and Legerski, 2006; Pearl, 2010; Shipley, 2016; Pearl and Mackenzie, 2018). Among them, structural equation modeling (SEM) assesses the goodness-of-fit of candidate Directed Acyclic Graphs (DAGs) and estimates path coefficients which can be decomposed into direct and indirect effects of one variable on another. Although their origins trace back to Wright (1920, 1921, 1923), these tools have been underused in population genetics, in part because of the positivist posture adopted by RA Fisher (Pearl and Mackenzie, 2018; Dong, 2024). The perceived subjectivity of causal claims, together with the demanding assumptions of causal modeling, contributed to Fisher’s dominance in shaping the methodological trajectory of population genetics (but see a recent application regarding the drift-barrier hypothesis, Weinstein and Roy, 2026). Yet Wright’s causal inference represents the gold standard for statistical inference in ecology and in the social sciences, where most of its modern development took place (Pugesek and Tomer, 1996; Shipley, 2016).

We represent the dynamics among population genetic processes as DAGs where nodes are variables that we can quantify. Barroso and Dutheil (2023) proposed that drift, recombination and selection modulate coalescence times (*τ*), which drive diversity together with mutation rates (*µ*) (Figure S22). They cast this model as a compact linear regression *π* = *β*_1_*µ* + *β*_2_*τ* + *β*_3_*µ* : *τ* + *ε* that estimates independent effects on a single outcome, *π*. Here we introduce two improvements to this framework. First, we adopt our explicit model of linked selection, moments++ (Barroso and Ragsdale, 2026). Second, we fit candidate DAGs using SEM, where genomic maps of recombination, mutation and constrained elements serve both to predict *B*-values and as nodes in causal graphs (Figure 2), capturing interactions among variables beyond their direct effects on *π*.

1. DAG 1 is inspired in the model of Barroso and Dutheil (2023), except *B* represents the expected reduction in *τ* due to linked selection, so that genealogical noise (drift) is relegated to the residuals. *µ* → *B* indicates that mutation rates modulate *B*-values because of their effect on deleterious diversity.
2. DAG 2 extends DAG 1 by including the density of constrained sites, *E*, averaged across elements. We consider various classes of elements, each following its own DFE. Direct negative selection affects diversity at constrained sites, decreasing the local average (*E* → *π*) and intensifying BGS (*E* → *B*).
3. DAG 3 includes recombination rate (*r*), which indirectly increases diversity via two pathways: alleviating linked selection (*r* → *B* → *π*); and increasing mutation rates (*r* → *µ* → *π*) (Hellmann et al., 2003).
4. DAGs 1-3 assert that exogenous variables (those without incoming arrows) have uncorrelated residuals. To formally test the hypothesis of differential mutation rates in constrained elements (Rodriguez-Galindo et al., 2020), we include a free covariance term between *E* and *µ* (DAG 4).

We first used forward simulations to benchmark SEM’s performance in disentangling patterns of variation along the genome (Supplemental section 4), then fit these candidate DAGs using human data.

### Chromosome-specific signatures of linked selection

In biological systems, observed variables are often noisy estimates of underlying constructs, and failing to account for this measurement error can bias downstream analyses (Gustafson, 2021). SEM addresses this problem by explicitly modeling latent variables and their associated error, allowing path coefficients and fit indices to be estimated with reduced bias when the measurement model is correctly specified. We addressed estimation error in landscapes with models that yielded latent representations of *µ* and *B*-value maps (Methods). We then fit candidate DAGs as path models (Figure 2) using observed diversity (*π̂*) in YRI, averaged in non-overlapping windows of 10 kb, 100 kb or 1 Mb.

Previous work quantified *R*^2^ as the squared correlation between genome-wide landscapes of diversity and *B*-values (Murphy et al., 2022; Buffalo and Kern, 2024). This approach obscures chromosome-specific effects driven by differences in *µ*, *r*, and *E* landscapes, or by unmeasured confounders. Using a likelihood ratio test (LRT), we found that chromosome-specific models fit the data significantly better than restricted models which estimate genome-wide effects (Table S5). The strong support for chromosome-specific models indicates that the relative contributions of mutation, recombination and selection to diversity differ among human autosomes. This can happen if the molecular architecture shaping these landscapes is such that their mean, variance and autocorrelation differ across chromosomes (Supplemental section 8.3). Dissecting these differences is left for future work, and further discussion can be found in Barroso and Dutheil (2023). Hereafter, we focus on chromosome-specific results.

### Large variation in 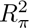 among autosomes

Using SEM, we computed 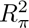 by tracing the composition of linear effects along causal chains (Bollen, 1987; Shipley, 2016). This metric is analogous to variance explained in linear regression but incorporates the full DAG structure through its dependence on the covariance matrix (Supplemental section 8.2). Hence 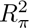 differs from the traditional *R*^2^ primarily because it accounts for the variance in diversity explained by the paths *µ* → *π* and *E* → *π* (Figure S24). SEM estimates the proportion of variance explained in each endogenous variable in the DAG, providing further insight into the explanatory power of the model (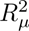, 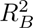, Figures S40, S39), though here we focus on 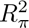.

We find large variation in 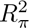 among autosomes (Figure 3A,B), ranging from 0.12 to 0.5 at 10 kb; from 0.25 to 0.7 at 100 kb; and from 0.37 to 0.86 at 1 Mb, depending on the DAG and annotations considered. At 10 kb and 100 kb, our causal models explain more variation in diversity than Murphy et al. (2022) as moments++ accurately predicts *B*-values in the weak-to-moderate selection regime and because our DAGs embed high-resolution genomic maps. On the other hand, at the 1 Mb scale, the high 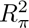 observed in chromosomes 8, 16, 18 and 22 (Figure S38) challenges Buffalo and Kern (2024)’s suggestion that drift imposes an upper limit of ≈ 0.67, a bound likely due to their derivation assuming a sample size of two genome copies. Like wider windows, larger sample sizes also reduce the contribution of genealogical noise to variation in *π*, which we confirm with simulations (Supplemental sections 4, 6, Figure S15). To assess the significance of the between-chromosome differences, we fit SEMs on resampled chromosome windows. Although bootstrapped distributions of 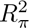 overlap at the 1 Mb scale (Figures S25–S27), presumably due to the small number of windows, they remain largely separate at 10 and 100 kb, suggesting that the contrast in the original data reflects biological differences among chromosomes and not stochastic fluctuations. This is corroborated by the homogeneous 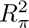 distributions on permuted chromosome labels (Figures S28–S30), motivating the search for biological explanations of the distinct explanatory power of candidate models across autosomes (Supplemental section 7).

**Figure 3:**
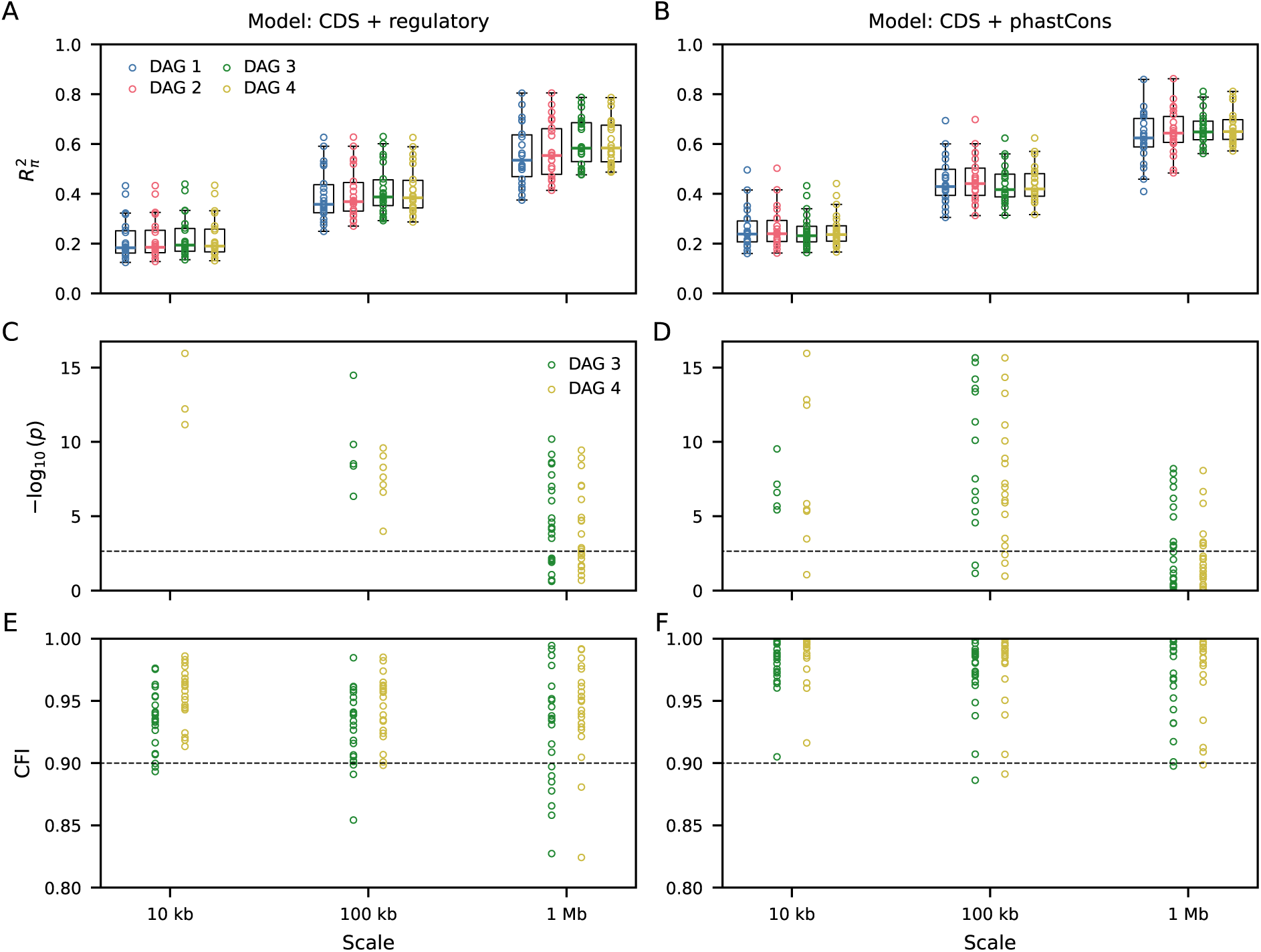
Summaries of structural equation models of genetic diversity. Points show results for each autosome, divided by group of elements and genomic scale. **A-B:** variance in genetic diversity explained by DAGs 1-4. We report 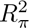, which differs from the squared correlation between E[*π*] and *π̂* (*R*^2^) as it takes the full DAG into account. **C-F:** goodness-of-fit measures of DAGs 3-4, which have sufficient degrees of freedom to assess model fit. C and D show *p*-values from the *χ*^2^ test (-*log*_10_ scale), where lower values indicate better agreement with data. Dashed lines show significance threshold after Bonferroni correction (0.05/22). E and F show comparative fit indices, where higher values indicate better agreement with data. Dashed lines show the rule-of-thumb for an acceptable fit (0.90). Figures S51–S54, S38 show results specified per chromosome label.

Chromosomes with lower 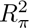 underscore limitations in available data and models. While Murphy et al. (2022) showed that selective sweeps have a negligible impact on the human landscape of *π* (Hernandez et al., 2011), selection on quantitative traits is widespread across the genome (Sella and Barton, 2019; Zeng et al., 2021; Simons et al., 2025). Stabilizing, fluctuating, or balancing selection each affect diversity differently than directional selection against deleterious mutations, and various modes of selection interact with background selection (Pardiñas et al., 2018; Li and Berg, 2026). Associative overdominance may also increase diversity in regions of low recombination and recessive deleterious variants (Ohta, 1971; Zhao and Charlesworth, 2016). Although we observe no clear discrepancy between *π̂* and E[*π*] in candidate regions identified by Gilbert et al. (2020) (Supplemental File), future efforts are needed to evaluate this and other explanations for the low 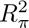 observed in some chromosomes, including missing functional annotations and estimation errors in the genomic maps.

### Proposed causal models are compatible with human data

While 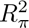 measures the ability of candidate models to predict diversity along the human genome, it is only an indirect assessment of model adequacy. SEM offers formal methods to quantify goodness-of-fit and perform model selection (Pearl, 2000; Shipley, 2016), such as the *χ*^2^ test and the Comparative Fit Index (CFI) which complement each other in model evaluation (Supplemental section 8). We find that the *χ*^2^ test is often unable to reject the proposed models at the 1 Mb scale (Figure 3C,D). DAG 4, which models spatial covariance between mutation rates and density of constrained sites, is compatible with the data from 15 out of 22 autosomes when considering *CDS* +*phastCons* elements. This includes not only short chromosomes (after filtering for coverage, 33 windows remain for chromosome 21) but also chromosomes 1-3 (*>* 150 windows). *CDS* +*regulatory* elements observe more frequent rejection at the 1 Mb scale. On the other hand, at 10 and 100 kb sample sizes increase drastically and the *χ*^2^ test captures small differences in the covariance matrices, leading to widespread rejection.

CFI is less sensitive to sample size, suggesting strong fits even at smaller scales (Figures 3E,F, S32), likely due to increased biological resolution. Averaging within larger windows reduces estimation noise on the one hand but dilutes local signals of joint heterogeneity on the other. For example, 10 kb windows densely packed with constrained sites leave a reliable signature in the covariance matrix that disappears at larger scales as neighboring sites regress to the mean. Such strong, short-range signals are overwhelmed by sample size in *χ*^2^ tests, but are rescued by CFI. Therefore, despite some mismatches, our causal models accurately describe structural relationships among human landscapes.

### Quantifying direct and indirect causal effects on diversity

Comparing the direct paths *B* → *π* and *µ* → *π*, we find that background selection is the primary driver of diversity along the human genome. However, the relative importance of BGS and mutation rate varies substantially by chromosome (Figure 4), ranging from equality in some to a ten-fold difference in others (Figure S42). Chromosomes 8 and 16 show particularly strong effects of mutation whereas chromosome 19 shows a negative *µ* → *π* coefficient (*CDS* +*phastCons* elements), a result that is biologically implausible and likely reflects estimation error. Otherwise, our results corroborate previous studies that underscored the importance of the mutation landscape to genetic variation (Harpak et al., 2016; Smith et al., 2018).

**Figure 4:**
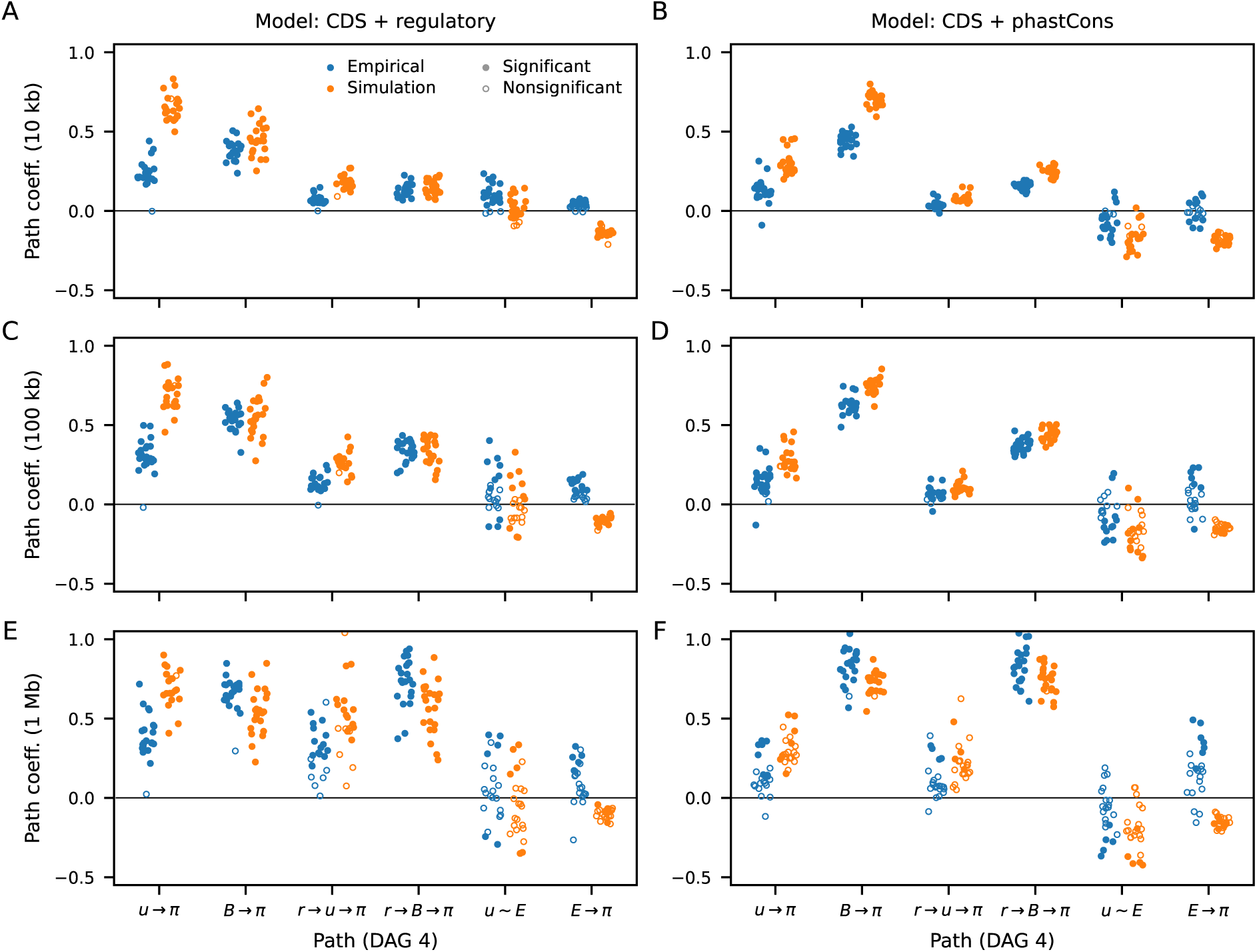
Estimated path coefficients across autosomes (DAG 4), divided by group of elements and genomic scales. *u* ∼ *E* stands for the free covariance between mutation rate and the density of constrained sites. Figures S41–S43 show estimated path coefficients computed separately within each chromosome.

Recombination affects *π* indirectly through both *r* → *B* → *π* or *r* → *µ* → *π*. Incorporating recombination into the causal models explains the positive covariance between *B* and *µ* (via the fork *µ* ← *r* → *B*), clarifying why the direct path coefficients *µ* → *π* are weaker than the correlations between mutation rate and diversity (Figures 4, S33–S37). We find that recombination affects *π* more strongly by alleviating background selection than by increasing mutation rates (Figures 4, S43). This happens despite its mutagenic effect being truly localized whereas its influence on *B*-values is long-ranged, and because linear effects, which are stronger along the BGS path, associate multiplicatively (*r* → *B* → *π* = *r* → *B* × *B* → *π* has a larger compound path coefficient than *r* → *µ* → *π* = *r* → *µ* × *µ* → *π*).

The positive *E* → *π* estimates alongside the negative *µ* → *π* in chromosome 19 are counterintuitive and suggest model mis-specification. To explore whether perfect knowledge of genomic maps and guaranteed adequacy of BGS would recover sensible estimates, we simulated mutational variance on top of E[*π*] and used it as observed diversity in SEMs (Methods). Candidate models fit better (as expected, Figure S46), and estimated path coefficients are biologically plausible (Figure 4): *E* → *π* is always negative whereas *µ* → *π* is always positive and even approaches *B* → *π* in magnitude. This consistency is also observed in individual-based forward simulations (Supplemental section 4). Therefore, the unusual estimates in human data probably derive from errors in the landscapes, from the influence of other types of selection, or both.

### Re-assessing the correlation between mutation rates and selected elements

Several models of evolutionary biology rest on the premise that *de novo* mutations occur independently from their fitness effects (Kimura, 1968; Bataillon and Bailey, 2014; Stoltzfus, 2021). Human somatic cells show evidence of reduced mutation rates in exons due to more efficient recruitment of mismatch repair machinery (Supek and Lehner, 2015; Frigola et al., 2017), although Rodriguez-Galindo et al. (2020) did not replicate this result in the germline. Conversely, whether mutation rates differ within and outside elements such as enhancers, promoters, and phastCons, has not been systematically tested. To quantify this relationship, we examined the covariance between *µ* and *E* within the SEM framework (Figure 2C-D). LRTs provide sparse support for DAG 4 at 1 Mb, but most chromosomes fit better under this model at 10 kb and 100 kb (Figures S31, S32, S47), particularly for *CDS* +*phastCons* where the estimated covariance with *µ* is negative (Figures 4, S48). This result mirrors simple correlations (Figures S33–S36), suggesting at face value that sites under long-term constraint also have lower mutation rates.

Yet the free covariance between *E* and *µ* in DAG 4 represents unmodeled causal structures that can take many forms. Resolving its nature will require explicit causal hypotheses and targeted data to discriminate among competing models. Without such rigorous treatment, models such as DAG 4 are provisional and we can only speculate about the origin of differential mutation rates in selected elements. In principle, despite their careful filtering of genetic variation, mutation maps could carry residual biases due to selection. Alternatively, the identification of phastCons from phylogenetic conservation may itself be affected by mutation rate variation. This could help explain why these sites are under weak selection in humans (Di et al., 2025, Table S4), why they have excelled at predicting local diversity (Murphy et al., 2022; Buffalo and Kern, 2024), and why the impact of mutation rate variation on *π* is much weaker in *CDS* +*phastCons* compared to *CDS* +*regulatory* models (Figures 4, S21, see also Supplemental section 5.6). Ongoing work is investigating these scenarios.

### Caveats and assumptions

A key assumption of studies of linked selection is that the available genomic maps accurately reflect the true landscapes of mutation, recombination and selective constraint, and that these have remained constant over recent evolutionary history. Although genomic maps are considerably advanced in humans, they still contain estimation noise and potential biases. These should reduce fit measures and 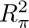, making our results (Figure 3) conservative rather than overstated. However, simulations indicate that map errors can also hinder the estimation of certain path coefficients. Addressing these limitations will require additional empirical data. Another challenge involves the distributional assumptions of SEM. At the 1 Mb scale, the smallest human chromosomes contain fewer windows (after filtering for coverage) than the recommended minimum of ∼ 5 − 10 data points per model parameter (Bentler and Chou, 1987). Even so, their estimated effects are broadly consistent with those at smaller scales and of larger chromosomes, so that small sample sizes should not strongly affect our conclusions. Similarly, the *B*-value of a focal site reflects the influence of several constrained elements whose impacts decay with recombination distance. Consequently, *B*-values are influenced not only by local *E*, *µ*, and *r*, but also by those of neighboring windows. SEM ignores such long-range dependencies, treating genomic windows as discrete, independent units. This incomplete representation is largely benign, as path coefficients going into *B* are not critical to our results (if anything, it makes the conclusion *r* → *B* → *π > r* → *µ* → *π* conservative). Nevertheless, the dependence of *B* on distant genomic regions can affect goodness-of-fit, contributing to rejection of causal models at scales *<* 1 Mb.

Finally, our findings are robust to empirical estimates of the DFE from the SFS (Supplemental section 4), and to the use of chromosome-specific *N_e_*, the assumption of demographic equilibrium, different thresholds for including noncoding phastCons, and the choice of recombination map (Supplemental sections 5.3–5.6).

## Conclusions

Here we used Sewall Wright’s method of path analysis (Wright, 1920, 1921, 1923) to study mechanisms shaping genetic diversity, a central topic in his own career (Wright, 1931). Although population genetics theory describes how micro-evolutionary processes jointly affect *π*, human genomic landscapes interact further. For example, recombination and mutation are causally connected since the repair of recombination-induced double-strand breaks increases mutation rates (Arbel-Eden and Simchen, 2019; Hinch et al., 2023). Causal inference clarifies the origins of observed correlations, teasing apart direct and indirect effects (Pearl, 2000; Pearl and Mackenzie, 2018; Shipley, 2016). We expect its application in population genetics to flourish in the coming years (see also Freudenberg et al., 2009; Weinstein and Roy, 2026).

Rather than accounting for all explainable variation in diversity, we found that *B*-values predicted on available annotations and DFEs do not correlate with *π* as strongly as previously suggested (Figures S21, S23). At the same time, the *B*-values we predicted here do not capture all signal of BGS in humans. A more comprehensive map of functional elements (including, for example, cryptic genes (Wright et al., 2022) and improved noncoding annotations (Benegas et al., 2025)) may strengthen the relationship between *B* and *π* even beyond previous estimates. Importantly, however, efforts to identify selection targets along the genome should be designed carefully to avoid absorbing confounding effects such as mutation rate variation.

We detected substantial impact of mutation rates on diversity in several autosomes. As this heterogeneity may affect evolutionary inferences, we advocate for incorporating mutation rate variation into the null model of human evolution. We further uncovered large variation in 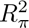, path coefficients and goodness-of-fit among chromosomes, raising questions about the quality of maps and the explanatory sufficiency of BGS in parts of the human genome. Simulations indicate that our proposed models are sound representations of causal relationships when complete information is available, motivating the search for biological factors that drive the discrepancies we identified in real data. For example, other types of selection (e.g., stabilizing selection on quantitative traits) may be particularly relevant in some regions of the genome. Therefore, DAGs 3 and 4 should be regarded as baseline models. Future extensions may include gene conversion (Pouyet et al., 2018) and efforts to resolve the covariance between mutation rates and density of constrained sites.

## Methods

### Predicting B-values and diversity in humans

We used moments++ to predict *B*-values in the weak and moderately deleterious regimes and the classic background selection module implemented in bgshr for strongly deleterious mutations (Barroso and Ragsdale, 2026). The two-locus system of moments++ is characterized by a deleterious mutation rate *µ*, a selection coefficient *s*, a recombination distance *r*, and a prescribed demographic model (or under equilibrium, a single *N_e_*). To make predictions for whole chromosomes, we created a lookup table containing expectations of *B* across a grid of *s* and *r* values at a unit *µ* (Supplemental section 1.1). We predicted *B* for focal sites positioned at regular 1 kb intervals along the chromosome by aggregating across their pairwise interactions with constrained elements, then obtained site *B* estimates using cubic splines interpolation between focal sites. We used mutation maps and estimates of direct selection to find expected diversity in the absence of background selection (*π*_0_), and predicted expected diversity per site as E[*π*] = *B* × E[*π*_0_]. We accounted for the effect of direct purifying selection on *π*_0_ within constrained elements by using the appropriate DFE to integrate across expected deleterious diversities calculated by moments++ for each *s*-value (Supplemental section 1.10). After filtering sequence data by combining the 1000 Genomes strict accessibility mask and the mutation map coverage, we further excluded sites within centromeres (Supplemental section 2).

To compute the pairwise *B*-value at a focal site linked to a single constrained element composed of multiple nucleotides, we first computed the recombination distance between the focal site and the center of the constrained element. To increase computational efficiency when modeling short, numerous elements (e.g., phastCons), we aggregated sites from potentially non-contiguous elements into regularly-spaced 1 kb windows. We calculated *B* given a unit mutation rate *µ* by interpolating between grid *r*-values for the required recombination distance. Taking the aggregate deleterious mutation rate of constrained sites in the window, *U*, we exponentiated *B* by *U/µ* to account for both element size and mutability; this is equivalent to taking the product of *B*-values associated with each constrained site in the window. We accounted for the distribution of selective constraint within each element by predicting *B*-values for each *s*-coefficient in the grid, then numerically integrating against a given DFE using weights assigned by discretizing the distribution. To find predicted *B* for a given focal site, we multiplied together the many pairwise *B*-values between the site and constrained windows along the chromosome. We generalized this approach to allow for multiple classes of constrained sites, each characterized by a distinct DFE (Supplemental section 1.2).

For each chromosome, we also predicted the relative reduction in diversity caused by strongly selected sites located in other chromosomes (*r* = 1*/*2, Charlesworth, 2012; Matheson and Masel, 2024). We incorporated such unlinked BGS effects both in the *N_e_*-fitting procedure (see below) and in our final model predictions.

#### Adjusting predictions due to selective interference

The procedure outlined above ignores the effects of background selection on constrained elements, or selective interference. To a first-order approximation, background selection modulates *N_e_*along the genome. To account for this effect, we performed several rounds of interference correction (Good et al., 2014; Barroso and Ragsdale, 2026), which is an iterative local rescaling of *s*, *r* and *U* by *B*-values estimated in the previous round of prediction to reflect the local reduction in *N_e_* (Supplemental section 1.9). We applied this correction to the entire recombination map by multiplying local recombination rates against local *B*. For each window harboring constrained sites, we rescaled *s* and *U* using window-specific average *B*-values. To account for interference from unlinked sites, we scaled local parameters by the unlinked *B*-value in the initial round of prediction and multiplied the unlinked effect against the *B*-map used for scaling in each subsequent iteration. As shown in Barroso and Ragsdale (2026), this procedure accurately recovers local *B*-values in equilibrium demographic scenarios.

#### Assigning DFEs

For all constrained elements, we based our predictions on SFS-based estimates of the DFE. Following the models presented in Di et al. (2025), we included the top 60% phastCons scores in putatively functional noncoding regions, computed in a 17-way primate phylogeny and divided into 12 classes by decreasing level of constraint. Our *regulatory* model included enhancers and promoters whose DFEs also follow Di et al. (2025). Exonic sites were either lumped under the Kim et al. (2017) DFE or split into deciles of constraint based on tolerance to loss-of-function mutations (Zeng et al., 2024). In the latter case, we fit SFS-based DFEs per decile using moments (Jouganous et al., 2017) (Supplemental section 1.3, Table S4).

#### Fitting the drift-effective population size

We fit the equilibrium drift-effective size (*N_e_*, Supplemental section 3), which moments++ uses to compute coalescence rates independently from background selection effects (Barroso and Ragsdale, 2026). Because this involves iteratively optimizing *N_e_*, and because BGS effects depend on compound parameters of *N_e_* with *s*, *r*, and *µ*, we rescaled the parameters of the lookup table (*s*, *r*, *µ*) by relative *N_e_* in each round (Supplemental section 1.5). We were therefore able to re-use the same lookup table for values of 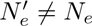 when fitting the drift-effective size. We used maximum likelihood to fit expected *π* landscapes to observations of *n_S_* and *n_D_*, the numbers of identical-by-state and different-by-state allele pairs, at the base pair resolution. Treating sites as independent, we applied a binomial likelihood function 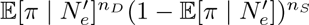, where 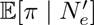 is the model prediction of site diversity, following Buffalo and Kern (2024). We used the fmin minimization function implemented in scipy (Nelder and Mead, 1965; Virtanen et al., 2020) to optimize the composite log-likelihood across accessible sites on a given chromosome (Supplemental section 2.2). We repeated this procedure for each chromosome, mutation map and constraint model.

### Structural equation models of genetic diversity

Structural equation models fit linked linear regressions by minimizing the distance between the observed covariance matrix and that implied by the model parameters. Since they assume independent and multivariate-normally distributed observations (genomic variables averaged within non-overlapping windows), we log-transformed and standardized landscapes before model-fitting. Standardization also renders variables unitless, allowing their relative contributions to be compared directly through the estimated path coefficients. We used latent representations of *B* and *µ*, and fit candidate DAGs as path models with lavaan (Rosseel, 2012). We filtered genomic windows based on their proportion of missing data (maximum of 20%, 50% and 75% at 10 kb, 100 kb and 1 Mb, respectively).

To account for spatial autocorrelation in 10 kb maps, we implemented a blockwise generalized least squares procedure before fitting SEMs. Due to coverage filtering (remaining windows are not necessarily adjacent in genomic coordinates), we first grouped windows into distance-aware blocks (maximum length of 500 kb). Within each block, we estimated pairwise covariances of residuals and modeled their decay as a function of genomic distance using an exponential autocorrelation function. The resulting covariance matrices were used to construct generalized least squares weights, and the adjusted covariance estimates were supplied for SEM.

#### Latent representations of genomic landscapes

Using lavaan (Rosseel, 2012), we fit chromosome-specific measurement models to synthesize the information from mutation maps (Supplemental section 8.1). We used the Roulette, gnomAD and Carlson maps as indicators (log-transformed and standardized), and predicted latent mutation maps that were used to fit candidate DAGs. For each chromosome, we generated twelve *B*-maps on a 3 × 4 grid of mutation maps and constrained elements. Within each group of elements, *B*-maps predicted under different mutation maps were effectively collinear (Pearson’s *r >* 0.99 for all pairs), leading measurement models into numerical instability. To obtain a latent estimate of *B*, we extracted the first principal component from the covariance matrix of the three *B*-maps. After computing PC loadings, we projected the standardized *B*-maps onto the vector to obtain a latent *B*-value for every genomic window. This procedure yielded synthetic *B*-values for each constraint model. Conversely, we based our predictions on the YRI-specific recombination map (Spence and Song, 2019) alone, so we used its estimates directly in the path analysis (see also Supplemental section 5.5).

#### Simulation study

To simulate genetic diversity under the assumption of error-free genomic landscapes, we added mutational variance on top of E[*π*] based on a sample size of 25 independent diploids. This layer of stochasticity scatters simulated *π* around *B* × E[*π*_0_], preventing numerical issues when fitting SEMs (Supplemental section 5.1). We used fwdpy11 (Thornton, 2014, 2019) to perform forward-in-time simulations incorporating population size changes, background selection as well as mutation and recombination rate variation (Supplemental section 4). From these, we evaluated the robustness of DFE fitting and downstream analyses, including *B*-value prediction and applications of SEM to simulated data.

## Supporting information

Visual landscapes and associative overdominance

## Data and software availability

The current source code for moments++ can be found at https://github.com/gvbarroso/momentspp. The current source code for bgshr can be found at https://github.com/apragsdale/bgshr. Data and scripts for reproducing figures and results, as well as fine-scale *B*-maps available as a resource, are provided at https://github.com/nwcol/bgs_lmr.

## Acknowledgments

We thank Armando Caballero and members of the Ragsdale lab for helpful discussions. Computational resources were generously provided through UW-Madison’s Center for High Throughput Computing. This work was supported by NIH Awards R35GM154962 to APR and R35GM119856 to KEL.

## Notes

### Competing Interest Statement

The authors have declared no competing interest.

### Summary of Updates

This version addresses comments from reviewers. It includes a substantial amount of new analyses, including validation with forward simulations, which can be found in the Supplement. It also improves the writing of the main text.

