## Supplemental material for "Causal inference clarifies the roles of background selection and mutation rate variation in shaping human genetic diversity"

September 9, 2026

**6 Evaluating the limits to $R^2_\pi$ imposed by genetic drift** **26**

**7 Predicting differences in $R^2_\pi$ among autosomes** **26**

**8 Tools and properties of genomic structural equation models** **27**

**9 Supplemental Tables** **34**

**10 Supplemental Figures** **37**

### 1 Modeling background selection and diversity landscapes

| Symbol | Meaning |
|---|---|
| $x$ | genomic position |
| $i$ | indexes DFE classes |
| $j$ | indexes constraint windows |
| $k$ | indexes $s$-grid |
| $s_k$ | element $k$ of the $s$-grid (sorted from $s_1 = 0$ to $s_n = 1$) |
| $w(s_k)$ | discretized DFE weight function |
| $b(s, u, r)$ | BGS effect for given parameters |
| $U_{i,j}$ | deleterious mutation rate of class $i$ in window $j$ |
| $\mu$ | mutation rate used to prepare lookup table |
| $u$ | site mutation rate |
| $r_{x,j}$ | map distance between site $x$ and window $j$ |

Table S1: A summary of notation used in this section.

#### 1.1 Overview

Our approach to predicting the $B$ landscape relied on a precomputed lookup table, which stored expected $B$-values across a discrete grid of deleterious selection coefficient ($s$) and recombination fraction ($r$) parameters

(Barroso and Ragsdale, 2026). We computed these tables using pure two-locus models (`moments++` in the domain where $2N_e s < 100$, and classic BGS (cBGS) theory where $2N_e s > 100$; we define the fitness of the heterozygote for the deleterious allele as $1-s$, so that $s \geq 0$), and extended them to our multi-locus framework by treating the effects of multiple constrained sites on a focal site multiplicatively. The lookup table used in this work contained 30 selection coefficients (log-spaced from $s = 0$ to 0.005) and 82 recombination fractions (log-spaced from $r = 0$ to 0.5). Our cBGS extension appended 20 further $s$-coefficients log-spaced between $s = 0.005$ and 1. We constructed the lookup table with a single deleterious mutation rate, $\mu = 10^{-8}$, and calculated reductions due to arbitrary mutation rates using a local rescaling.

For each chromosome, we predicted $B$ at focal sites regularly spaced at 1 kb intervals and used non-negative cubic splines interpolation to estimate $B$-values at intermediate sites. Because some of the constrained elements that we considered occur in very small fragments (e.g., phastCons sites), we grouped elements together in non-overlapping 1 kb windows. Where an element crossed the boundary between two or more windows, we broke it into multiple segments and assigned each to the appropriate window. For computational efficiency, we aggregated constrained elements into these 1 kb windows and computed recombination distances with respect to the midpoint of each window, since predicted $B$ is nearly invariant over small differences in recombination distances (with differences in $r \ll 1/N_e$) (Barroso and Ragsdale, 2026). The compound effect of hundreds of 1 kb windows on the $B$-value of a focal site overshadows the small imprecision in the recombination distance to individual constrained sites that is introduced by this simplification.

To predict the pairwise B-value at a focal site linked to one such window, we computed the recombination distance $r$ between the focal site and the center of the window containing constrained sites. To account for the number of constrained sites in a window and their mutation rates, we calculated $b(r, s, U)$ using the sum of deleterious mutation rates across the window, $U$, repeating for each $s$-coefficient. To integrate across the discrete $s$-grid of the lookup table, we discretized the element-specific gamma-distributed DFEs, using numerical integration to find a probability mass for each $s$ coefficient. We then took a product across table predictions for each element of the $s$-grid, weighting each by raising it to power of the appropriate probability mass. We generalized this approach to allow several classes of functional elements, each characterized by a set of genomic intervals specifying contiguous constrained sites and a DFE. To account for selective interference between constrained sites, we performed several rounds of iterative adjustments to local parameters $r$, $s$, $\mu$, using $B$-values obtained in the prior round of prediction as scaling factors (proxies of local $N_e$). To further reduce computational burden, we imposed an upper bound on interaction distance for each $s$ coefficient in the grid. For a given $s$, we considered effects at recombination distances where $1 - b < 10^{-10}$ to be negligible. For the strongest selection coefficients, this distance was $r = 0.5$ and no upper bound was applied.

#### 1.2 Obtaining and preparing data

##### 1.2.1 Constraint annotations

Throughout, we used genomic maps and annotations in genome build GRCh38. We represented regions under selective constraint using BED files, which we obtained from sources listed in this section. When combining annotations from independent sources, a small number of overlaps, where sites were assigned to more than one annotation, occurred. We removed any overlaps at computation runtime, prioritizing DFE classes manually so that DFEs with the highest mean $s$ had higher priority.

**CDS elements.** We downloaded exon data from the consensus coding sequence (CCDS) category in Ensembl release 114 (`https://ftp.ensembl.org/pub/release-114/`). From these, we removed sites which were not translated in any transcript, leaving only protein-coding sequence (comprising 33 Mb, 189,000 elements). We assigned coding exons to genes using Ensembl gene identifiers and partitioned them into deciles of constraint according to gene $sh$ scores from Zeng et al. (2024). We lumped genes not present in the data of Zeng et al. (2024) into a further eleventh category.

**Regulatory elements.** We obtained annotations for promoters (29 Mb, 78,000 elements) and enhancers (239 Mb, 610,000 elements) after Di et al. (2025), who assembled these elements by combining promoter and enhancer-associated chromatin states inferred by Vu and Ernst (2022) using chromHMM (Ernst and Kellis, 2012). Promoter elements comprised chromatin states PromF1–PromF7 (flanking promoters) and enhancers comprised states EnhA1–EnhA20 (active enhancers) from Vu and Ernst (2022). For each annotation, only sites which passed the 1000 Genomes Project strict mask were retained.

**Non-coding phastCons elements.** We obtained phastCons annotations after Di et al. (2025), who divided genome-wide phastCons conservation scores (17-way scores among primates, Siepel et al., 2005) into bins representing $\sim$5% quantiles. They then intersected each bin with the set of candidate functional noncoding sites, defined as noncoding sites that were not classified as heterochromatin or quiescent chromatin states (regions with weak epigenetic marking) by Vu and Ernst (2022), and removed sites which did not pass the 1000 Genomes Project strict mask. These elements include the chromatin classes grouped into promoter and enhancer categories, as well as other regulatory elements and transcribed non-coding regions (introns, untranslated regions). In our analyses, we used the top 12 candidate functional noncoding phastCons bins (756 Mb, 237 million elements), which contain the $\sim$60% highest phastCons scores among candidate functional noncoding sites and account for $\sim$88% of their total inferred deleterious mutation rate (Di et al., 2025).

##### 1.2.2 Mutation maps

We obtained site-resolution Carlson (Carlson et al., 2018), gnomAD (Karczewski et al., 2020) and Roulette (Seplyarskiy et al., 2023) mutation rates in `.vcf` format from `https://genetics.bwh.harvard.edu/downloads/Vova/Roulette/`. These files contained mutation rates from the Hg38 reference nucleotide to the three possible alternate nucleotides. To calculate a site-resolution average rate, we took the sum of rates at each site and applied the scaling factors provided at `https://github.com/vseplyarskiy/Roulette`.

We incorporated the specific coverage of each mutation map into the genetic masks used to filter sequence data. We also used these genetic masks to determine the sites at which to predict expected diversity, so that $\mathbb{E}[\pi]$ at each site was calculated with a site-specific mutation rate estimate. Consequently, the set of sites for which we computed $\mathbb{E}[\pi]$ corresponded exactly to the sites at which we measured observed diversity.

When tabulating the deleterious mutation rates of constrained sites to predict $B$, we never masked mutation rates. In cases where mutation data for an element was missing, we imputed the rate using the following procedure: if an entire contiguous element lacked mutation data, we assigned its sites the chromosome-wide average mutation rate. If only some sites in a contiguous element were missing data, we assumed that their mutation rate was the average across non-missing sites in the same element. This step preceded the aggregation of contiguous elements into 1 kb windows.

##### 1.2.3 Recombination maps

We downloaded human recombination maps inferred by Spence and Song (2019) with `pyrho` (Kamm et al., 2016) from `https://zenodo.org/records/11437540`. We used the maps inferred for the YRI population. These maps assign piecewise-constant recombination rates to genomic intervals with irregular lengths. To calculate the map positions of arbitrary sites, we used linear interpolation between cumulative map distances at the breakpoints between piecewise-constant segments.

We downloaded the deCODE recombination map (Halldorsson et al., 2019) from `https://www.science.org/doi/full/10.1126/science.aau1043`, and used it to fit alternative models (Section 5.5).

##### 1.2.4 Obtaining regularized genomic maps

Prior to fitting measurement and path models with `lavaan`, we regularized all genomic landscapes so that they are presented in non-overlapping intervals of equal length. We first computed interval averages of genetic diversity (both observed and expected), recombination, mutation, $B$-values and density of constrained sites within 1 kb windows. We then further binned these windows to obtain maps at 10 kb, 100 kb and 1 Mb scales by computing averages weighted by 1 kb coverage. For both estimating correlations and fitting SEM, the density of constrained sites, $E$, represents the sum of constrained sites across all elements in a given model (*CDS+phastCons* or *CDS+regulatory*), divided by the number of callable sites. As such, its estimated effects ignore differences in the (average) strength of negative selection among deleterious sites in different windows.

#### 1.3 Choice of DFE parameters

Throughout, we used gamma or gamma-neutral (a gamma distribution with a point-mass of neutrality) DFEs, with parameters independently inferred using the SFS. To control for population size history and background selection, each DFE was fit conditional on a demography inferred with putatively neutral sites located near selectively constrained loci. We converted inferred scale parameters, which are typically presented in units scaled by the ancestral size ($2N_A$), to physical units by dividing by the appropriate $2N_\mathrm{A}$ value. The full array of DFE parameters is presented in Table S4.

**Merged CDS elements.** For coding exons, we used the DFE inferred for nonsynonymous sites by Kim et al. (2017) using Fit*dadi* (Kim et al., 2017; Gutenkunst et al., 2009) on the LuCamp dataset (Lohmueller et al., 2013). To account for the synonymous fraction of mutations in CDS elements, we used the parameters of the pure gamma DFE from Kim et al. (2017) to construct a gamma-neutral DFE with a neutral probability mass of $p_\mathrm{neu} = L_\mathrm{S}/(L_\mathrm{NS} + L_\mathrm{S})$, where $L_\mathrm{NS}$ is the nonsynonymous sequence length and $L_\mathrm{S}$ the synonymous length. Using $L_\mathrm{NS} = 2.31 \times L_\mathrm{S}$ (Huber et al., 2017) yielded $p_\mathrm{neu} \approx 0.302$.

**Fitting DFEs across deciles of CDS constraint** Using Ensembl gene identifiers, we assigned CDS exons to *sh* scores available from Zeng et al. (2024) and divided them into deciles of decreasing levels of constraint. We filtered out exonic sites that were not amino-acid coding (untranslated regions) in any available transcript, and computed synonymous and nonsynonymous mutation rates per gene according to the Roulette mutation map (Seplyarskiy et al., 2023). We fit two-epoch demographic models on the synonymous SFS of each decile to control for population size history and background selection, then fit gamma-distributed DFEs to all nonsynonymous mutations independently for each decile.

**Regulatory and noncoding phastCons elements.** DFE estimates for promoters, enhancers, and binned phastCons annotations were made in Di et al. (2025) using Fit*dadi* (Kim et al., 2017). Briefly, in that work the SFS was generated from the 108 1000 Genomes Project YRI genomes (Byrska-Bishop et al., 2022). To control for demography and the effects of linked selection, parameters were fit conditional on three-epoch demographic models inferred with variation at nearby putatively nonfunctional sites; these were sites annotated with the quiescent chromatin state by Vu and Ernst (2022) that fell within 100 kb of two or more constrained elements. Local mutation rate variation was accounted for by scaling the putatively neutral SFS by the ratio of the total constrained mutation rate to the total putatively neutral rate, computed with the Roulette mutation map (Seplyarskiy et al., 2023).

#### 1.4 Discretizing the DFE

We discretized DFEs (either gamma or gamma-neutral) in order to integrate across $B$-values predicted on the discrete lookup table $s$-grid. The DFE mass corresponding to a particular $s$ coefficient is the probability that a mutation has a selective effect closer to $s$ than to any other value in the grid. Gamma-neutral DFEs place a discrete mass representing the probability that a mutation is selectively neutral, $p_\mathrm{neu}$, on $s = 0$. Let $k$ index the $s$-grid, which has length $n$ and is sorted so that its first element is $s_1 = 0$ and the last is $s_n = 1$, and let $F$ be the cumulative distribution function (CDF) of the desired pure gamma distribution. Also let $z_k = (s_k + s_{k+1})/2$ be the midpoint between $s_k$ and $s_{k+1}$, defined for $1 \leq k < n$; then $z_{k-1}$ and $z_k$ delimit a bin about $s_k$. For a gamma distribution without neutral mass, the discrete weights $w(s_k)$ are:

$$w(s_k) = \begin{cases} 0, & k = 1 \\ F(z_k), & k = 2 \\ F(z_k) - F(z_{k-1}), & 2 < k < n \\ 1 - F(z_{k-1}), & k = n. \end{cases} \tag{S1}$$

Note that $s_1 = 0$ receives no probability mass, while the smallest nonzero grid point $s_2$ receives all the density in $0 < s < z_2$. The gamma-neutral discretization is:

$$w(s_k) = \begin{cases} p_{\text{neu}}, & k = 1 \\ (1 - p_{\text{neu}})F(z_k), & k = 2 \\ (1 - p_{\text{neu}})(F(z_k) - F(z_{k-1})), & 2 < k < n \\ (1 - p_{\text{neu}})(1 - F(z_{k-1})), & k = n. \end{cases} \tag{S2}$$

**Integrating across the DFE** To integrate across the DFE, we took a weighted product of $B$-values corresponding to each element of the $s$-grid. This is equivalent to multiplicatively combining effects from the proportion of mutations that belong to each $s$ bin, so that

$$B = \prod_k^n b(s_k)^{w(s_k)}. \tag{S3}$$

Suppose that $b(s_k) = \exp(-h(s_k))$ for some function $h$. Then, our integration scheme is equivalent to taking a weighted sum of $h(s_k)$ within the exponent,

$$B = \exp\left(-\sum_k^n w(s_k)h(s_k)\right). \tag{S4}$$

In practice, we summed over bins $k > 1$ corresponding to $s > 0$, as mutations with $s = 0$ make no contribution to background selection.

#### 1.5 Scaling lookup tables to the drift-effective population size

To both predict $B$-values and estimate effective population sizes, we assembled an equilibrium lookup table using a drift-effective size ($N_e^{\text{ref}}$) of $10^4$. This value is lower than the the drift-effective size for which we typically aimed to make predictions (Figure S57). To adjust the $B$-value predictions in the lookup table to correspond to an arbitrary drift-effective size $N_e^{\text{target}}$, we scaled table $s$, $r$, and $\mu$ values by the factor $c = N_e^{\text{ref}}/N_e^{\text{target}}$. The rationale is that at demographic equilibrium, $B$ dynamics are determined by the population-scaled parameters $\theta = 4N_e\mu$, $\rho = 4N_e r$, $\alpha = 2N_e s$. We transformed physical rates into population-scaled ones according to the reference size, $\theta = 4N_e^{\text{ref}}\mu$, then back into physical units corresponding to the target size with $\mu' = \theta/4N_e^{\text{target}} = c\mu$. Hence, by multiplying physical parameters by $c$ while holding the lookup table predictions of $B$, $\pi_{0,L}$ (diversity at constrained loci), and $\pi_{0,R}$ (diversity at neutral loci) constant, we can transform a prediction from $b(s, r, u, N_e^{\text{ref}})$ to $b(cs, cr, cu, N_e^{\text{target}})$. After scaling, we used the classic background selection module in `bgshr` (Barroso and Ragsdale, 2026) (which is independent of $N_e$) to extend the lookup table selection coefficient grid up to $s = 1$.

**Transforming recombination fractions to map units.** For computational efficiency, we transformed the lookup table recombination parameter grid from $r$ to Morgans using Haldane's inverse mapping function, $d = -\log(1 - 2r)/2$. This makes calculating recombination distances faster – we only need to compute the map distance between two sites using a genetic map expressed in Morgans, rather than applying Haldane's map function to each map distance considered during multi-locus prediction. Because $r = 1/2$ corresponds to an infinite map distance, we inserted several large map distances to the $M$-grid obtained by transforming $r$ (e.g., $M = 10$) to prevent interpolation from extending beyond the grid. When $c < 1$, which is the case since $N_{\text{ref}} = 10^4$ and $N_{\text{target}} \approx 3\times10^4$, this procedure yielded $B$-value predictions in the weak and moderate selection regimes that did not extend to the maximum recombination fraction of 0.5. This gap is inconsequential, however, as diversity reduction is negligible for these selection coefficients at large recombination distances.

#### 1.6 Extending prediction with classic BGS theory

Charlesworth (2012) derived an expression for cBGS under arbitrary recombination distances – we specifically used the expression $B \approx \prod_i \{(1 - q_i(\delta_{c_i}^2 + \delta_{r_i}^2)\}$ from the Appendix of that work, without making the assumption that $s$ or $r$ is small. Substituting in the variables defined by Charlesworth (2012) yields

$$b(s, r', q) = \exp\left(-\frac{qs^2(1 + [2r' + s]^2)}{(r' + s)^2}\right). \tag{S5}$$

Here $q$ is the equilibrium frequency of the deleterious allele and $r' \approx r(1 - s)$ is the effective rate of recombination, which reflects the loss of heterozygous carriers of the deleterious mutation due to selection. Substituting $q \approx \mu/s$ and $r' \approx r(1 - s)$, we obtain

$$b(s, r, \mu) = \exp\left(-\frac{s\mu(1 + [2r(1 - s) + s]^2)}{(r[1 - s] + s)^2}\right). \tag{S6}$$

Since cBGS predictions are invariant to $N_e$, we do not apply $N_e$ scaling to predictions where $2N_e s > 100$.

#### 1.7 Modeling unlinked background selection

Substituting $r = 1/2$ into Equation S6, we can find a simpler expression for unlinked BGS. Let $U = \bar{\mu} \times L$ be a total deleterious mutation rate. Then if every mutation has selection coefficient $s$,

$$b_{\text{unlinked}}(s, U) = \exp\left(-\frac{8sU}{(1 + s)^2}\right). \tag{S7}$$

We precomputed unlinked effects for each combination of chromosome and model, and used the tabulated reductions when predicting $B$ landscapes. To compute the effect on chromosome $a$, we took the product of DFE-weighted effects from each other chromosome. Suppose that $U_{c,i}$ is the total mutation rate of elements in DFE class $i$ on chromosome $c$, then

$$B_{\text{unlinked}}(a) = \prod_{c \neq a} \prod_i \prod_k^{n-1} b_{\text{unlinked}}(s_k, U_{c,i})^{w_i(s_k)}. \tag{S8}$$

Because effects from weakly selected mutations are are expected to be negligible, we ignored unlinked BGS effects for $s$ which were not calculated using cBGS.

#### 1.8 Multi-locus prediction

For a focal site $x$, $B(x)$ is the product of linked and unlinked diversity reductions. Then, since $B_{\text{unlinked}}$ is equal for all sites on a given chromosome,

$$B(x) = B_{\text{unlinked}} \times B_{\text{linked}}(x) \tag{S9}$$

To compute $B(x)$ due to linked sites associated with DFE $i$, we take a product of background selection effects exerted by each constraint window, indexed by $j$. The total linked $B$-value is then the product of DFE-specific reductions, each integrated across its discretized weight function $w_i$. Let $U_{i,j}$ be the deleterious mutation rate of element type $i$ in window $j$, and $r_{x,j}$ be a precomputed genetic map distance between focal site $x$ and the center of constraint window $j$. Then

$$B_{\text{linked}}(x) = \prod_i \prod_k^{n-1} \left[ \prod_j b(s_k, r_{x,j}, U_{i,j}) \right]^{w_i(s_k)}. \tag{S10}$$

We employed linear interpolation across the $r$-grid to find $b$ as a function of $r_{x,j}$. To find the effect for mutation rate $U$, we use

$$b(s_k, r_{x,j}, U) = b(s_k, r_{x,j}, \mu)^{U/\mu}, \tag{S11}$$

where $\mu = 10^{-8}$ is the mutation rate present in the lookup table. Assuming that $U = L \times \mu$, this is analogous to modeling the effects of $L$ sites with mutation rate $\mu$ using $b(\mu)^L$.

#### 1.9 Interference correction

For each round of interference correction, we used parameters which were locally rescaled by the $B$-map predicted in the previous round, $s'$, $r'$, and $U'$, to calculate a new map $B'$, so that

$$B'_{\text{linked}}(x) = \prod_i \prod_k^{n-1} \left[ \prod_j b(s'_k, r'_{x,j}, U'_{i,j}) \right]^{w_i(s_k)}. \tag{S12}$$

Using cubic interpolation, we estimated $B$ at every position along the genome using our grid of focal predictions. Define a function $\bar{B}(x_0, x_1)$ that averages the predicted $B$-map over a half-open interval $[x_0, x_1)$, so that

$$\bar{B}(x_0, x_1) = \frac{1}{x_1 - x_0} \int_{x_0}^{x_1} B(x)dx. \tag{S13}$$

The application of $\bar{B}(x_0, x_1)$ to rescale local parameters is shown below. For human-like parameters, convergence is generally observed in $< 10$ iterations (Barroso and Ragsdale, 2026); we used 10 iterations in this work, except when fitting $N_e$, where we use 5 due to computational constraints. To model interference from unlinked sites, we performed the initial round of prediction using an interference correction with a uniform $B$-map equal to $B_{\text{unlinked}}$.

**Scaling recombination maps** To model the effects of interference on recombination, we broke the recombination map into segments with endpoints determined by (1) the piecewise-constant-rate regions of the initial recombination map, and (2) the positions at which $B$ was predicted in the last round. Let $R(x)$ specify the map position of site $x$ in Morgans, and let $x_0$ and $x_1$ be the start and end position of a given segment. The average recombination rate of the segment is

$$\bar{r} = \frac{R(x_1) - R(x_0)}{x_1 - x_0}. \tag{S14}$$

We assigned the scaled rate across the segment to be $r' = \bar{r}\bar{B}(x_1, x_0)$. We then assembled the scaled map out of all such piecewise-constant rescaled segments and used it to compute distances $r'_{x,j}$.

**Scaling mutation rates** We scaled mutation maps by computing average predicted $B$ across each 1 kb window and multiplying it by the unscaled rate. Let $x_j$ be the start position of the $j$th window. Then

$$U'_{i,j} = U_{i,j}\bar{B}(x_j, x_{j+1}). \tag{S15}$$

**Scaling selection coefficients** To scale selection coefficients associated with a constraint window $j$, we left the DFE discretization unaltered, but adjusted the value of each grid point with

$$s'_k = s_k \bar{B}(x_j, x_{j+1}). \tag{S16}$$

We used linear interpolation across the $s$-grid in the lookup table to calculate the corresponding effect $b(s'_k)$.

##### 1.10 Computing expected diversity

Expected diversity in the absence of linked selection ($\mathbb{E}[\pi_0]$) for neutral sites is readily available in the lookup table (neutral $\mathbb{E}[\pi_0] = 4N_e\mu$ for demographies with equilibrium size $N_e$). We scaled expected neutral diversity to the site mutation rate $u$ using

$$\mathbb{E}[\pi_0] = \frac{u}{\mu}\mathbb{E}[\pi_0 \mid \mu]. \tag{S17}$$

Conversely, we calculated expected deleterious $\pi_0$ by integrating across $E[\pi_0|s_k, \mu]$ presented in the lookup table with element-specific DFEs and scaling by the local mutation rate, so that

$$\mathbb{E}[\pi_0] = \frac{u}{\mu}\sum_{k=1}^{n} w(s_k)E[\pi_0|s_k, \mu]. \tag{S18}$$

Then, $\mathbb{E}[\pi] = B \times \mathbb{E}[\pi_0]$. This procedure applies to both equilibrium and non-equilibrium models, since the lookup table contains expected neutral and deleterious $\pi_0$ under the prescribed demography. For a given mutation model, we computed $\mathbb{E}[\pi]$ only at sites that passed the 1000 Genomes strict callability mask (Consortium et al., 2015) and where a mutation rate estimate was available. The set of sites where we calculated expected diversity was therefore the same as the set of sites where we estimated $\hat{\pi}$ (see Section 2).

To visualize $\mathbb{E}[\pi]$ in regions putatively experiencing associative overdominance (Supplemental File), we downloaded the candidate 2 Mb windows identified by Gilbert et al. (2020) and lifted them over to GRCh38 using the `R` package `rtracklayer` (Lawrence et al., 2009).

#### 2 Obtaining variation data

##### 2.1 Sequence data

We obtained sequence variation data from 1000 Genomes samples resequenced to 30x coverage and aligned to the GRCh38 reference genome by the New York Genome Center Byrska-Bishop et al. (2022). To estimate base pair-resolution $\hat{\pi}$, we downloaded these samples in gVCF format (which contain records for all sites) and estimated diversity using methods after Buffalo and Kern (2024).

##### 2.2 Filtering accessible sites

When fitting models and computing average $\hat{\pi}$ across genomic windows, we masked out sites which were filtered by the 1000 Genomes strict callability mask or which lacked mutation rate data. Because we filtered out sites which lacked mutation data, masks were mutation-model specific (although the Roulette and gnomAD mutation maps have equivalent genomic coverage). We further masked out sites within annotated centromeres (labeled `acen` in the chromosome banding table `cytoBand.txt` downloaded from `http://hgdownload.cse.ucsc.edu/goldenPath/hg38/database/`; cf. Buffalo and Kern 2024, who applied a 5 Mb padding to each side of the centromere) and the neighborhood of the HLA complex on chromosome 6, as determined with the UCSC Genome Browser (positions `28500000` – `33500000`; see Figure S1 for a view of this region without filtering). Because we modeled direct purifying selection when computing $\mathbb{E}[\pi]$, we did not filter sites by putative selective constraint.

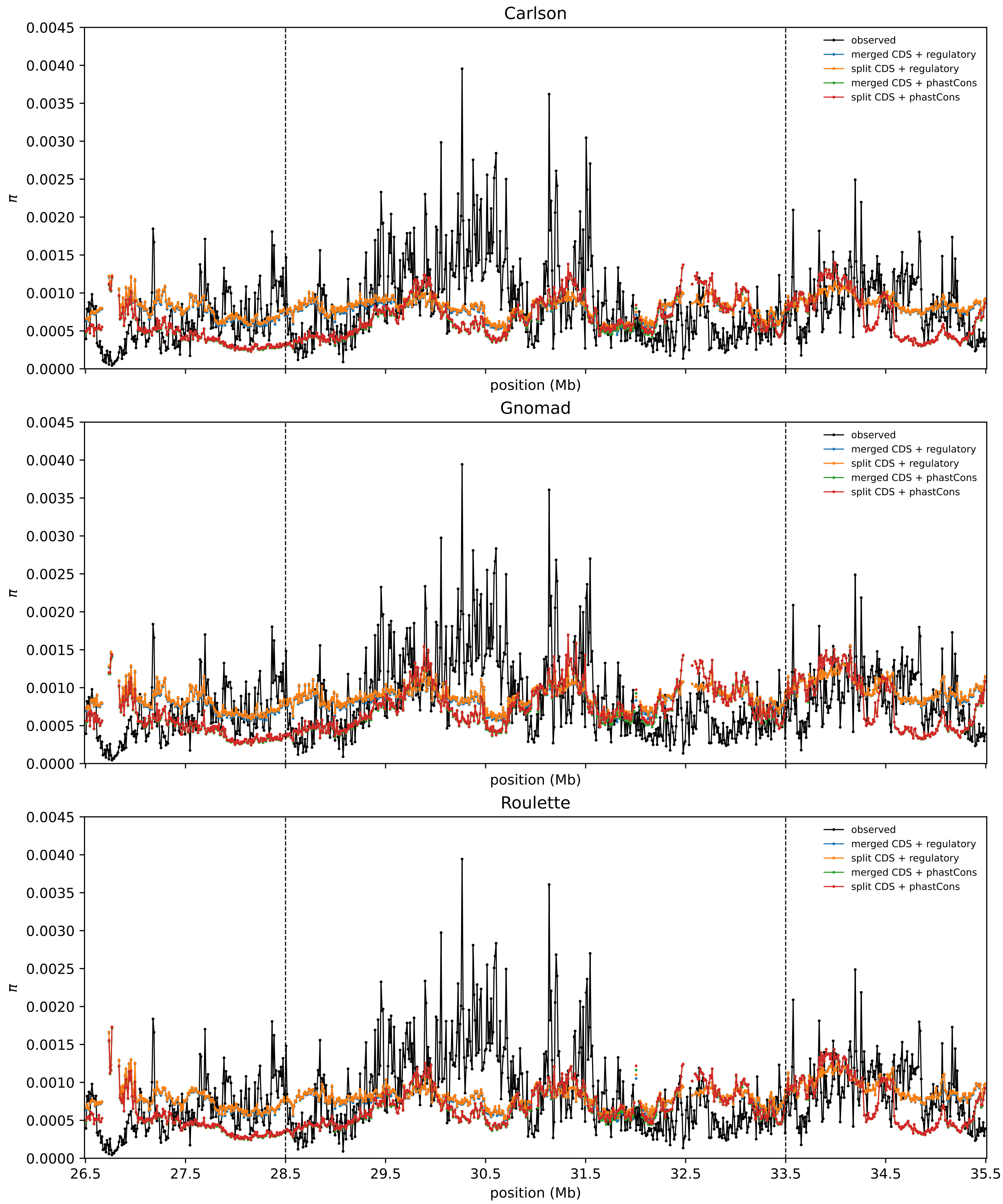

Figure S1: Observed and expected $\pi$ in the vicinity of the HLA (Human Leukocyte Antigen) locus on chromosome 6, at 10kb window size. Broken vertical lines indicate the edges of the HLA region masked in analyses. All windows with $\geq 1$ accessible sites are plotted.

#### 2.3 Estimating observed diversity

**Sample-level filtering** We extracted an array holding allele counts (reference/alternate) at each site from sample-specific gVCF files. We counted alleles only at sites with quality scores `GQ` $\geq$ `30`, `QUAL` $\geq$ `50`. Where samples did not meet these criteria, we left 0 entries in sample-level allele count matrices; hence, the number of samples counted varied from site to site. By summing together sample-specific allele count matrices, we found the total numbers of filter-passing reference ($n_R$) and alternate ($n_A$) alleles at each site. At sites with more than one alternate allele, we summed the counts of all alternate alleles to get $n_A$.

**Estimators of diversity** The total number of unique allele copy pairs that can be drawn from $n = n_A + n_R$ allele copies is $n_T = \binom{n}{2}$. From the counts of reference and alternate alleles at a site, we computed the number of distinct-by-state or heterozygous allele copy pairs as

$$n_D = n_R \times n_A. \tag{S19}$$

The number of identical-by-state or homozygous allele copy pairs is $n_S = n_T - n_D = \binom{n_R}{2} + \binom{n_A}{2}$. We estimated pairwise nucleotide diversity at a site with

$$\hat{\pi} = \frac{n_D}{n_T}. \tag{S20}$$

Which is equivalent to the typical measure of diversity;

$$\hat{\pi} = \frac{2n_D}{n(n-1)} \tag{S21}$$

#### 2.4 Computing windowed summaries

When averaging across sites to compute windowed summaries (Section 1.2.4), we weighted all sites equally, counting only sites which passed the appropriate genetic mask (see Section 2.2). This was in contrast to Buffalo and Kern (2024), who weighted site $\hat{\pi}$ by the number of allele copy pairs observed at each site.

### 3 Estimating $N_e$ across chromosomes and models

We estimated the drift-effective $N_e$ (in the absence of direct or linked selection) by minimizing the distance between the observed landscape of diversity and that predicted by our model. At a site, the likelihood of a given $N_e$ is the binomial likelihood of observing $n_D$ heterozygous sample pairs when the probability of sampling such a pair is $\mathbb{E}[\pi \mid N_e]$.

$$\mathcal{L}(N_e) \propto \mathbb{E}[\pi \mid N_e]^{n_D}(1 - \mathbb{E}[\pi \mid N_e])^{n_S} \tag{S22}$$

We drop the constant factor $\binom{n_T}{n_D}$. Here, $N_e$ influences $\mathbb{E}[\pi \mid N_e]$ both directly via $\mathbb{E}[\pi_0]$ and indirectly through its effects on $B$-values via `moments++`'s prediction.

Estimating $N_e$ using aggregated genome-wide diversity data is computationally unfeasible. We therefore estimated $N_e$ per chromosome by maximizing the composite log-likelihood across sites, yielding 22 $N_e$ estimates per model set. Let $x$ index sites that pass the mask on some chromosome; then

$$\ell(N_e) = \sum_x \left[n_{D,x} \log(\mathbb{E}[\pi_x \mid N_e]) + n_{S,x} \log(1 - \mathbb{E}[\pi_x \mid N_e])\right]. \tag{S23}$$

We optimized this function using the `fmin` function from SciPy, using $N_e$ = 30,000 as an initial guess. To speed up the inference procedure, we ran fits with relatively loose convergence criteria and computed focal $B$-values on a 10kb, rather than 1kb, grid. For each chromosome, we also incorporated the effect of unlinked BGS arising from strongly deleterious sites in other autosomes (Table S6). $N_e$ estimates were largely consistent across chromosomes (see below), and the modest amount of heterogeneity in inferred $N_e$ across chromosomes likely represents estimation noise and varying degrees of model mis-specification.

Our approach has the advantage of being model-specific as we fit $N_e$ individually for each combination of constrained element and mutation map. By construction, the $N_e$-fitting procedure aligns the *scale* of the predicted and observed landscapes of diversity (Supplemental File) but does not strongly influence the broad correlation patterns at window sizes $\geq$ 10 kb that are used in downstream analyses.

Figure S57 shows that depending on the constrained elements considered, estimated $N_e$ ranges across chromosomes from 22,546 to 36,398 (Roulette), from 22,500 to 38,273 (gnomAD) and from 17,812 to 30,284 (Carlson, where the average estimated mutation rate is higher, hence a smaller $N_e$ is needed to explain levels of genetic diversity). Note that because these values represent estimates of the drift-effective population size, they are substantially higher than the typical $N_e$ estimated from genome-wide average diversity in humans ($N_e = \hat{\theta}/4\mu \approx 15{,}000$), which is depressed by linked selection. In principle, our fitted $N_e$ should be closer to $N_e$ obtained from genetic diversity when $\theta = 4N_e\mu$ is estimated exclusively from sites where linked BGS is negligible ($B \approx 1$), although unlinked BGS also reduces diversity at these sites (Table S6).

### 4 Evaluating SEM with forward-in-time simulations

Fitting structural equation models (SEM) to human genomic landscapes involves important challenges. First, available maps of recombination, mutation, and $B$-values—as well as the genomic positions of functionally constrained elements—are estimated with some degree of error. Even if genome annotations were perfect, DFE inference from the SFS may introduce inaccuracies that propagate into $B$-value predictions. These errors can plausibly distort the estimation of path coefficients by SEM and worsen fit measures of candidate DAGs. Second, although our models are grounded in genome biology, they do not capture every process shaping human genetic diversity. Models that fit human data reasonably well do not rule out alternative causal structures that may also be compatible–or even yield better fits. Finally, binned genomic maps are autocorrelated, especially at 10 kb, introducing non-independence between the data points (genomic windows) used to fit SEM. Similar considerations prevented Weinstein and Roy (2026) from using SEM, as it is difficult to account for correlation patterns induced by phylogenetic relationships.

Collectively, these challenges may complicate the interpretation of SEM applied to human genomic landscapes, as poor fits or unrealistic estimates can arise from any combination of these factors. To address some of these concerns, we evaluated our SEM framework under idealized conditions without estimation errors and where only drift, mutation and (background) selection affect diversity. The goal of these simulations is to assess whether SEM is able to recover known population-genetic relationships in the absence of confounding effects. We conducted two sets of simulations using `fwdpy11` (Thornton, 2014) in which the population size history was piecewise constant, with more epochs than are typically assumed in DFE inference, so that fitting a two- or three-epoch model introduces some model mis-specification at that stage.

1. **Simulation set A:**
   - 100 independent chromosomes, 10 Mb each (1 Gb in total length), without scaling simulated parameters.
   - Baseline mutation rates were drawn in 1 Mb segments, uniform in (0, 2e-8), and then rates were drawn within 1 kb windows with mean equal to the baseline rate and standard deviation 2e-9. This created fine-scale variation and large-scale autocorrelation.
   - Baseline recombination rates were drawn independently of mutation rates, following the same parameters.
   - Two gamma-distributed DFEs, one stronger (with mean $s = 0.01$, shape parameter = 0.2) and one weaker (mean $s = 0.002$, shape = 0.1). 25% of all 1 kb regions were drawn randomly without replacement and assigned one of the two DFEs, with 25% chance the stronger DFE and 75% chance the weaker DFE. Within selected regions, 70% of new mutations had selection coefficients drawn from the DFE, while 30% were neutral.

2. **Simulation set B:**

   - 10 independent chromosomes, 100 Mb each (1 Gb total)
   - Due to the prohibitive computational burden of simulating large chromsomes without scaling, we applied a 5x down-scaling (in which $N$ was reduced 5-fold, $u$, $s$, and $r$ increased 5-fold)
   - Mutation and recombination rates were drawn as in set A, but with different parameters: base mutation rates were drawn from (1e-8, 1.5e-8), with standard deviation of 5e-9 within 1 kb windows. The Base mutation rates were drawn from (0, 2e-8), with a 1kb standard deviation of 1e-8.
   - Selection followed the same procedure as above.

Specifically, our analyses of these simulated data address: 1) the accuracy of $B$-value predictions when the DFE is inferred from the SFS (as opposed to predicting $B$ with the true DFE parameters); 2) whether equilibrium $B$-values are a reasonable approximation when population sizes fluctuate moderately; 3) whether DAGs 3-4 fit the landscapes well and are therefore adequate representations of causal effects in standard population-genetic scenarios; 4) whether SEM has enough power to distinguish between slightly different models (when we know which one is correct). We fit SEM with the same settings as we used for the analyses of human data, including correction for autocorrelation at 10 kb (see main text Methods and Section 8).

#### 4.1 $B$-value prediction is robust to empirical estimates of the DFE

At the end of the forward-time simulations, we sampled 100 diploid individuals and aggregated the observed site frequency spectra in neutral and selected regions, partitioning the SFS in selected regions between neutral (mimicking synonymous) and selected (mimicking nonsynonymous) mutations. We followed the procedure in which a demographic model is first fit to putatively neutral mutations, and then a DFE is fit to the selected alleles while fixing the inferred demography (Figure S2). Overall, this procedure has been shown to work well in practice (with simulated data), and results are consistent across independent datasets when applied to humans (e.g., Kim et al., 2017; Huber et al., 2017; Ragsdale, 2022). Here, we similarly find accurate recovery of the DFE, except when a steady-state demographic model is assumed (Table S2).

| | Shape | Scale | Mean |
|---|---|---|---|
| Truth (weak DFE) | 0.1 | 0.02 | 0.002 |
| Steady state | 0.367 | 0.00085 | 0.00031 |
| Two epoch | 0.103 | 0.0195 | 0.00201 |
| Three epoch | 0.104 | 0.0190 | 0.00197 |
| Truth (strong DFE) | 0.2 | 0.05 | 0.01 |
| Steady state | 0.436 | 0.00345 | 0.00151 |
| Two epoch | 0.204 | 0.0459 | 0.00937 |
| Three epoch | 0.204 | 0.0461 | 0.00941 |

Table S2: Re-inferred DFEs from simulations (simulation set B), using the SFS with 100 diploid samples. The DFE is poorly estimated when assuming steady-state demography, but both a two- and three-epoch demographic model control allows for accurate recovery of the true DFE.

We next asked how well our predicted $B$-values match simulated $\pi$. We predicted $B$-values under the "true model", which used the input simulated demographic history and input DFE parameters, and twice under steady-state demography that allowed for interference correction: first, a "good" DFE fit (under the three-epoch model) that recovers the input parameters accurately, and a "bad" DFE fit (under the equilibrium demographic model) that is biased in the shape and scale of the reinferred DFE. We found predictions to be highly correlated (Figure S3), even with the poorly reconstructed DFE. In the larger-chromosome simulation (set B), selective interference is more pronounced, and the "true" model predicts larger values of $\pi$ than observed, likely due to not accounting for interference between selected alleles.

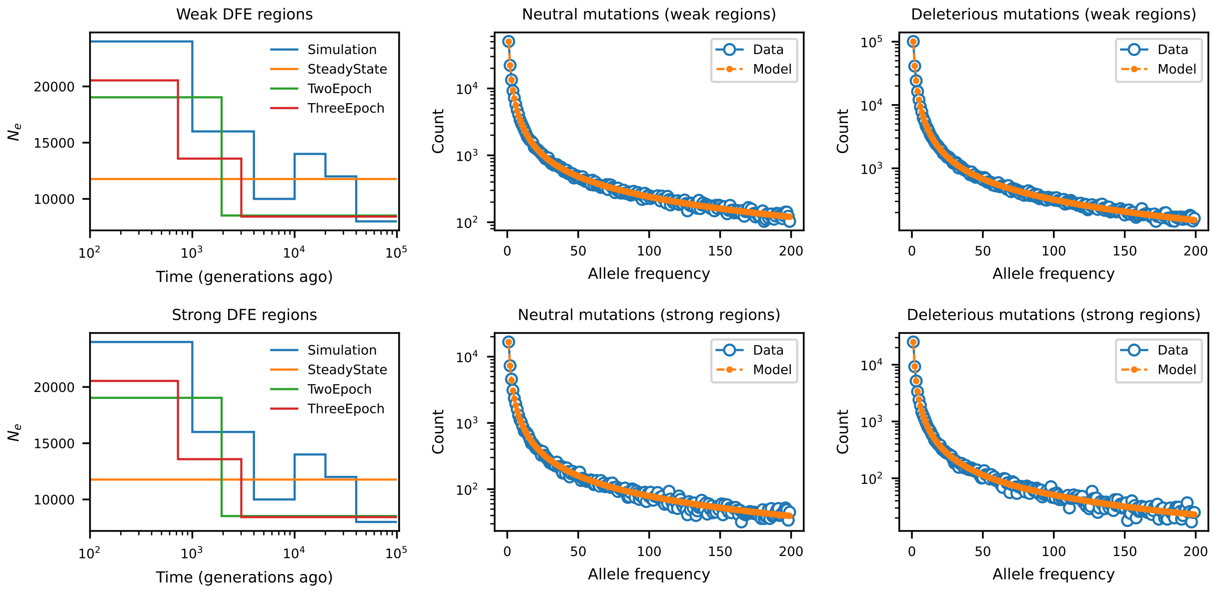

Figure S2: Left: Demographic models for the simulated data with two DFE classes, showing fits for a steady-state model, a two-epoch model with a single size change, and a three epoch model with two size changes. Center and right: fits to observed data under the three-epoch model. The model for selected alleles follows the DFE inferred, shown in Table S2.

#### 4.2 Equilibrium $B$-values fit the simulated data well

A plausible concern is that past fluctuations in human population size may preclude prediction of $B$-values under equilibrium and therefore the use of interference correction (which relies on constant $N_e$ through time). We show elsewhere (Barroso and Ragsdale, 2026, and Section 5.4) that the particular history of the Yoruba population does not introduce strong, transient BGS dynamics, so the assumption of equilibrium should be valid. Here, we explicitly investigate non-equilibrium departures in controlled conditions.

In the simulated scenario (Figure S2), we predict similar equilibrium and non-equilibrium $B$-values (Figure S3), with the difference that equilibrium models can better describe dips in diversity in simulation set B due to interference correction. Likewise, both kinds of $B$-maps lead to quantitatively similar SEM estimates (Figure S4). While there are some demographic scenarios for which assuming equilibrium will result in biased predictions of $B$-values (Barroso and Ragsdale, 2026), for modest size changes (as inferred in the YRI data) we expect steady-state predictions to be accurate and lead to robust SEM analyses (Figure S4).

#### 4.3 DAGs 3-4 are adequately represent causal effects in population genetics

When the landscape of genetic diversity is simultaneously influenced by drift (including population size changes), mutation rate variation and negative selection (both direct and indirect), without other factors such as biased gene conversion and other forms of selection, our proposed models fit the data remarkably well (Figure S5). Moreover, the distribution of $R^2_\pi$ across simulated 100 Mb chromosomes is essentially flat (Figure S6). These results help us interpret the human data estimates, which point to a poor fit in some chromosomes and genomic scales as well as large between-chromosome heterogeneity in $R^2_\pi$. Rather than implying that the evolutionary (causal) relationships among variables are incorrectly specified, low $R^2_\pi$ and poor model fit in some chromosomes can be explained by local estimation errors in the genomic maps and by unmodeled biological processes that are heterogeneous along the genome. Notably, $R^2_\pi$ stands above 80% when SEM is fit to error-free landscapes, similar to our empirical estimates in chromosomes 16 and 22,

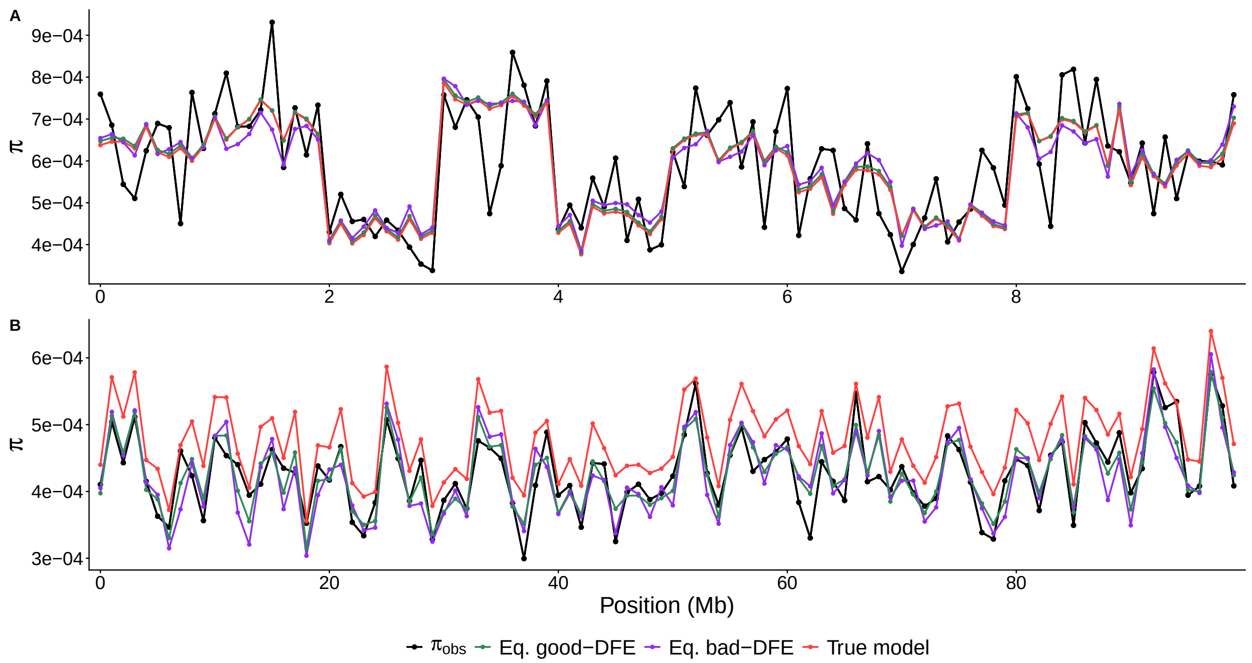

Figure S3: Observed (black) and predicted ($\mathbb{E}[\pi] = B \times \mathbb{E}[\pi_0]$, color, with $\pi_0$ using the simulated mutation map) landscapes of diversity in forward-in-time simulations. Red: $B$-values predicted using true model parameters (demography, shape and scale of the Gamma-distributed DFE). Green: $B$-values predicted under equilibrium, using demography-aware DFE parameters inferred from the SFS. Purple: $B$-values predicted under equilibrium, using DFE parameters inferred from the SFS assuming equilibrium. A: chromosome 1 from simulation set A (100x 10-Mb chromosomes), shown at the 100 kb scale. B: chromosome 1 from simulation set B (10x 100-Mb chromosomes), shown at the 1 Mb scale. In panel B, the non-equilibrium $B$-values overshoot the predictions of $\pi$ because they do not correct for selective interference.

showing that models that account for mutation rate variation can explain a large proportion of variation at the 1 Mb scale (see also Barroso and Dutheil (2023)).

Lastly, simulations set A leads to a curious negative correlation between $B$-values and $\pi$ (Figure S49). This apparent puzzle is solved once the DAG structure is taken into account. Mutation rates have a *positive* impact on $\pi$ but a *negative* impact on $B$, so the fork $\pi \leftarrow \mu \rightarrow B$ induces a negative correlation between the child nodes of $\mu$. In many biological systems (including humans), the effect of BGS on diversity is strong enough to counteract this and elevate the correlation of $B$ and $\pi$ above zero. In these simulations, however, with the particular parameters chosen, mutation rate variation dominates BGS. Because the causal path $B \rightarrow \pi$ is small, its contribution to the covariance matrix is overshadowed by the influence of mutation rates, leaving the simple correlation between $B$ and $\pi$ negative. Nevertheless, as DAGs 3-4 model these relationships, SEM correctly recovers a positive $B \rightarrow \pi$ coefficient (red circles in Figure S4).

#### 4.4 SEM chooses the correct model

By construction, our simulated landscapes of recombination, mutation, and selective constraint are independent of each other. Accordingly, in the vast majority of cases, model choice between DAGs 3 and 4 using the likelihood ratio test favors the simpler model which lacks the free covariance between $E$ and $\mu$ (Figure S7), except for three replicates from simulation set B where DAG 4 is preferred at the 10 kb scale. These could be due to forward-in-time scaling, by residual correlations among the maps (Figure S50), or simply by chance. We conclude that SEM can recover a correct parsimonious representation of evolutionary interactions when the input data is error-free.

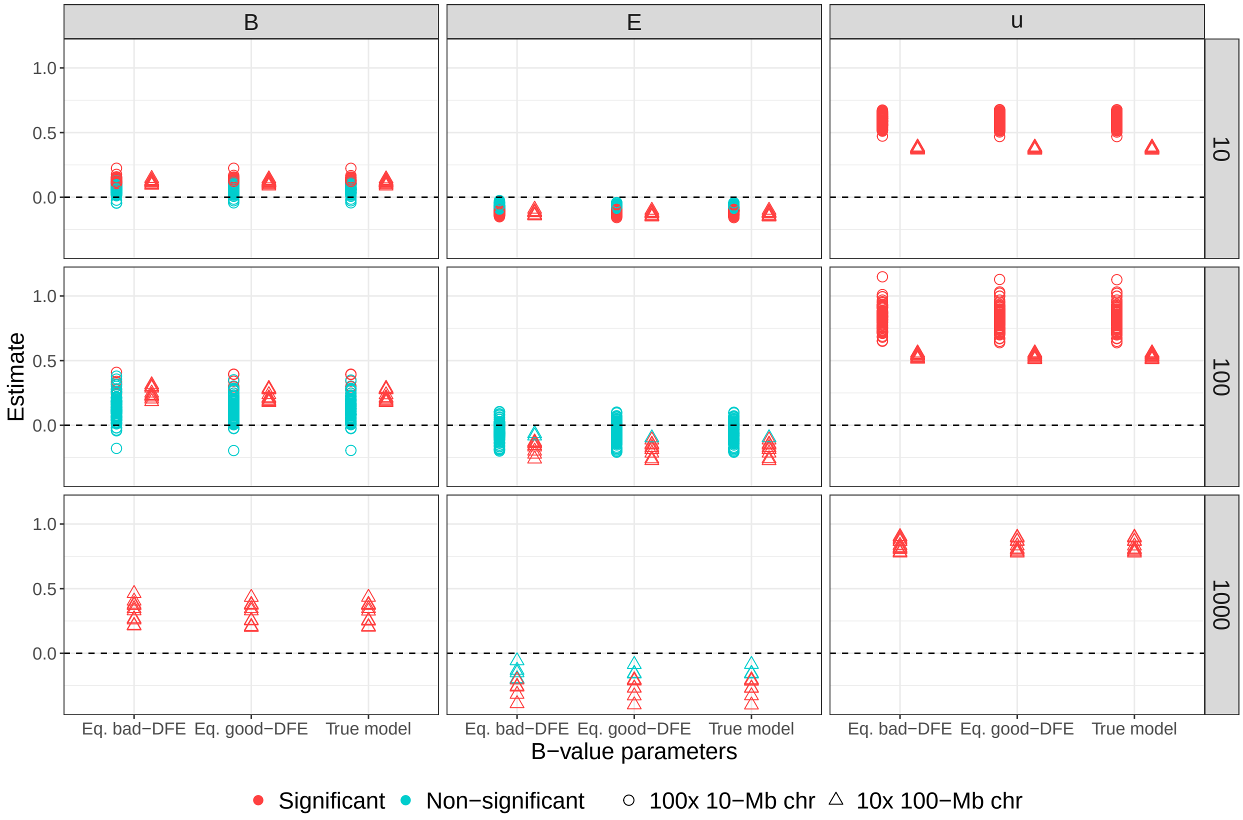

Figure S4: Inferred path coefficients $\mu \to \pi$, $B \to \pi$ and $E \to \pi$ from DAG 4 in forward-in-time simulations. X-axis divides results according to $B$-maps, which can be predicted either using the true model parameters (demography, shape and scale of the Gamma-distributed DFE); assuming equilibrium and using demography-aware DFE parameters inferred from the SFS; or assuming equilibrium and using DFE parameters inferred from the equilibrium SFS. Circles: simulation set A, 100 chromosomes of length 10 Mb each. Triangles: simulation set B, 10 chromosomes of length 100 Mb each. Row panels denote genomic scale (kb), with circles omitted from 1-Mb resolution due to insufficient data.

### 5 Assessing robustness to modeling assumptions

#### 5.1 Model mis-specification

DAGs 1-4 assume that the empirical maps of mutation, recombination and functional annotation–together with the background selection model based on unconditionally deleterious mutations–are adequate representations of the evolutionary process that generated the observed landscape of diversity. Inferred human landscapes deviate from these assumptions, leading to model mis-specification that lowers fit indices and biases estimates of some path coefficients. To evaluate our proposed causal models in scenarios without any model mis-specification, we simulated the landscape of diversity directly from population-genetic expectations obtained with `moments++`. Unlike the forward-in-time simulations described in Section 4, these simulations draw from the model predictions and are therefore tailored to the human data we analyze in the main text.

Let $w$ index non-overlapping windows of fixed length $L$ and average predicted diversity per site $\mathbb{E}[\pi_w]$. In each, we drew the number of polymorphic sites from a Binomial distribution with $p = \mathbb{E}[\pi_w]$ and averaged over all sites:

$$X_{w,s} \sim \text{Binomial}(L, \mathbb{E}[\pi_w]), \tag{S24}$$

$$\hat{\pi}^{sim}_{w,s} = \frac{X_{w,s}}{L}. \tag{S25}$$

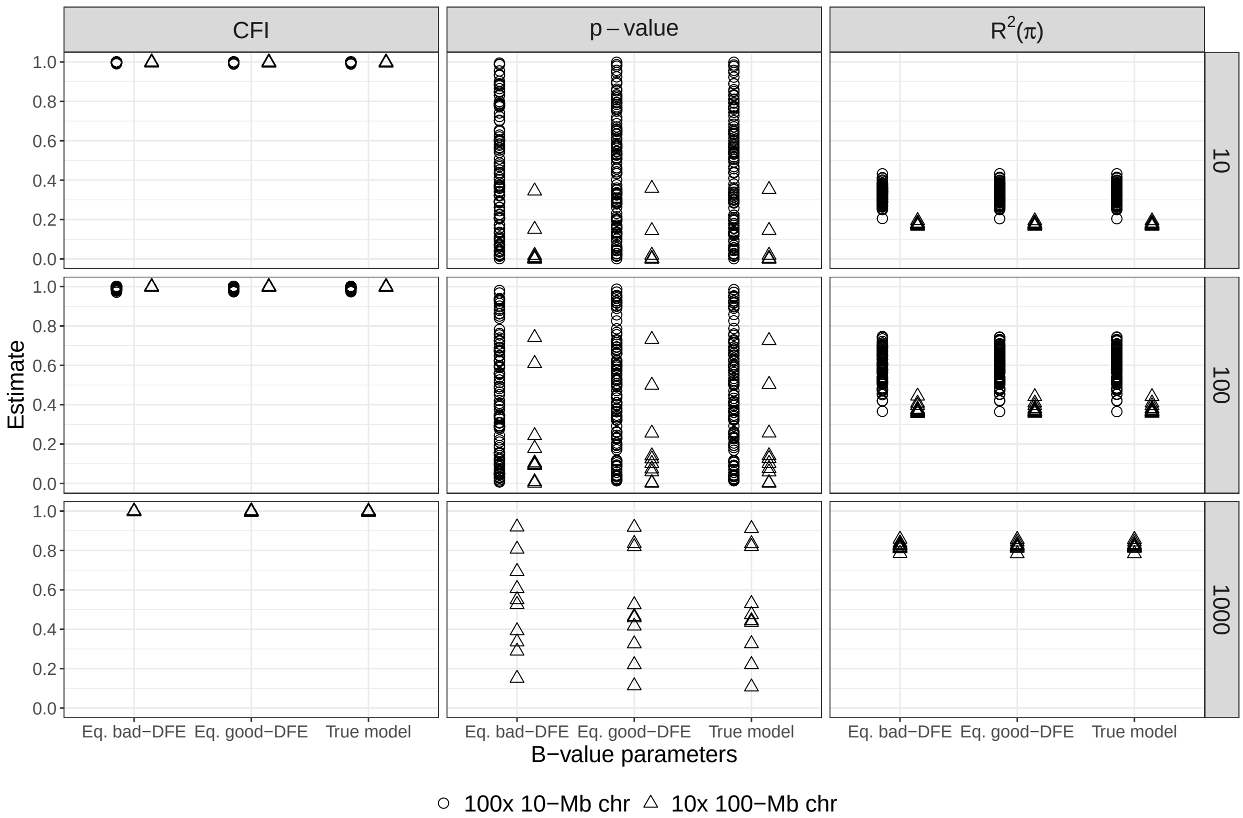

Figure S5: Conditional fit index (CFI), $p$-values and $R^2_\pi$ of DAG 4 in forward-in-time simulations. X-axis divides results according to $B$-maps, which can be predicted either using the true model parameters (demography, shape and scale of the Gamma-distributed DFE); assuming equilibrium and using demography-aware DFE parameters inferred from the SFS; or assuming equilibrium and using DFE parameters inferred from the equilibrium SFS. Circles: simulation set A, 100 chromosomes of length 10 Mb each. Triangles: simulation set B, 10 chromosomes of length 100 Mb each. Row panels denote genomic scale (kb), with circles omitted from 1-Mb resolution due to insufficient data.

This amounts to simulating heterozygosity in one diploid genome. We then averaged over $S$ replicate diploids:

$$\pi_w^{sim} = \frac{1}{S}\sum_{S} \hat{\pi}_{w,s}^{sim}. \tag{S26}$$

We used $S = 25$ and $L = 10^4$, then further averaged the maps to obtain simulated diversity at 100 kb and 1 Mb. These simulations ignore masking patterns observed in real data. They further neglect linkage and the shared genealogical relationships among sites (which are simulated independently). Our goal was merely to generate sampling noise around $\mathbb{E}[\pi_x]$, avoiding a perfect match between $B$, $\mu$ and $\pi$ which would lead to numerical instability such as non-invertible matrices when fitting SEM.

We then fit SEMs using the landscapes of $\pi^{sim}$ instead of $\pi^{obs}$. Note that this procedure is not equivalent to simulating variables directly from the DAGs; for example, the average $B$-value of a focal window still depends on $\mu$, $r$, $E$ in distant genomic regions. As such, our simulation guarantees that the path coefficients directly associated with $\pi$ ($B \to \pi$, $E \to \pi$ and $\mu \to \pi$) faithfully represent the data-generating process. But the overall adequacy of the DAGs remains to be tested as it relies on the entire covariance matrix, hence on accurate representations of all causal relationships. We found that DAGs 3-4 provide an excellent fit to these simulated data (Figure S46), and that estimated path coefficients agree with biological intuition and forward simulations (Figure 4).

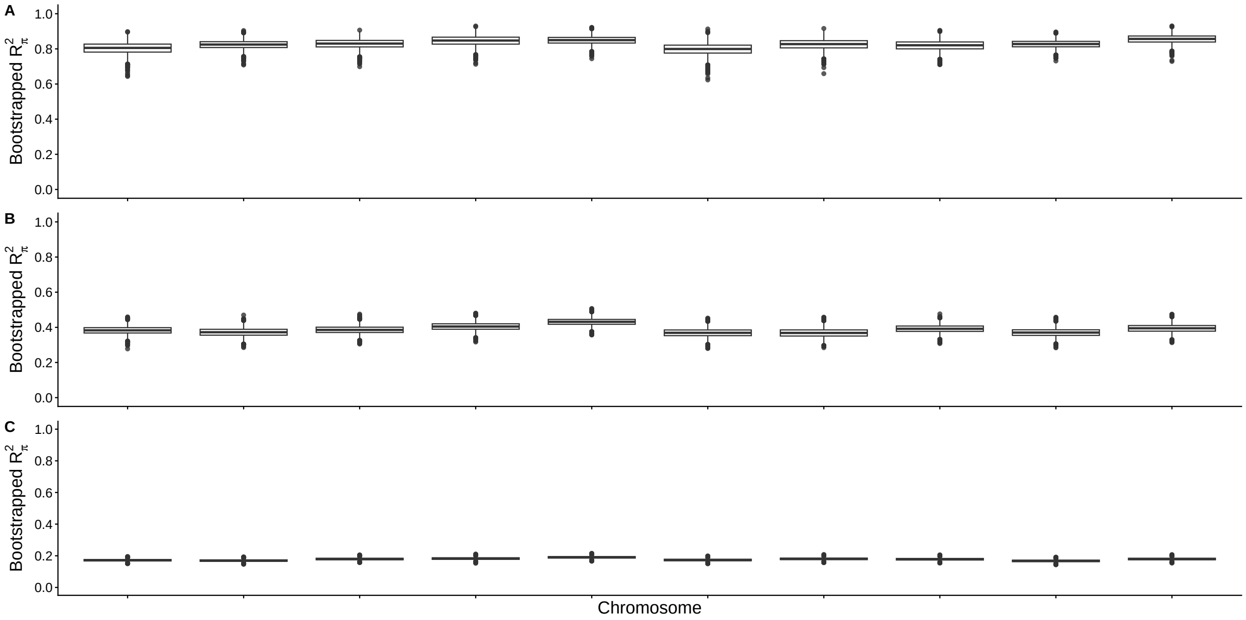

Figure S6: Bootstrapped $R^2_\pi$ across ten independently simulated chromosomes, each of length 100 Mb. A: 1 Mb scale. B: 100 kb scale. C: 10 kb scale. Each boxplot summarizes $R^2_\pi$ among 5,000 bootstrap replicate fits under DAG 3. Shown are SEM results obtained with $B$-values computed at steady state (with interference correction) using accurately inferred DFEs (with the three-epoch demographic control during DFE inference), but the other two $B$-maps lead to similar $R^2_\pi$.

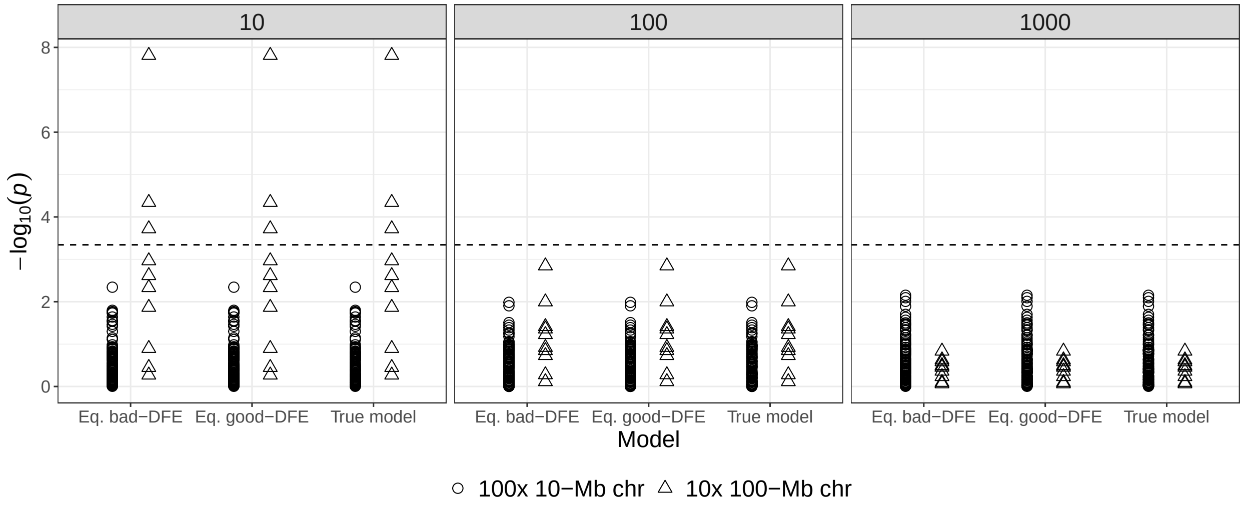

Figure S7: Model choice between DAGs 3 and 4 in forward-in-time simulation sets (shape). Y-axis shows $-log_{10}(p)$ of the likelihood ratio tests, with lower value indicating that the simpler (and correct) model (DAG 3) is preferred.

#### 5.2 Sensitivity to alternative causal models

Our simulation study (Sections 4, 5.1) demonstrates that DAGs 3 and 4 are reasonable descriptions of the causal relationships among genomic landscapes, particularly in the absence of map errors. Here, we test whether alternative representations, which are overall incorrect but still capture important causal effects,

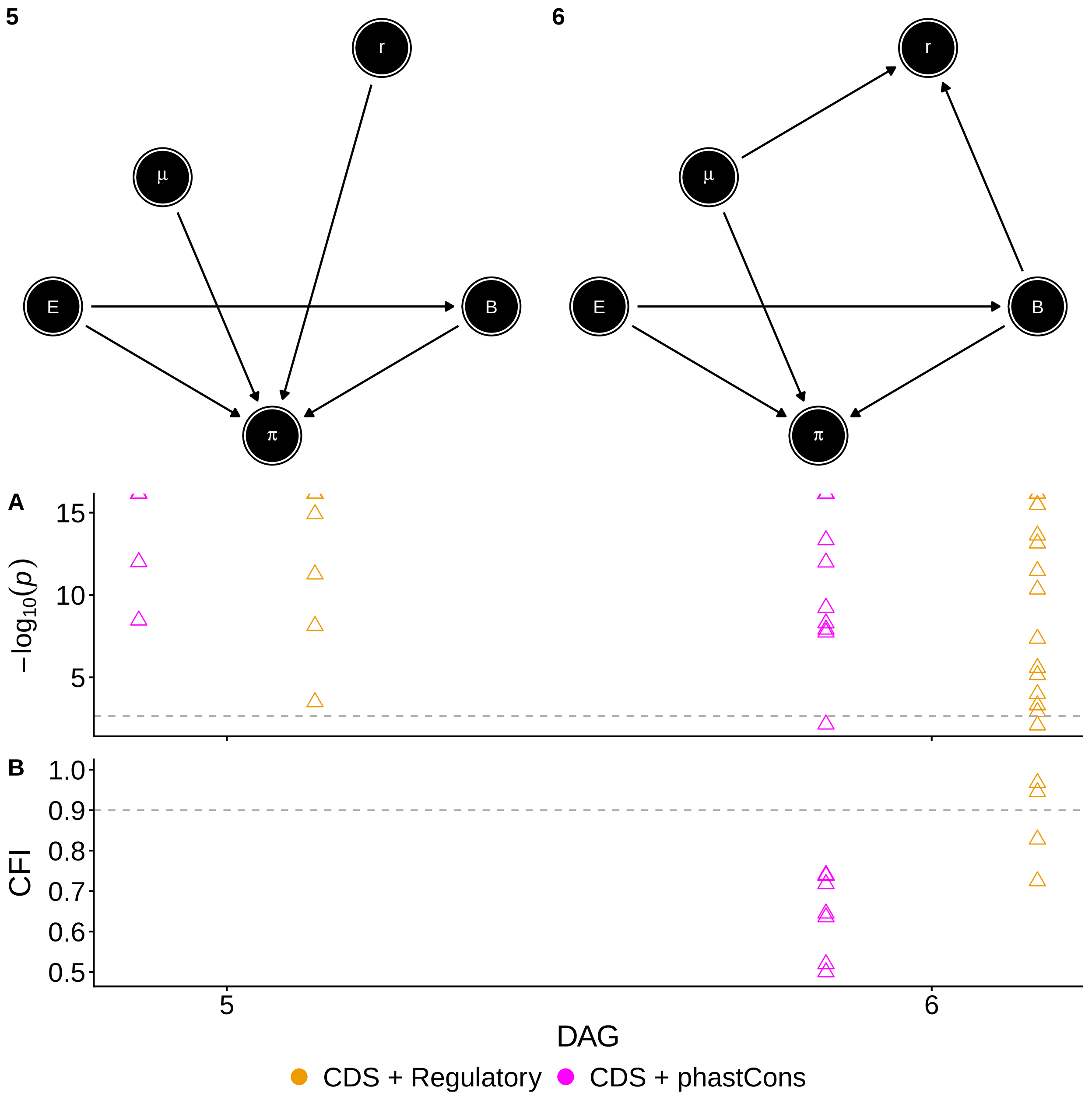

Figure S8: Goodness-of-fit of alternative causal models (DAGs 5 and 6) to 1-Mb human landscapes. Points show results for each autosome, divided by annotation group (color). A: $p$-values from the $\chi^2$ test on a $-log_{10}$ scale where lower values indicate better agreement with data. Dashed line shows significance threshold after Bonferroni correction (0.05/22). Several points cluster at the top. B: comparative fit indices, where higher values indicate better agreement with data. Dashed line shows the rule-of-thumb for an acceptable fit (0.90). The poor fit in several autosomes prevents `lavaan` from reporting CFIs, which are omitted from the plot. Exonic DFEs are split among deciles of constraint.

are able to fit the data well (Figure S8). Namely, DAG 5 treats recombination as exerting a direct effect on diversity (bypassing the chains $r \to \mu \to \pi$ and $r \to B \to \pi$), but fails to describe the correlation of $r$ with $\mu$ and $B$. By contrast, DAG 6 reverses the causal direction between recombination and its child nodes: it reproduces the observed covariance of $r$ with $\mu$ and $B$, but implies that recombination has no effect on $\pi$.

We find that DAGs 5 and 6 fit poorly to human data under both *CDS+phastCons* and *CDS+regulatory* annotation sets. The $\chi^2$ test rejects DAG 5 across all autosomes, while DAG 6 is rejected in all but one of them (Figure S8A). Meanwhile, the poor fits of these models prevent `lavaan` from even reporting CFI in most cases (Figure S8B). Taken together, these results provide examples of SEM rejecting models that are clearly incorrect representations of causal relationships (by design), reinforcing SEM's ability to identify the causal relationships that best explain the covariance structure in observed data.

This exercise also suggests that we have sufficient statistical power to reject models at the 1 Mb scale, where several chromosomes provide limited sample size. Because candidate models can only be falsified (instead of proven correct), model compatibility strictly indicates that the data do not provide sufficient evidence to reject them. Fit indices from DAGs 5-6 suggest that 1 Mb landscapes have enough data to reject models that partially describe important relationships among variables yet are incorrect overall. These results corroborate the reasonable agreement between human data and DAGs 3-4 that we find in the main text.

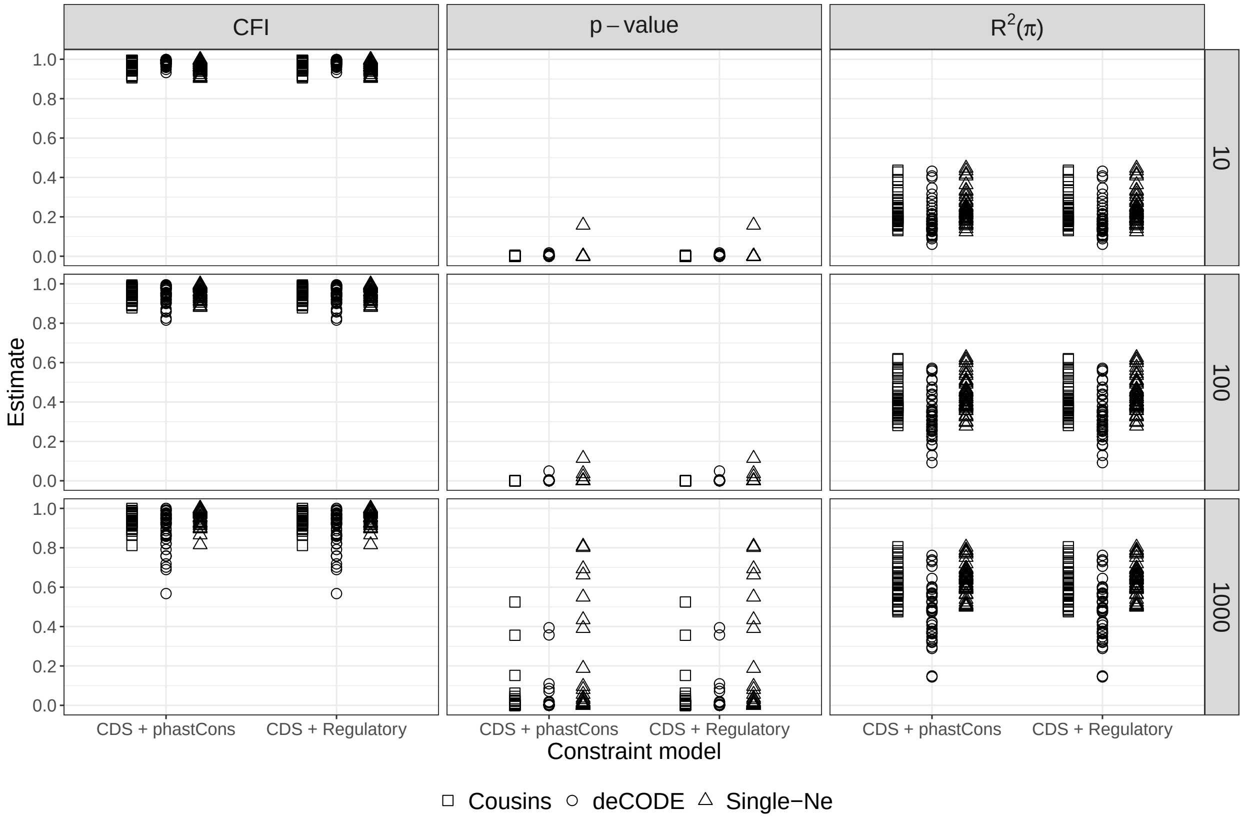

Figure S9: Conditional fit index (CFI), $p$-values and $R^2_\pi$ of DAG 4 under alternative models. Circles: predictions using a 24-epochs approximation of the Cousins et al. (2024) demography. Triangles: predictions using the same equilibrium $N_e$ across autosomes (weighted average of chromosome-specific estimates). Rows denote genomic scale (kb). Exonic DFEs are split among deciles of constraint. Although $R^2_\pi$ and CFI as similar among alternative models, equilibrium $B$-values lead to higher $p$-values, likely due to interference correction.

#### 5.3 Chromosome-specific $N_e$

Our main results are obtained with $B$-value predictions that use chromosome-specific $N_e$ estimates. It is in principle plausible that the chromosome-specific signatures that we find (regarding model choice, $R^2_\pi$ and estimated path coefficients) are artifacts from the use of chromosome-specific $\hat{N}_e$. To assess the robustness of

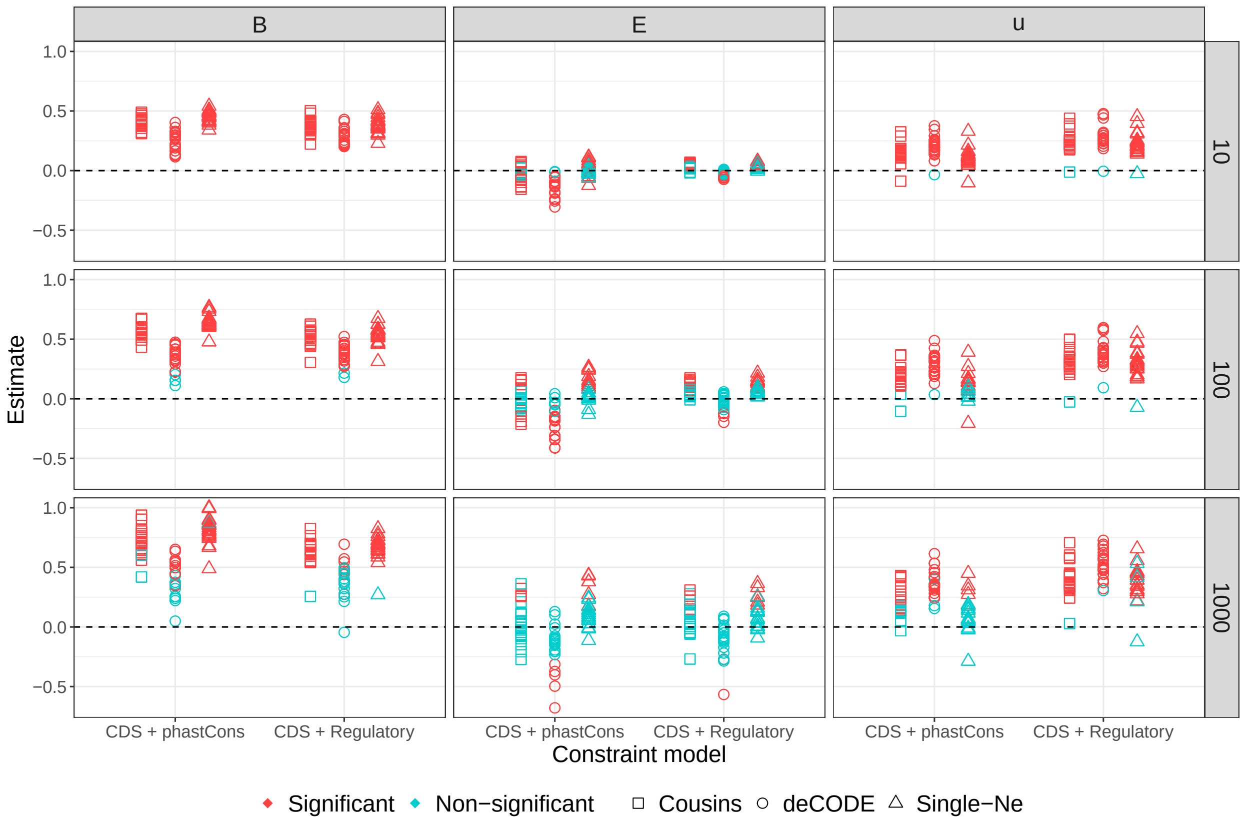

Figure S10: Inferred path coefficients $B \to \pi$, $E \to \pi$, and $\mu \to \pi$ from DAG 4 under alternative models. Squares: predictions using a 24-epochs approximation of the Cousins et al. (2024) demography. Circles: prediction under equilibrium demography, using the deCODE (rather than `pyrho`) recombination map. Triangles: predictions using the same equilibrium $N_e$ across autosomes (weighted average of chromosome-specific $\hat{N}_e$). Rows denote genomic scale (kb). Colors indicate significance after Bonferroni correction (0.05/22). Exonic DFEs are split among deciles of constraint.

our results, we repeat the data analyses with $B$ and $\mathbb{E}[\pi_0]$ predicted across the genome using the autosome-wide average $\hat{N}_e$ weighted by chromosome length.

Results obtained with these homogeneous $N_e$ models (triangles in Figures S9, S10) are consistent with our main findings (Figures 3, 4), indicating that the strong support for chromosome-specific effects that we find in the main text are not due to our use of chromosome-specific $\hat{N}_e$ in predictions, but instead represent real biological differences.

#### 5.4 Non-equilibrium demography

We have previously shown that population size changes can dramatically affect $B$-values, especially under strong and relatively recent bottlenecks (Barroso and Ragsdale, 2026). However, in that same study we find that equilibrium demography is a good approximation in some scenarios. Since assuming equilibrium allows us to correct for interference among constrained sites (our heuristic requires that $N_e$ is constant over time), there is a trade-off between equilibrium and non-equilibrium models of background selection. In the case of YRI, for which inferred historical size changes are not extreme (Cousins et al., 2024), the balance tips off in favor of interference correction, which increases the accuracy of $B$-value prediction in regions of strong BGS.

We aimed to assess how departures from equilibrium might influence our results. To capture temporal fluctuations in the drift-effective $N_e$, we approximated the Cousins et al. (2024) demographic curve as a

24-epochs model (Figure S11). We used their estimates from regions with $0.9 < B \leq 1$ so that $N(t)$ is not strongly affected by BGS. We used this model to construct a demography-aware lookup table, and predicted $B$-maps for the six combinations of mutation maps crossed with *CDS+phastCons* and *CDS+regulatory* annotations. We found that SEM results under the Cousins et al. (2024) demography are largely consistent with equilibrium estimates (squares in Figures S9, S10), although fit indices are slightly worse (particularly $p$-values from the $\chi^2$ test evaluating the DAG), likely due to the non-equilibrium model not correcting for selective interference.

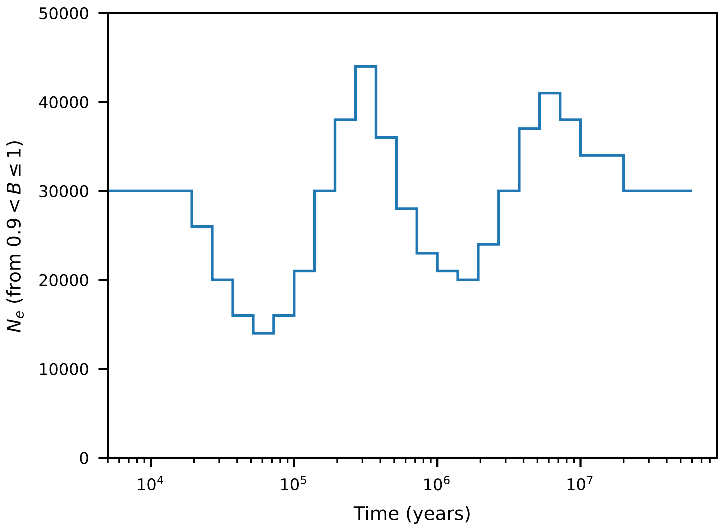

Figure S11: The approximated Cousins et al. (2024) demographic model (see Figure 2 therein).

#### 5.5 Choice of recombination map

`pyrho` estimates recombination rates from observed patterns of linkage disequilibrium (LD) between all pairs of loci across the genome, accounting for single-population demographic history (Spence and Song, 2019). However, its estimates derive from neutral models that ignore the effects of selection, and its reliance on observed SNPs might introduce circularity in our framework. Therefore, we evaluated the robustness of our results using a pedigree-based map from the deCODE project (Halldorsson et al., 2019). We used this map to both fit $N_e$ and predict $B$-values in each chromosome, then fit DAGs 3 and 4 using SEM.

A straightforward comparison is difficult because the deCODE recombination map has lower coverage than the `pyrho` map, especially in the distal ends of chromosomes. In regions lacking coverage, predicted $B$-values are artificially lowered due to using lower recombination rates (sometimes zero) than the actual rates in those regions (Figures S12, S19). Thus, fitting SEM with deCODE-derived maps results in somewhat worse fit indices and lower $R^2_\pi$ than other models (Figure S9). Estimated path coefficients are qualitatively similar, so despite the poorer fit, we draw similar conclusions to those using the `pyrho` map (Figure S10; the path coefficient $E \rightarrow \pi$ is always positive, in agreement with both simulations and intuition). In contrast to the historical estimates of recombination reported by `pyrho`, deCODE estimates current rates of crossing over in the Icelandic population (Halldorsson et al., 2019). This makes it less prone to the confounding of selection, but perhaps less applicable to our case study.

Although $B$-maps estimated with the YRI-`pyrho` and the deCODE recombination maps generally agree, their differences near the ends of chromosomes are noticeable (Figures S12, S19). Excluding genomic windows where deCODE estimates are $< 10^{-9}$ increases the correlations between `pyrho`-derived and deCODE-derived $B$-maps from $\approx 0.88$ to $\approx 0.93$ (Figures S55, S56). Yet these correlations are still substantially lower than between $B$-maps predicted for the same annotation set but using different mutation maps, demographic models or DFEs, which were all $> 0.99$. This suggests that, among genomic features relevant to BGS, the recombination map has the strongest influence on the $B$-value landscape, at least in humans.

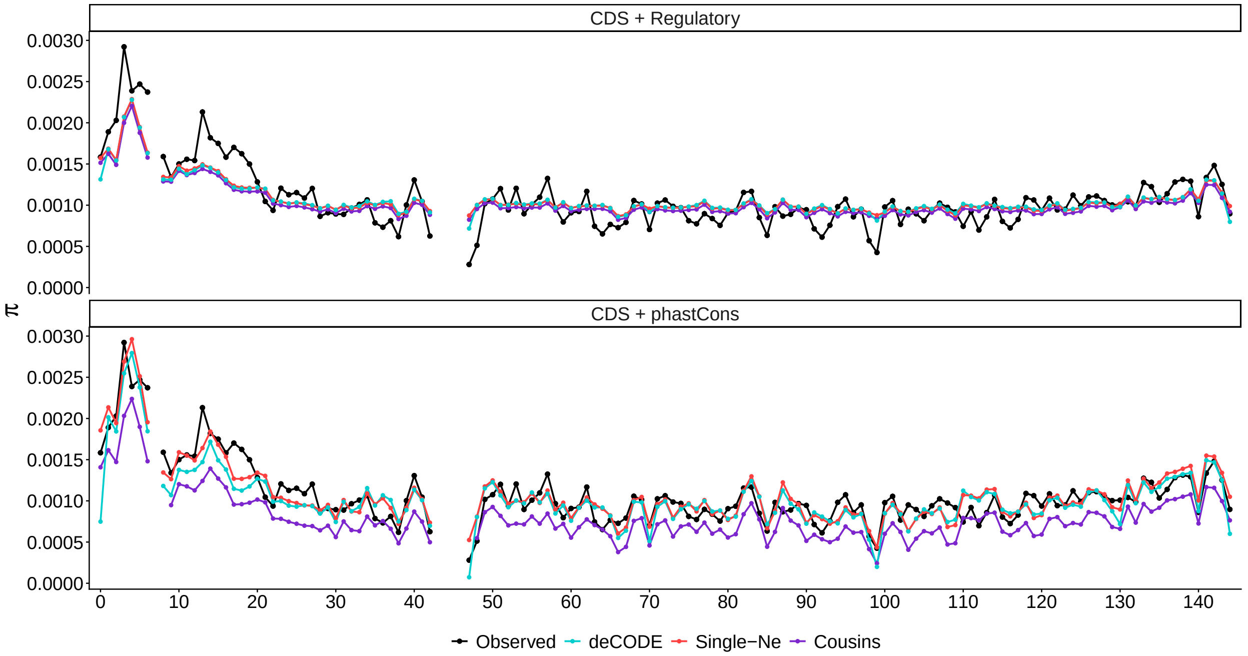

Figure S12: Observed ($\hat{\pi}$, black) and predicted ($\mathbb{E}[\pi] = B \times \mathbb{E}[\pi]$, color) diversity landscapes under alternative models, here shown for the Roulette mutation map in chromosome 8 (1 Mb scale). Red: $B$-values assuming the genome-wide average estimate $N_e$ across autosomes. Purple: $B$-value assuming the Cousins et al. (2024) demography instead of the fit $N_e$. Light blue: $B$-value assuming the deCODE recombination map and its corresponding chromosome-specific $N_e$ estimates (Figure S58). Note that deCODE estimates are typically zero at the tips of the chromosomes, predicting very low $B$-values in some cases. Although non-equilibrium models have no free parameters, they still capture the absolute scale of the diversity landscape reasonably well. Here the exonic DFE is split into deciles of constraint. See figure S19 for chromosome 22 at 100 kb scale.

#### 5.6 Choice of phastCons cut-off

In their recent SFS-based inference of the DFE from candidate functional noncoding phastCons, Di et al. (2025) found statistical support for negative selection acting on the top 60% scores ($\approx 30\%$ of the human genome). Therefore, our main results obtained under the *CDS+phastCons* models are based on these elements, which represent substantially more constrained sites than the phastCons annotations included by Murphy et al. (2022) as direct targets of selection ($\approx 6\%$ of the genome).

To investigate the impact of different thresholds for including phastCons elements in our analyses, we fit models using the top 5%, 10%, 20% 30% or 40% noncoding scores. For each of these re-defined *CDS+phastCons* annotations, we fit $N_e$ (Figure S59), predicted $B$-values and $\mathbb{E}[\pi]$, and executed the SEM pipeline. We found that results are in close agreement with the full *CDS+phastCons* models, with a few notable exceptions that clarify the counterintuitive estimates of path coefficients in the main text (Figure 4, Section 5.1). Namely, $\mu \to \pi$ (chromosome 19) is only significantly negative after including 20% noncoding phastCons or more, while $E \to \pi$ only shows positive estimates after including 30% noncoding phastCons or more (Figure S13). Somewhat surprisingly, $R^2_\pi$ and the conditional fit index (CFI) are mostly unchanged as we include more or fewer phastCons bins (Figure S14). On the one hand, this suggests that our inferences are robust to the precise threshold chosen for the inclusion of phastCons elements. But on the other hand,

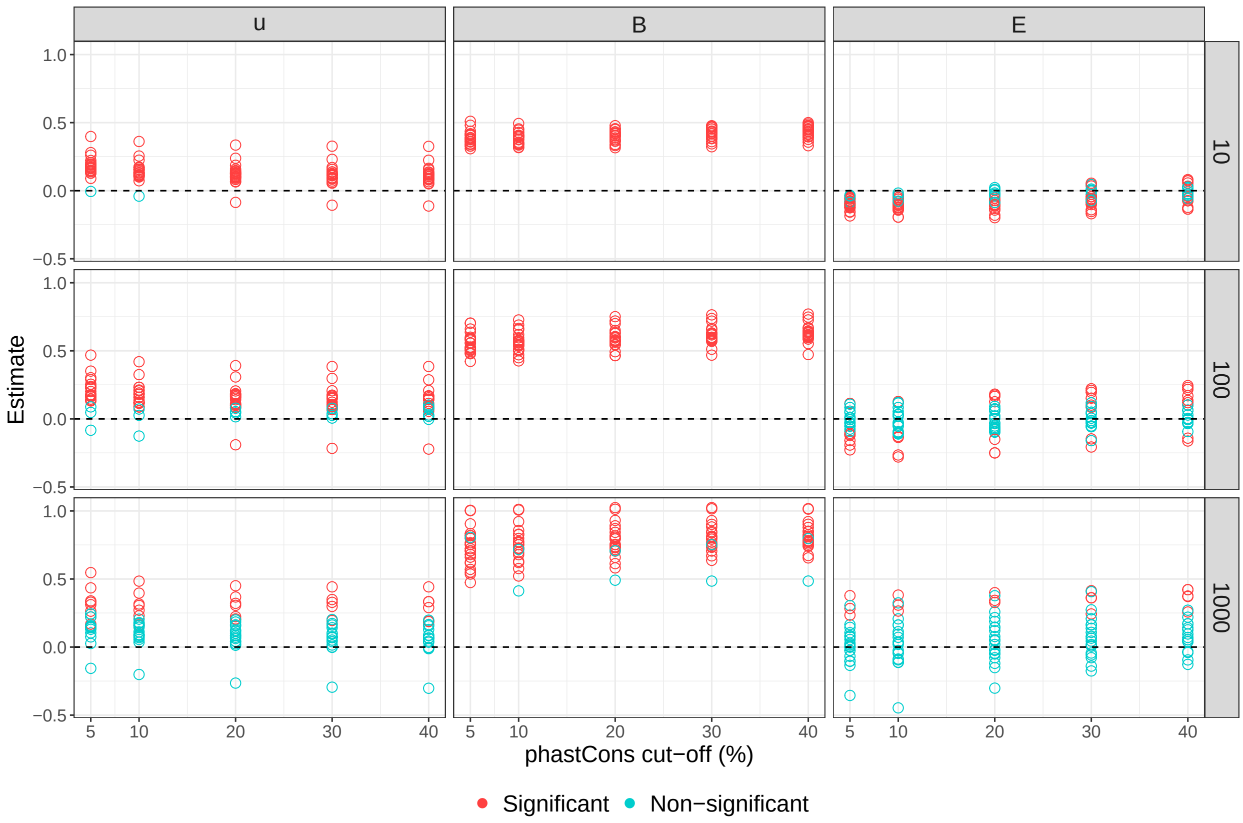

Figure S13: Inferred path coefficients $\mu \to \pi$, $B \to \pi$ and $E \to \pi$ of DAG 4, here for different amounts of noncoding phastCons included as targets of direct selection. Rows denote genomic scale (kb). Colors indicate significance after Bonferroni correction (0.05/22). The exonic DFEs are split among deciles of constraint.

that $R^2_\pi$ and CFI do not increase with the inclusion of more putatively constrained sites suggests either that the predominant BGS effect in the noncoding genome is contributed by the most constrained bins, or that uncertainty in the identification of truly constrained elements increases as more bins are included. Because phastCons elements are based on measured constraint across a phylogeny, rather than functional predictions, more work is needed to evaluate noncoding annotations in the human genome, especially given that many annotated phastCons elements fall outside of putatively regulatory regions (Section 5.7).

#### 5.7 Overlap of regulatory and phastCons annotations

The final concern we address is that some of the noncoding elements we analyze may not be direct targets of negative selection. The two sets of noncoding annotations we considered were *regulatory*, i.e., promoters and enhancers, or *phastCons* (bins 0-60, corresponding to $\approx 30\%$ of the callable noncoding genome, Section 1.2.1). Promoters and enhancers have been annotated by a hidden Markov model that classifies discrete 200 base-pair windows into putatively functional, noncoding categories (Ernst and Kellis, 2012; Vu and Ernst, 2022, Section 1.2.1). These calls may include neutral sites due to their limited resolution, but we would expect at least some portion of regulatory-annotated elements to be subject to negative selection, as suggested by DFE inference (Di et al., 2025). We observe 54.9% of promoters to be annotated as phastCons and 59.6% of enhancers annotated as phastCons. Both these proportions are much larger than the $\sim 30\%$ expected if those regions were randomly distributed with respect to phastCons scores, confirming that regulatory elements are indeed under selective constraint.

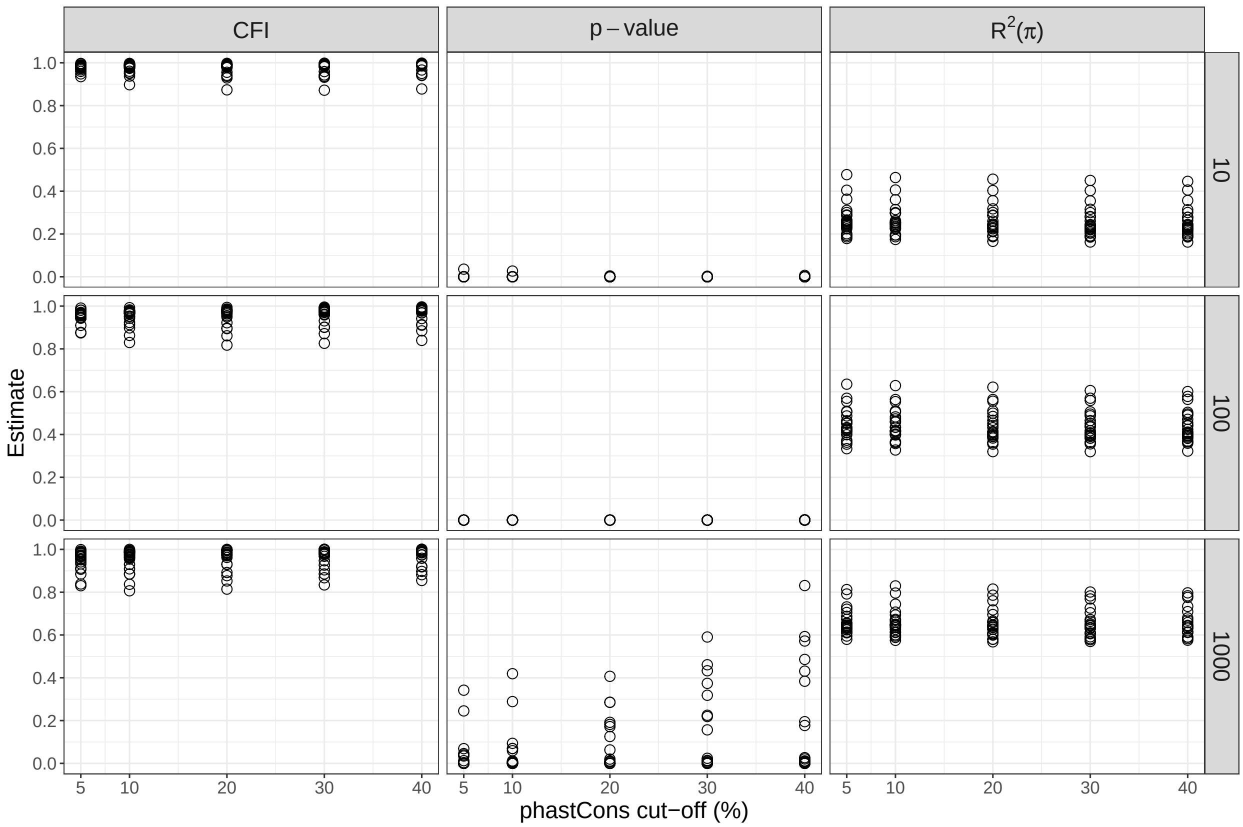

Figure S14: Conditional fit index (CFI), $p$-values and $R^2_\pi$ of DAG 4 for different amounts of noncoding phastCons included as targets of selection. Rows denote genomic scale (kb). Exonic DFEs are split among deciles of constraint. The higher $p$-value (better fit of the DAG) of higher cut-offs (1 Mb) corroborates some of the weaker phastCons calls.

| phastCons | Prop. in promoters | Prop. in enhancers |
|---|---|---|
| All | 0.0212 | 0.1885 |
| Bin 0-5 | 0.0323 | 0.2403 |
| Bin 5-10 | 0.0226 | 0.1947 |
| Bin 10-15 | 0.0196 | 0.1846 |
| Bin 15-20 | 0.0188 | 0.1819 |
| Bin 20-25 | 0.0186 | 0.1807 |
| Bin 25-30 | 0.0186 | 0.1804 |
| Bin 30-35 | 0.0188 | 0.1801 |
| Bin 35-45 | 0.0191 | 0.1799 |
| Bin 45-50 | 0.0198 | 0.1799 |
| Bin 50-55 | 0.0204 | 0.1800 |
| Bin 55-60 | 0.0209 | 0.1803 |

Table S3: Proportion of phastCons elements (bins 0-60) that are annotated as promoters or enhancers.

There is enrichment for both promoter and enhancer elements among the strongest-constrained phastCons bins (0-5 and 5-10) (Table S3). But overall, there are many more sites annotated as phastCons compared to regulatory elements (at least when using the 0-60 range of phastCons scores), and $\approx 80\%$ of such noncoding phastCons are non-regulatory (at least, not annotated as either promoters or enhancers). Thus, much of the signal of linked selection in the phastCons models are due to linked selection from non-regulatory regions.

### 6 Evaluating the limits to $R^2_\pi$ imposed by genetic drift

In section 3.8 of their Supplemental Material, Buffalo and Kern (2024) develop an approximation to the theoretical limit in $R^2$ imposed by the stochastic fluctuations in coalescence times across the genome. They find this limit to be roughly 67% in the Yoruba population at the 1 Mb scale (based on their model predictions), and conclude that their implementation of the Santiago and Caballero (2016) background selection model accounts for almost all variation in diversity that can possibly be explained (the remainder $\approx 33\%$ would be attributed to genealogical noise, or drift). However, their derivation assumes a sample of size two, where the variance in coalescence times is maximized. In contrast, estimating the average pairwise genetic diversity ($\hat{\pi}$) using sequence data from a cohort of 106 individuals from the Yoruba population (Byrska-Bishop et al., 2022) should result in weaker effects of drift. That is, a higher sample size could in principle allow statistical models to explain more than 67% of the variation in diversity at the 1 Mb scale, in agreement with our findings.

Although Buffalo and Kern (2024) are careful in caveating their approximation, our contrasting results (with $R^2_\pi > 70\%$ in some models at the 1 Mb scale) motivated us to explore the limits of $R^2_\pi$ using simulations. We used `msprime` (Kelleher et al., 2016; Baumdicker et al., 2022) to simulate windows of varying size ($L$ base pairs), assuming steady-state demography with $N_e = 10^4$ and both per-base $u$ and $r$ of $10^{-8}$. The estimated $R^2_{\text{drift}}$ from Supplementary section 3.8 in Buffalo and Kern (2024) relies on the expected number of segregating sites in a sample of size two haploids (or a single diploid) for a region with given $\theta = 4N_e\mu L$ and $\rho = 4N_e rL$, given by Wakeley (2009) as

$$\text{Var}(S_2) = \theta + \frac{2\theta^2}{\rho^2}\int_0^\rho (\rho - x) f_2(x)dx,$$

where

$$f_2(\rho) = \frac{\rho + 18}{\rho^2 + 13\rho + 18}.$$

Before considering BGS, we first compared this prediction to observed variance in the total diversity in a sample of a given number of diploids ($n$) ranging from one to over 100 (Figure S15A). We find that this provides a good approximation for $n = 1$ and for small window sizes, but begins to break down for larger window sizes ($L \gtrsim 10^4$ base pairs) and overestimates $\text{Var}(\pi)$ for larger $n$.

To demonstrate the dependence of $R^2_{\text{drift}}$ on $n$, for a given window size $L$, we simulated $m = 10^9/L$ windows. Within each window, we assigned a randomly drawn $B$ value with mean $\bar{B} = 0.7$ and $\text{Var}(B) = 0.0075$, restricted to $[0.1, 1]$ (this gave an $R^2_{\text{drift}} \approx 0.68$ at 1 Mb scale, but was arbitrarily chosen otherwise). Sequence data was simulated with population size $N_e \times B$ to approximate the effect of background selection within that window. We then computed theoretical $R^2_{\text{drift}} = 1 - \frac{V_{\text{res}}}{V_{\text{tot}}}$ (in the notation of Buffalo and Kern, 2024), and compared to observed $R^2$ by computing the squared correlation between $B$ values within each window and observed $\pi$. Across scales, we find that observed $R^2$ depends sensitively on $n$, especially for small $n$, and it exceeds the theoretical prediction as $n$ becomes large due to reduced residual error from drift (Figure S15B).

The theoretical prediction appears to be biased by both the breakdown of predictions for large window sizes in a sample size of two haploid copies (Figure S15A) and the reduction in variance of observed $\pi$ for larger sample sizes, making it difficult to draw firm conclusions about the true theoretical bounds of $R^2_{\text{drift}}$. The simulations here also assess $R^2$ in a simplified scenario. Including additional causal parents of $\pi$ in forward simulations with selection and mutation rate variation (Section 4.3) further increase the share of variance in diversity that can be explained by the models, leaving less room for drift.

### 7 Predicting differences in $R^2_\pi$ among autosomes

Are there genomic features that predict the large heterogeneity in $R^2_\pi$ among chromosomes (Figure S38)? We tested whether chromosomal differences in average mutation, recombination, GC content, and density of

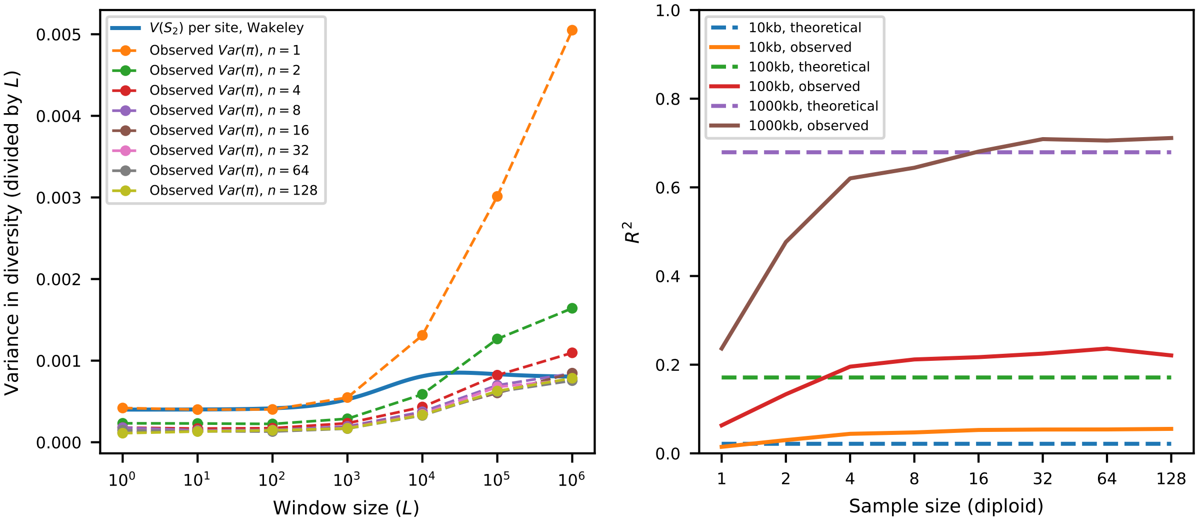

Figure S15: Left (A): Observed variance in $\pi$ across replicates of simulations of a given window size, ranging from one to one million base pairs. The theoretical prediction is valid for $n = 1$ diploid, but breaks down for large window sizes. Right (B): Observed $R^2$ against the theoretical limit of $R^2_{\text{drift}}$, which can be exceeded for larger sample sizes. In both, $N_e = 10^4$ and $u = r = 1e-8$, and see Section 6.

constrained sites could be associated with our differential ability to explain genetic diversity across autosomes. Since these variables are themselves correlated (Freudenberg et al., 2009; Romiguier et al., 2010), we used linear regression to estimate independent effects,

$$R^2_\pi = \beta_1 \overline{GC\%} + \beta_2 \overline{E} + \beta_3 \overline{\mu} + \beta_4 \overline{r} + \varepsilon,$$

which estimates the linear effects of features on the variance in diversity explained by DAG 4 in each chromosome. Although there are visual trends (Figures S51–S54), the small sample size (22 autosomes) limits our power to detect significant associations. The average mutation rate stands out as the only significant predictor of $R^2_\pi$. Yet the linear models explain much of the variance in $R^2_\pi$ among chromosomes at scales $< 1$ Mb (Figure S16). An in-depth investigation of such inter-chromosomal differences is beyond our scope.

### 8 Tools and properties of genomic structural equation models

#### 8.1 Measurement models and goodness-of-fit

Structural equations models (SEM) are able to take measurement error of observed variables into account (Shipley, 2016; Gustafson, 2021). This is done with *indicators* (or measurements) from a latent variable we are trying to represent. In the context of genomic landscapes, an example latent would be the true (unknown) mutation map, with empirical estimates (Roulette, Carlson and gnomAD) functioning as its indicators. Measurement models separately estimate the error in each indicator. Similarly to principal component analysis, they can then construct a synthetic latent variable. This is useful not only because of estimation noise but also when available maps capture distinct aspects of of the underlying construct.

Since four indicators are required to both estimate parameters and evaluate the goodness-of-fit of the measurement model, we first fit a model (1 Mb) that included the iSMC mutation map, which is available for chromosome 1 (Barroso and Dutheil, 2026): $\mu = \lambda_1\, \mu_{\text{roulette}} + \lambda_2\, \mu_{\text{carlson}} + \lambda_3\, \mu_{\text{gnomad}} + \lambda_4\, \mu_{\text{ismc}}$. This served as a test case to evaluate the suitability of measurement models of mutation, where we expect estimation noise in the indicators to be more pronounced. After including a covariance term between Carlson and iSMC

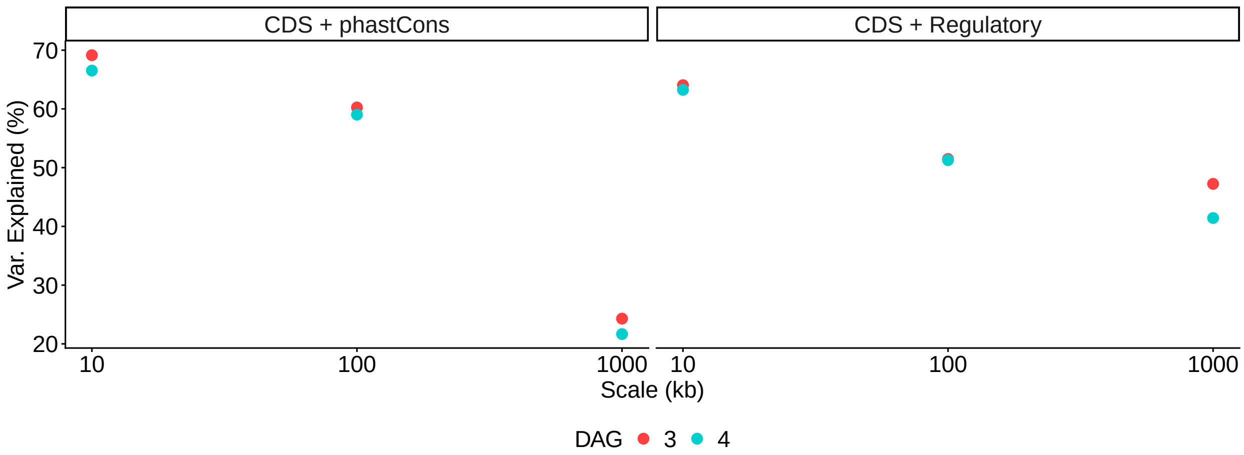

Figure S16: Variance in $R^2_\pi$ across chromosomes explained by a linear model using average chromosome features (GC%, mutation, recombination and proportion of deleterious sites) as log-transformed predictors. Y-axis shows the variance explained (across chromosomes) in the variance (across the $\pi$ landscape) per DAG (color).

maps, the model fit well ($p$-value=0.206, CFI=0.998), indicating that the empirical mutation maps reflect a common latent construct. For each autosome, we then fit measurement models using exclusively Roulette, Carlson and gnomAD as indicators, and reconstructed the latent $\mu$ from the estimated loadings ($\lambda$'s).

Another key advantage of structural equation models over traditional regression is the availability of rigorous tools for assessing goodness-of-fit and performing model selection (Shipley, 2016). Throughout the main text we use two complementary metrics for evaluating the goodness-of-fit: the $\chi^2$ test and the CFI.

The $\chi^2$ test contrasts the covariance matrix predicted by the fitted model with that observed in data, providing a comprehensive evaluation of causal structures hypothesized by the DAG. Because it tests the null hypothesis that the model is exactly compatible with the data, $\chi^2$ $p$-values are highly sensitive to sample size (number of genomic windows). Alternatively, fit measures such as the Comparative Fit Index (CFI) give a more practical assessment by comparing the specified model to a null (independence) model, with values above 0.90 typically treated as acceptable and above 0.95 indicating a good fit. CFI is less sensitive to sample size than the exact $\chi^2$ test, complementing it in evaluating model performance at different genomic scales, as wider windows contain less noisy estimates at the cost of smaller sample sizes.

#### 8.2 Why DAGs 3 and 4 yield higher squared correlation than DAGs 1 and 2

In the main text, we find that the more complex causal models (DAGs 3 and 4) explain, on average, more variation in diversity ($R^2_\pi$) than the simpler DAGs 1 and 2 (Figure S38). This happens despite DAGs 3 and 4 not including any additional direct parent of $\pi$ relative to DAG 2. However, by including recombination in the model, DAGs 3 and 4 explain covariance patterns better and allow variation to propagate between nodes differently. In other words, although $\pi$ experiences no direct effect from recombination, including $r$ can change the variances and covariances of $\pi$'s parents and the estimated path coefficients, so the modeled variance (and therefore $R^2_\pi$) reported by SEM can differ.

**Variance decomposition in SEM.** Let $\pi$ be an endogenous variable with parents $X_1, \ldots, X_k$ and residual $\varepsilon$. The structural equation for $\pi$ is

$$\pi = \sum_{i=1}^{k} b_i X_i + \varepsilon.$$

Its variance decomposes as

$$\mathrm{Var}(\pi) = \mathrm{Var}\Big(\sum_{i=1}^{k} b_i X_i\Big) + \mathrm{Var}(\varepsilon) + 2\,\mathrm{Cov}\Big(\sum_{i=1}^{k} b_i X_i,\ \varepsilon\Big).$$

Under the usual SEM assumption that residuals are uncorrelated with predictors, this simplifies to

$$\mathrm{Var}(\pi) = \mathrm{Var}\Big(\sum_{i=1}^{k} b_i X_i\Big) + \mathrm{Var}(\varepsilon).$$

The model $R^2$ for $\pi$ is

$$R_\pi^2 \;=\; \frac{\mathrm{Var}\big(\sum_{i=1}^{k} b_i X_i\big)}{\mathrm{Var}(\pi)}.$$

**Effect of adding a top-layer variable $r$.** Suppose $r$ affects some parent $X_j$ with coefficient $a_j$, so

$$X_j \;=\; a_j r \;+\; z_j,$$

where $z_j$ collects other causes and noise. Substituting into $\pi$'s equation gives

$$\pi = \sum_{i \neq j} b_i X_i + b_j(a_j r + z_j) + \varepsilon = \Big(\sum_{i \neq j} b_i X_i + b_j z_j\Big) + (b_j a_j)\, r + \varepsilon.$$

Thus $r$ contributes indirectly to $\pi$ via the term $(b_j a_j) r$. More generally, any change in $\mathrm{Var}(X_i)$ or $\mathrm{Cov}(X_i, X_{i'})$ affects $\mathrm{Var}\big(\sum b_i X_i\big)$ and therefore $R_\pi^2$.

**Indirect and total effects.** The indirect effect of $r$ on $\pi$ through the chain $r \to X_j \to \pi$ equals the product of path coefficients:

$$\mathrm{Indirect}_{r \to \pi} \;=\; a_j \cdot b_j.$$

If multiple directed paths exist from $r$ to $\pi$, the total effect is the sum of products along all directed paths; these effects determine how much of $\pi$'s variance becomes attributable (directly or indirectly) to $r$.

#### 8.3 Wright's rule and variance explained at different genomic scales

Wright's rule of path analysis states that every path connecting two variables contributes to their correlation. Each path contributes a value equal to the product of the coefficients along that route. The total correlation is the sum of all such products.

In our genomic SEMs, as we increase window sizes, the total variance explained in a given endogenous variable (e.g., $R_\pi^2$) can be seen as depending on two components. First, a biological component given by the path coefficients ($\beta$), which represent causal strengths. Second, a purely statistical component: the variance retained in the predictors at the given scale. This follows from Wright's rule:

$$\mathrm{Cov}(A, B) = \beta \cdot \mathrm{Var}(A)$$

For a single predictor $A \to B$, the path coefficient is

$$\beta_{BA} = \frac{\mathrm{Cov}(A, B)}{\mathrm{Var}(A)}.$$

If $A$ and $B$ are standardized (mean 0, variance 1), this reduces to

$$\beta_{BA} = r_{AB},$$

where $r_{AB}$ is the correlation between $A$ and $B$.

With two predictors $A$ and $C$ for $B$, the path coefficient for $A \to B$ is

$$\beta_{BA} = \frac{r_{AB} - r_{AC}\, r_{BC}}{1 - r_{AC}^2}.$$

In the general case with predictors $X_1, X_2, \ldots, X_k$, the vector of path coefficients is

$$\boldsymbol{\beta}_B = \mathbf{R}_{XX}^{-1}\, \mathbf{r}_{XB},$$

where $\mathbf{R}_{XX}$ is the correlation matrix among predictors and $\mathbf{r}_{XB}$ is the vector of correlations between each predictor and $B$.

Wright's rule of path analysis states that the covariance between any two variables equals the sum, over all distinct paths connecting them, of the product of coefficients along each path, with exactly one variance or covariance term included per route:

$$\mathrm{Cov}(X_i, X_j) \;=\; \sum_{p \in \mathcal{P}(i \to j)} \left( \prod_{\ell \in p} \beta_\ell \right) \sigma_{z(p)},$$

where:

- $\mathcal{P}(i \to j)$ is the set of all distinct compound paths connecting $X_i$ and $X_j$,
- $\beta_\ell$ are the directed path coefficients along path $p$,
- $\sigma_{z(p)}$ is the single variance or covariance term included in path $p$.

Summing all such terms yields Wright's decomposition of $\mathrm{Cov}(X_i, X_j)$ into compound path products with exactly one variance or covariance per path.

**Example: Chain $X \to Y \to Z$** For example, the covariance between $X$ and $Z$ is

$$\begin{aligned}
\mathrm{Cov}(X, Z) &= \mathrm{Cov}\big(X, \beta_{ZY} Y\big) \\
&= \beta_{ZY}\, \mathrm{Cov}(X, Y) \\
&= \beta_{ZY} \beta_{YX}\, \mathrm{Var}(X) \\
&= \beta_{ZY} \beta_{YX} \sigma_X^2,
\end{aligned}$$

corresponding to the single compound path $X \to Y \to Z$, with one variance term $\sigma_X^2$ and the product of coefficients $\beta_{ZY}\beta_{YX}$.

**Relation to $R^2$ of $Z$ (Single Predictor Chain)** The variance explained in $Z$ (its $R^2$) is the proportion of variance in $Z$ accounted for by its predictors:

$$R_Z^2 = \frac{\mathrm{Var}(\hat{Z})}{\mathrm{Var}(Z)}.$$

For the chain $X \to Y \to Z$:

$$\mathrm{Var}(\hat{Y}) = \beta_{YX}^2 \sigma_X^2,$$

$$\mathrm{Var}(\hat{Z}) = \beta_{ZY}^2\, \mathrm{Var}(\hat{Y}) = \beta_{ZY}^2 \beta_{YX}^2 \sigma_X^2.$$

Thus,

$$R_Z^2 = \frac{\beta_{ZY}^2 \beta_{YX}^2 \sigma_X^2}{\sigma_Z^2}.$$

**Multiple predictors causally affecting $Z$** Suppose $Z$ has two direct causal parents, $X$ and $Y$. Then

$$Z = \beta_{ZX} X + \beta_{ZY} Y + \varepsilon_Z,$$

where $\varepsilon_Z$ is the residual variance in $Z$.
The variance explained in $Z$ is

$$\mathrm{Var}(\hat{Z}) = \beta_{ZX}^2 \sigma_X^2 + \beta_{ZY}^2 \sigma_Y^2 + 2\beta_{ZX}\beta_{ZY}\,\mathrm{Cov}(X,Y).$$

Therefore,

$$R_Z^2 = \frac{\beta_{ZX}^2 \sigma_X^2 + \beta_{ZY}^2 \sigma_Y^2 + 2\beta_{ZX}\beta_{ZY}\,\mathrm{Cov}(X,Y)}{\sigma_Z^2}.$$

And the variance explained in $Z$ depends on biological (causal) and statistical components:

$$R_Z^2 \;\propto\; \underbrace{\beta_{ZX}^2, \beta_{ZY}^2, \beta_{ZX}\beta_{ZY}}_{\text{causal strengths}} \times \underbrace{\sigma_X^2, \sigma_Y^2, \mathrm{Cov}(X,Y)}_{\text{variance retained at scale}}\;.$$

In brief,

- Each predictor contributes according to its path coefficient squared times its variance.
- Shared variance between predictors contributes through the cross-term $2\beta_{ZX}\beta_{ZY}\,\mathrm{Cov}(X,Y)$.
- The total $R_Z^2$ depends jointly on biological causal strengths and retained statistical variance at each scale.

Trevor Cousins, Daniel Tabin, Nick Patterson, David Reich, and Arun Durvasula. Accurate inference of population history in the presence of background selection. *bioRxiv*, 2024.

Chenlu Di, Swetha Ramesh, Jason Ernst, and Kirk E Lohmueller. The landscape of fitness effects of putatively functional noncoding mutations in humans. *bioRxiv*, pages 2025–05, 2025.

Jason Ernst and Manolis Kellis. ChromHMM: automating chromatin-state discovery and characterization. *Nature Methods*, 9(3):215–216, 2012.

Jan Freudenberg, Mingyi Wang, Yaning Yang, and Wentian Li. Partial correlation analysis indicates causal relationships between GC-content, exon density and recombination rate in the human genome. *BMC Bioinformatics*, 10(Suppl 1):S66, 2009.

Kimberly J Gilbert, Fanny Pouyet, Laurent Excoffier, and Stephan Peischl. Transition from background selection to associative overdominance promotes diversity in regions of low recombination. *Current Biology*, 30(1):101–107, 2020.

Paul Gustafson. The impact of unacknowledged measurement error. In *Handbook of measurement error models*, pages 37–52. Chapman and Hall/CRC, 2021.

Ryan N Gutenkunst, Ryan D Hernandez, Scott H Williamson, and Carlos D Bustamante. Inferring the joint demographic history of multiple populations from multidimensional snp frequency data. *PLoS Genetics*, 5 (10):e1000695, 2009.

Bjarni V Halldorsson, Gunnar Palsson, Olafur A Stefansson, Hakon Jonsson, Marteinn T Hardarson, Hannes P Eggertsson, Bjarni Gunnarsson, Asmundur Oddsson, Gisli H Halldorsson, Florian Zink, et al. Characterizing mutagenic effects of recombination through a sequence-level genetic map. *Science*, 363(6425):eaau1043, 2019.

Christian D Huber, Bernard Y Kim, Clare D Marsden, and Kirk E Lohmueller. Determining the factors driving selective effects of new nonsynonymous mutations. *Proceedings of the National Academy of Sciences*, 114(17):4465–4470, 2017.

John A Kamm, Jeffrey P Spence, Jeffrey Chan, and Yun S Song. Two-locus likelihoods under variable population size and fine-scale recombination rate estimation. *Genetics*, 203(3):1381–1399, 2016.

Konrad J Karczewski, Laurent C Francioli, Grace Tiao, Beryl B Cummings, Jessica Alföldi, Qingbo Wang, Ryan L Collins, Kristen M Laricchia, Andrea Ganna, Daniel P Birnbaum, et al. The mutational constraint spectrum quantified from variation in 141,456 humans. *Nature*, 581(7809):434–443, 2020.

Jerome Kelleher, Alison M Etheridge, and Gilean McVean. Efficient coalescent simulation and genealogical analysis for large sample sizes. *PLoS computational biology*, 12(5):e1004842, 2016.

Bernard Y Kim, Christian D Huber, and Kirk E Lohmueller. Inference of the distribution of selection coefficients for new nonsynonymous mutations using large samples. *Genetics*, 206(1):345–361, 2017.

Michael Lawrence, Robert Gentleman, and Vincent Carey. rtracklayer: an r package for interfacing with genome browsers. *Bioinformatics*, 25(14):1841, 2009.

Kirk E Lohmueller, Thomas Sparsø, Qibin Li, Ehm Andersson, Thorfinn Korneliussen, Anders Albrechtsen, Karina Banasik, Niels Grarup, Ingileif Hallgrimsdottir, Kristoffer Kiil, et al. Whole-exome sequencing of 2,000 danish individuals and the role of rare coding variants in type 2 diabetes. *The American Journal of Human Genetics*, 93(6):1072–1086, 2013.

David A Murphy, Eyal Elyashiv, Guy Amster, and Guy Sella. Broad-scale variation in human genetic diversity levels is predicted by purifying selection on coding and non-coding elements. *Elife*, 12:e76065, 2022.

Aaron P Ragsdale. Local fitness and epistatic effects lead to distinct patterns of linkage disequilibrium in protein-coding genes. *Genetics*, 221(4):iyac097, 2022.

Jonathan Romiguier, Vincent Ranwez, Emmanuel JP Douzery, and Nicolas Galtier. Contrasting gc-content dynamics across 33 mammalian genomes: relationship with life-history traits and chromosome sizes. *Genome Research*, 20(8):1001–1009, 2010.

Enrique Santiago and Armando Caballero. Joint prediction of the effective population size and the rate of fixation of deleterious mutations. *Genetics*, 204(3):1267–1279, 2016.

Vladimir Seplyarskiy, Evan M Koch, Daniel J Lee, Joshua S Lichtman, Harding H Luan, and Shamil R Sunyaev. A mutation rate model at the basepair resolution identifies the mutagenic effect of polymerase III transcription. *Nature Genetics*, 55(12):2235–2242, 2023.

Bill Shipley. *Cause and correlation in biology: A user's guide to path analysis, structural equations and causal inference with R*. Cambridge university press, 2016.

Adam Siepel, Gill Bejerano, Jakob S Pedersen, Angie S Hinrichs, Minmei Hou, Kate Rosenbloom, Hiram Clawson, John Spieth, LaDeana W Hillier, Stephen Richards, et al. Evolutionarily conserved elements in vertebrate, insect, worm, and yeast genomes. *Genome research*, 15(8):1034–1050, 2005.

Jeffrey P Spence and Yun S Song. Inference and analysis of population-specific fine-scale recombination maps across 26 diverse human populations. *Science Advances*, 5(10):eaaw9206, 2019.

Kevin R Thornton. A c++ template library for efficient forward-time population genetic simulation of large populations. *Genetics*, 198(1):157–166, 2014.

Ha Vu and Jason Ernst. Universal annotation of the human genome through integration of over a thousand epigenomic datasets. *Genome Biology*, 23(1):9, 2022.

John Wakeley. *Coalescent Theory: An Introduction*. Roberts and Company Publishers, 2009.

Brooke Weinstein and Scott William Roy. Controlling for life-history traits in vertebrates reveals that effective population size does not affect mutation rate or genome size. *Proceedings of the National Academy of Sciences*, 123(6):e2519649123, 2026.

Tony Zeng, Jeffrey P Spence, Hakhamanesh Mostafavi, and Jonathan K Pritchard. Bayesian estimation of gene constraint from an evolutionary model with gene features. *Nature Genetics*, 56(8):1632–1643, 2024.

### 9 Supplemental Tables

Table S4: DFE parameter estimates for constrained elements included in *CDS+phastCons* and *CDS+regulatory* models. When exons are divided into categories, the mean strength of selection against missense mutations increases with the intolerance to nonsense mutations (as expected).

| **Name** | **Type** | **Shape** | **Scale** | $p_{\text{neu}}$ | $s_{\text{mean}}$ |
|---|---|---|---|---|---|
| **CDS** | | | | | |
| exons | gamma_neutral | 2.15e-1 | 2.40e-2 | 3.02e-1 | 3.60e-3 |
| exons_1 | gamma_neutral | 2.95e-1 | 2.06e-1 | 2.78e-1 | 4.39e-2 |
| exons_2 | gamma_neutral | 2.56e-1 | 1.69e-1 | 2.83e-1 | 3.10e-2 |
| exons_3 | gamma_neutral | 2.42e-1 | 8.50e-2 | 2.83e-1 | 1.48e-2 |
| exons_4 | gamma_neutral | 2.12e-1 | 8.54e-2 | 2.82e-1 | 1.30e-2 |
| exons_5 | gamma_neutral | 1.63e-1 | 1.38e-1 | 2.79e-1 | 1.62e-2 |
| exons_6 | gamma_neutral | 1.66e-1 | 2.65e-2 | 2.77e-1 | 3.19e-3 |
| exons_7 | gamma_neutral | 1.02e-1 | 1.10e-1 | 2.78e-1 | 8.10e-3 |
| exons_8 | gamma_neutral | 1.09e-1 | 4.49e-2 | 2.78e-1 | 3.54e-3 |
| exons_9 | gamma_neutral | 1.18e-1 | 2.06e-2 | 2.80e-1 | 1.74e-3 |
| exons_10 | gamma_neutral | 1.36e-1 | 8.74e-3 | 2.80e-1 | 8.58e-4 |
| exons_11 | gamma_neutral | 8.93e-2 | 5.60e-2 | 2.89e-1 | 3.56e-3 |
| **Regulatory** | | | | | |
| promoters | gamma | 1.11e-1 | 5.00e-4 | 0.0 | 5.54e-5 |
| enhancers | gamma | 1.01e-2 | 2.27e-1 | 0.0 | 2.28e-3 |
| **Non-coding phastCons (0–60%)** | | | | | |
| phastCons_0–5 | gamma | 1.69e-1 | 1.78e-3 | 0.0 | 3.01e-4 |
| phastCons_5–10 | gamma | 1.27e-1 | 1.15e-3 | 0.0 | 1.46e-4 |
| phastCons_10–15 | gamma | 1.13e-1 | 1.05e-3 | 0.0 | 1.19e-4 |
| phastCons_15–20 | gamma | 9.23e-2 | 1.36e-3 | 0.0 | 1.25e-4 |
| phastCons_20–25 | gamma | 8.12e-2 | 1.39e-3 | 0.0 | 1.13e-4 |
| phastCons_25–30 | gamma | 7.26e-2 | 1.52e-3 | 0.0 | 1.11e-4 |
| phastCons_30–35 | gamma | 5.43e-2 | 2.91e-3 | 0.0 | 1.58e-4 |
| phastCons_35–40 | gamma | 4.77e-2 | 3.28e-3 | 0.0 | 1.56e-4 |
| phastCons_40–45 | gamma | 5.01e-2 | 2.29e-3 | 0.0 | 1.15e-4 |
| phastCons_45–50 | gamma | 3.85e-2 | 4.25e-3 | 0.0 | 1.64e-4 |
| phastCons_50–55 | gamma | 3.42e-2 | 4.35e-3 | 0.0 | 1.49e-4 |
| phastCons_55–60 | gamma | 3.05e-2 | 4.52e-3 | 0.0 | 1.38e-4 |

Table S5: Likelihood ratio tests comparing genome-wide and chromosome-specific models of diversity.

| **Constraint model** | **DAG** | **Scale** | **P-value** |
|---|---|---|---|
| split *CDS+phastCons* | 1 | 10 kb | $< 1 \times 10^{-300}$ |
| split *CDS+regulatory* | 1 | 10 kb | $< 1 \times 10^{-300}$ |
| split *CDS+phastCons* | 2 | 10 kb | $< 1 \times 10^{-300}$ |
| split *CDS+regulatory* | 2 | 10 kb | $< 1 \times 10^{-300}$ |
| split *CDS+phastCons* | 3 | 10 kb | $< 1 \times 10^{-300}$ |
| split *CDS+regulatory* | 3 | 10 kb | $< 1 \times 10^{-300}$ |
| split *CDS+phastCons* | 4 | 10 kb | $< 1 \times 10^{-300}$ |
| split *CDS+regulatory* | 4 | 10 kb | $< 1 \times 10^{-300}$ |
| split *CDS+phastCons* | 1 | 100 kb | $4.096\,788 \times 10^{-108}$ |
| split *CDS+regulatory* | 1 | 100 kb | $4.757\,204 \times 10^{-158}$ |
| split *CDS+phastCons* | 2 | 100 kb | $5.378\,837 \times 10^{-153}$ |
| split *CDS+regulatory* | 2 | 100 kb | $1.019\,143 \times 10^{-211}$ |
| split *CDS+phastCons* | 3 | 100 kb | $1.516\,175 \times 10^{-265}$ |
| split *CDS+regulatory* | 3 | 100 kb | $4.850\,423 \times 10^{-210}$ |
| split *CDS+phastCons* | 4 | 100 kb | $2.138\,158 \times 10^{-265}$ |
| split *CDS+regulatory* | 4 | 100 kb | $9.044\,428 \times 10^{-214}$ |
| split *CDS+phastCons* | 1 | 1 Mb | $4.029\,012 \times 10^{-14}$ |
| split *CDS+regulatory* | 1 | 1 Mb | $1.502\,838 \times 10^{-14}$ |
| split *CDS+phastCons* | 2 | 1 Mb | $1.537\,601 \times 10^{-21}$ |
| split *CDS+regulatory* | 2 | 1 Mb | $6.750\,590 \times 10^{-14}$ |
| split *CDS+phastCons* | 3 | 1 Mb | $1.061\,088 \times 10^{-48}$ |
| split *CDS+regulatory* | 3 | 1 Mb | $6.432\,885 \times 10^{-23}$ |
| split *CDS+phastCons* | 4 | 1 Mb | $2.278\,674 \times 10^{-49}$ |
| split *CDS+regulatory* | 4 | 1 Mb | $3.124\,356 \times 10^{-24}$ |

Note: Extremely small p-values originally reported as 0 are displayed as $< 1 \times 10^{-300}$ for clarity.

Table S6: Autosome-wide unlinked $B$ estimates across constraint and mutation models.

| **Constraint model** | **Mutation model** | **Autosome-wide** $B_{\text{unlinked}}$ |
|---|---|---|
| merged *CDS+regulatory* | Carlson | 0.943 |
| | gnomAD | 0.952 |
| | Roulette | 0.952 |
| split *CDS+regulatory* | Carlson | 0.911 |
| | gnomAD | 0.920 |
| | Roulette | 0.922 |
| merged *CDS+phastCons* | Carlson | 0.974 |
| | gnomAD | 0.977 |
| | Roulette | 0.977 |
| split *CDS+phastCons* | Carlson | 0.940 |
| | gnomAD | 0.944 |
| | Roulette | 0.947 |

Table S7: Autosome-wide log-likelihoods. $\Delta LL$ for Carlson models is calculated separately from Roulette and gnomAD $LL$, as the Carlson map has different genomic coverage than the Roulette and gnomAD maps.

| **Mutation model** | **Constraint model** | **Accessible sites** | $LL$ | $\Delta LL$ |
|---|---|---|---|---|
| Carlson | merged *CDS+regulatory* | 1,968,399,089 | $-3.396618 \times 10^{11}$ | $5.557216 \times 10^{8}$ |
| | split *CDS+regulatory* | 1,968,399,089 | $-3.396689 \times 10^{11}$ | $5.627897 \times 10^{8}$ |
| | merged *CDS+phastCons* | 1,968,399,089 | $-3.391252 \times 10^{11}$ | $1.912630 \times 10^{7}$ |
| | split *CDS+phastCons* | 1,968,399,089 | $-3.391061 \times 10^{11}$ | 0 |
| gnomAD | merged *CDS+regulatory* | 2,164,673,425 | $-3.840134 \times 10^{11}$ | $1.378994 \times 10^{9}$ |
| | split *CDS+regulatory* | 2,164,673,425 | $-3.840302 \times 10^{11}$ | $1.395750 \times 10^{9}$ |
| | merged *CDS+phastCons* | 2,164,673,425 | $-3.831838 \times 10^{11}$ | $5.494125 \times 10^{8}$ |
| | split *CDS+phastCons* | 2,164,673,425 | $-3.831728 \times 10^{11}$ | $5.384178 \times 10^{8}$ |
| Roulette | merged *CDS+regulatory* | 2,164,673,425 | $-3.831660 \times 10^{11}$ | $5.316324 \times 10^{8}$ |
| | split *CDS+regulatory* | 2,164,673,425 | $-3.831732 \times 10^{11}$ | $5.388178 \times 10^{8}$ |
| | merged *CDS+phastCons* | 2,164,673,425 | $-3.826512 \times 10^{11}$ | $1.682158 \times 10^{7}$ |
| | split *CDS+phastCons* | 2,164,673,425 | $-3.826344 \times 10^{11}$ | 0 |

### 10 Supplemental Figures

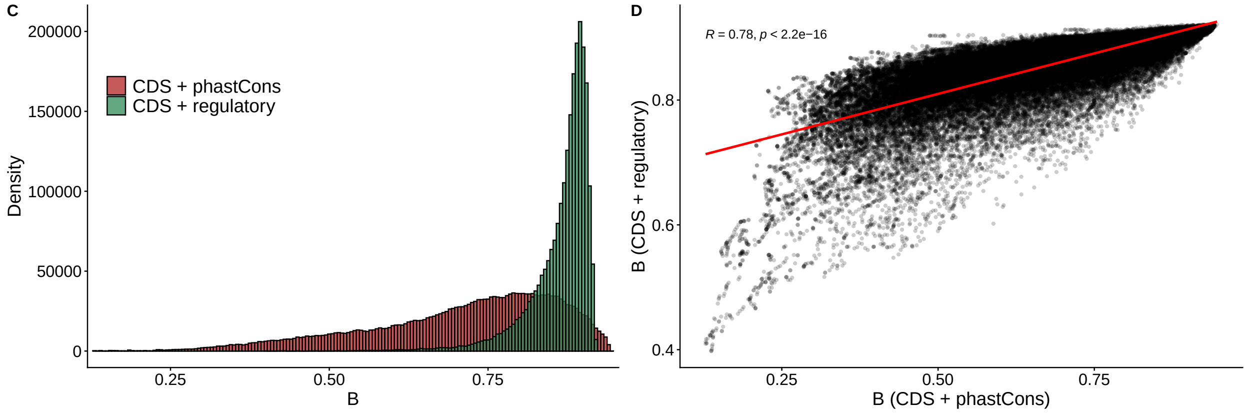

Figure S17: Fine-scale $B$-values according to *CDS+phastCons* and *CDS+regulatory* elements. Here 1 kb predictions are shown for models where the exonic DFE is split into deciles. **A:** histograms of $B$-value distributions. **B:** relationship between $B$-values predicted under both groups of elements (down-sampled to 5%), with the best fit linear model in red.

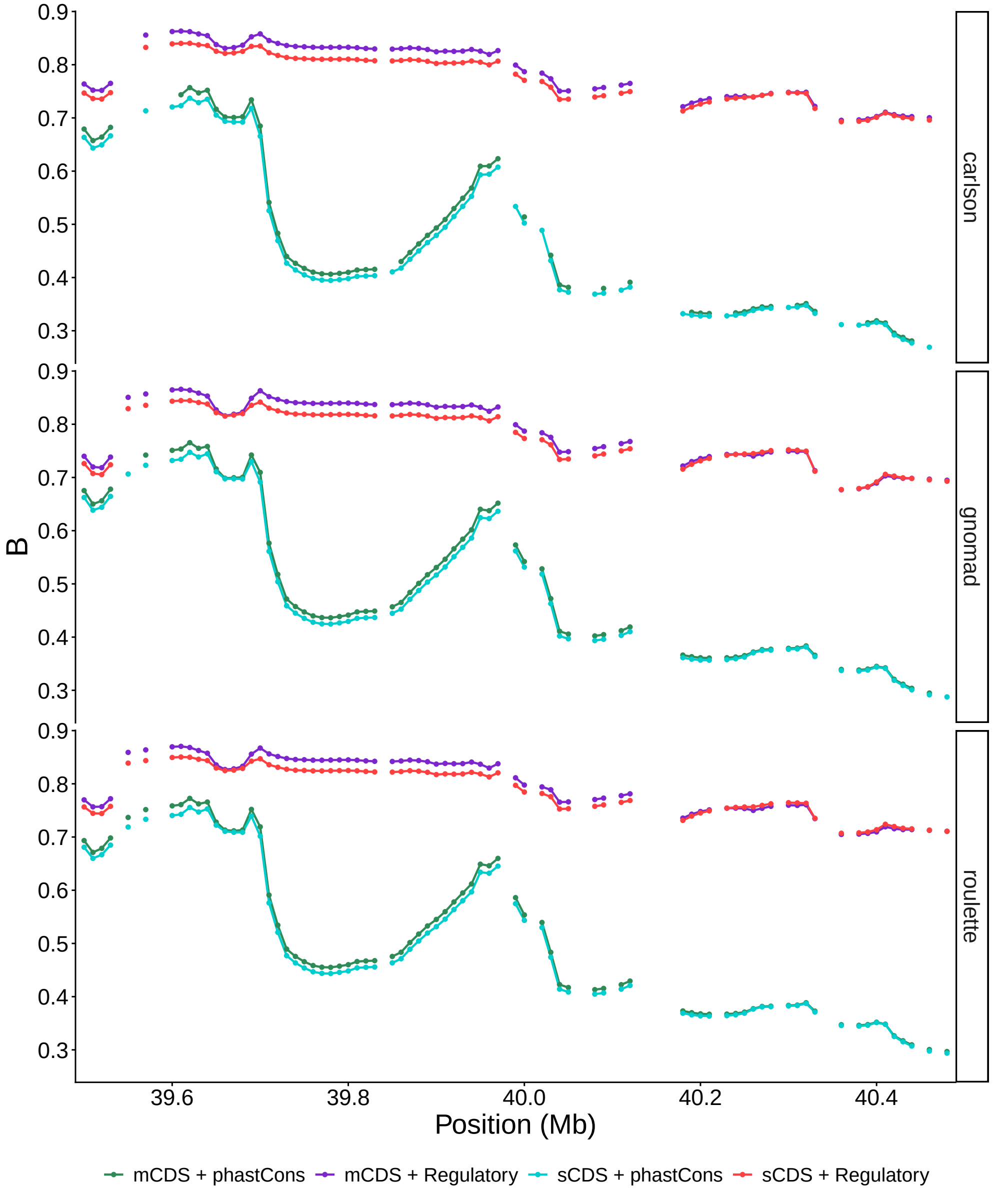

Figure S18: Predicted $B$-values across mutation maps (panels) and groups of elements (color). To increase resolution, 1 kb predictions are shown for a small segment of chromosome 22. Green: *CDS+phastCons* elements with exons merged under the Kim et al. (2017) DFE. Purple: *CDS+regulatory* elements with exons merged under the Kim et al. (2017) DFE. Cyan: *CDS+phastCons* elements with exons split into deciles of constraint. Red: *CDS+regulatory* elements with exons split into deciles of constraint. $B$-values are nearly invariant to the mutation map and to discretization of the exonic DFE.

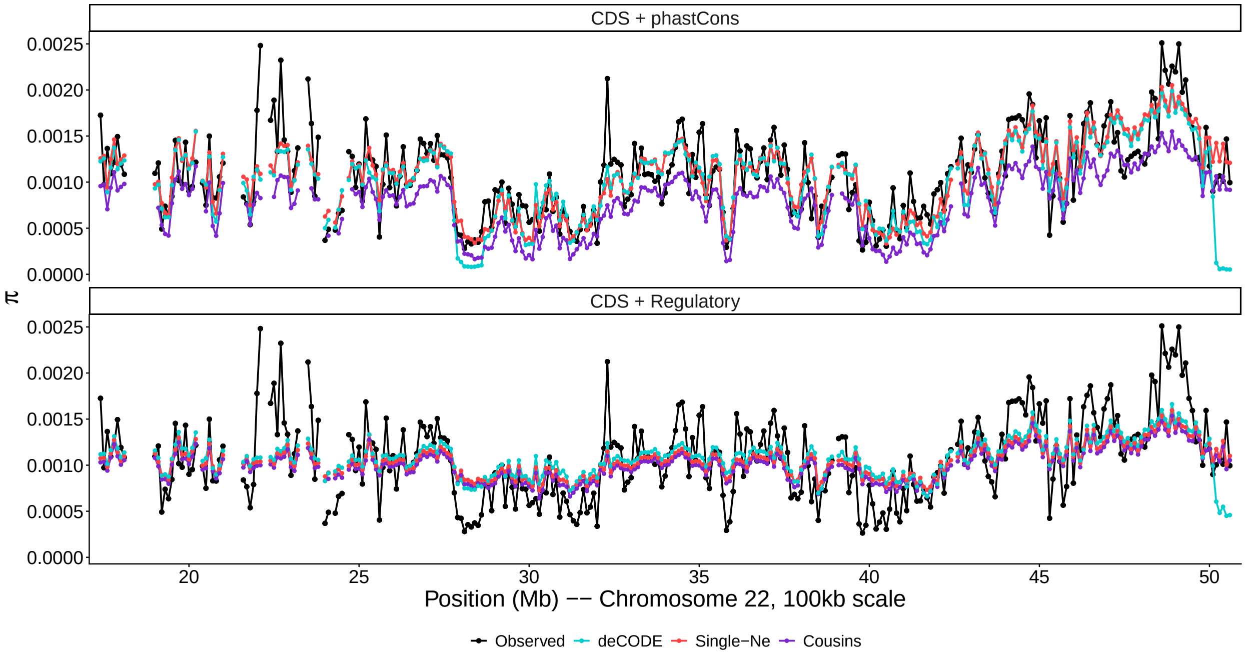

Figure S19: Observed ($\hat{\pi}$, black) and predicted ($\mathbb{E}[\pi] = B \times \mathbb{E}[\pi]$, color) diversity landscapes under alternative models, here shown for the Roulette mutation map in chromosome 22 (100 kb scale). Red: $B$-values assuming the genome-wide average estimate $N_e$ across autosomes. Purple: $B$-value assuming the Cousins et al. (2024) demography instead of the fit $N_e$. Light blue: $B$-value assuming the deCODE recombination map and its corresponding chromosome-specific $N_e$ estimates (Figure S58). Note that deCODE estimates are typically zero at the tips of the chromosomes, predicting very low $B$-values in some cases. Although non-equilibrium models have no free parameters, they still capture the absolute scale of the diversity landscape reasonably well. Here the exonic DFE is split into deciles of constraint.

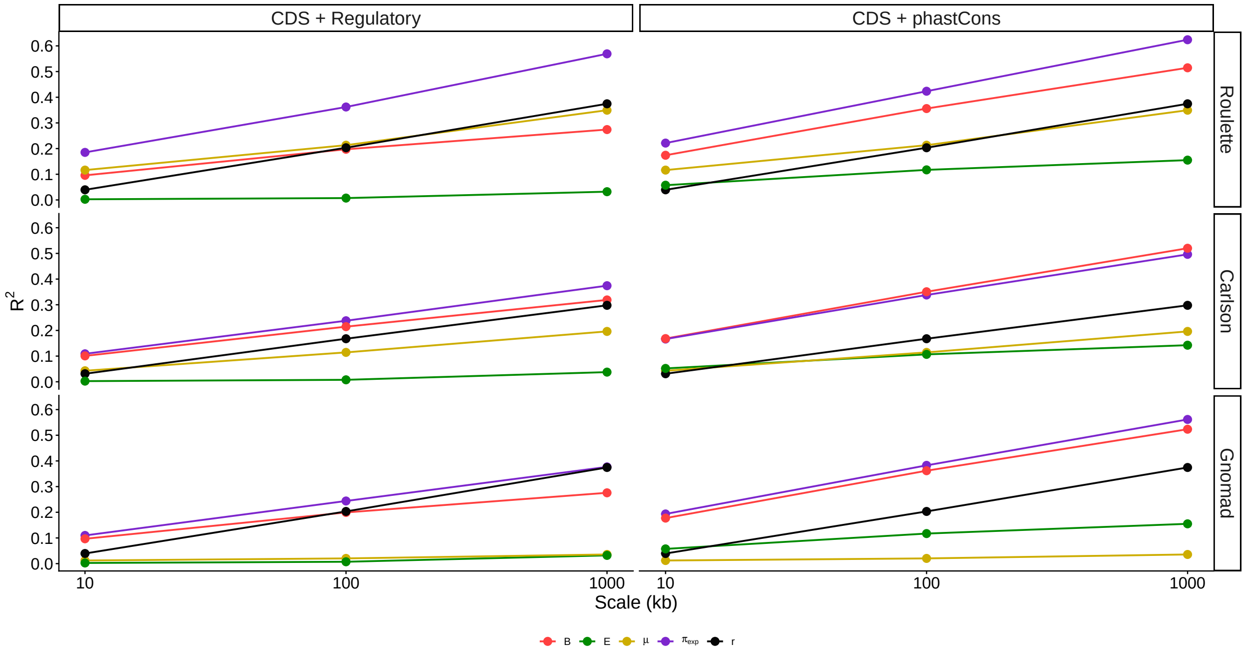

Figure S20: Genome-wide squared correlations between $\pi$ and individual landscapes (color). Previous studies focused on either $B$ (red) or $\mu$ (yellow). In purple, $\pi_{exp} = B \times \pi_0$, with both variables influenced by mutation rate variation. The density of constrained elements ($E$, green) is the ratio of the count of constrained and callable sites in each window. Squared correlations for $r$ (black) and $E$, which are both independent of mutation, differ slightly for Carlson map due to its distinct coverage. The raw predictive power of $E$ is much higher in phastCons than in genes. These squared correlations cannot be interpreted as the proportion of variance in $\pi$ explained by each predictor, as their sum can exceed 1. See Figures S37 and S23 for chromosome-specific correlations.

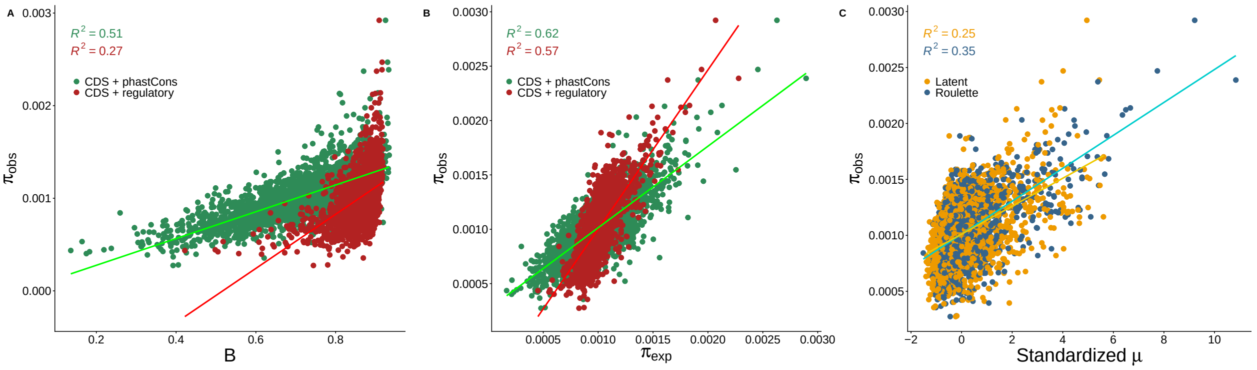

Figure S21: Relationship between genome-wide observed diversity (1 Mb scale) and either $B$-values (**A**), $\mathbb{E}[\pi] = B \times \mathbb{E}[\pi_0]$, with $\pi_0$ incorporating the Roulette map (**B**), or mutation rates (**C**). $R^2$ values denote the squared Pearson correlation per group.

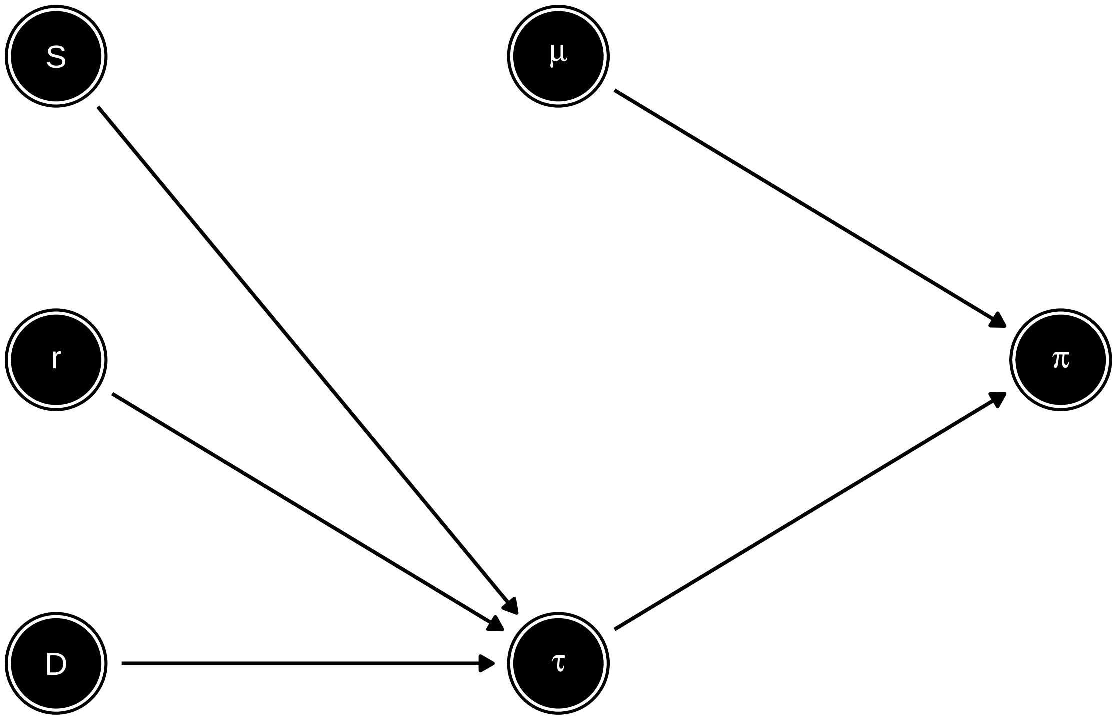

Figure S22: Causal model proposed by Barroso and Dutheil (2023). Since the impact of selection (S), recombination (r) and genetic drift (D) on diversity ($\pi$) is mediated by the average coalescente times ($\tau$), they fit this model to *Drosophila melanogaster* data using multiple linear regression with $\tau$ and mutation rate ($\mu$) as predictors.

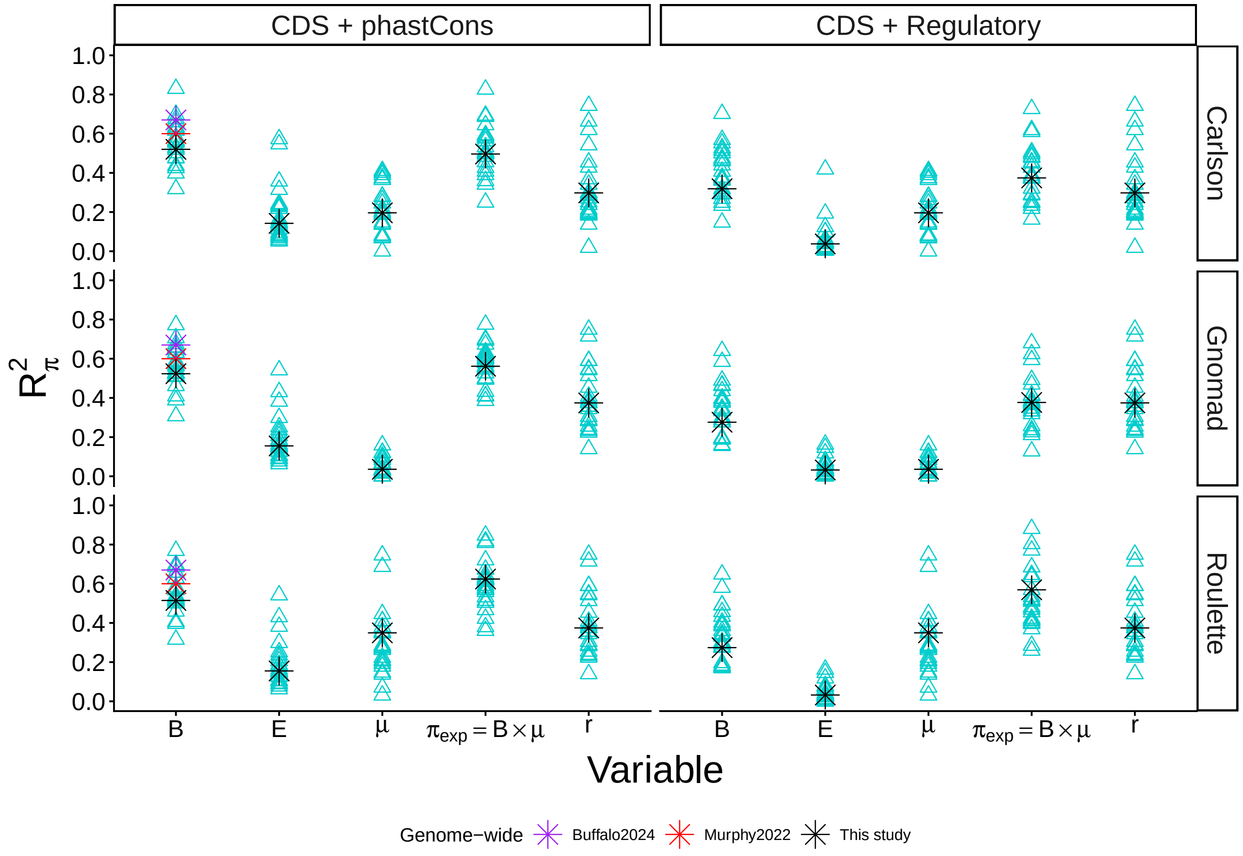

Figure S23: Squared correlations between $\pi$ and individual landscapes at the megabase scale. Blue triangles show distributions across human autosomes whereas black stars show genome-wide estimates. As reference points, red and purple stars show results obtained by Murphy et al. (2022) and Buffalo and Kern (2024), respectively, although they do not incorporate mutation maps and use different genome annotations. $\pi_{exp} = B \times \pi_0$, with both variables influenced by mutation rate variation. The density of constrained elements ($E$) is the ratio of the count of constrained and callable sites in each window. Results using $r$ and $E$,which are independent from mutation, differ slightly for Carlson map due to its distinct coverage. See Figure S37 for more details.

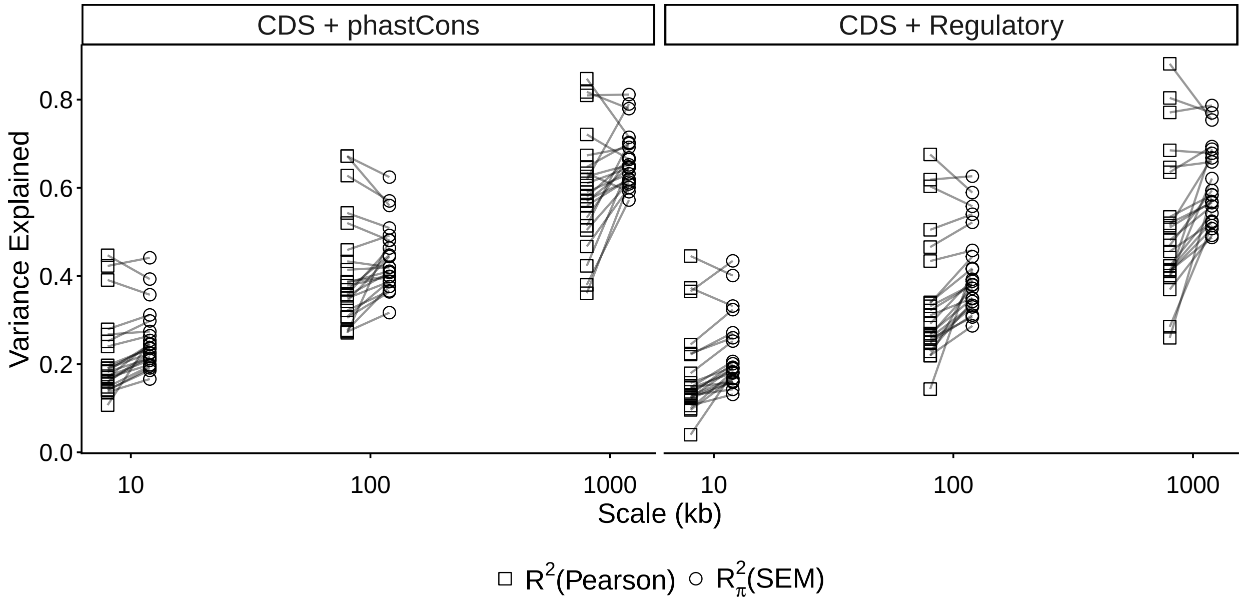

Figure S24: Comparison between measures of variance in diversity that is explained by each model. $R^2$: squared Pearson correlation between $\pi_{obs}$ and $\mathbb{E}[\pi] = B \times \mathbb{E}[\pi_0]$. $R^2_\pi$: estimate of the total variance in diversity explained by DAG 4. Shown are (paired) results per chromosome and scale.

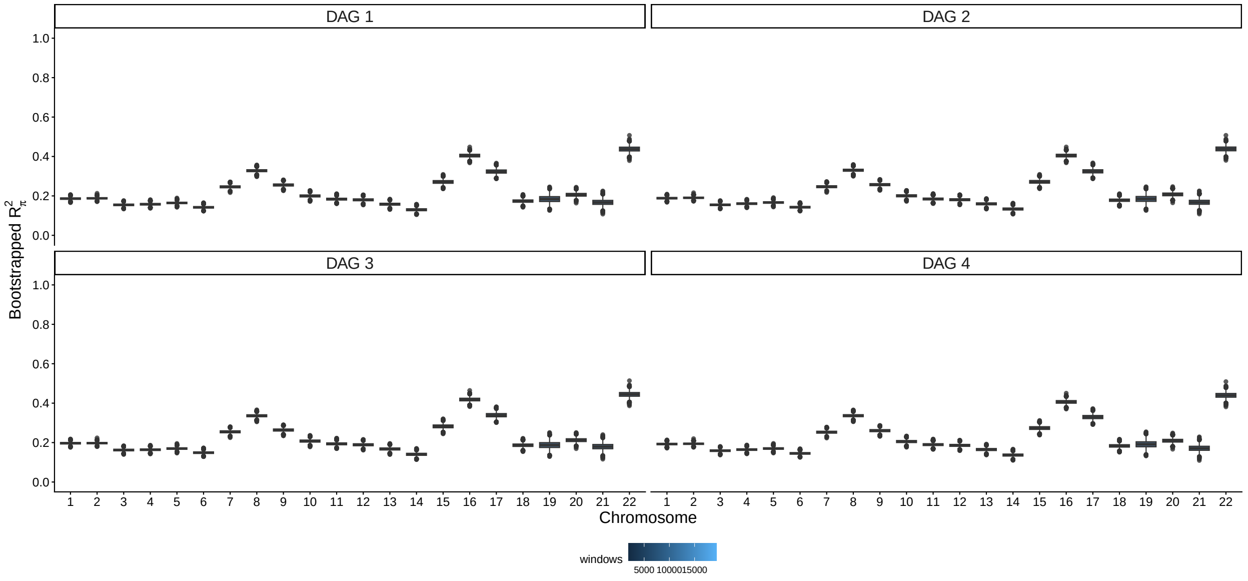

Figure S25: Distribution of $R^2_\pi$ after 5,000 bootstrap replicates of SEM, here shown for the *phastCons* model at 10 kb scale. Color key shows number of windows after filtering for minimum of 75% coverage.

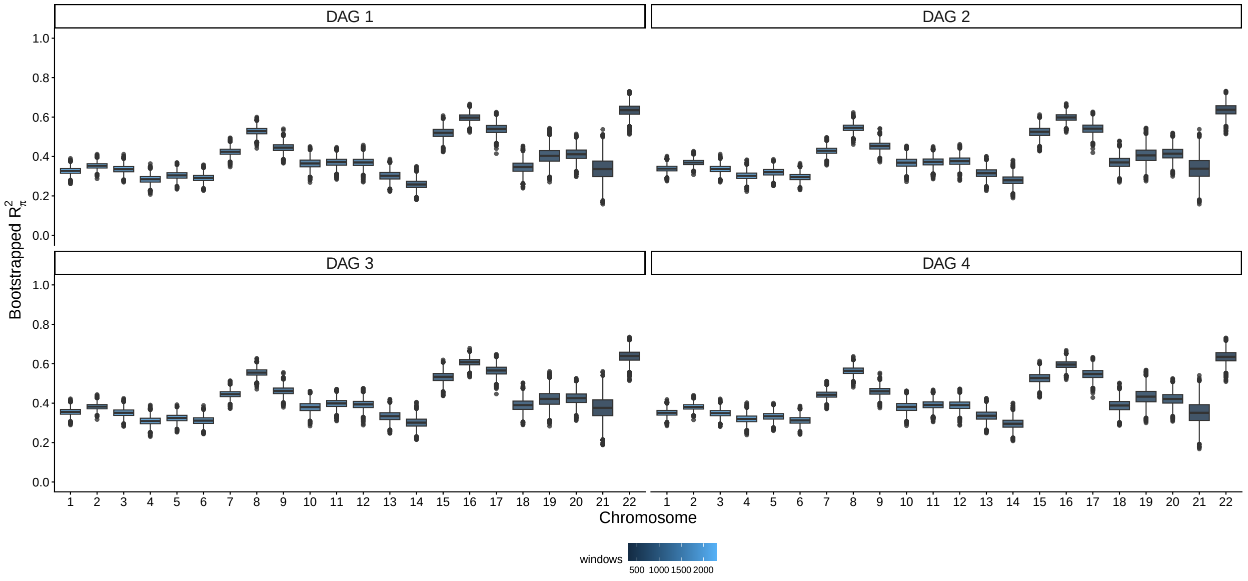

Figure S26: Distribution of $R_\pi^2$ after 5,000 bootstrap replicates of SEM, here shown for the *phastCons* model at 100 kb scale. Color key shows number of windows after filtering for minimum of 50% coverage.

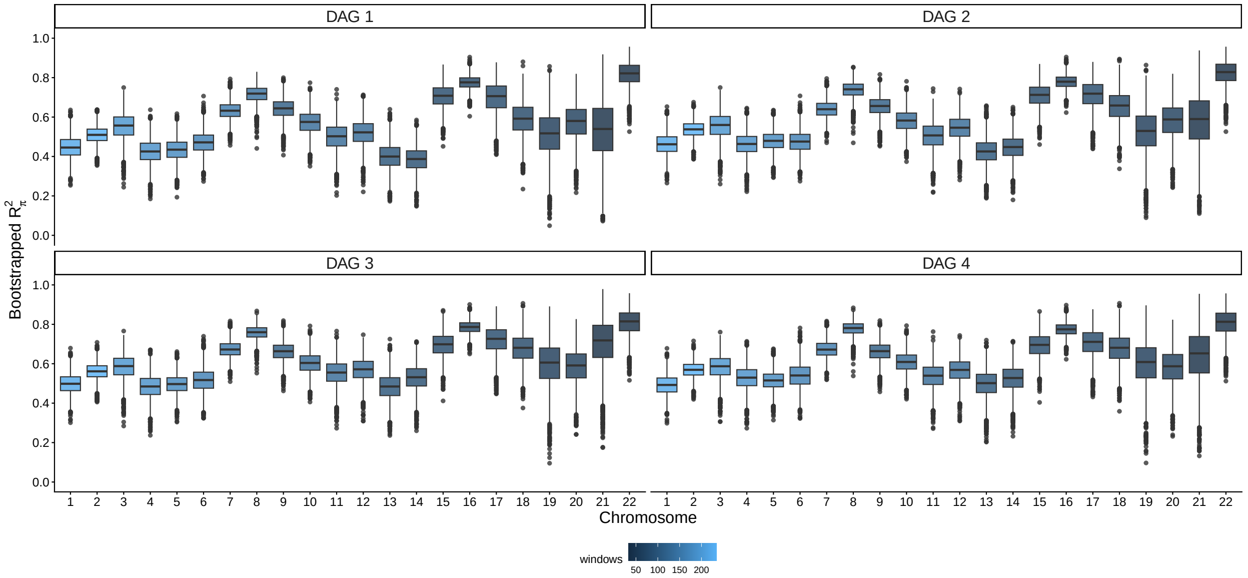

Figure S27: Distribution of $R_\pi^2$ after 5,000 bootstrap replicates of SEM, here shown for the *phastCons* model at 1 Mb scale. Color key shows number of windows after filtering for minimum of 20% coverage.

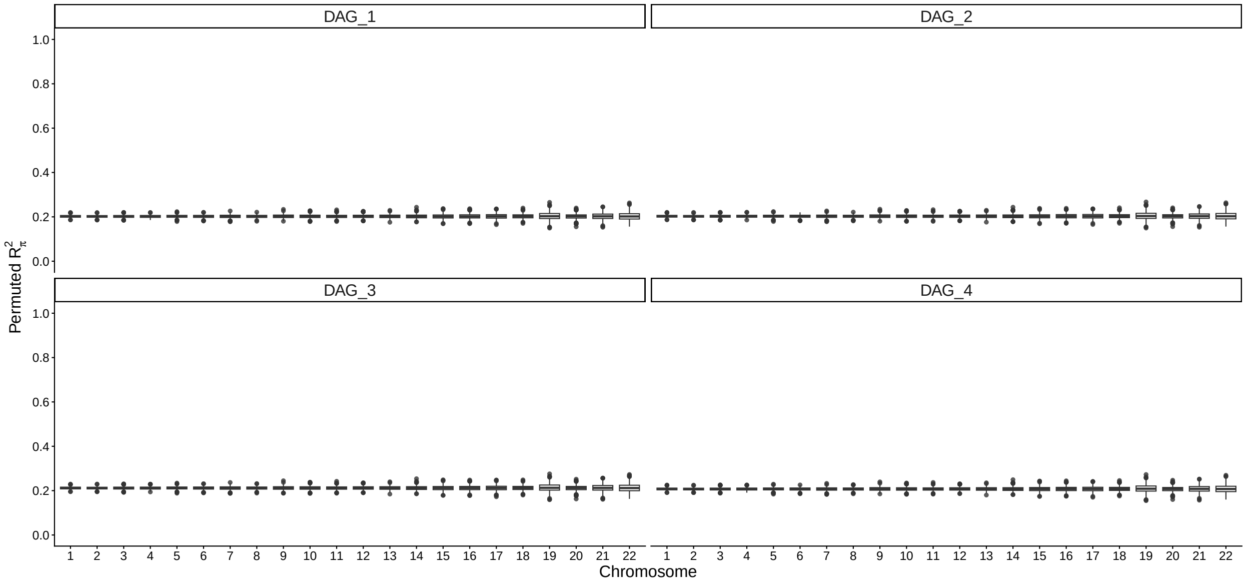

Figure S28: Distribution of $R^2_\pi$ after 1,000 permutations of chromosome labels, here shown for the *phastCons* model at 10 kb scale.

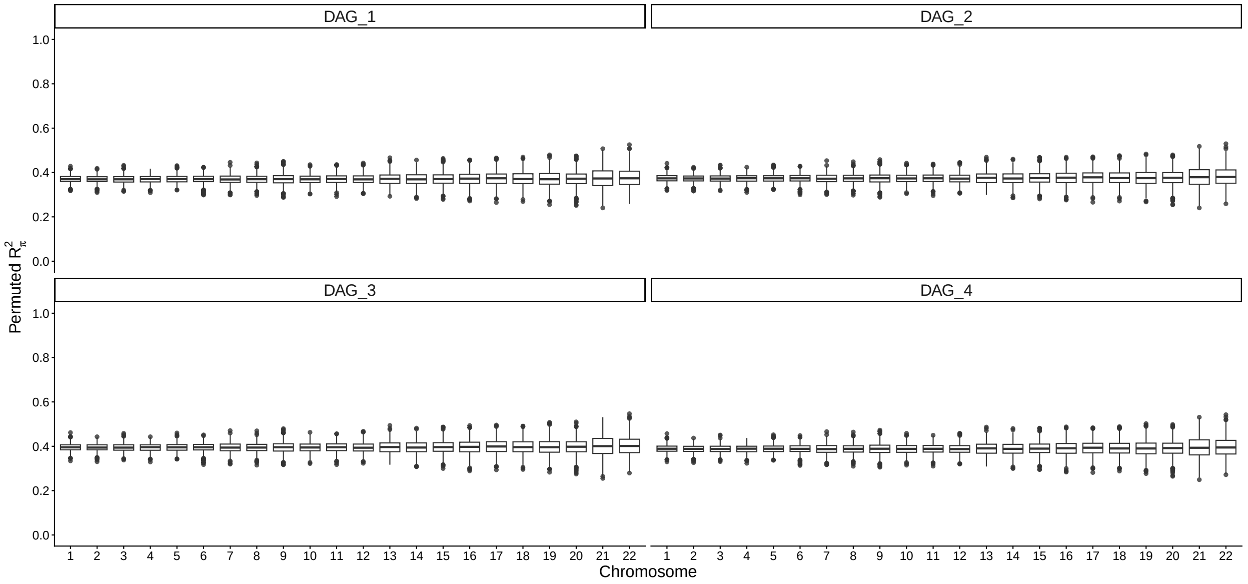

Figure S29: Distribution of $R^2_\pi$ after 1,000 permutations of chromosome labels, here shown for the *phastCons* model at 100 kb scale.

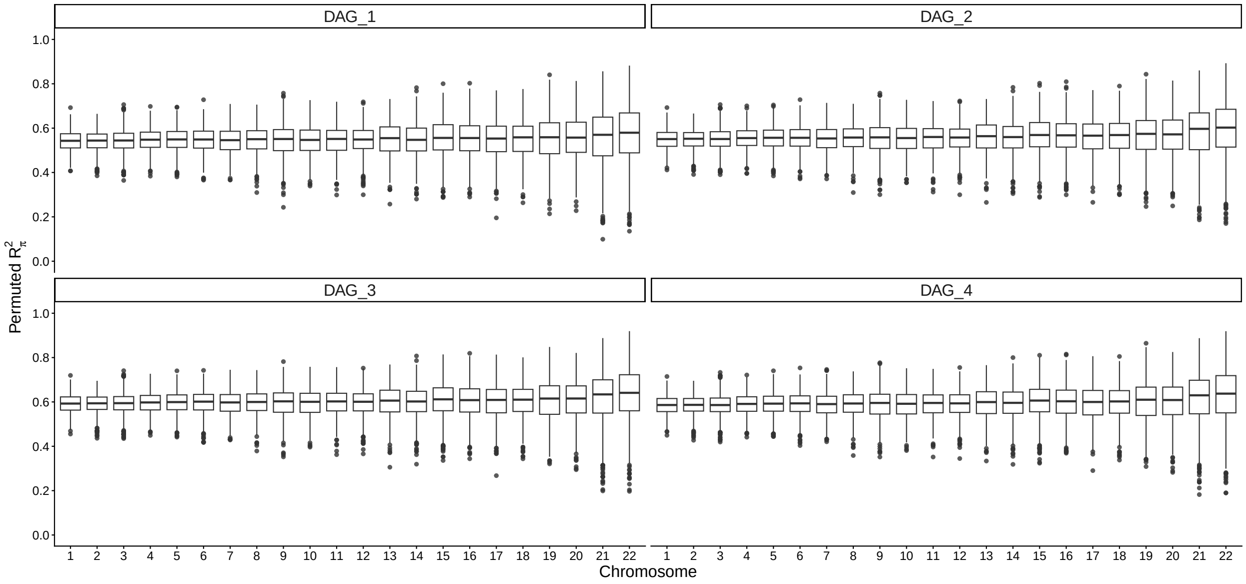

Figure S30: Distribution of $R^2_\pi$ after 1,000 permutations of chromosome labels, here shown for the *phastCons* model at 1 Mb scale.

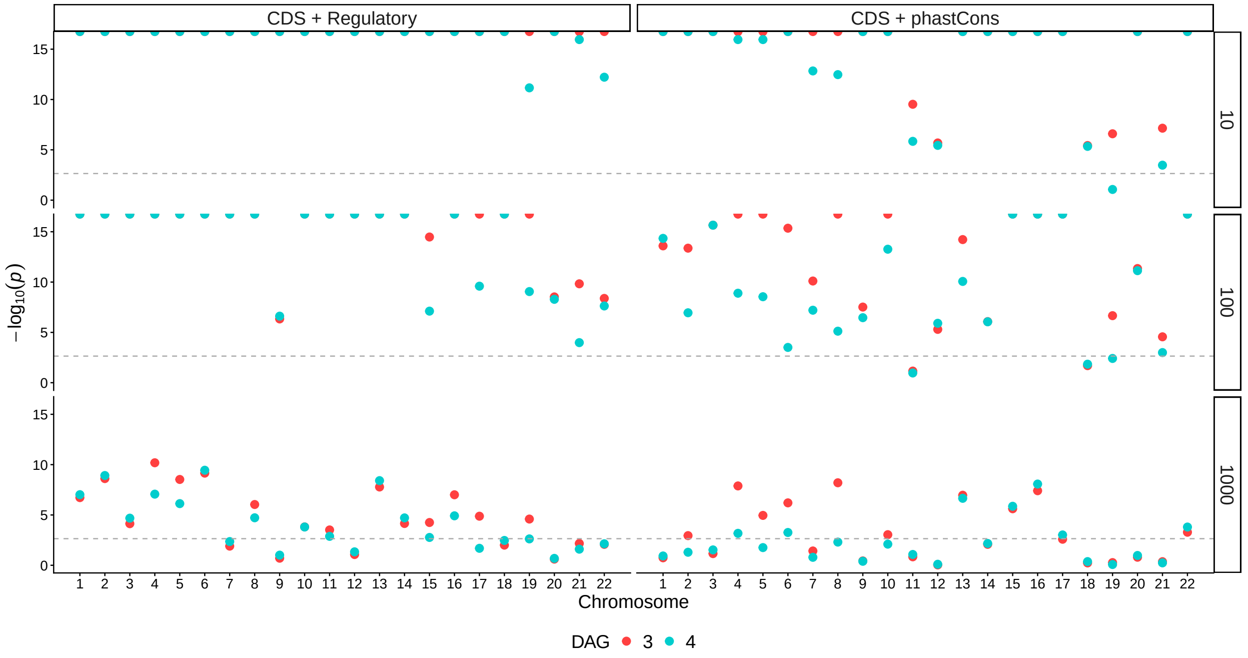

Figure S31: DAGs 3 and 4 are broadly compatible with human data (part I). $\chi^2$ $p$-values testing the fit of the DAGs in each autosome, for different functional elements (column panels) and window sizes (kb, row panels). Dashed grey lines show significance threshold after Bonferroni correction over chromosomes (0.05/22).

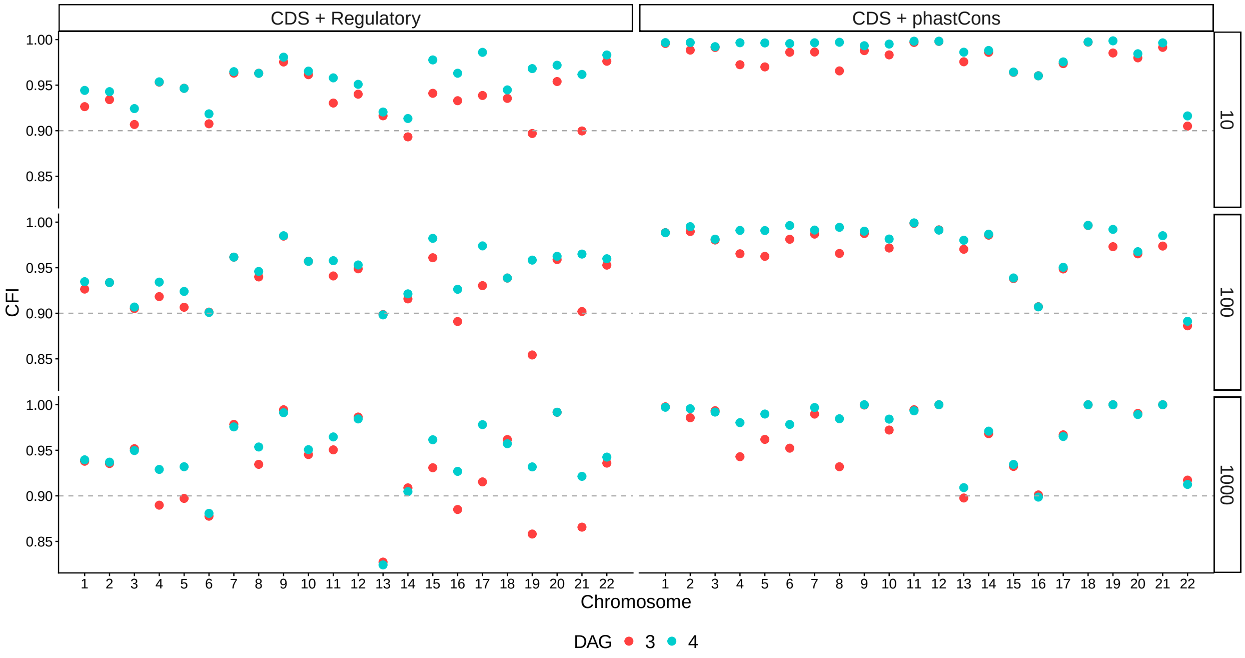

Figure S32: DAGs 3 and 4 are broadly compatible with human data (part II). Comparative Fit Indices for each chromosome, different functional elements (column panels) and window sizes (kb, row panels). Dashed grey lines show the rule-of-thumb for a acceptable fit (0.90).

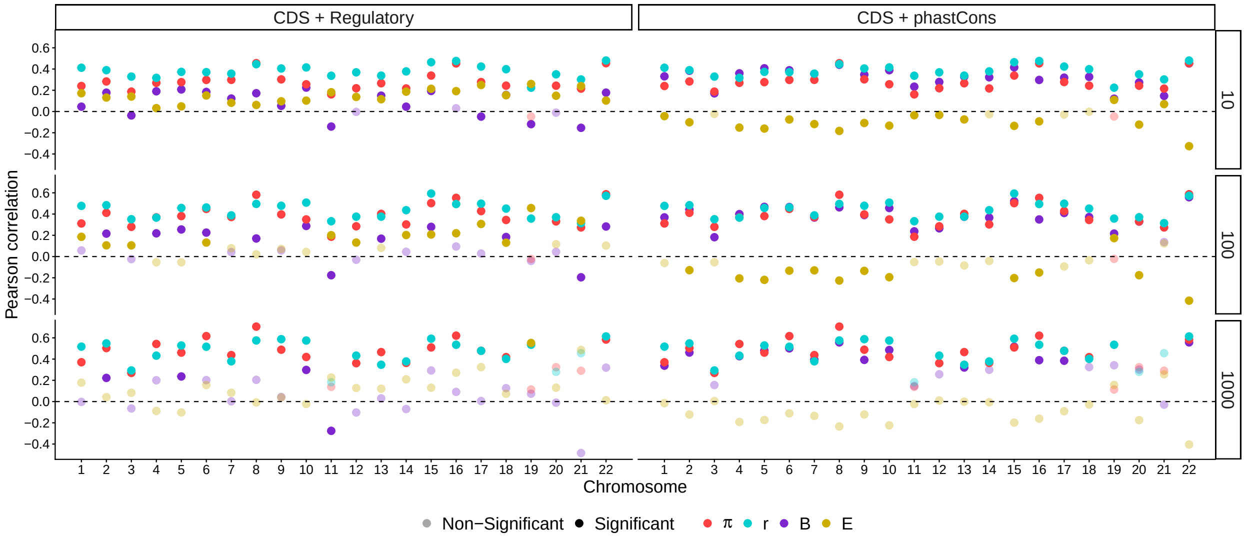

Figure S33: Pearson correlations between mutation rate (latent predictor) and other variables involved in our DAGs, shown for different functional elements (column panels) and window sizes (kb, row panels). Alpha scale denotes significance after Bonferroni correction over chromosomes ($p < 0.05/22$).

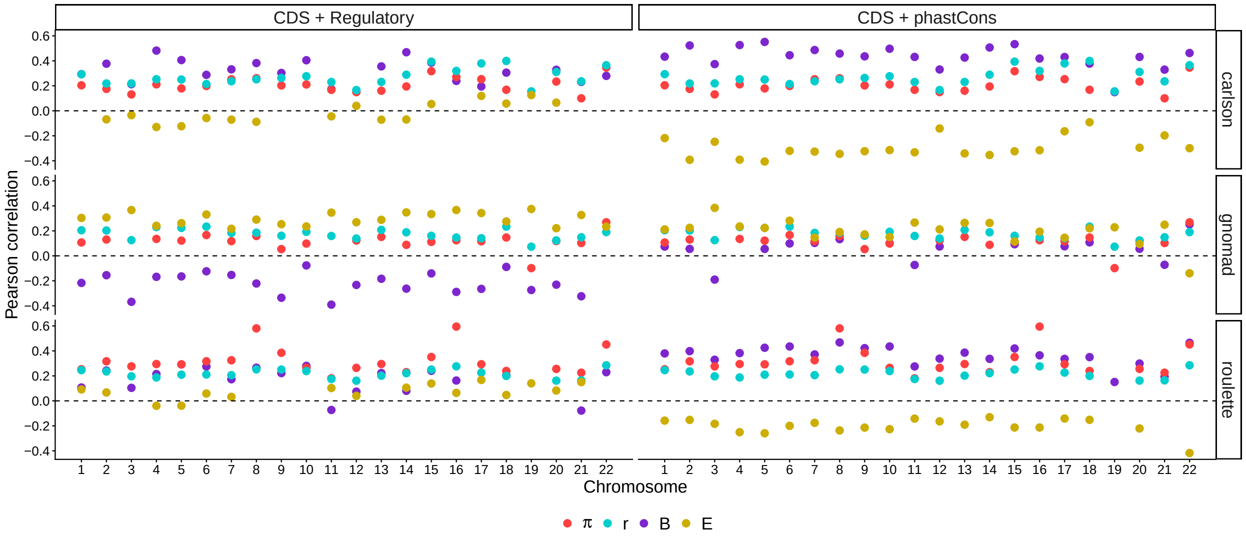

Figure S34: Pearson correlations between mutation rate (raw maps, row panels) and other variables involved in our DAGs (color), shown for different functional elements (column panels). Only correlations significant after Bonferroni correction over chromosomes are plotted ($p < 0.05/22$). Here 10 kb scale.

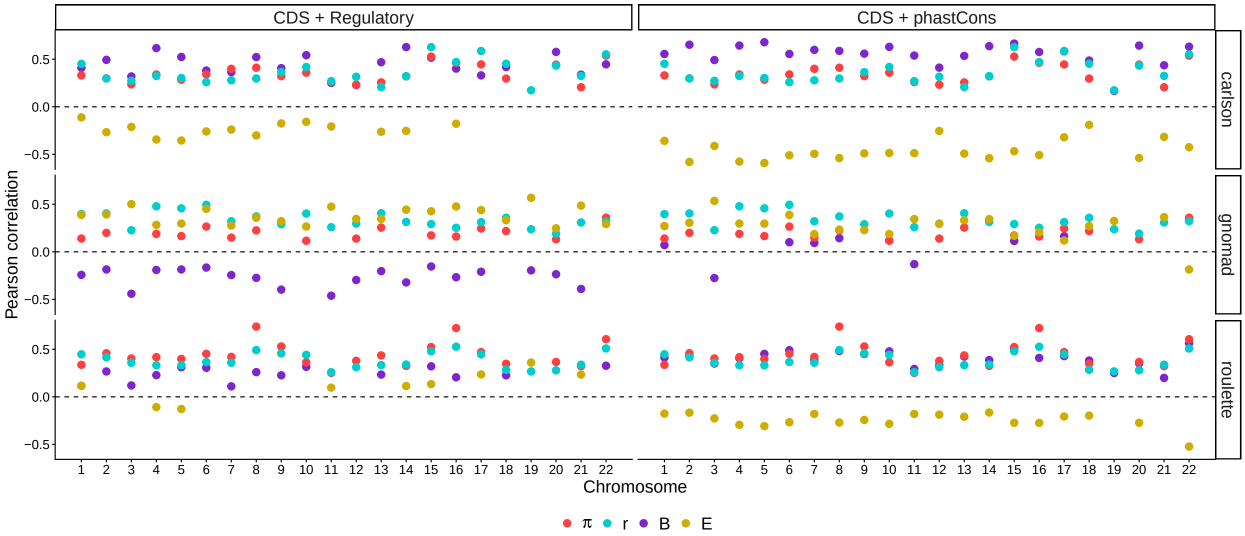

Figure S35: Pearson correlations between mutation rate (raw maps, row panels) and other variables involved in our DAGs (color), shown for different functional elements (column panels). Only correlations significant after Bonferroni correction over chromosomes are plotted ($p < 0.05/22$). Here 100 kb scale.

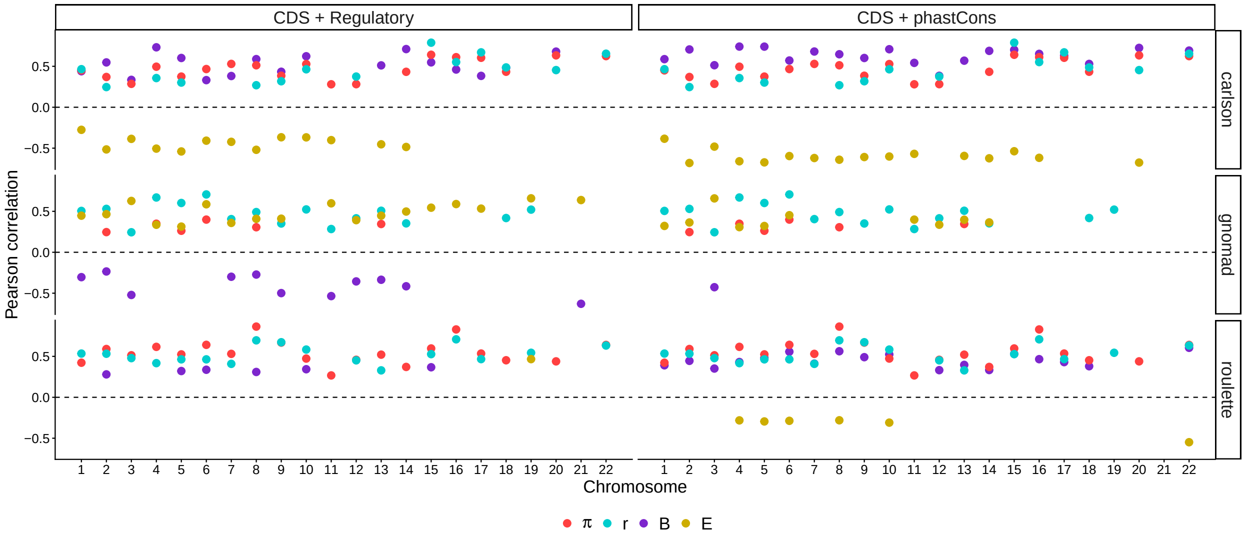

Figure S36: Pearson correlations between mutation rate (raw maps, row panels) and other variables involved in our DAGs (color), shown for different functional elements (column panels). Only correlations significant after Bonferroni correction over chromosomes are plotted ($p < 0.05/22$). Here 1 Mb scale.

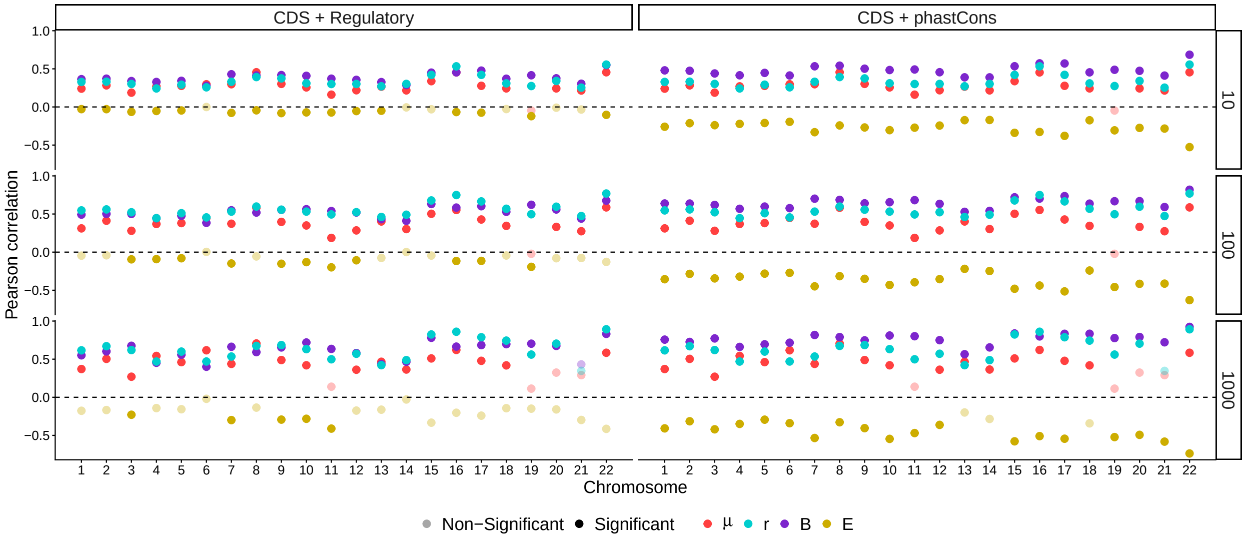

Figure S37: Pearson correlations between $\hat{\pi}$ and other variables involved in our DAGs (color), shown for different functional elements (column panels) and window sizes (kb, row panels). Alpha scale denotes significance after Bonferroni correction over chromosomes ($p < 0.05/22$). Latent constructs are used for $\mu$, $B$ and $r$.

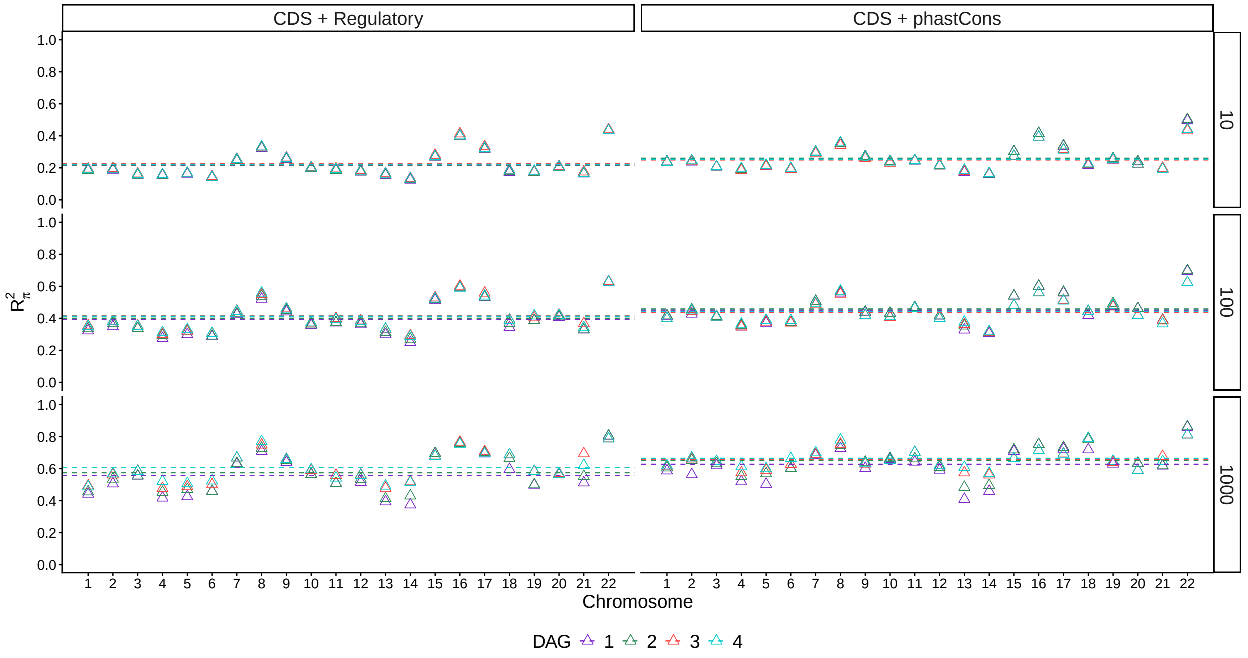

Figure S38: Variation in $R^2_\pi$ among autosomes, shown for different DAGs (colors), constrained elements (column panels) and window sizes (kb, row panels). Dashed horizontal lines show genome-wide averages weighted by chromosome length.

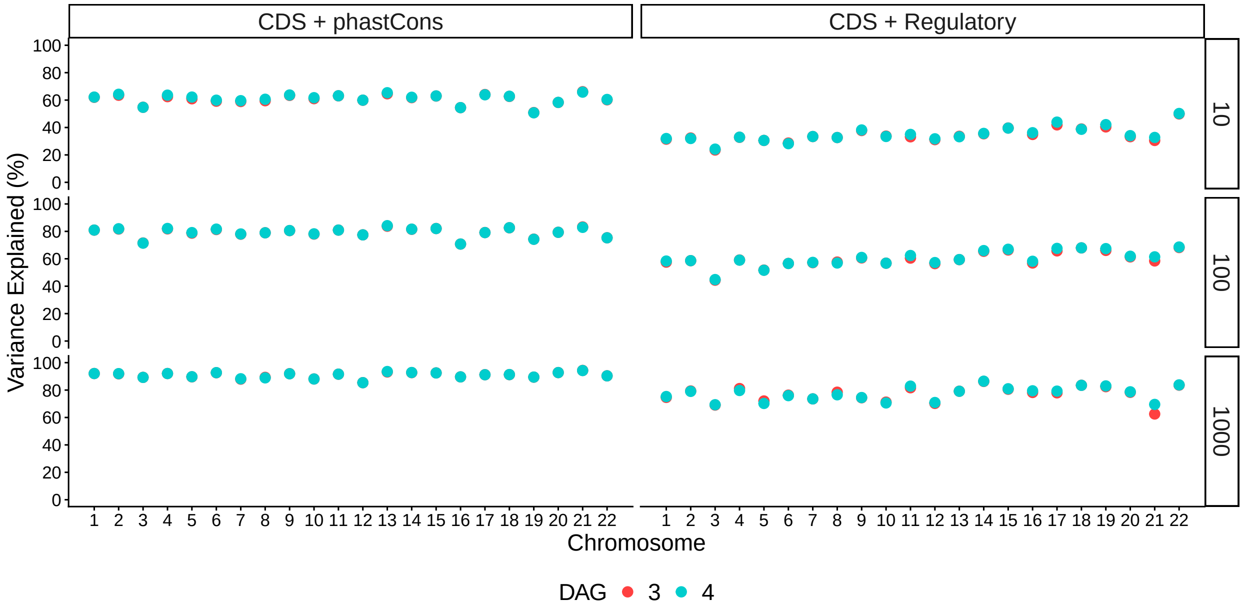

Figure S39: Variation in $R_B^2$ (the variance explained in $B$-values) among chromosomes, shown for different DAGs (colors), constrained elements (column panels) and window sizes (kb, row panels).

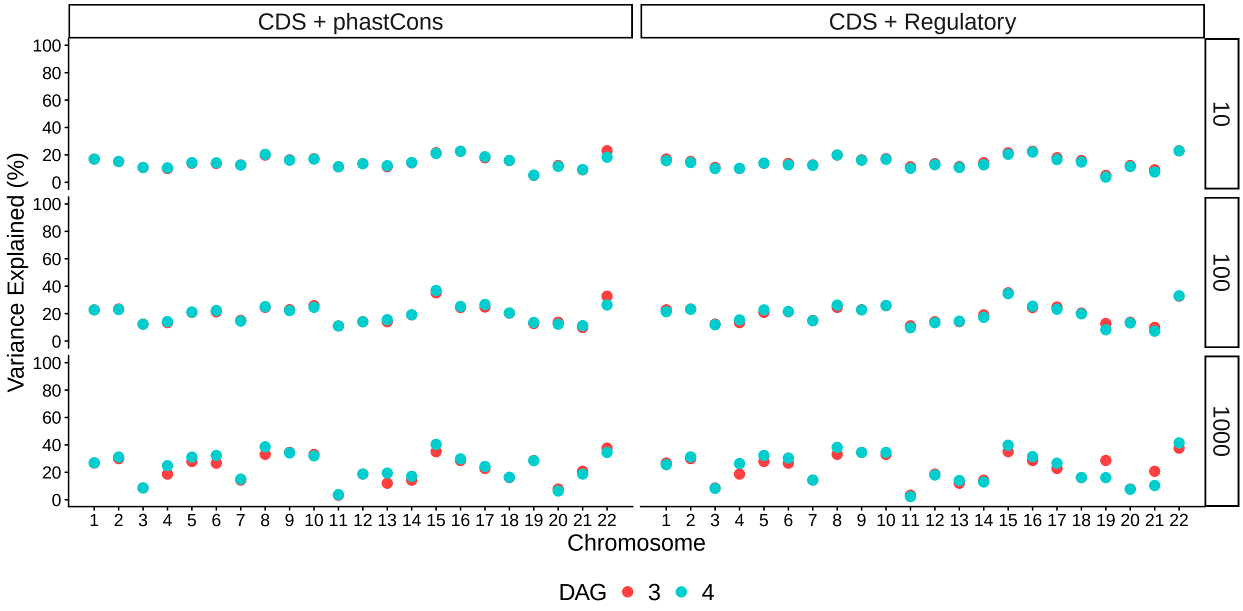

Figure S40: Variation in $R^2_\mu$ (the variance explained in mutation rates) among chromosomes, shown for different DAGs (colors), constrained elements (column panels) and window sizes (kb, row panels).

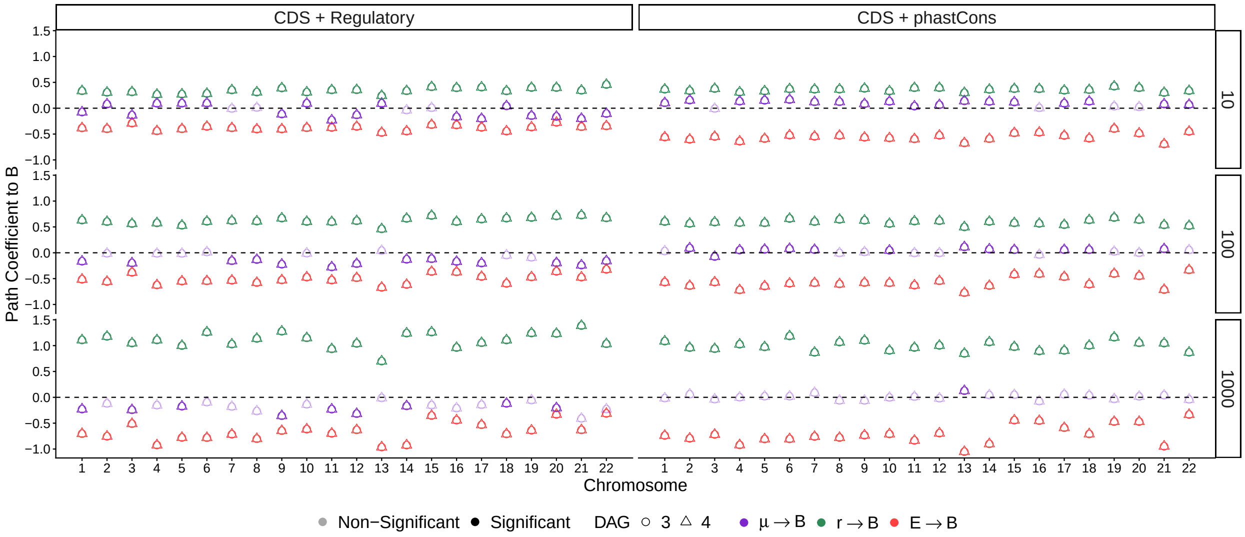

Figure S41: Estimated path coefficients (color) going into $B$, for each autosome, constrained elements (column panels) and window sizes (kb, row panels). Alpha scale denotes significance after Bonferroni correction over chromosomes ($p < 0.05/22$). Circles denote DAG 3 and triangles denote DAG 4.

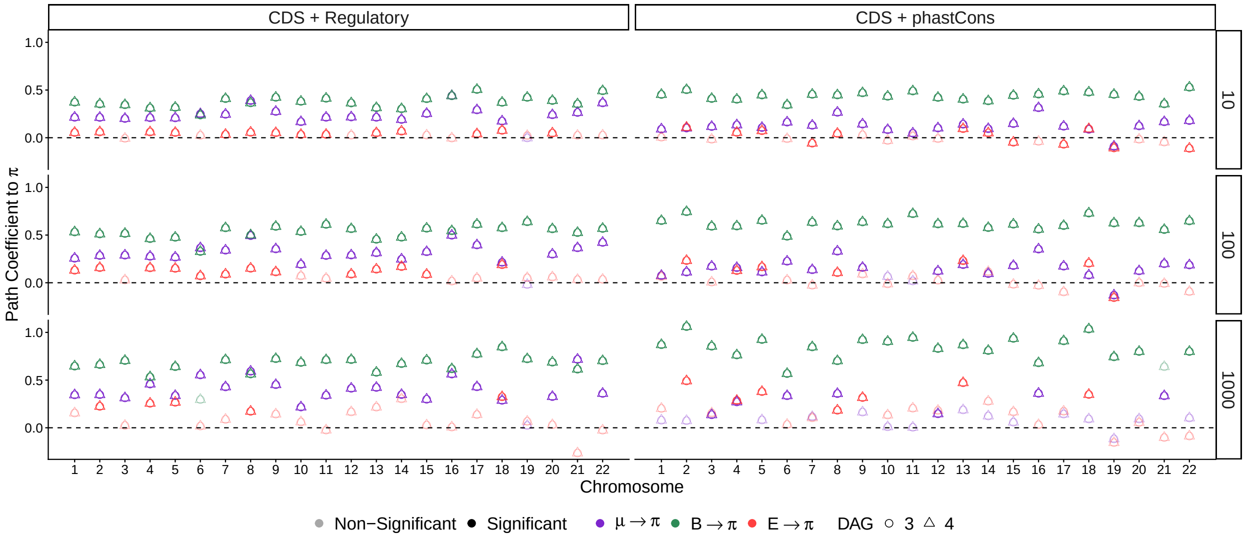

Figure S42: Estimated path coefficients (color) going into $\pi$, for each autosome, constrained elements (column panels) and window sizes (kb, row panels). Alpha scale denotes significance after Bonferroni correction over chromosomes ($p < 0.05/22$). Circles denote DAG 3 and triangles denote DAG 4.

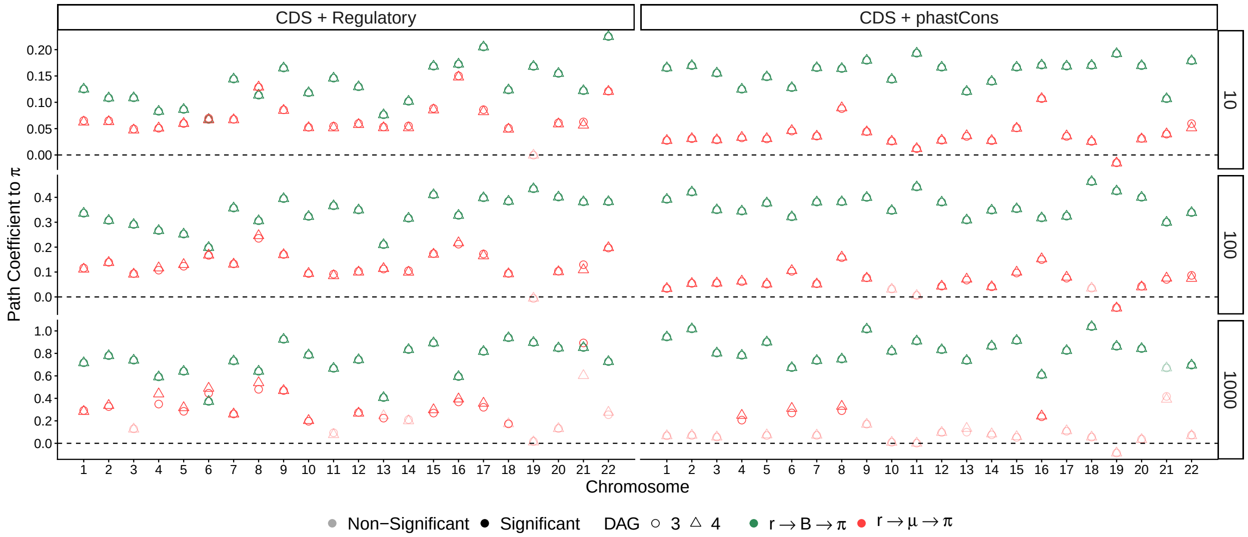

Figure S43: Estimated path coefficients from $r$ to $\pi$ (color), for each autosome, constrained elements (column panels) and window sizes (kb, row panels). Alpha scale denotes significance after Bonferroni correction over chromosomes ($p < 0.05/22$). Circles denote DAG 3 and triangles denote DAG 4.

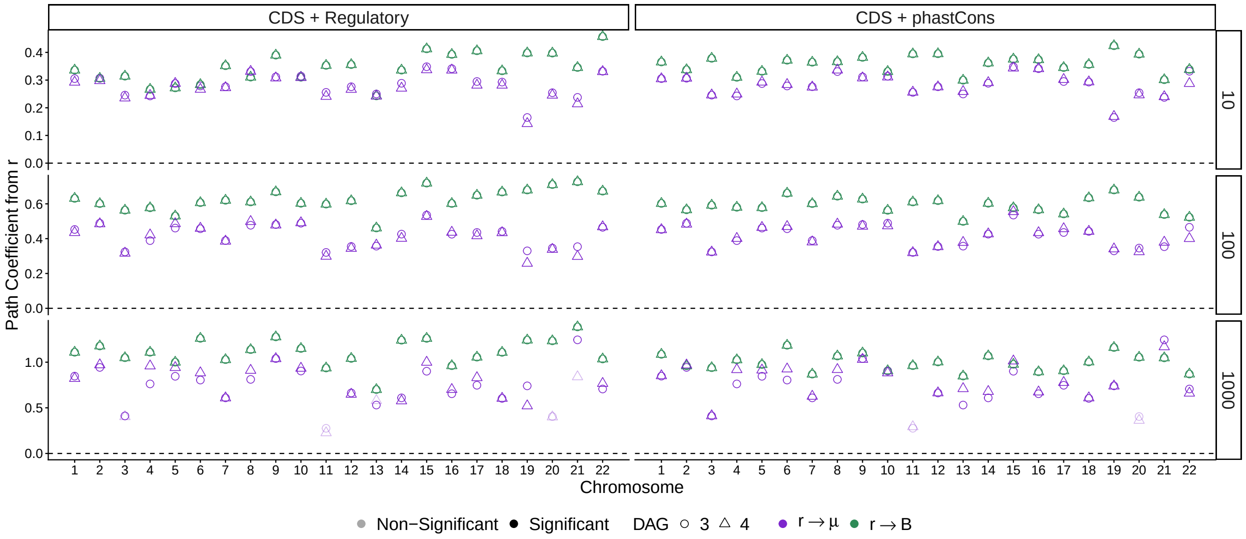

Figure S44: Estimated (direct) path coefficients from $r$ (color), for each autosome, constrained elements (column panels) and window sizes (kb, row panels). Alpha scale denotes significance after Bonferroni correction over chromosomes ($p < 0.05/22$). Circles denote DAG 3 and triangles denote DAG 4.

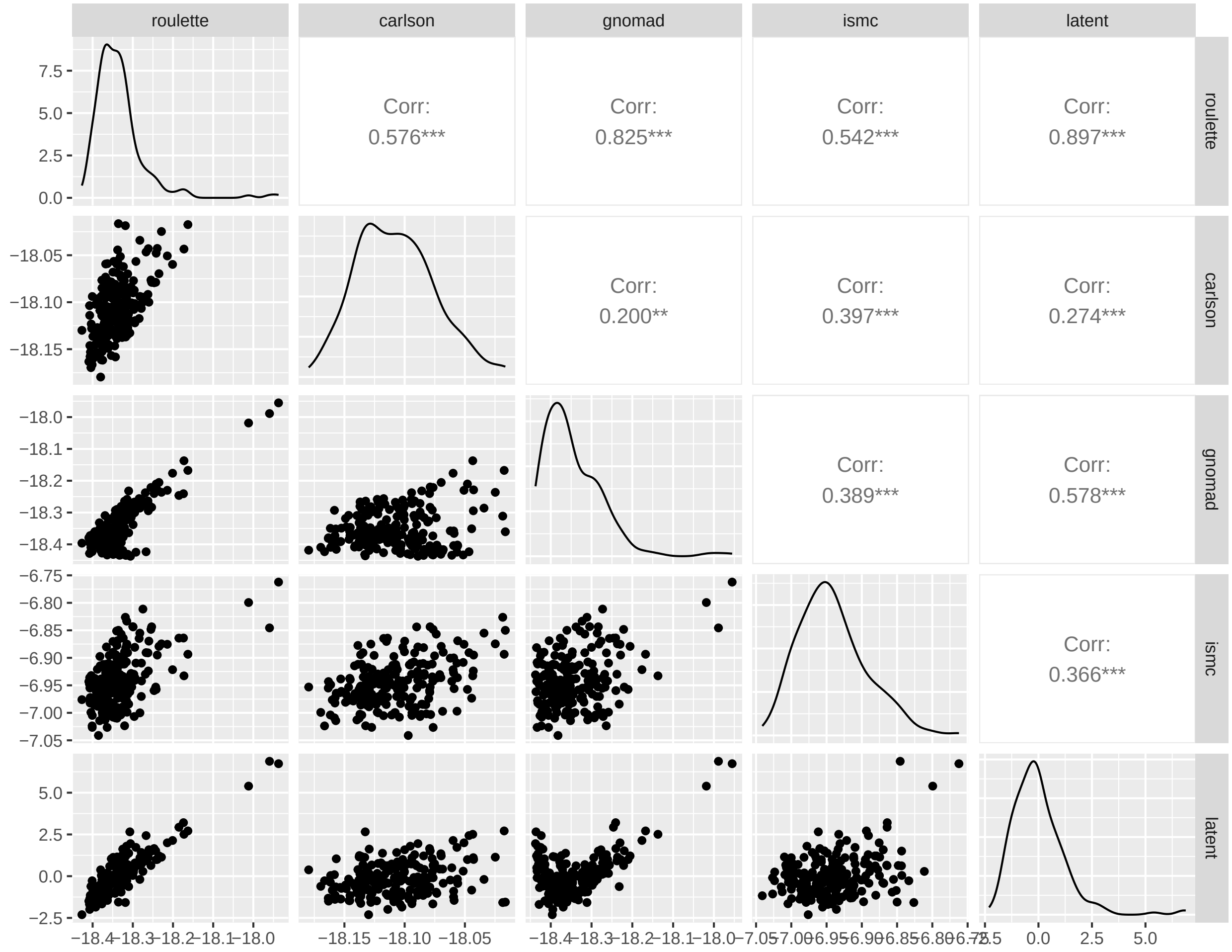

Figure S45: Visual summary of the measurement model of mutation rate variation for chromosome 1 at the 1 Mb scale. Shown are pairwise relationships between all maps (indicators and the latent construct).

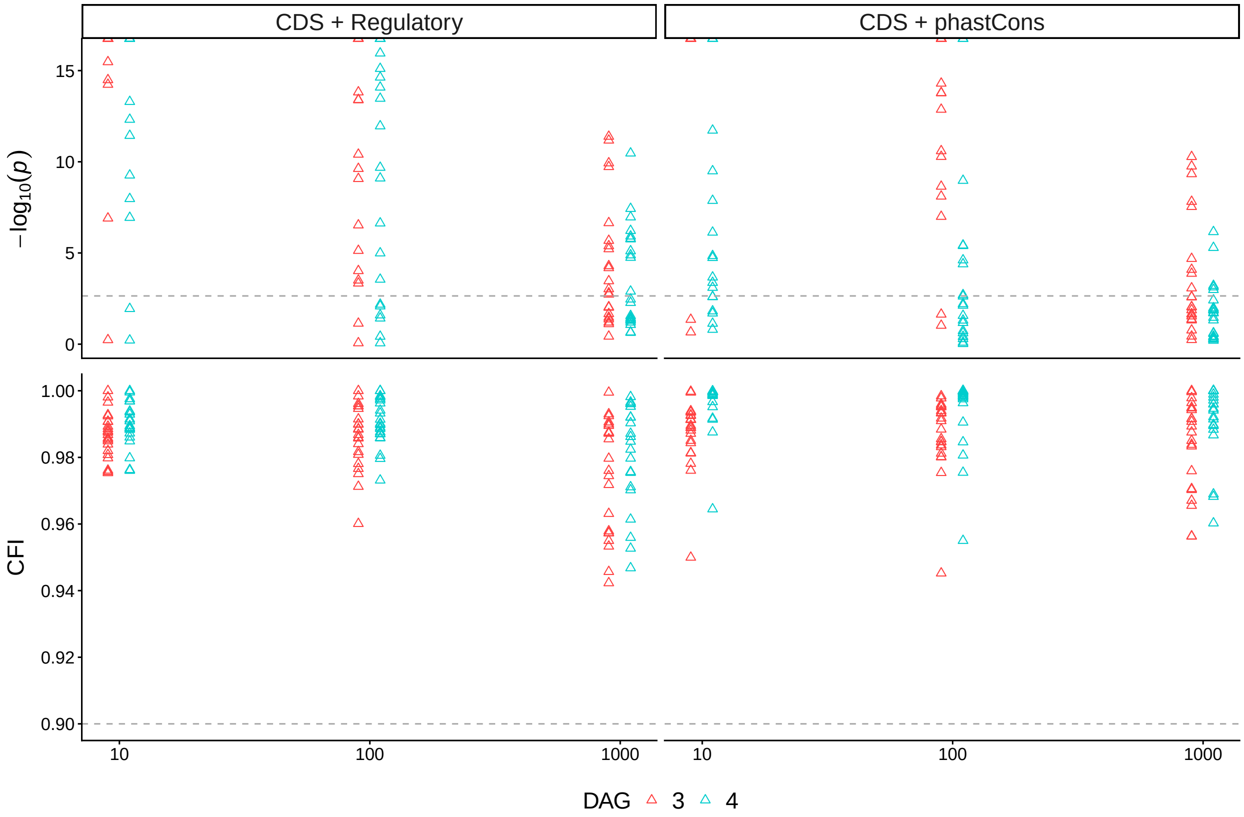

Figure S46: Goodness-of-fit of models 3 and 4 when $\pi$ is simulated from $\mathbb{E}[\pi]$ (Section 5.1). **A:** $\chi^2$ $p$-values across autosomes. Dashed lines show significance threshold after Bonferroni correction over chromosomes (0.05/22). Very small $p$-values are pressed against the top of each panel. **B:** Comparative Fit Indices across chromosomes. Dashed grey lines show the rule-of-thumb for a acceptable fit (0.90).

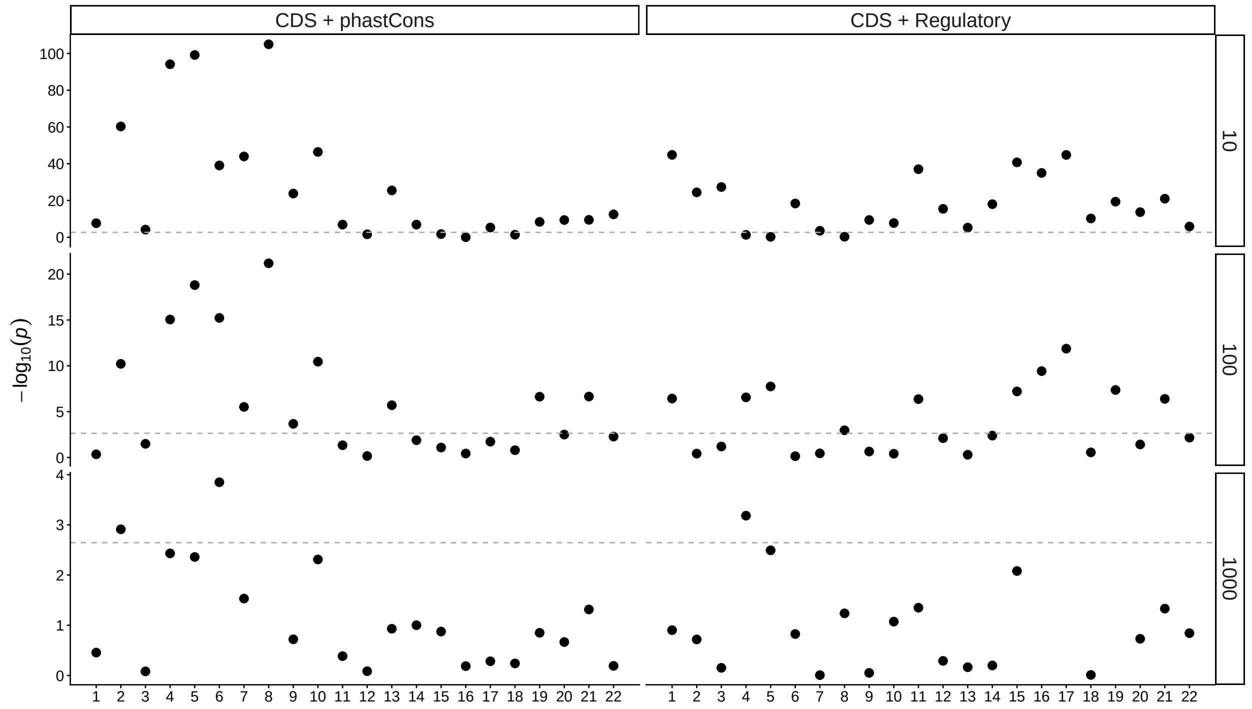

Figure S47: Model choice of nested DAGs 3 and 4. Shown are $p$-values from likelihood ratio tests for each chromosome, different functional elements (column panels) and window sizes (kb, row panels). The dashed horizontal line shows the significance threshold after Bonferroni correction over chromosomes (0.5/22).

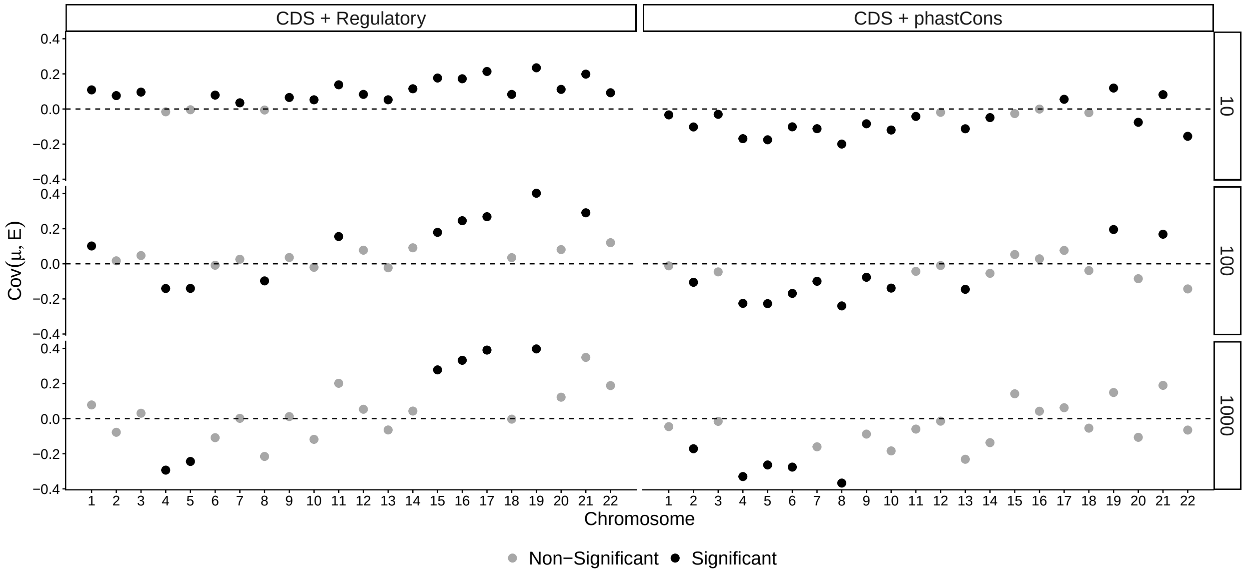

Figure S48: Estimated covariance between mutation rate and density of constrained elements, a free parameter in DAG 4. Estimates are shown for each chromosome, constrained elements (column panels) and window sizes (kb, row panels). Alpha scale denotes significance after Bonferroni correction over chromosomes ($p < 0.05/22$).

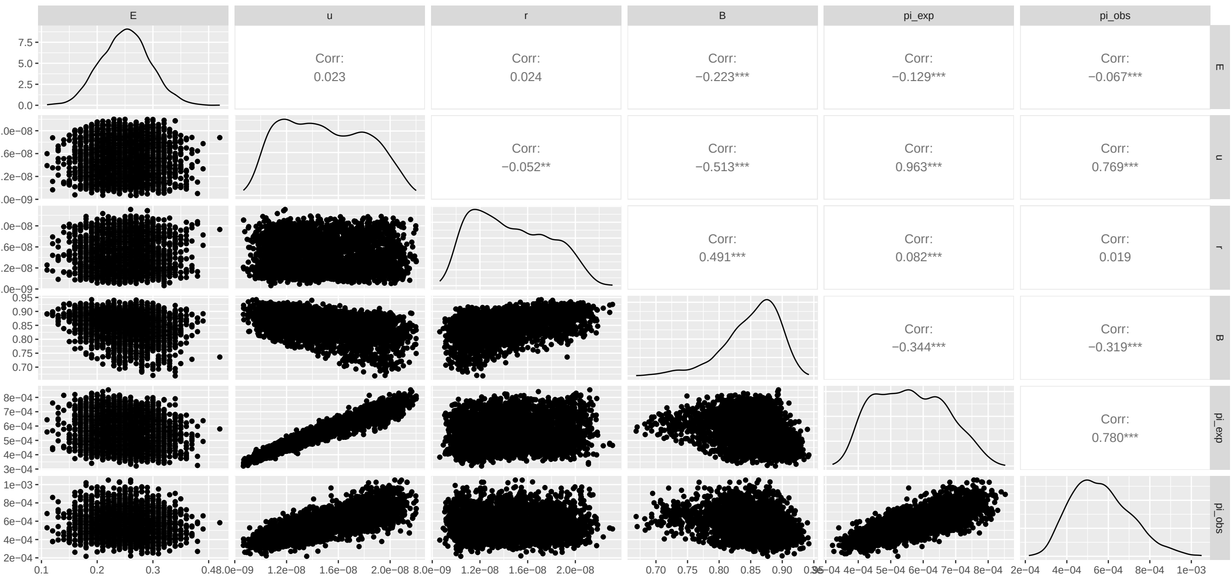

Figure S49: Pairwise correlations among genomic landscapes in simulation set A: 100x 10-Mb chromosomes. Scatter plots show data (100 kb windows, down-sampled to 10%) across all chromosomes. There is a significantly negative correlation between $\mu$ and $r$, despite independent draws to build these maps. Note also the strong negative correlation between $B$ and $\pi$ that arises through their common parent, $\mu$, which affects them with opposite sign effects.

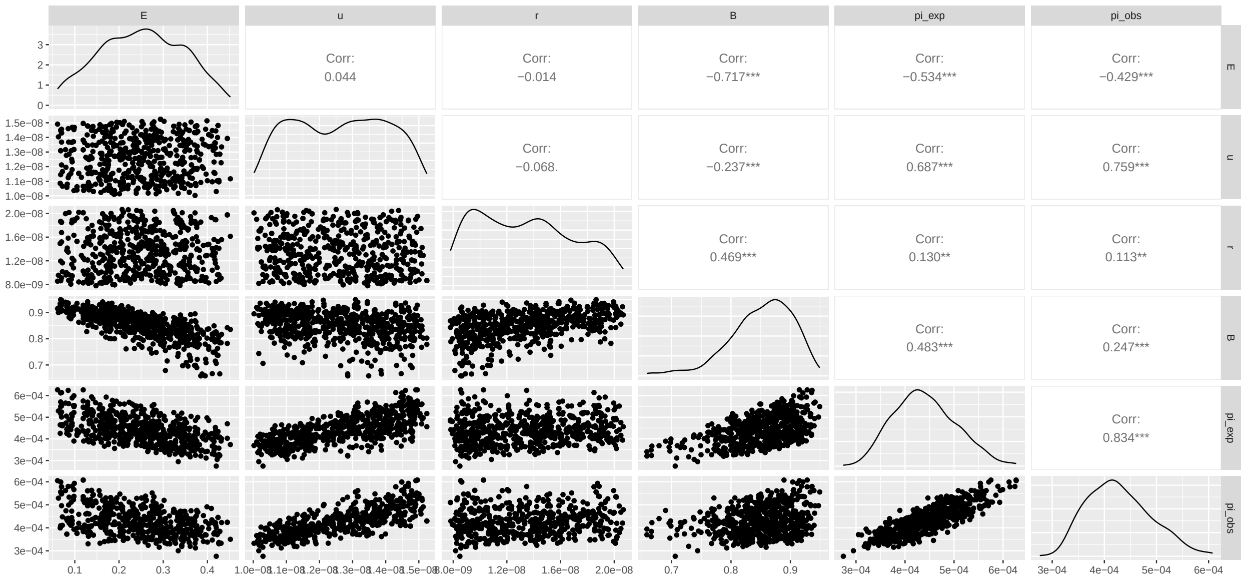

Figure S50: Pairwise correlations among genomic landscapes in simulation set B: 10x 100-Mb chromosomes. Scatter plots show data (1 Mb windows, down-sampled to 20%) across all chromosomes.

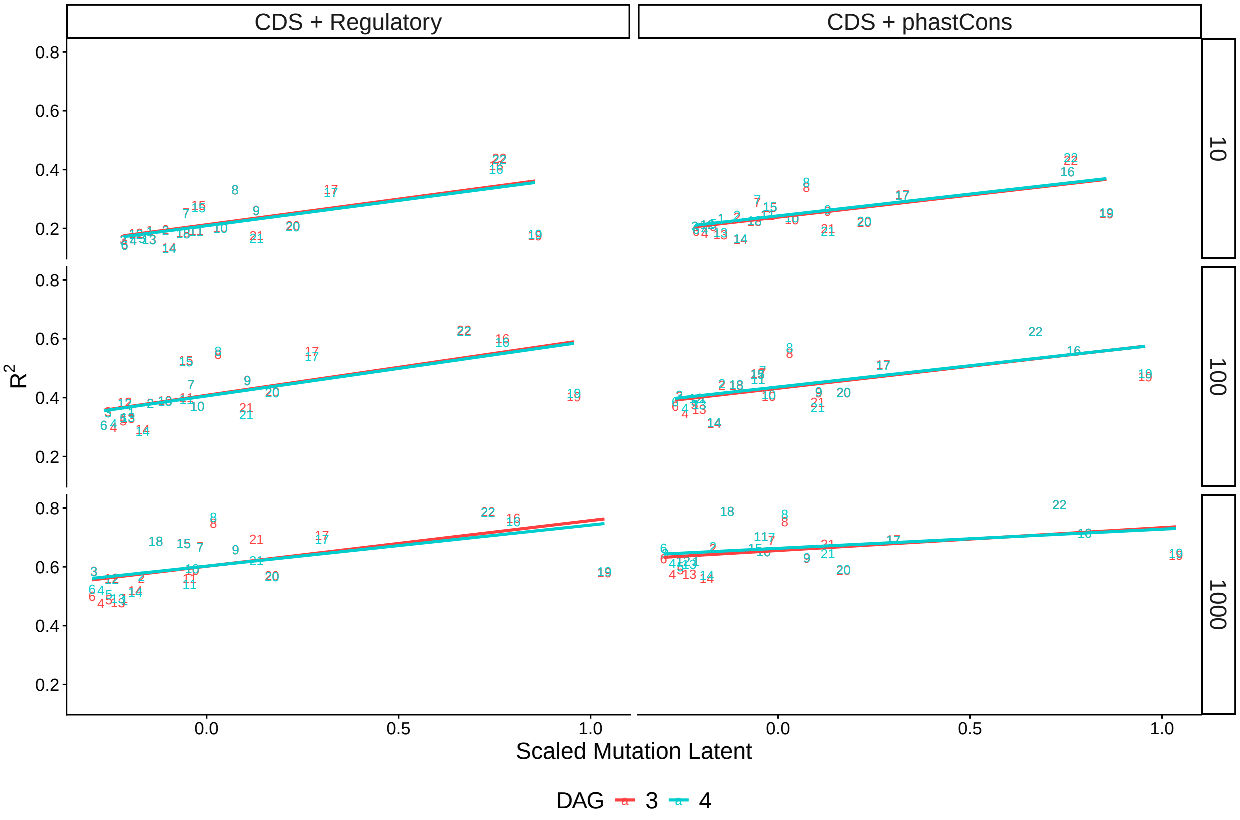

Figure S51: $R^2_\pi$ vs average mutation rate per chromosome per model, with least-squares regression lines shown.

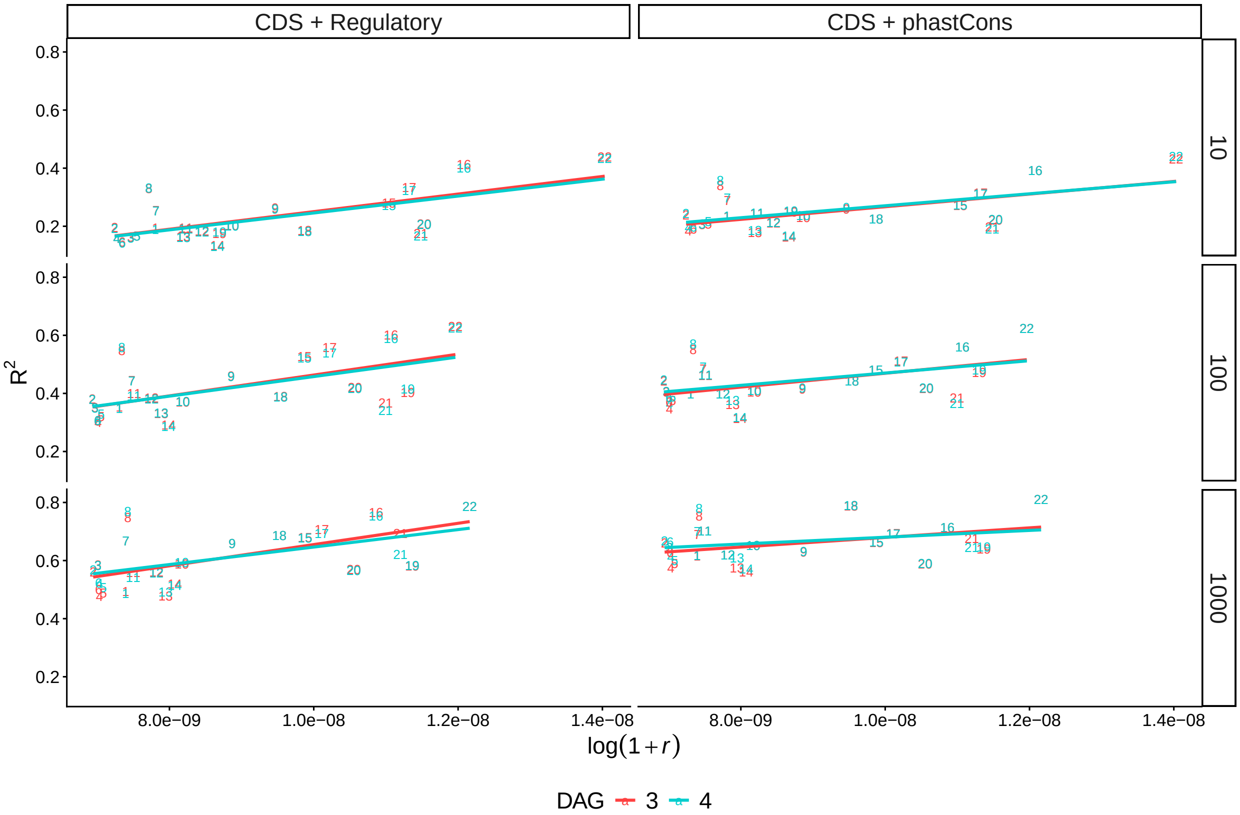

Figure S52: $R^2_\pi$ vs average recombination rate per chromosome per model, with least-squares regression lines shown.

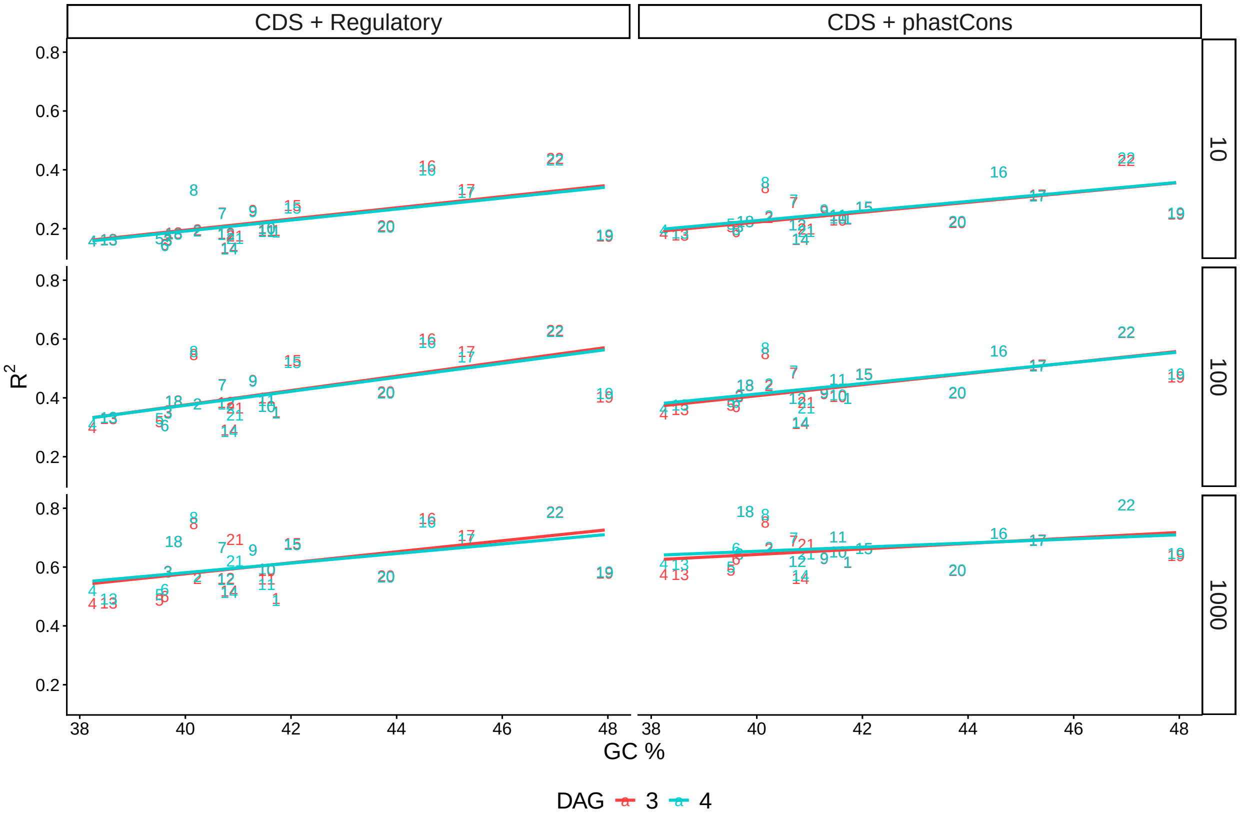

Figure S53: $R^2_\pi$ vs GC content per chromosome per model, with least-squares regression lines shown.

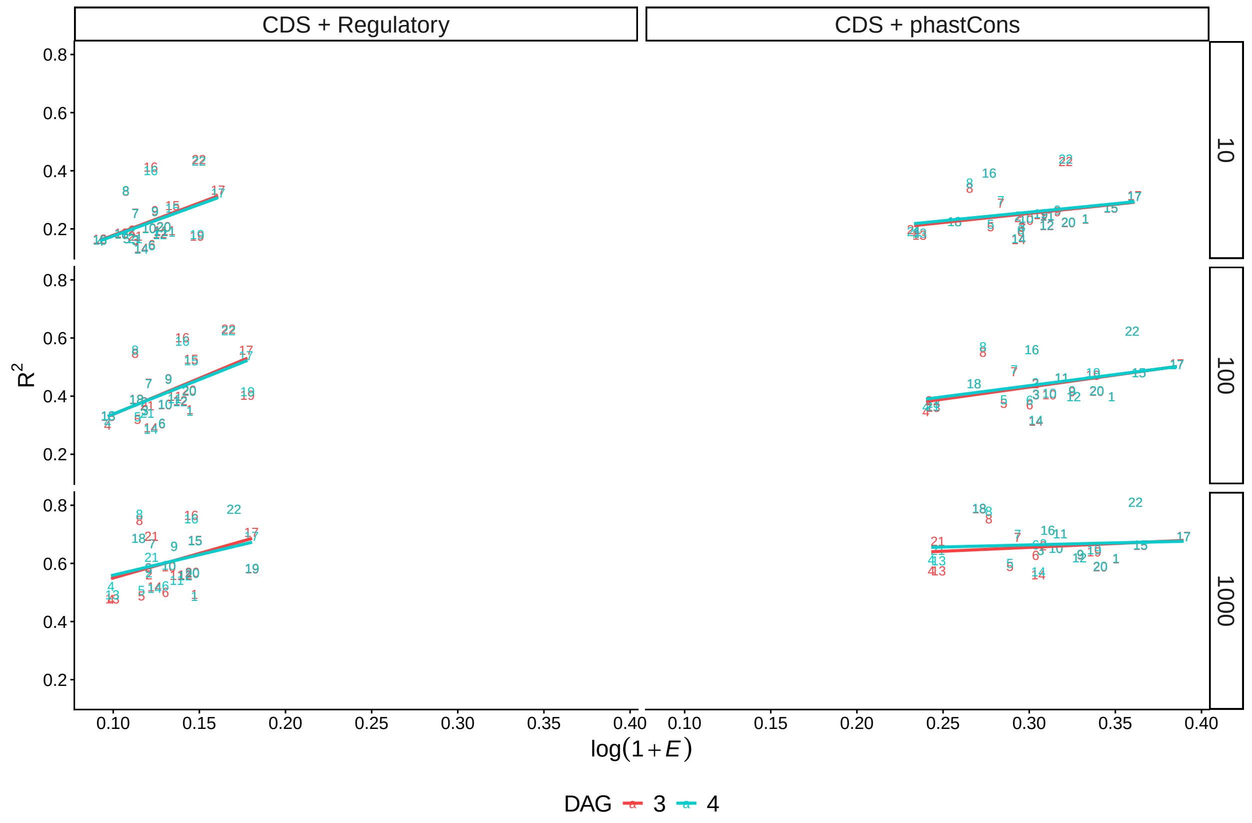

Figure S54: $R^2_\pi$ vs proportion of deleterious sites per chromosome per model, with least-squares regression lines shown.

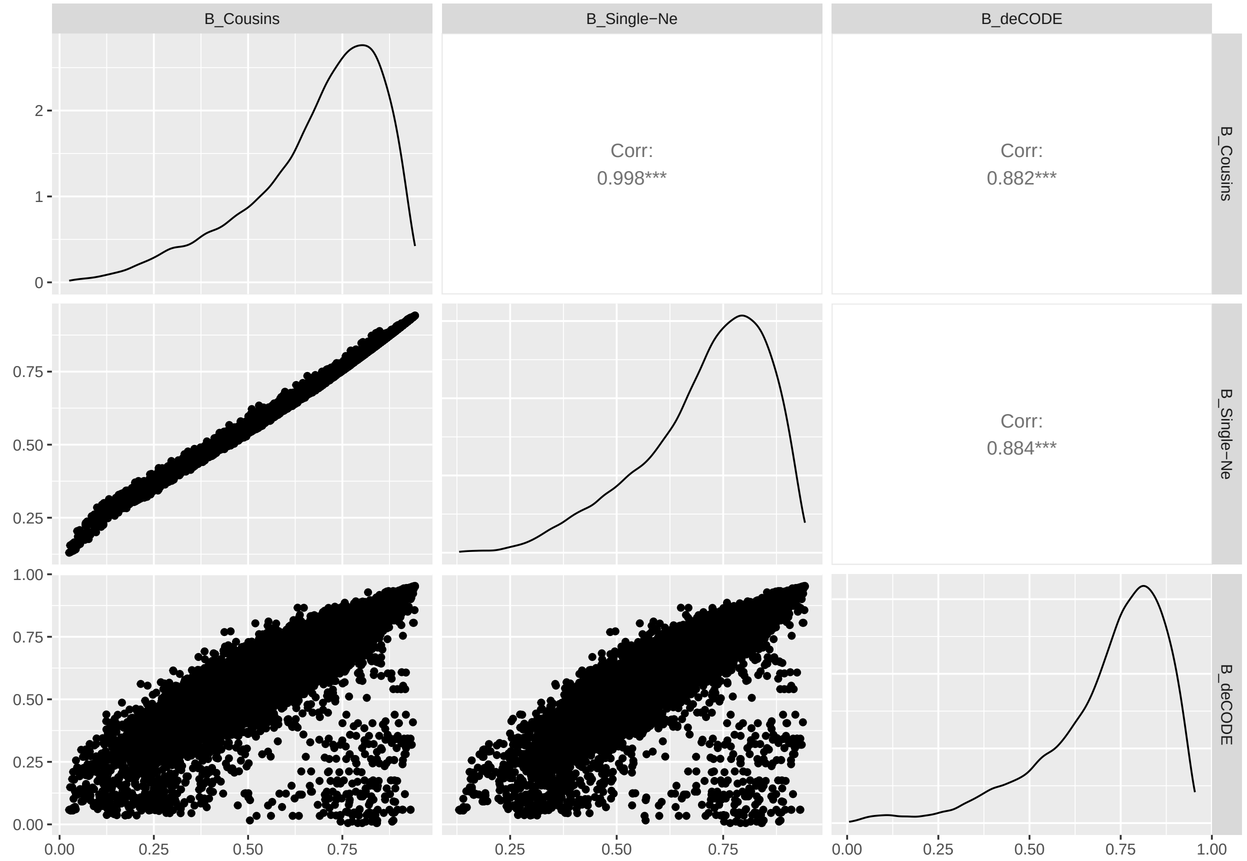

Figure S55: Pairwise correlations between $B$-values predicted under alternative models in humans (Sections 5.3–5.5). Here plotted at the 100 kb scale, down-sampled to 20% of windows across the genome.

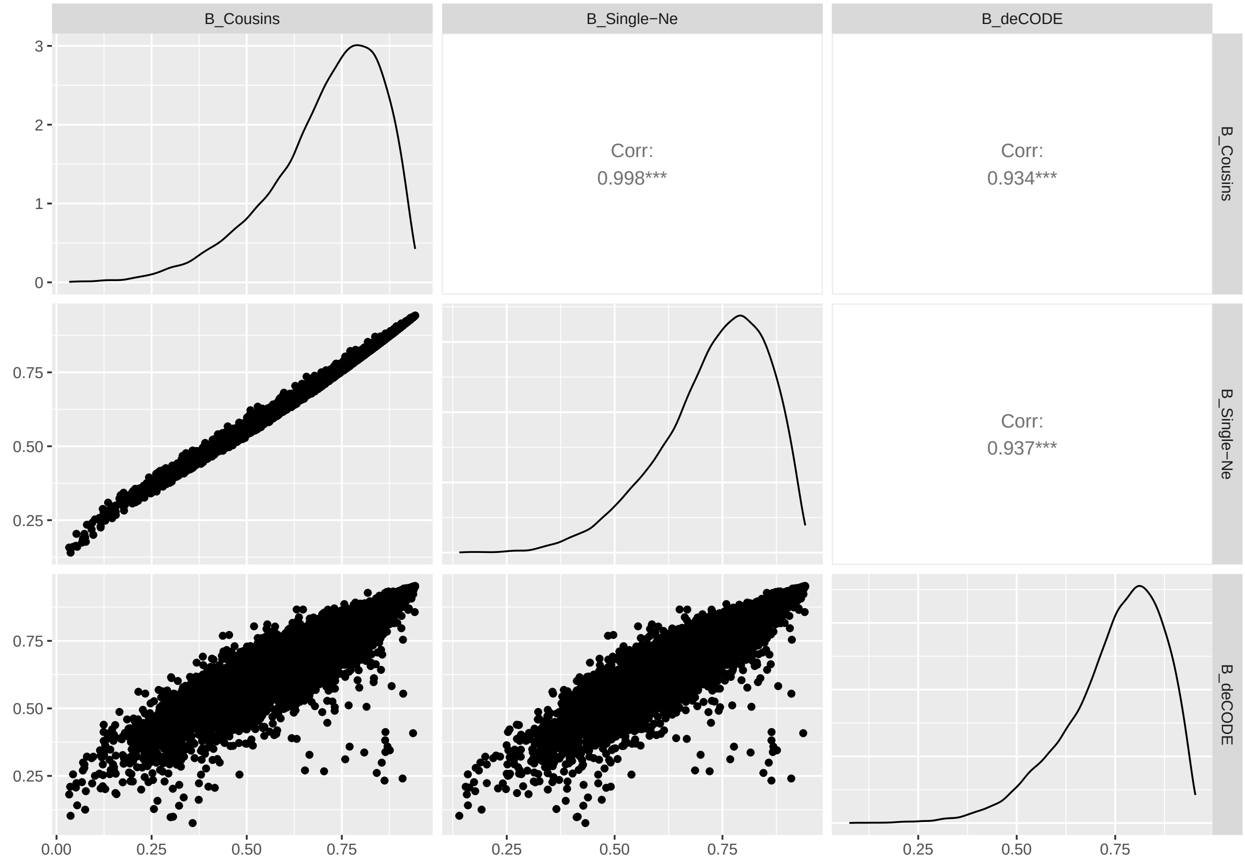

Figure S56: Pairwise correlations between $B$-values predicted under alternative models in humans (Sections 5.3–5.5). Here plotted at the 100 kb scale, for windows where the deCODE estimates of recombination are $> 10^{-9}$, down-sampled to 20% of windows across the genome.

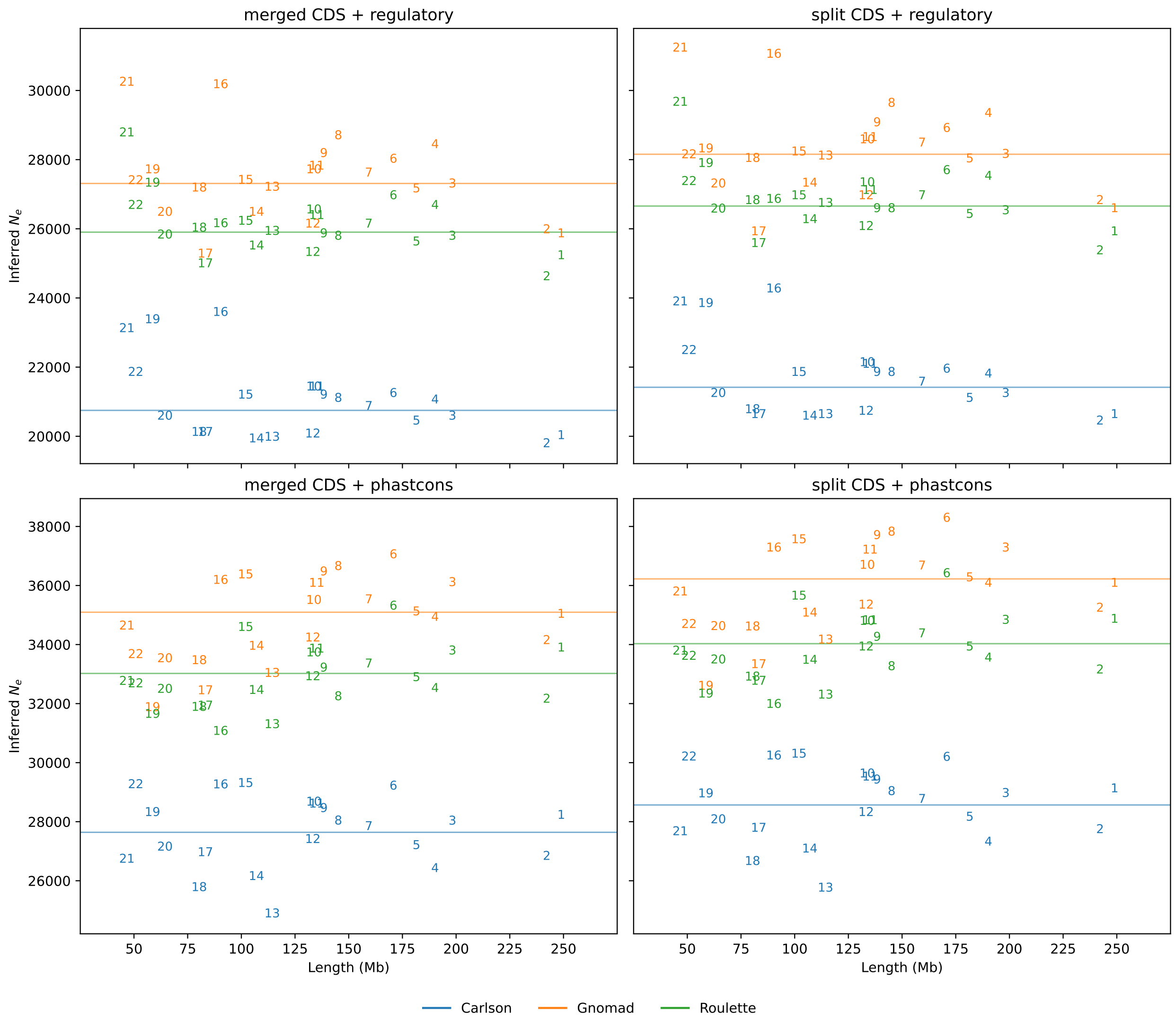

Figure S57: Estimated $N_e$ vs. chromosome length across constraint and mutation models. Markers indicate chromosome number and horizontal lines show average $N_e$, weighted by the number of accessible sites on each chromosome. For each combination of constrained model and mutation map, $N_e$ estimates are consistent across chromosomes. Here shown for the `pyrho` recombination map estimated for the Yoruba population.

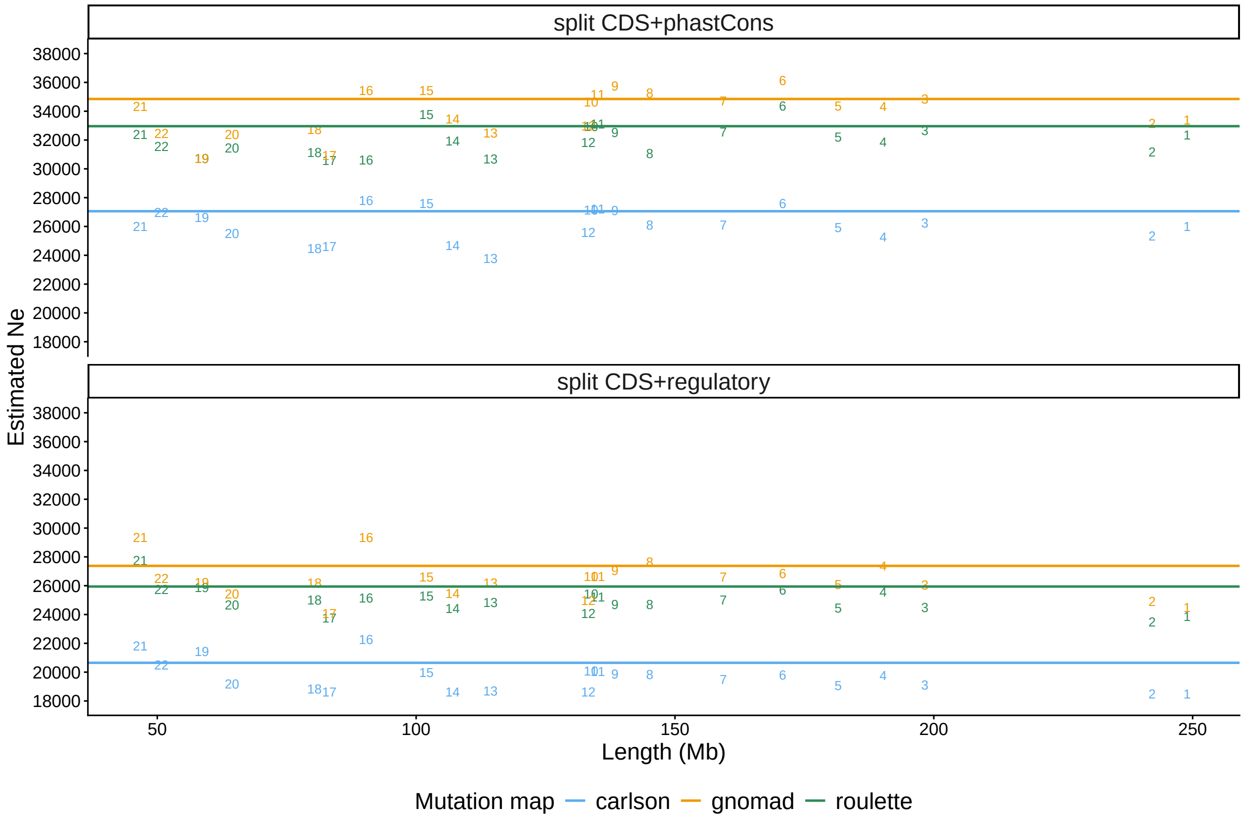

Figure S58: Estimated $N_e$ vs. chromosome length across constraint and mutation models. Markers indicate chromosome number and horizontal lines show average $N_e$, weighted by the number of accessible sites on each chromosome. For each combination of constrained model and mutation map, $N_e$ estimates are consistent across chromosomes. Here shown for the deCODE recombination map.

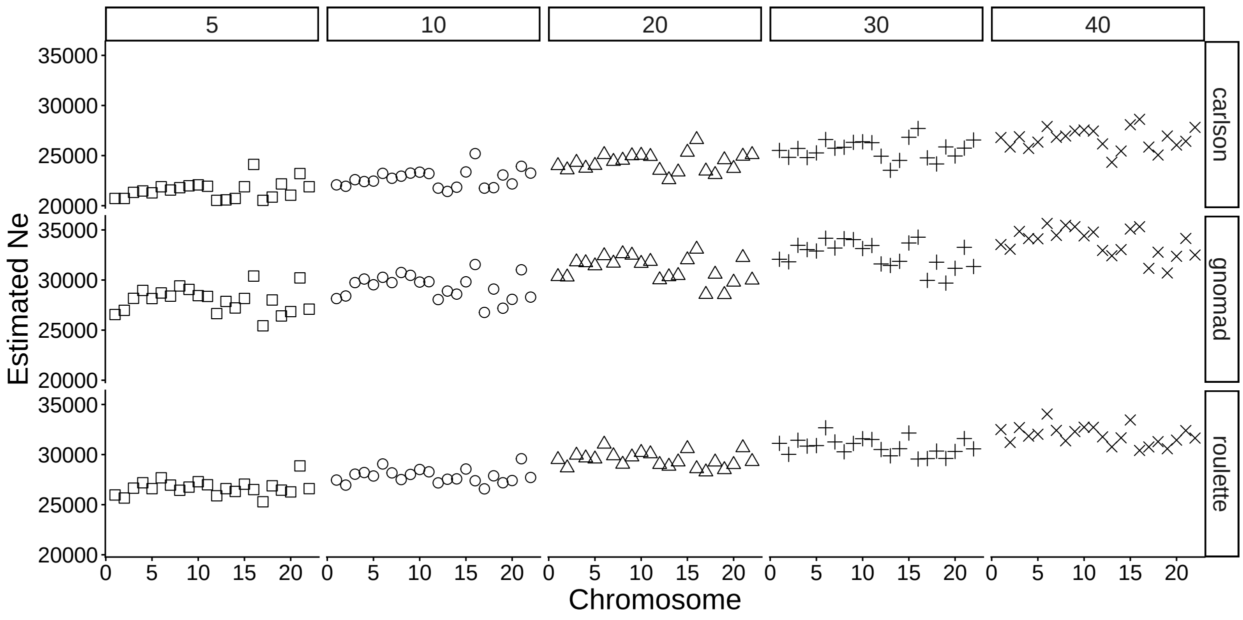

Figure S59: Estimates of the drift-effective population size ($N_e$) across autosomes, for different mutation maps (rows) and thresholds for including phastCons as targets of direct selection (columns). Exonic DFEs are split among deciles of constraint. Less stringent cut-offs (higher threshold) predict stronger reductions in diversity due to direct and background selection, thus compensating with higher estimated $N_e$ to explain $\hat{\pi}_{obs}$ (via $\mathbb{E}[\pi_0] = 4N_e\mu$).
