## Supplementary figures and images for "Causal inference clarifies the roles of background selection and mutation rate variation in shaping human genetic diversity"

### aov_1Mb_chr1.pdf

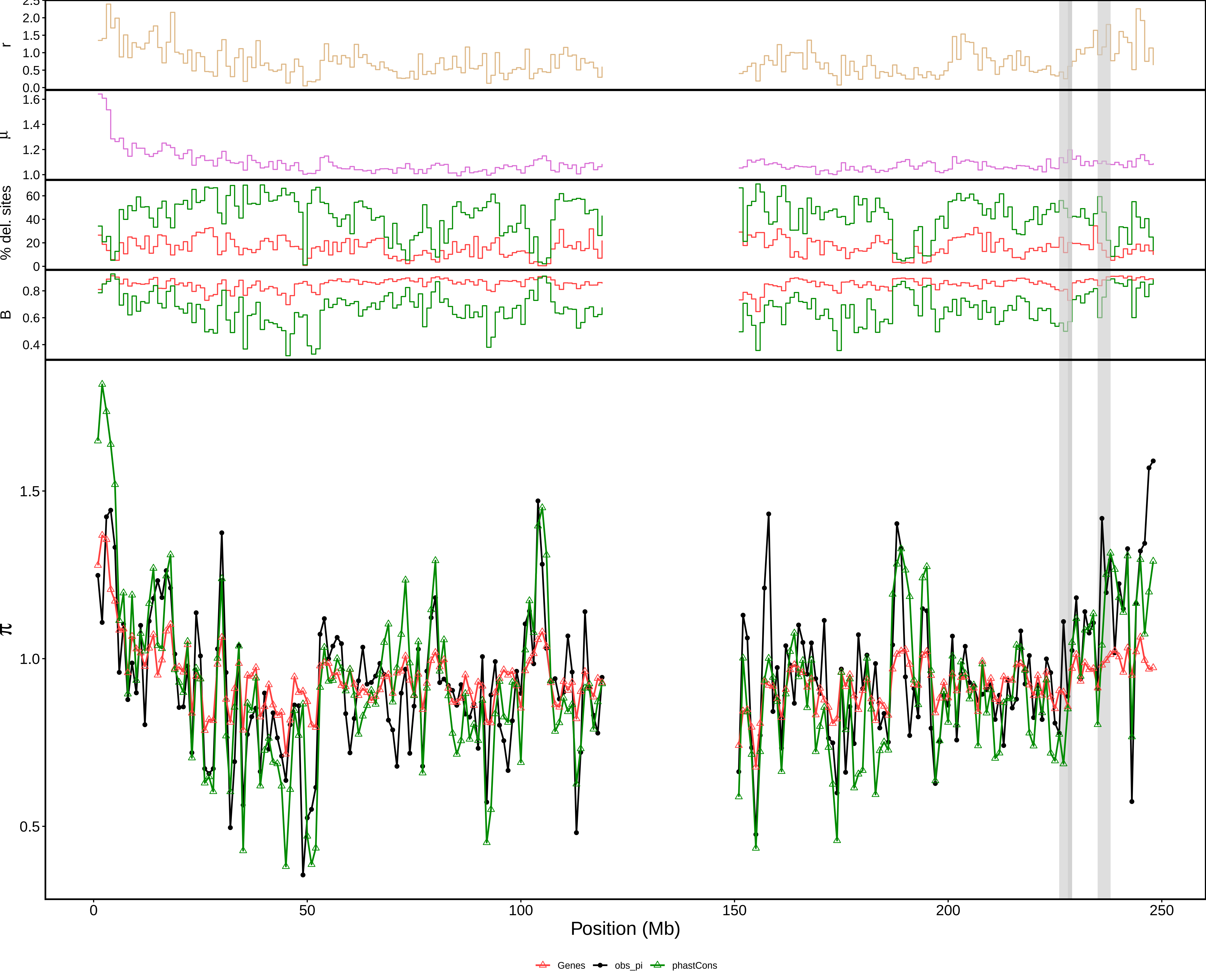

### aov_1Mb_chr2.pdf

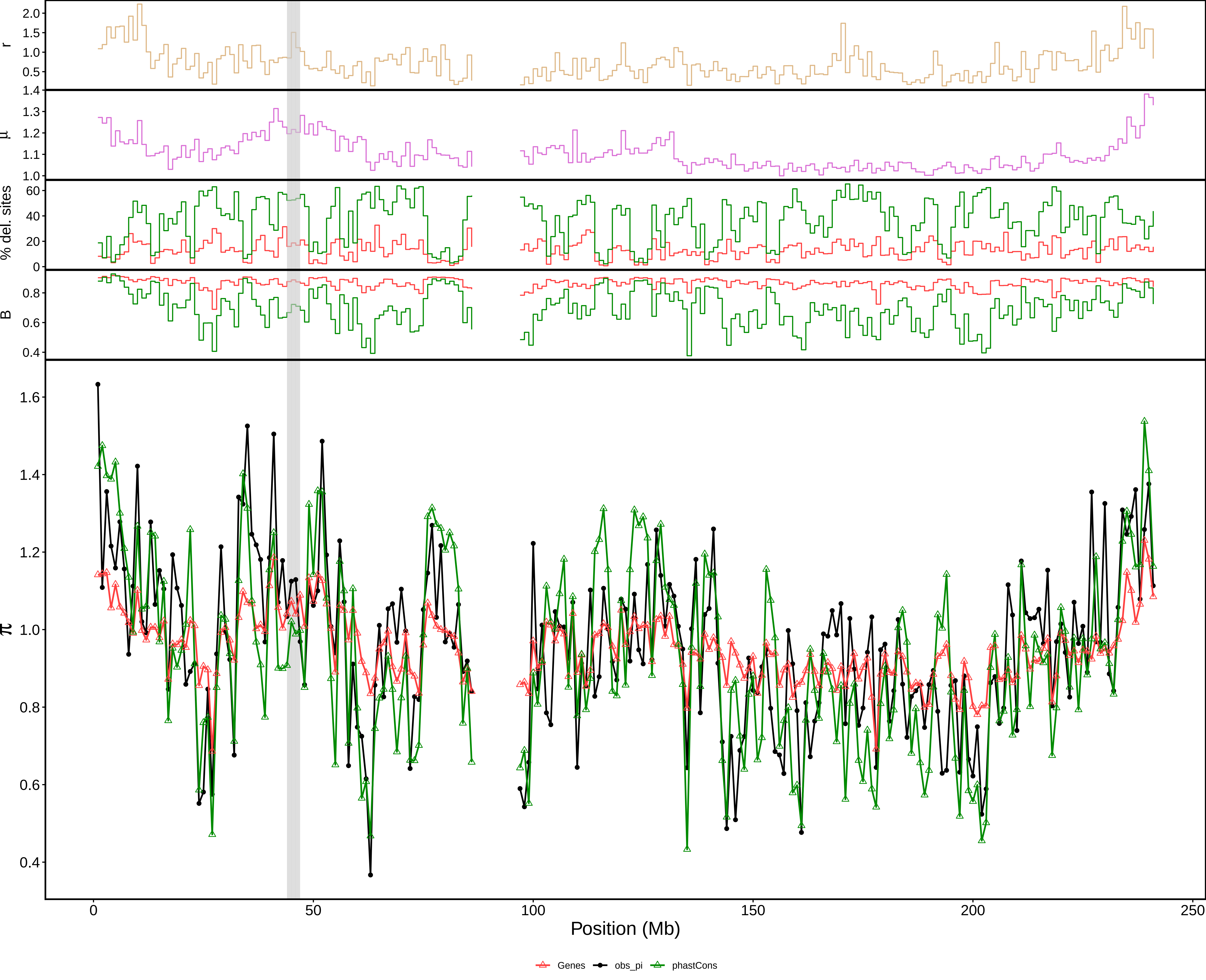

### aov_1Mb_chr3.pdf

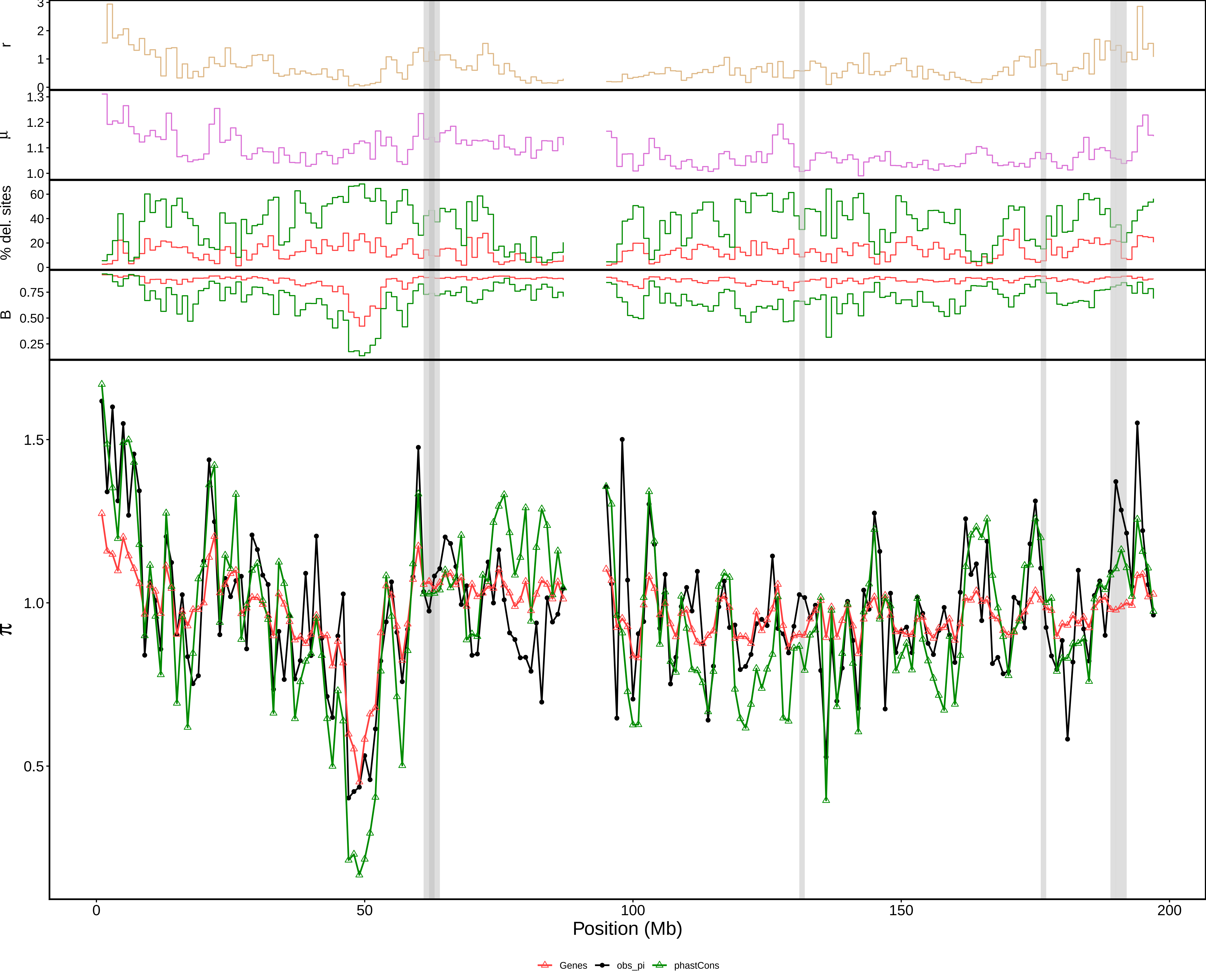

### aov_1Mb_chr4.pdf

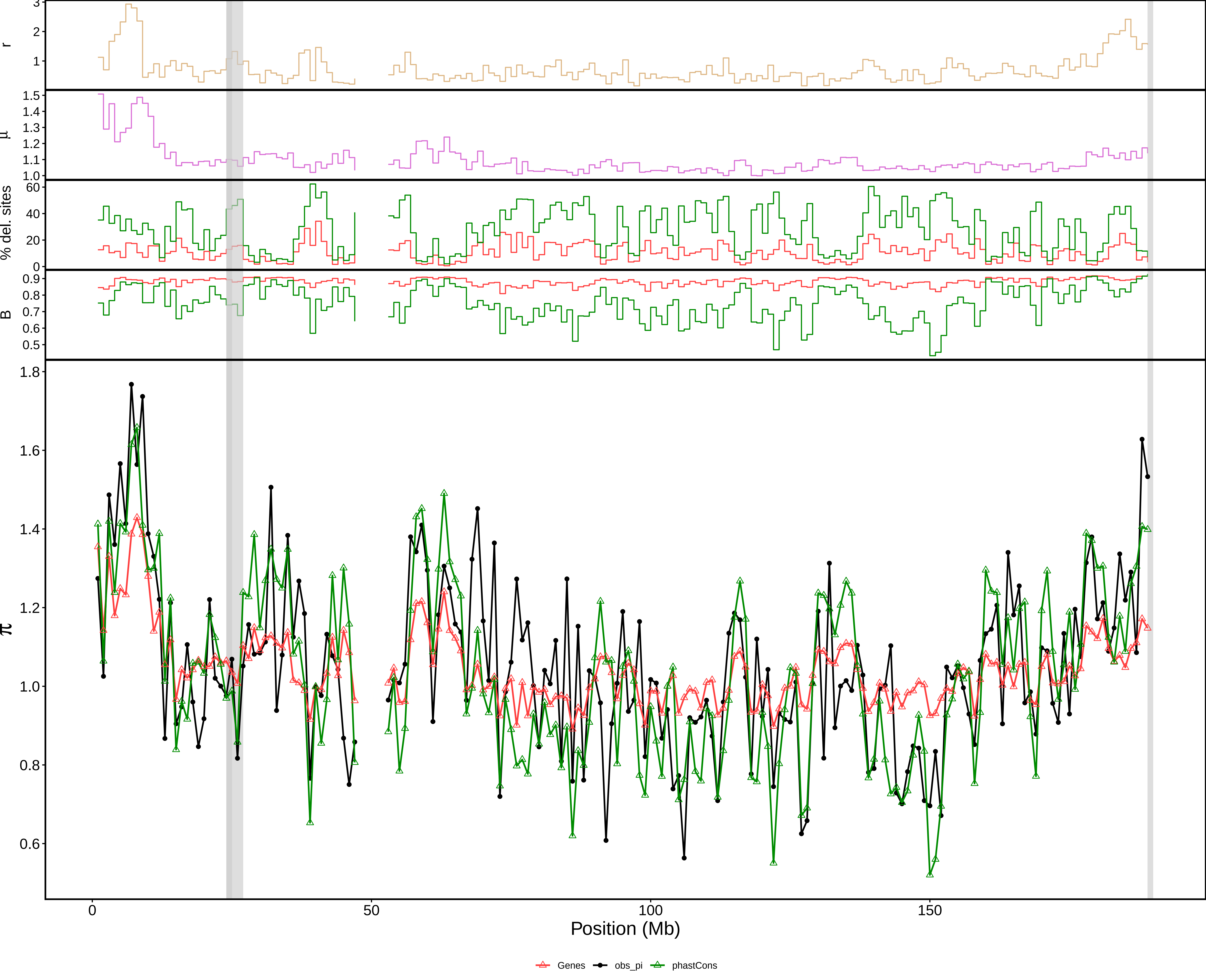

### aov_1Mb_chr5.pdf

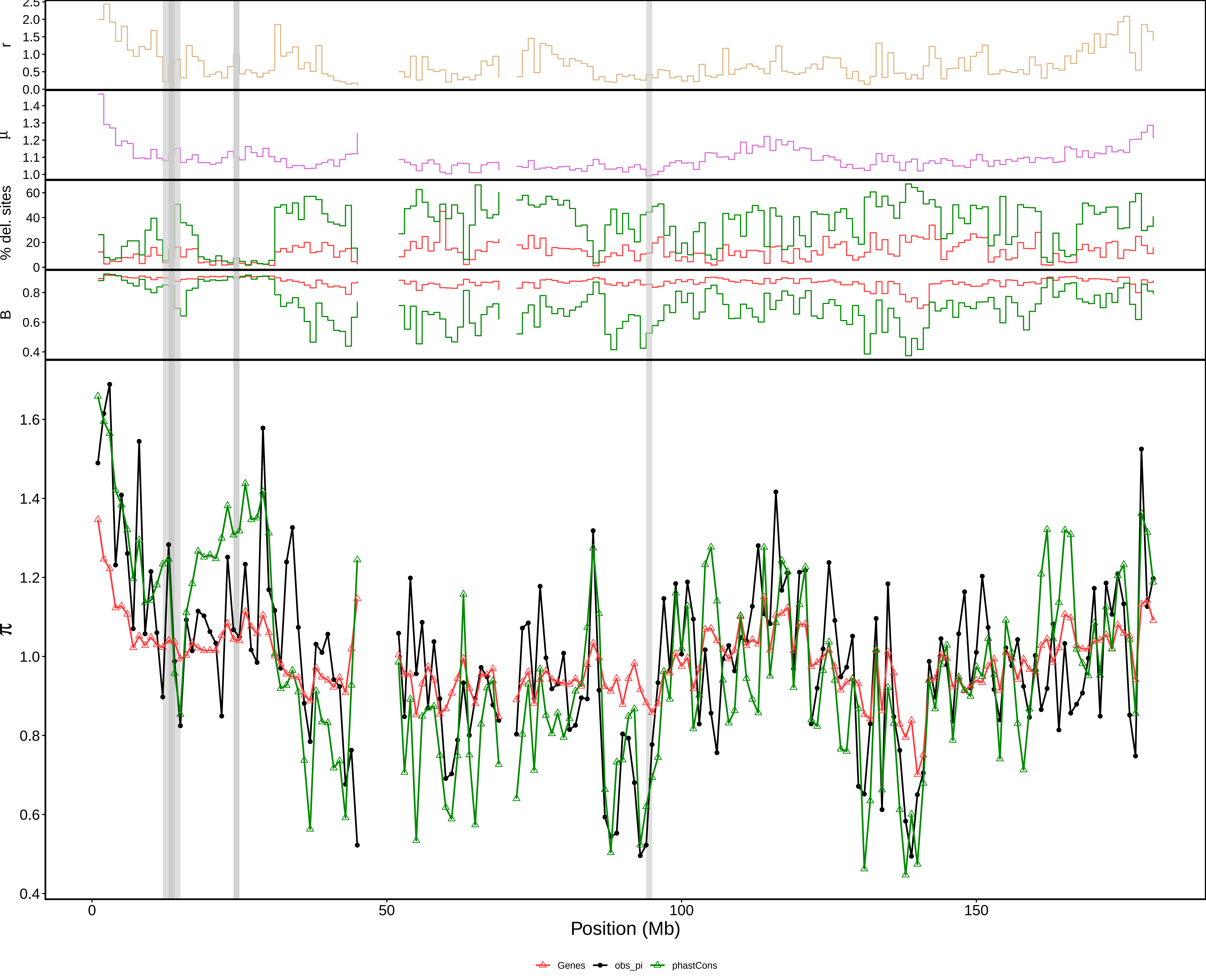

### aov_1Mb_chr6.pdf

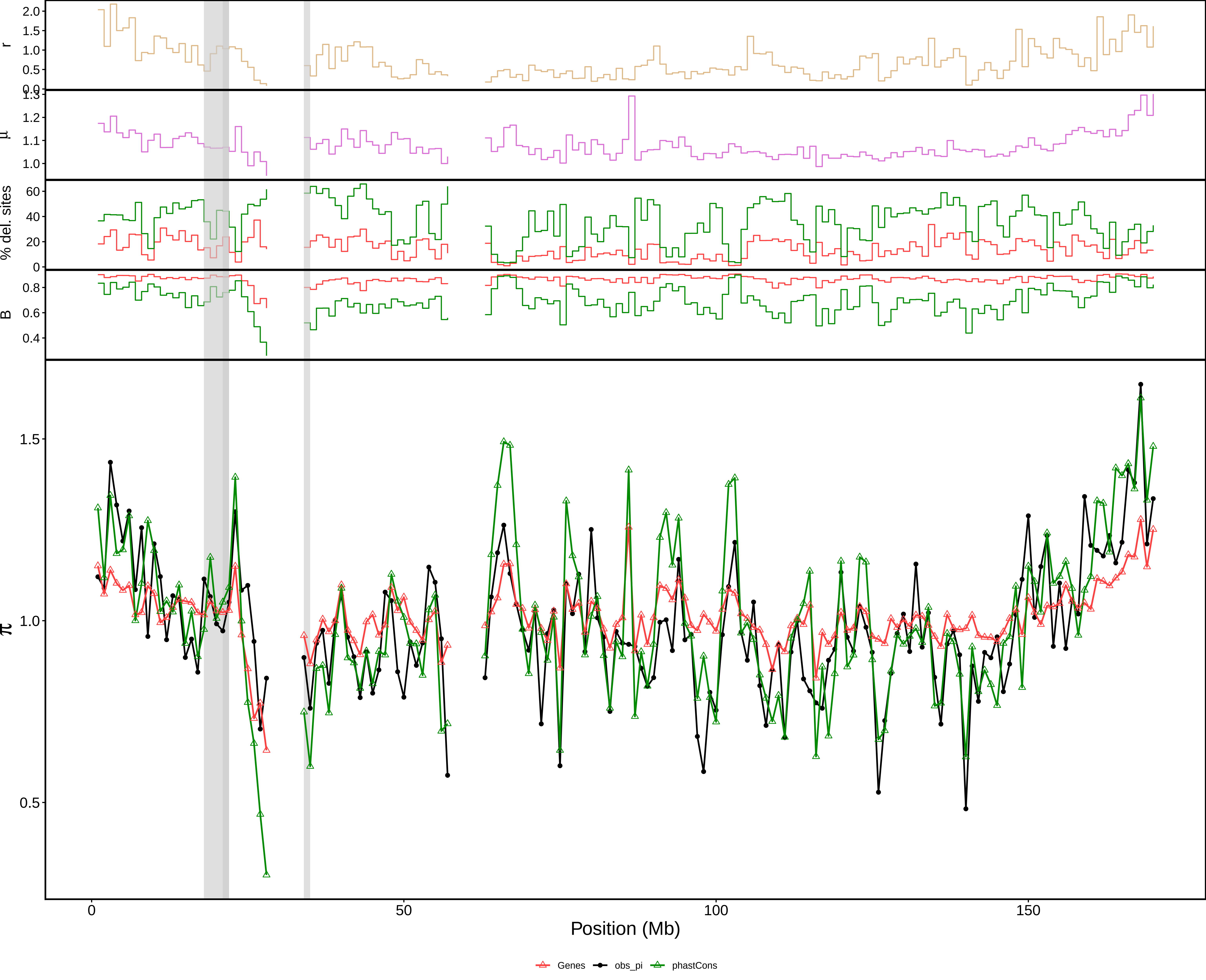

### aov_1Mb_chr7.pdf

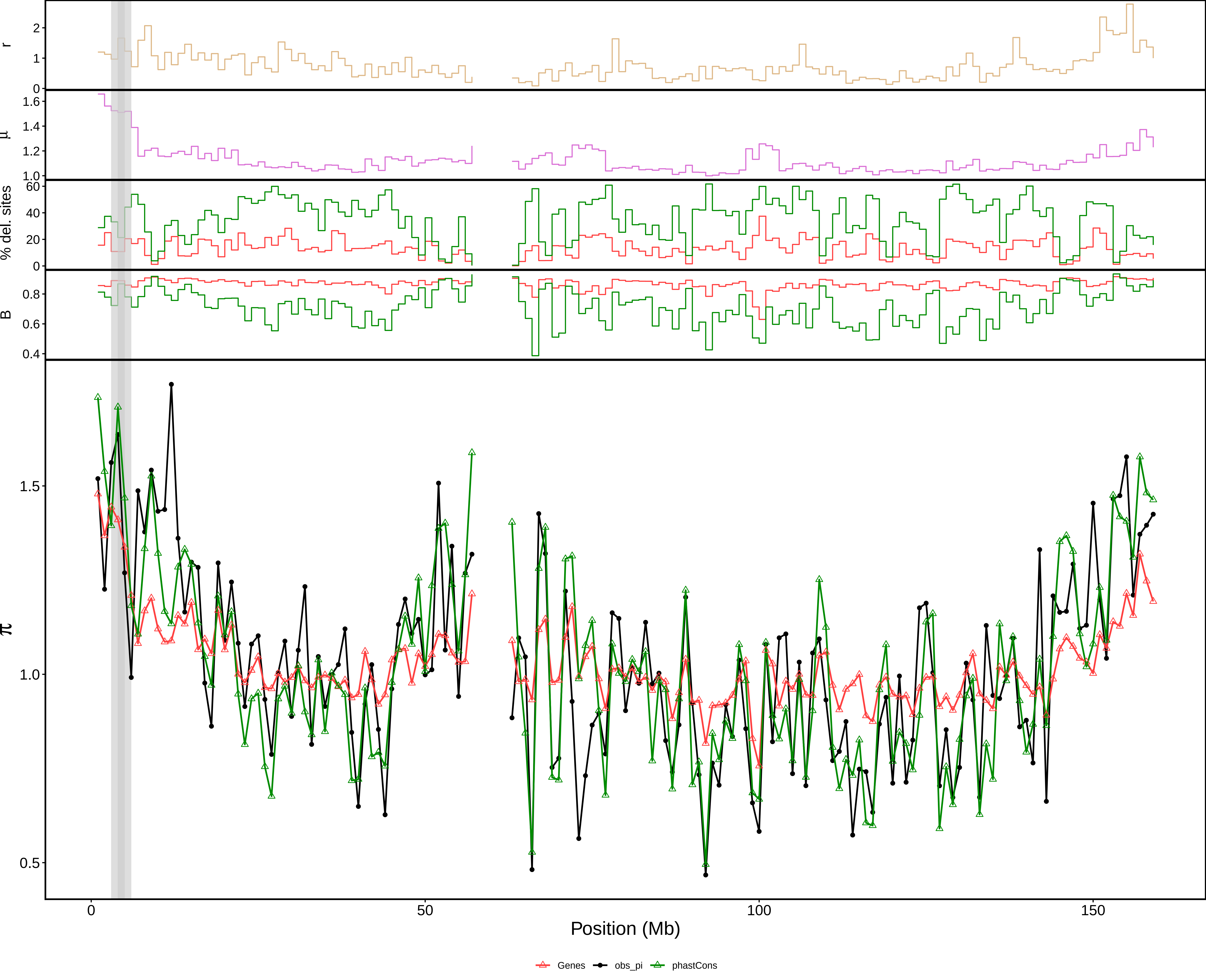

### aov_1Mb_chr8.pdf

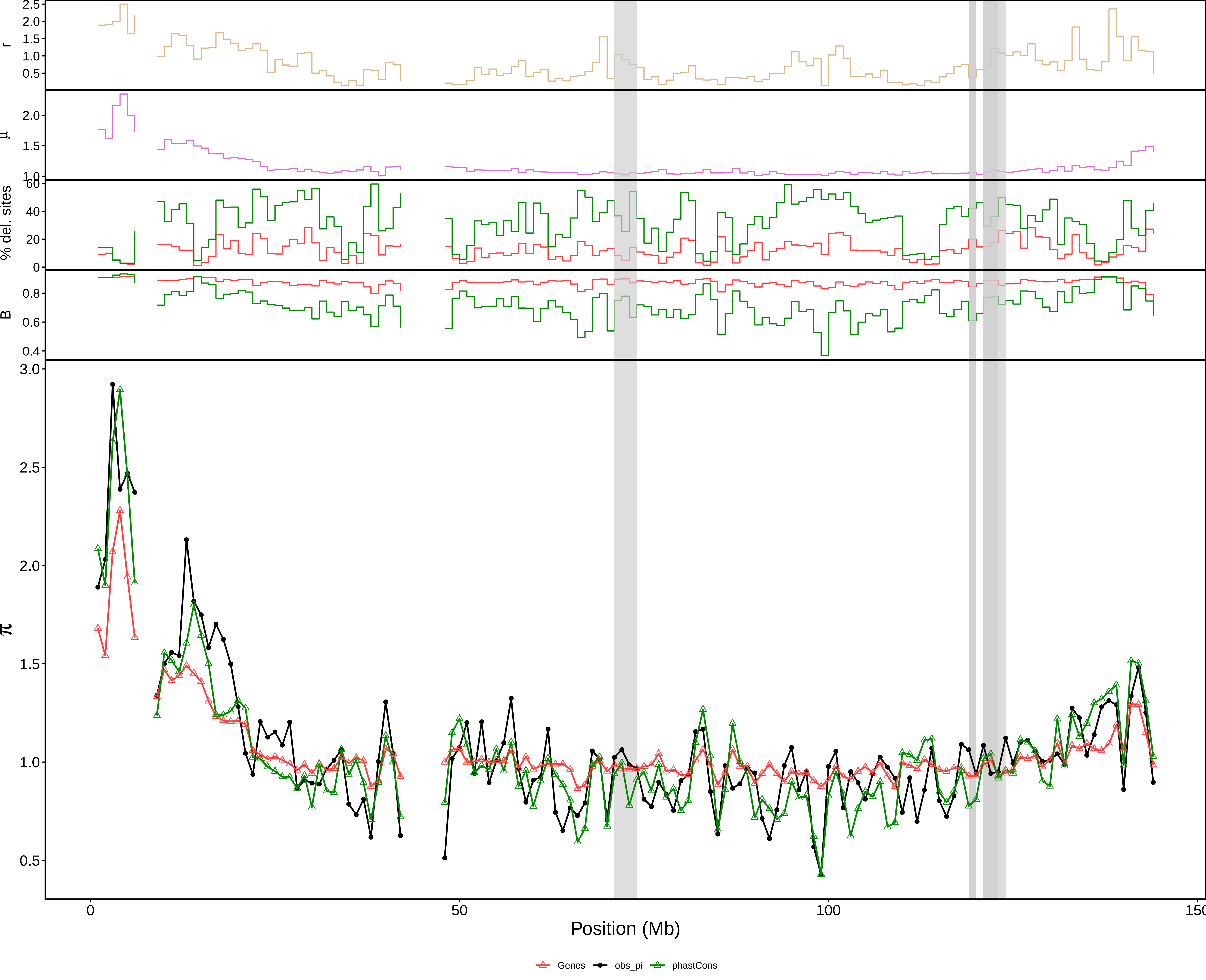

### aov_1Mb_chr9.pdf

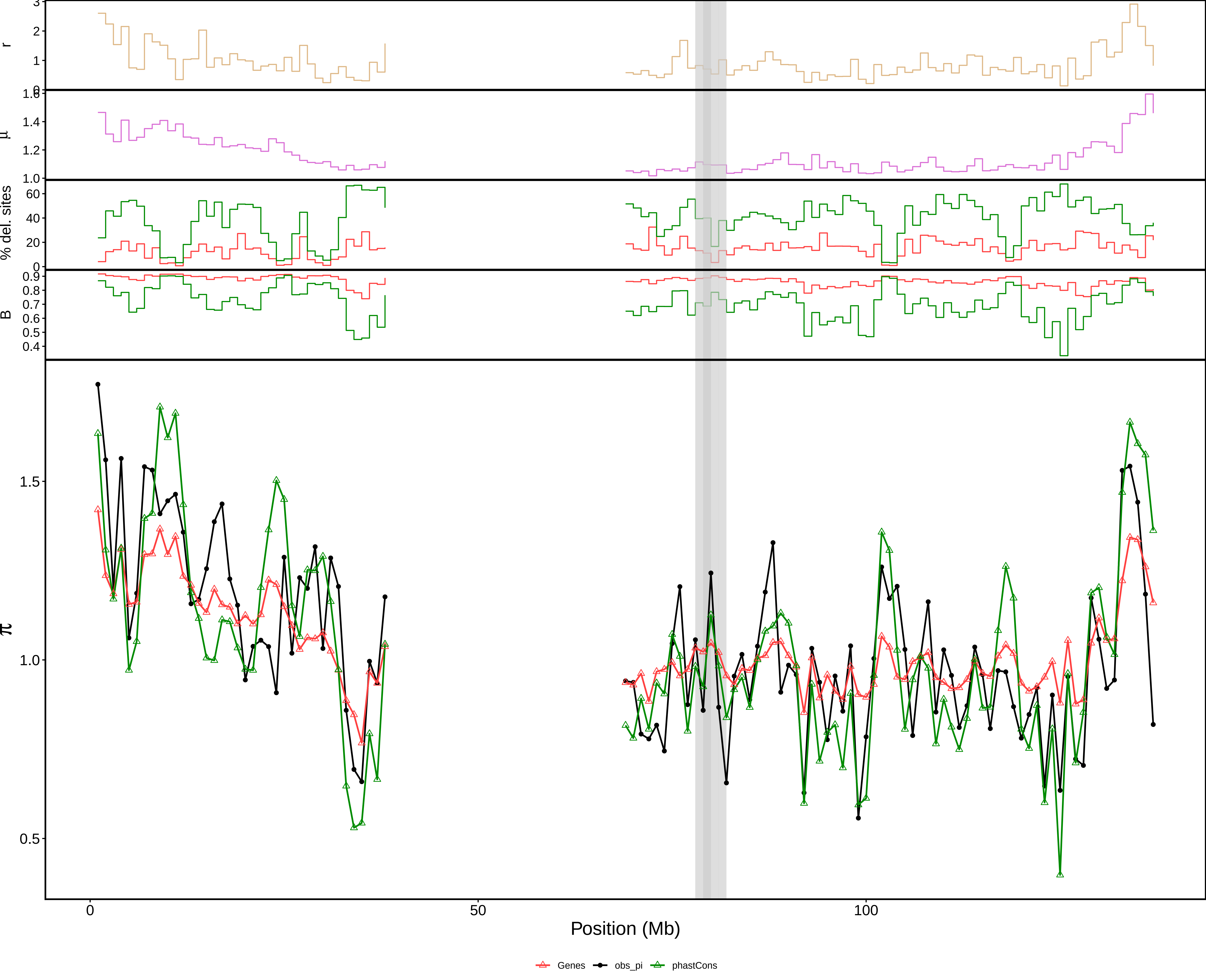

### aov_1Mb_chr10.pdf

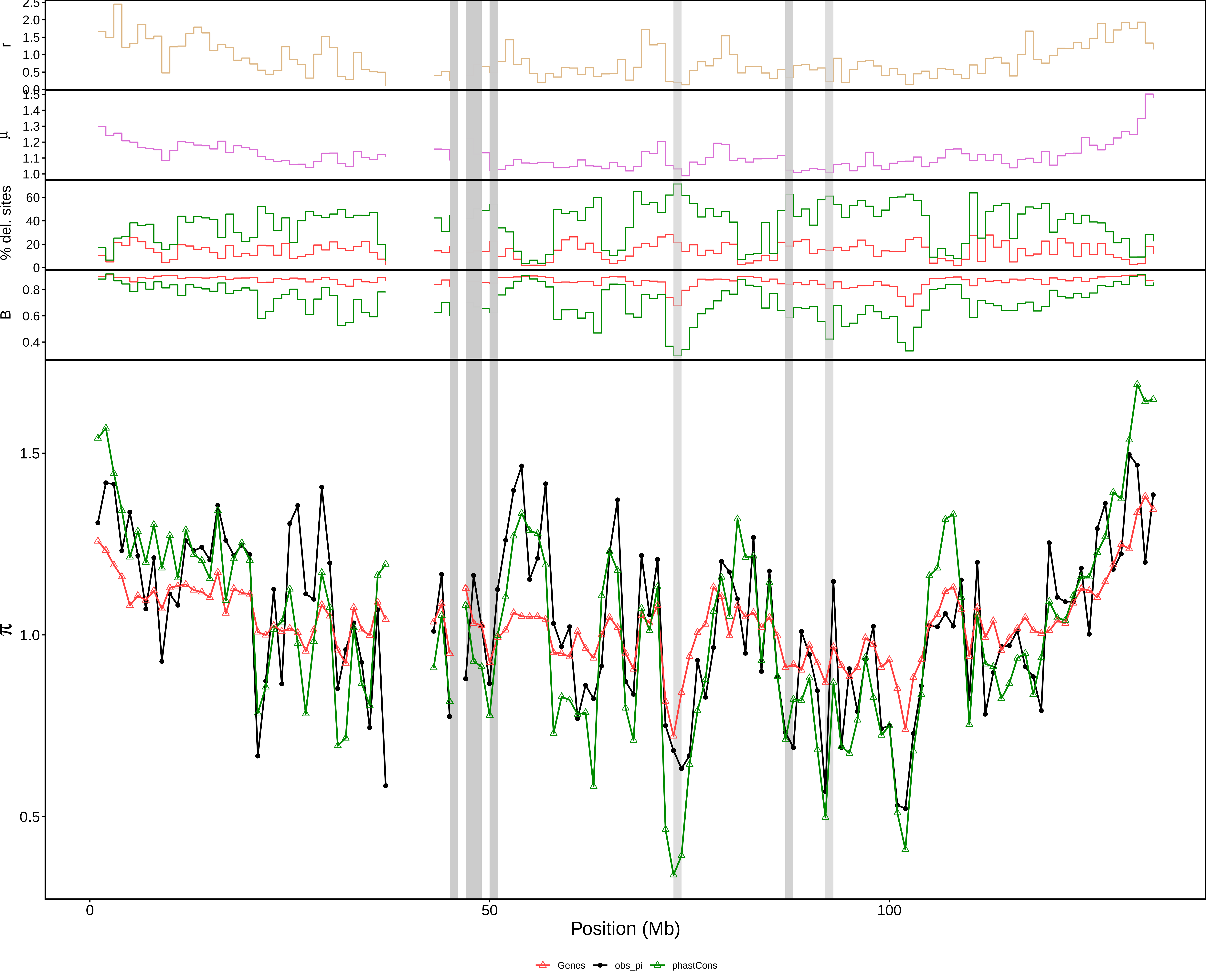

### aov_1Mb_chr11.pdf

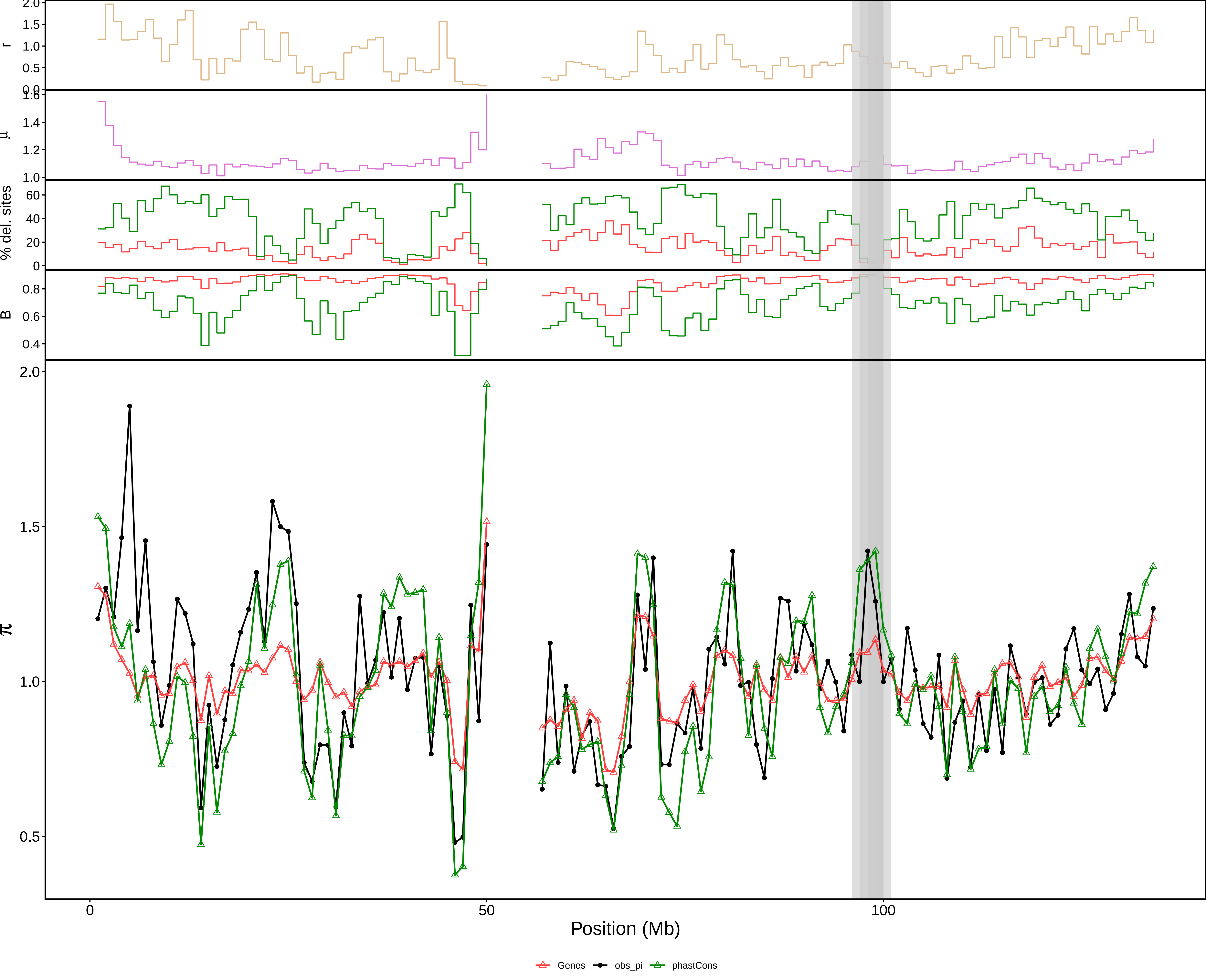

### aov_1Mb_chr12.pdf

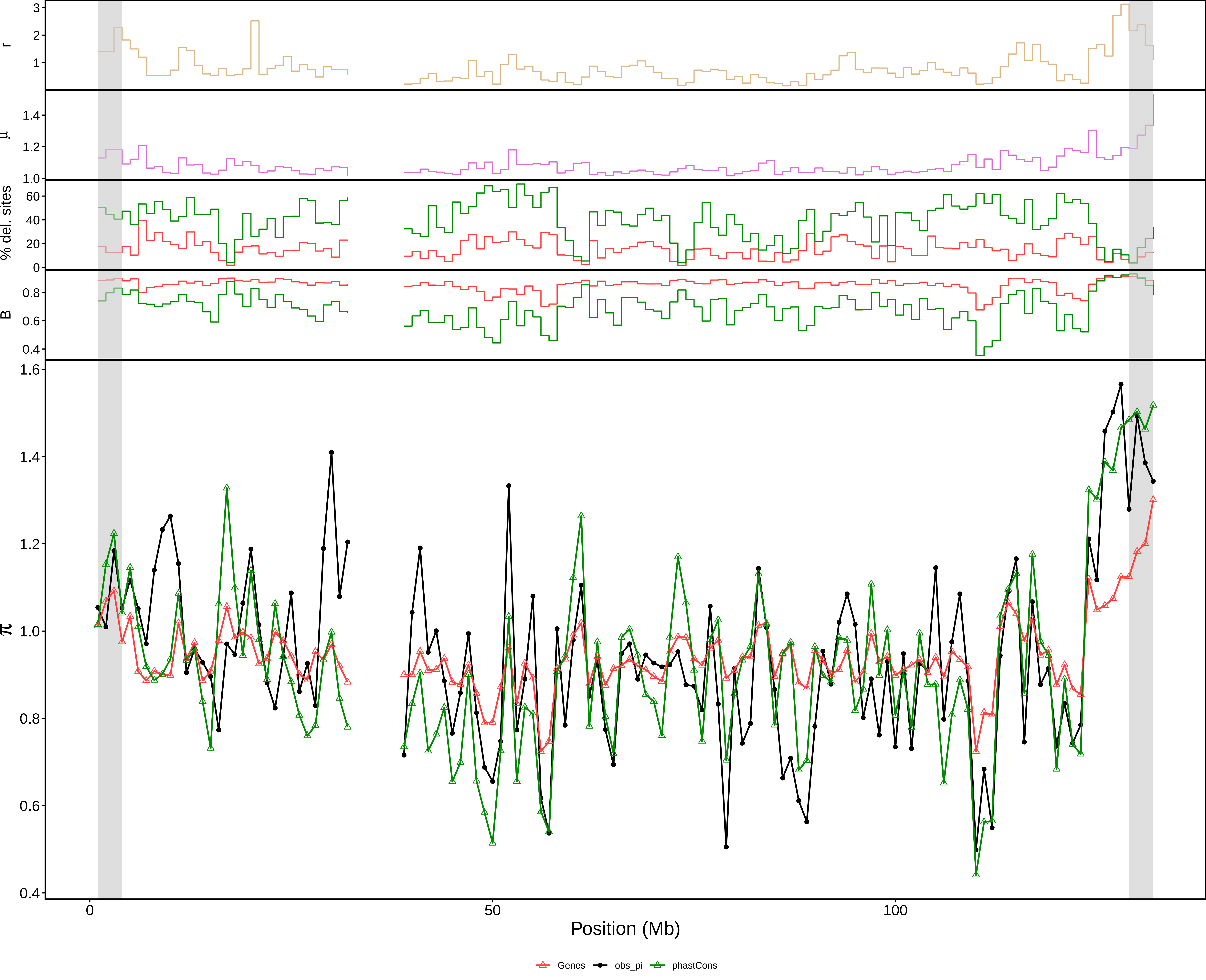

### aov_1Mb_chr13.pdf

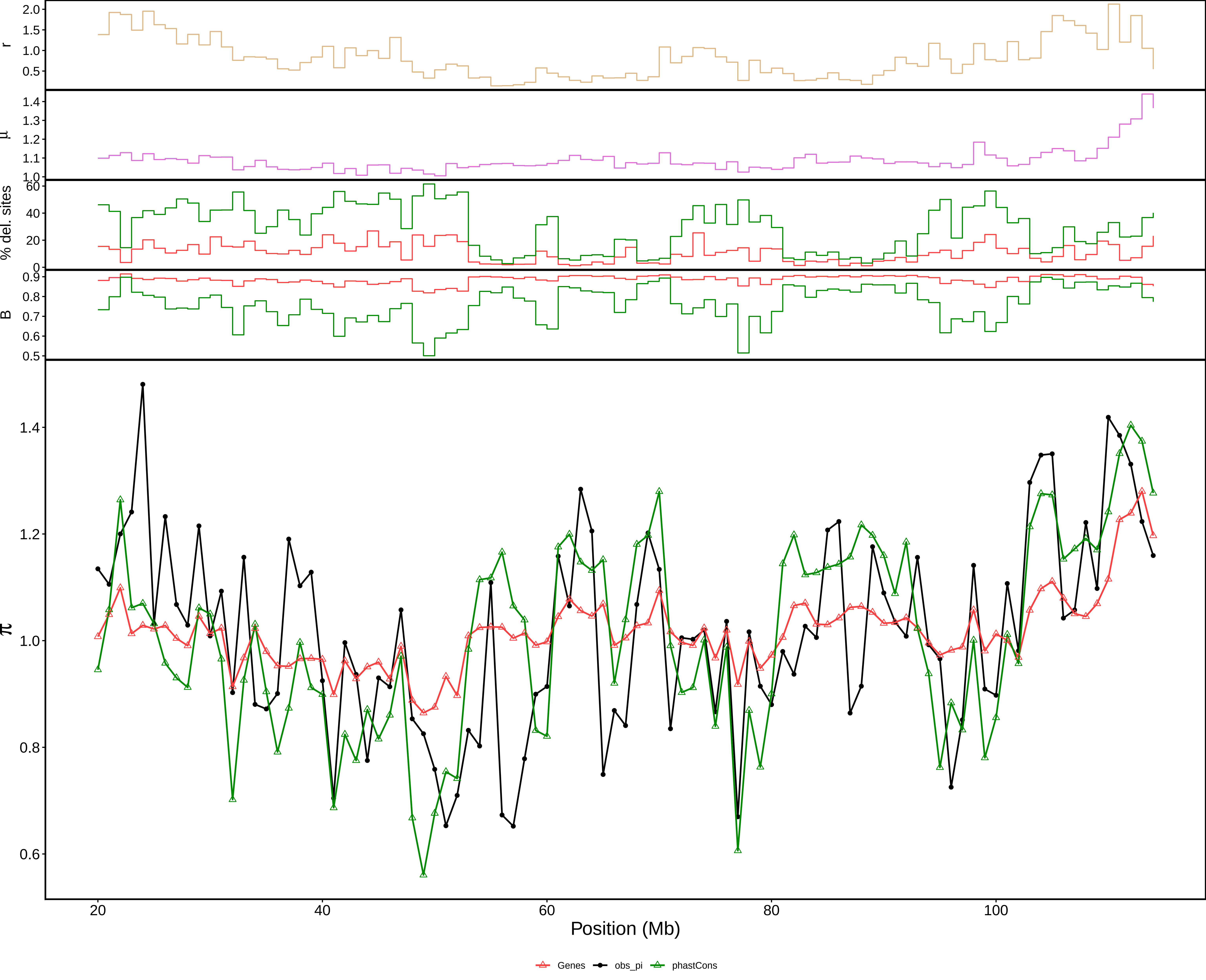

### aov_1Mb_chr14.pdf

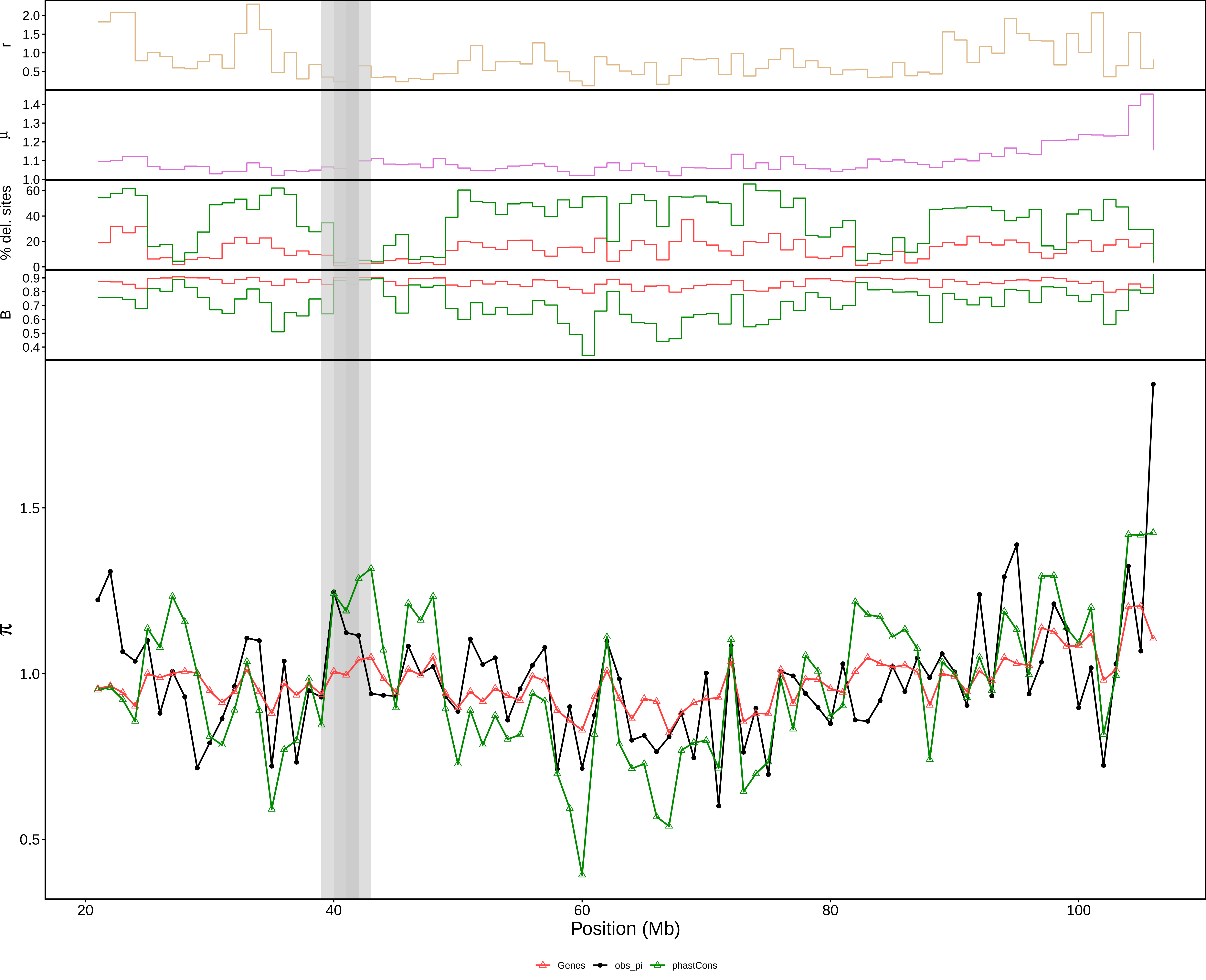

### aov_1Mb_chr15.pdf

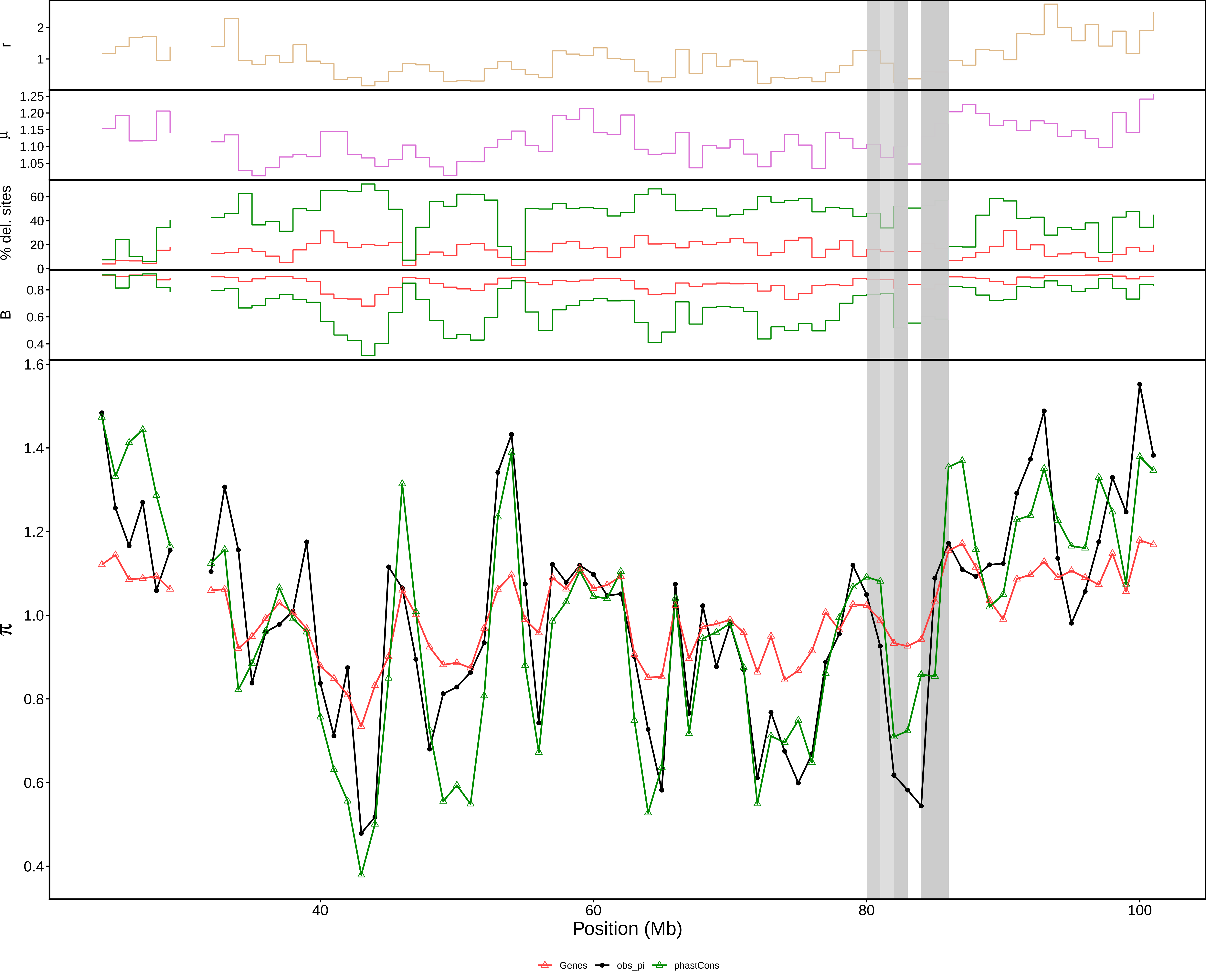

### aov_1Mb_chr16.pdf

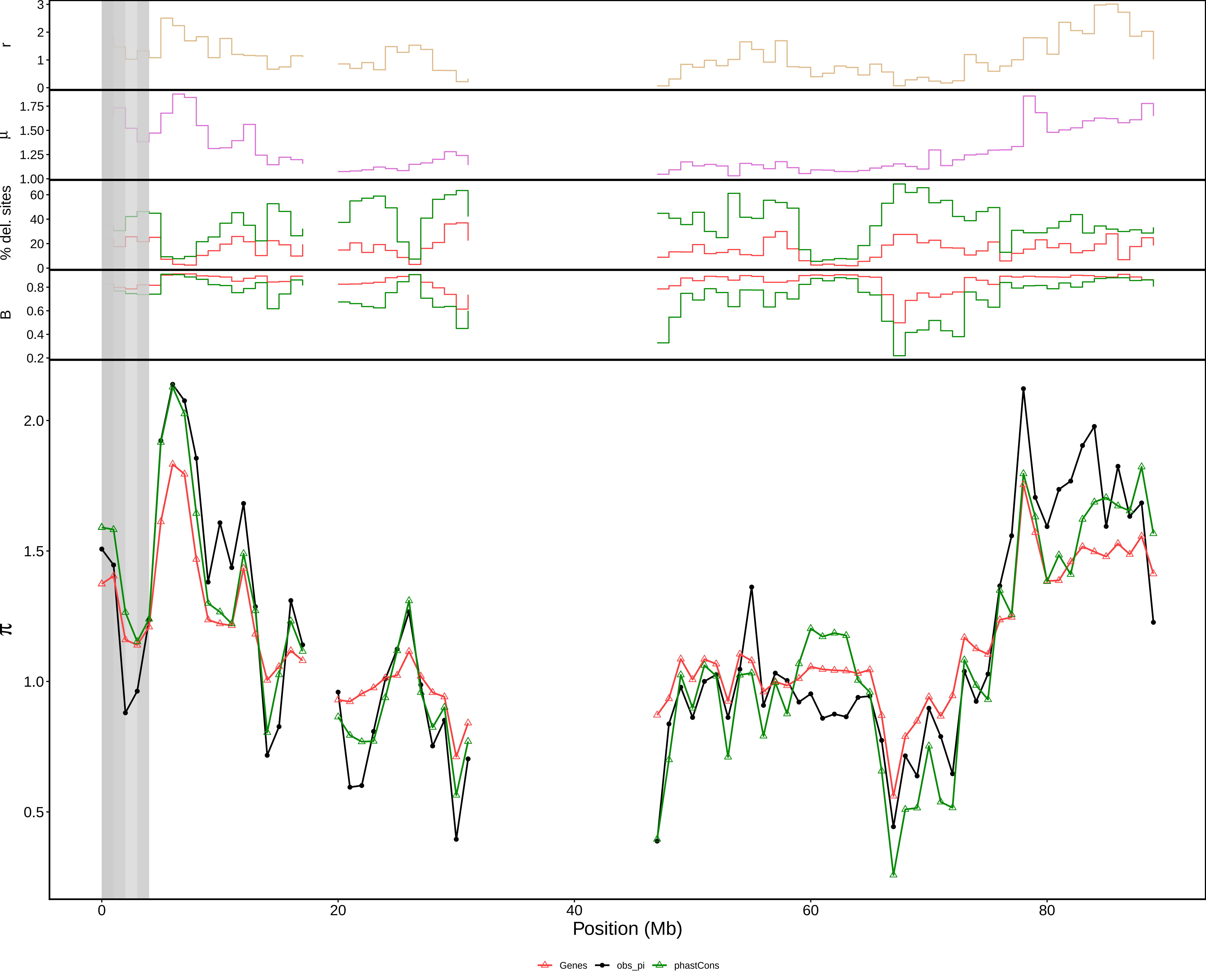

### aov_1Mb_chr17.pdf

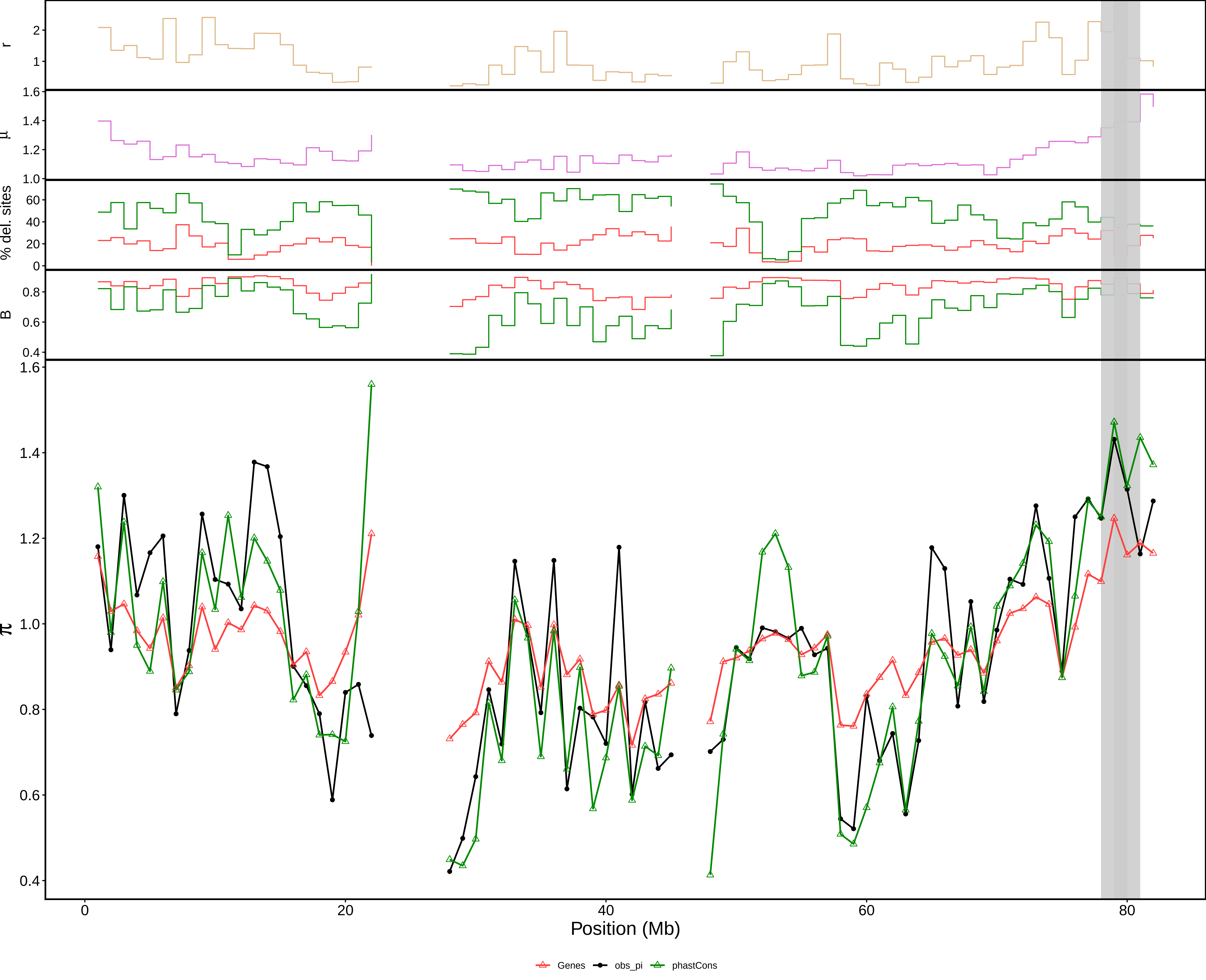

### aov_1Mb_chr18.pdf

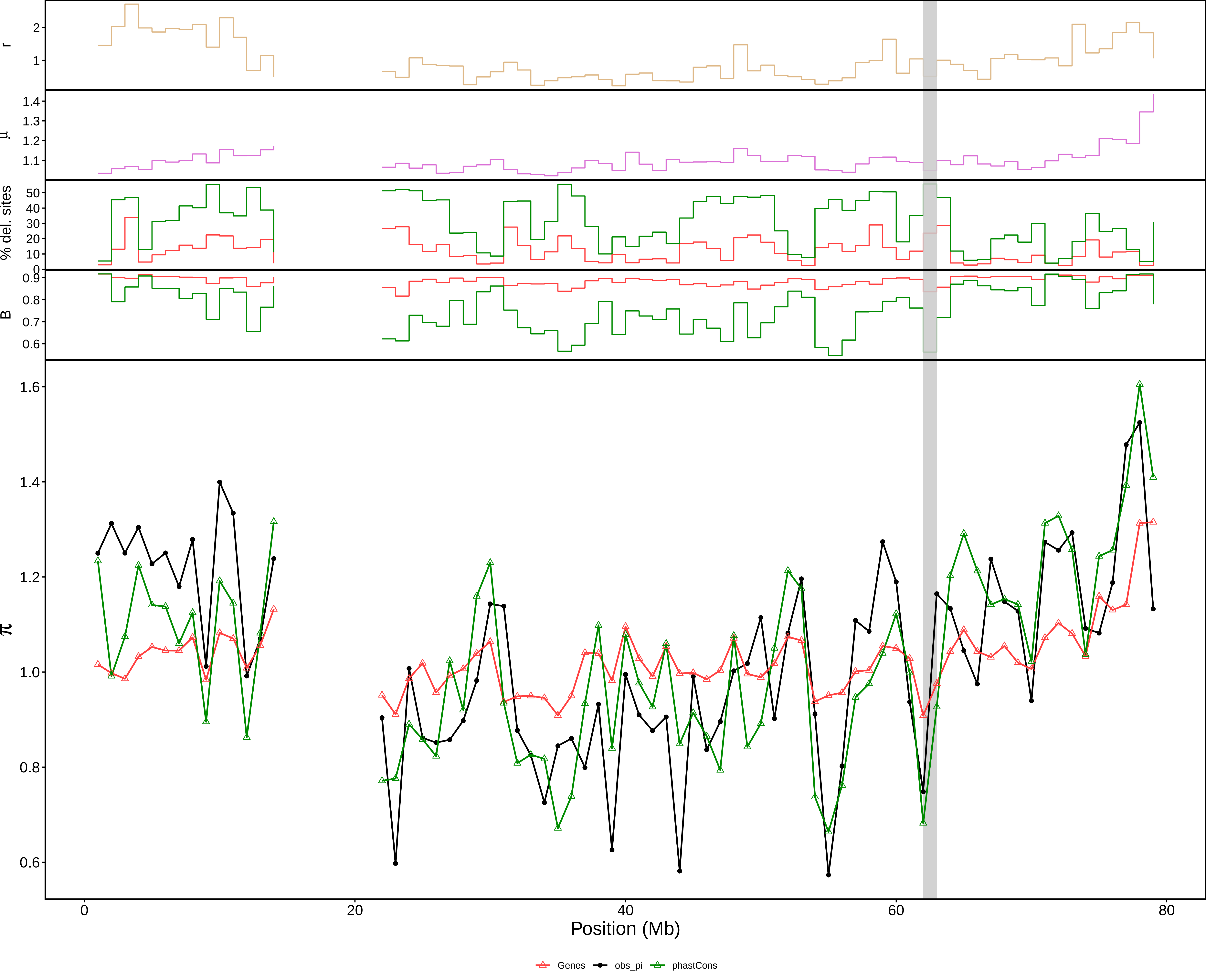

### aov_1Mb_chr19.pdf

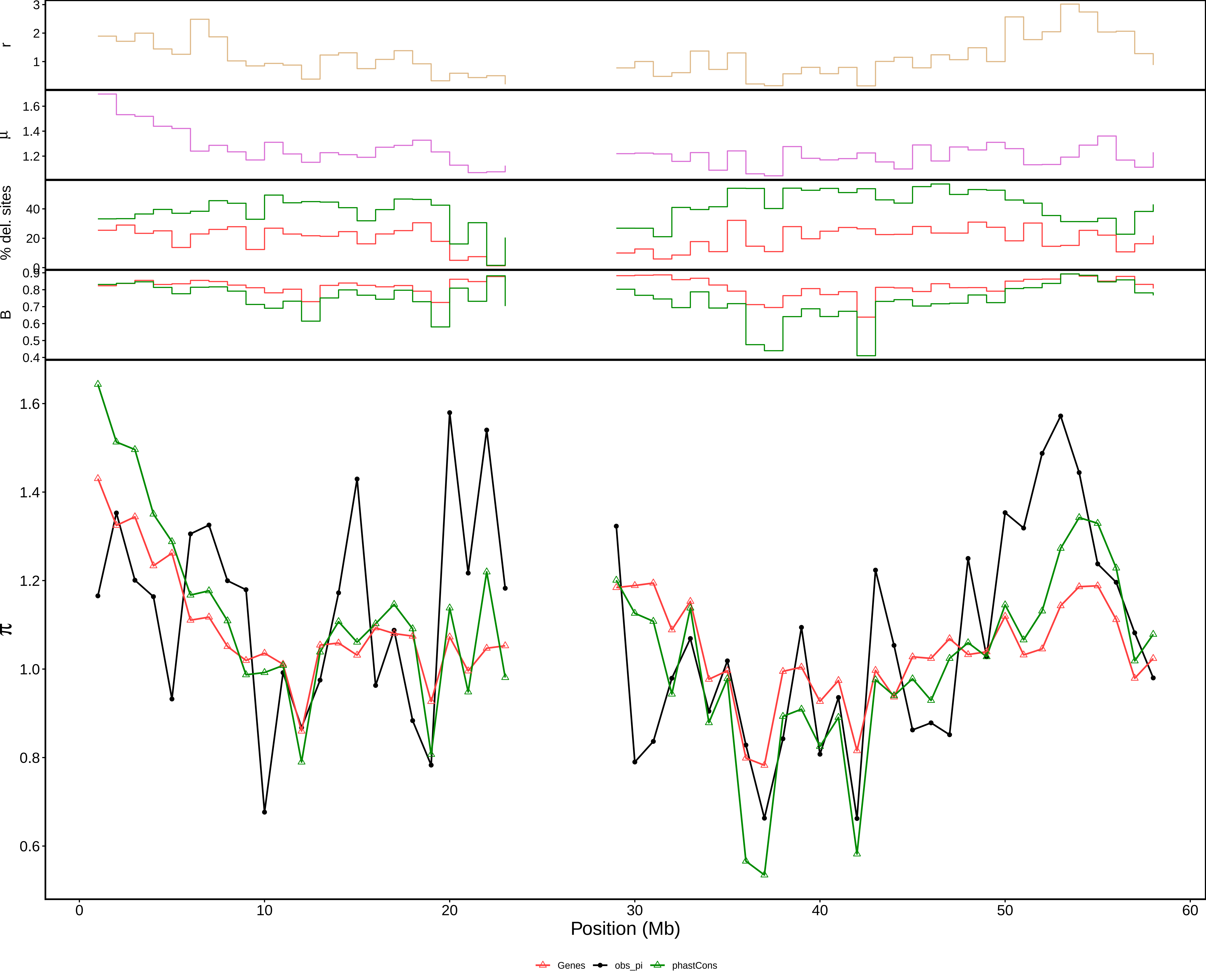

### aov_1Mb_chr20.pdf

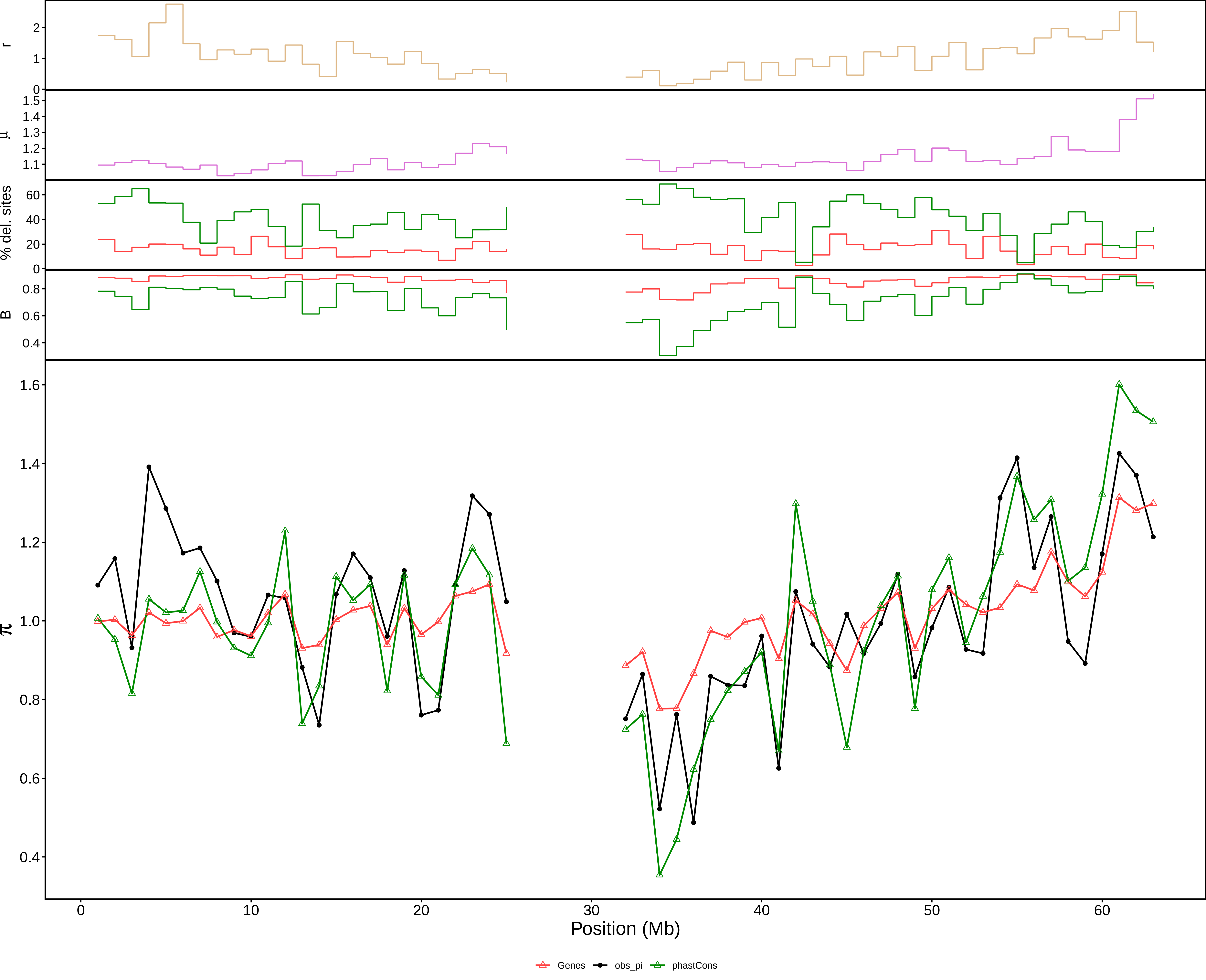

### aov_1Mb_chr21.pdf

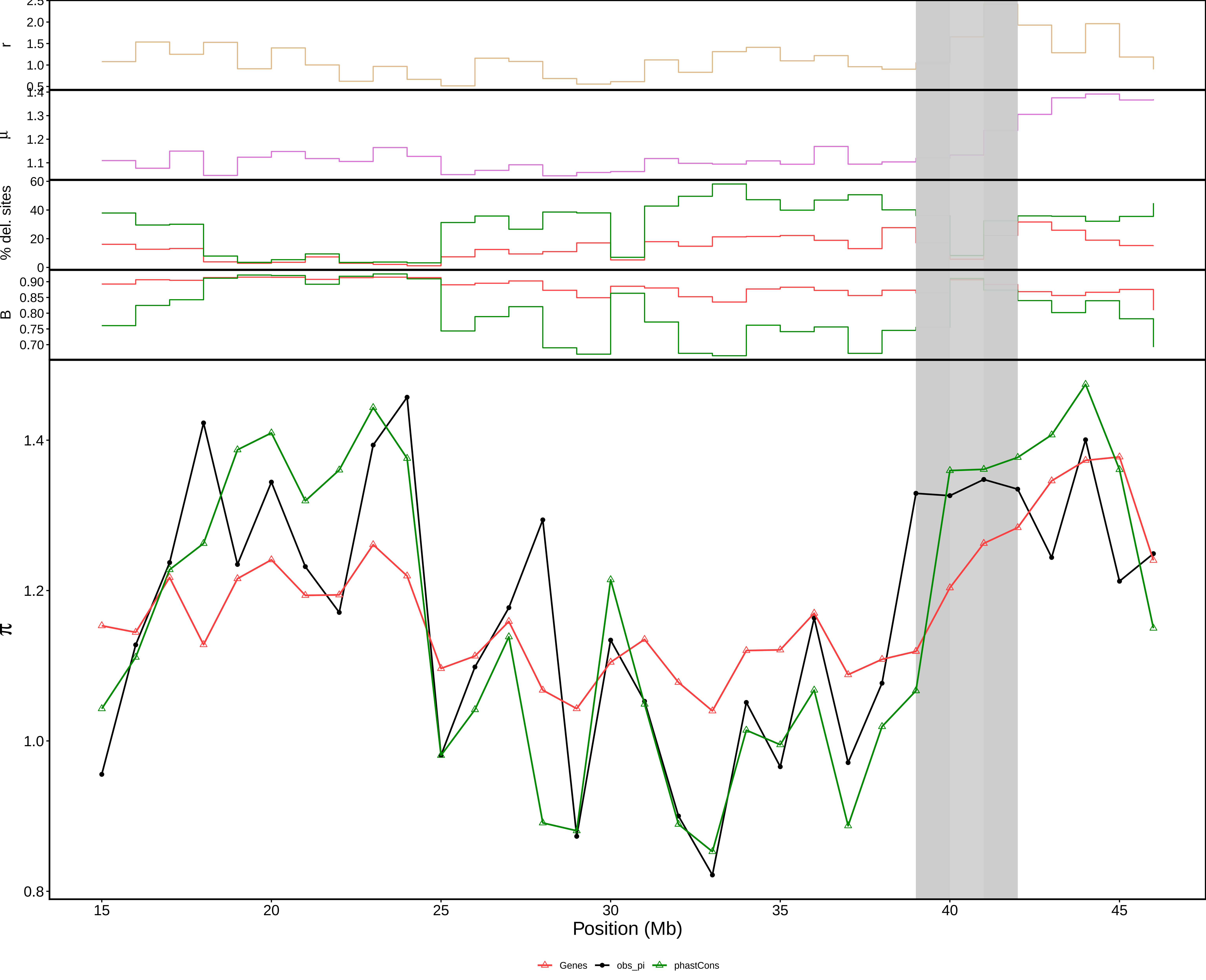

### aov_1Mb_chr22.pdf

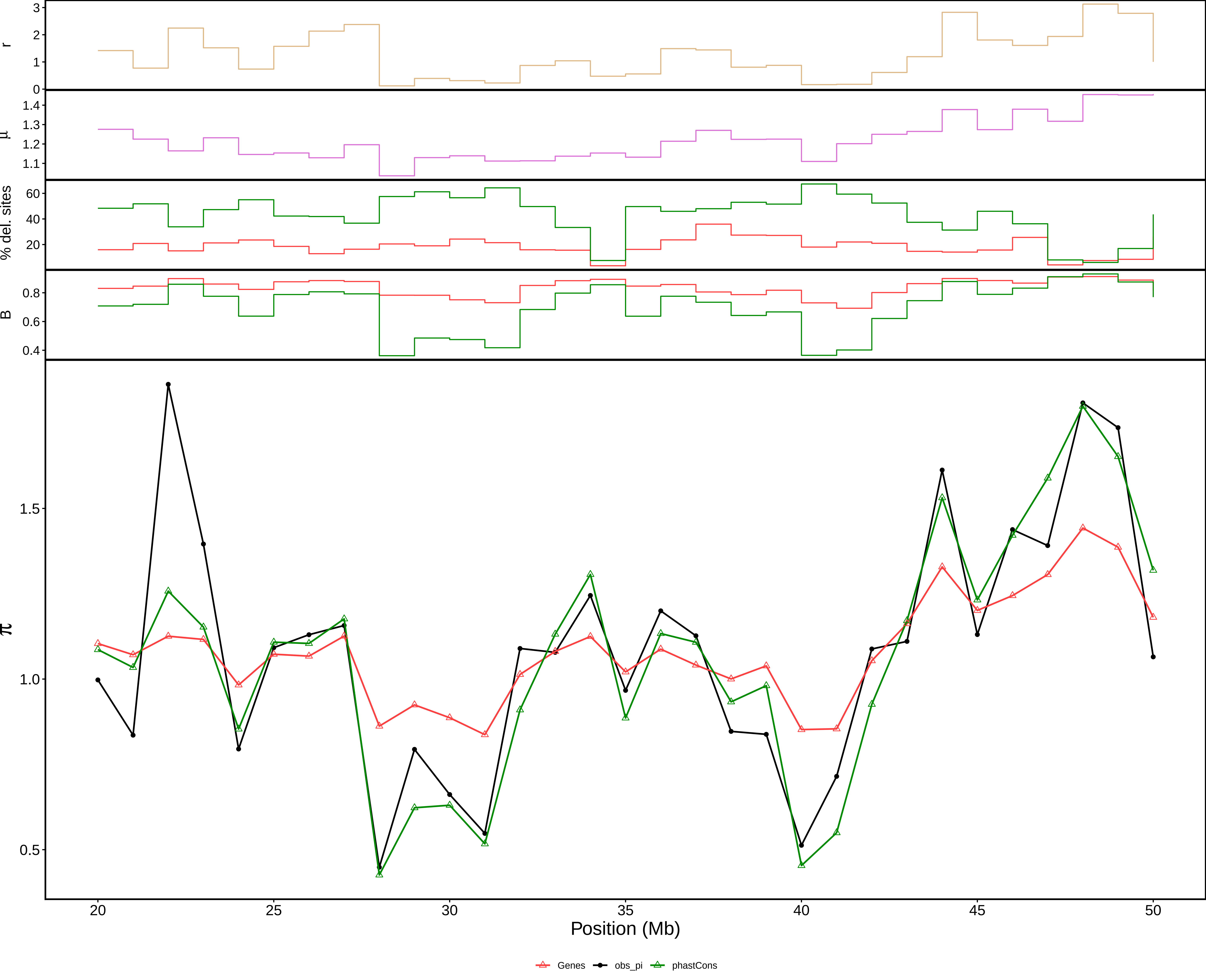

### pi_c1_1Mb.pdf

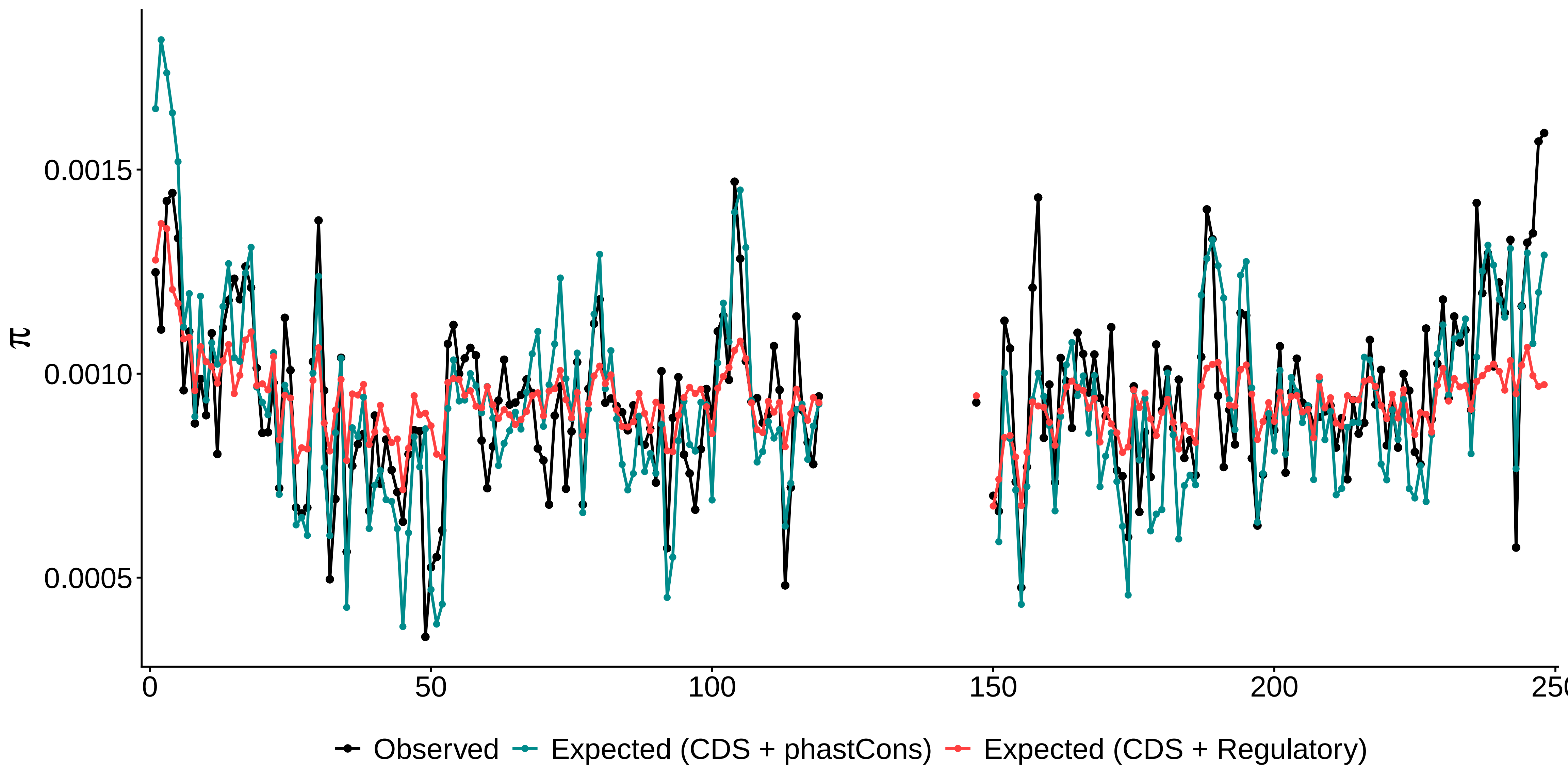

### pi_c2_1Mb.pdf

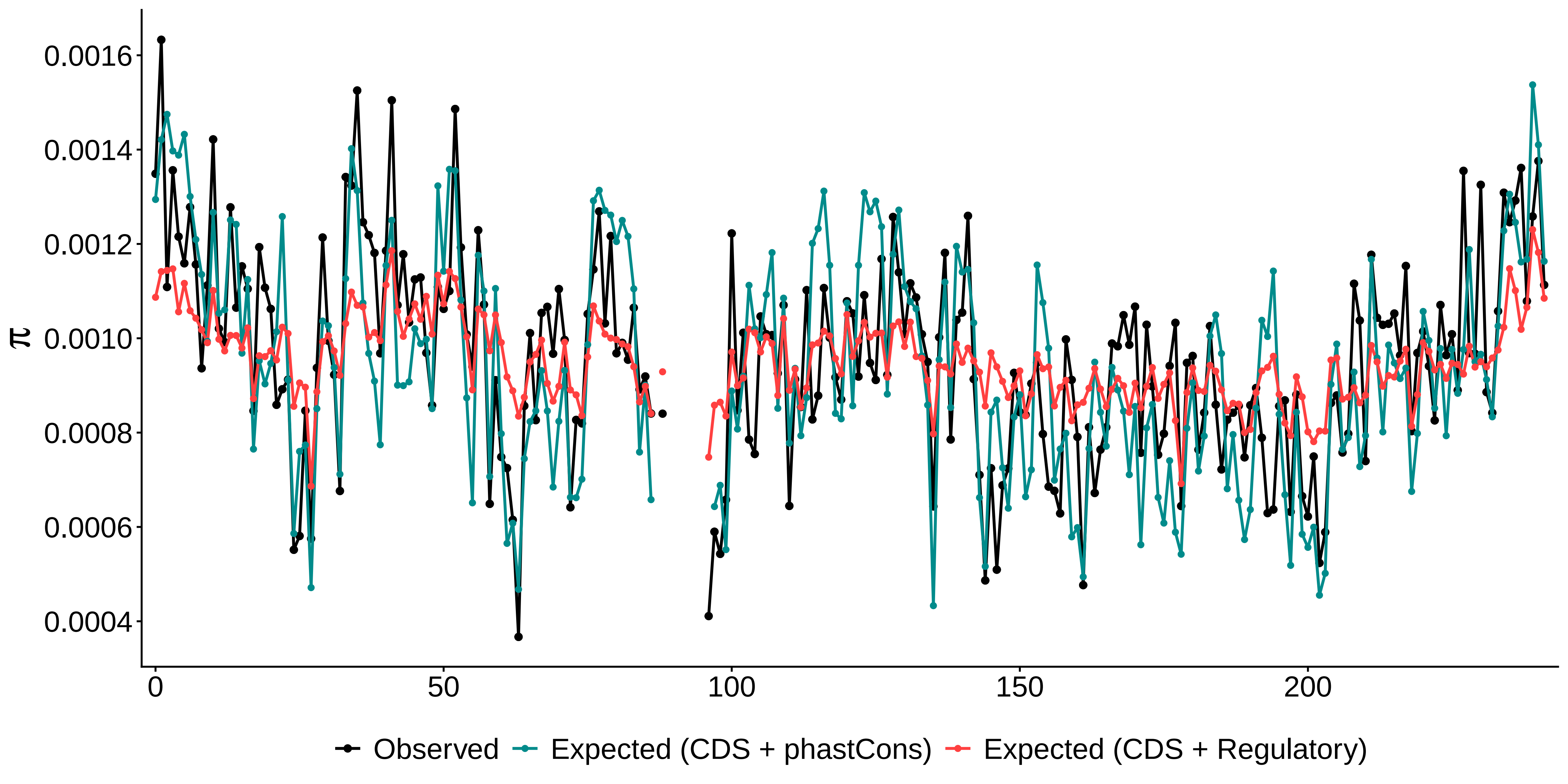

### pi_c3_1Mb.pdf

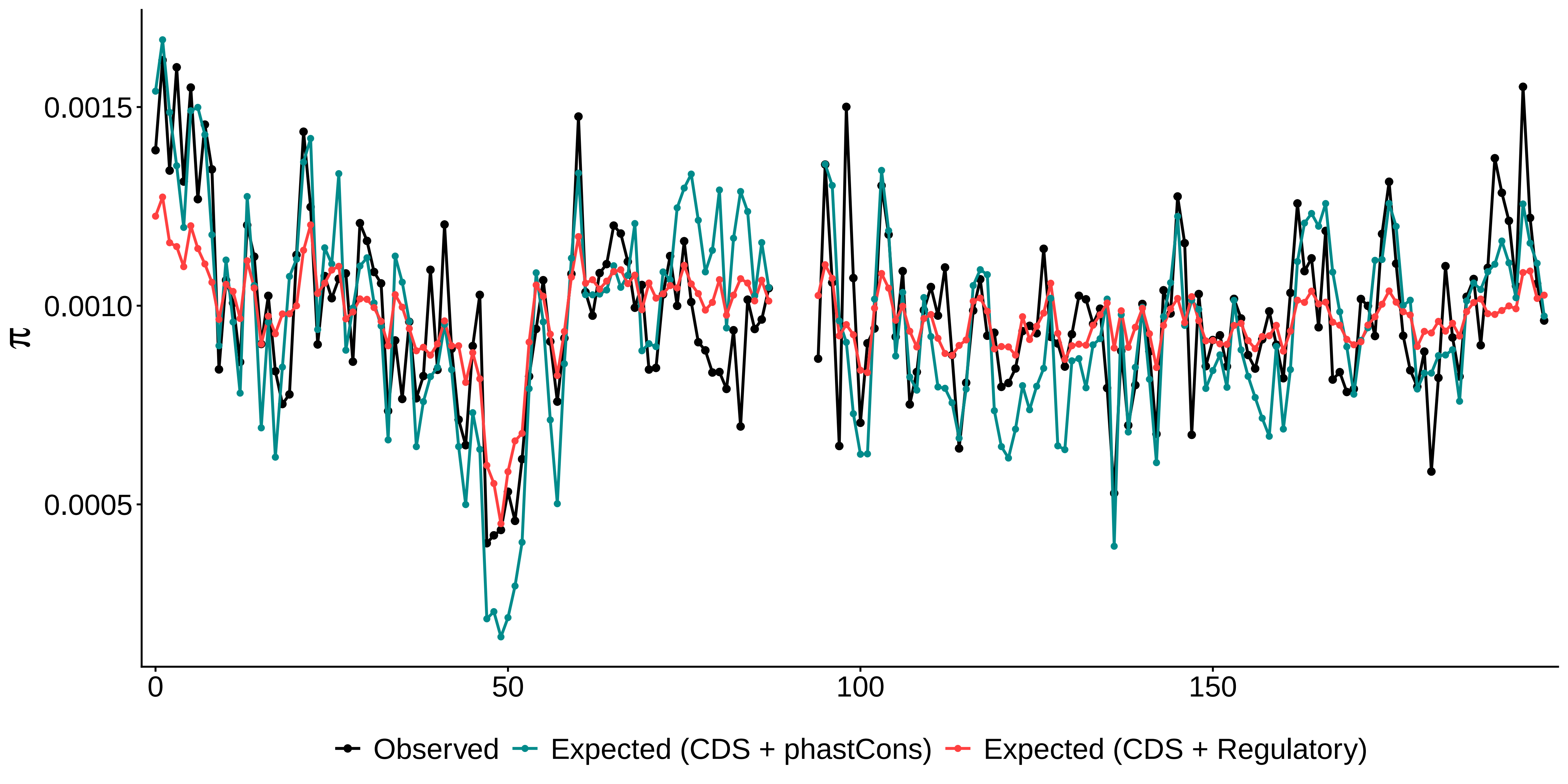

### pi_c4_1Mb.pdf

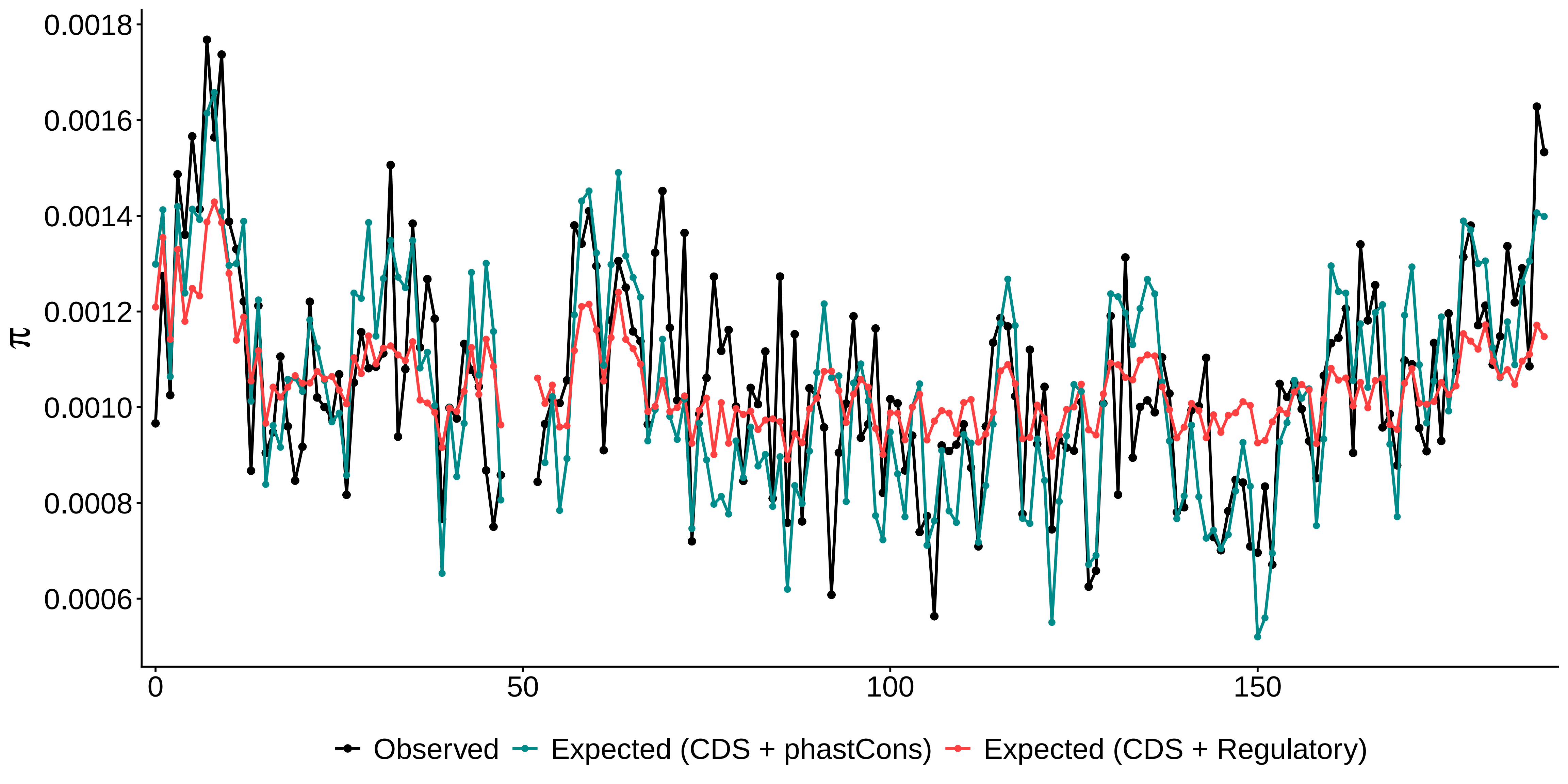

### pi_c5_1Mb.pdf

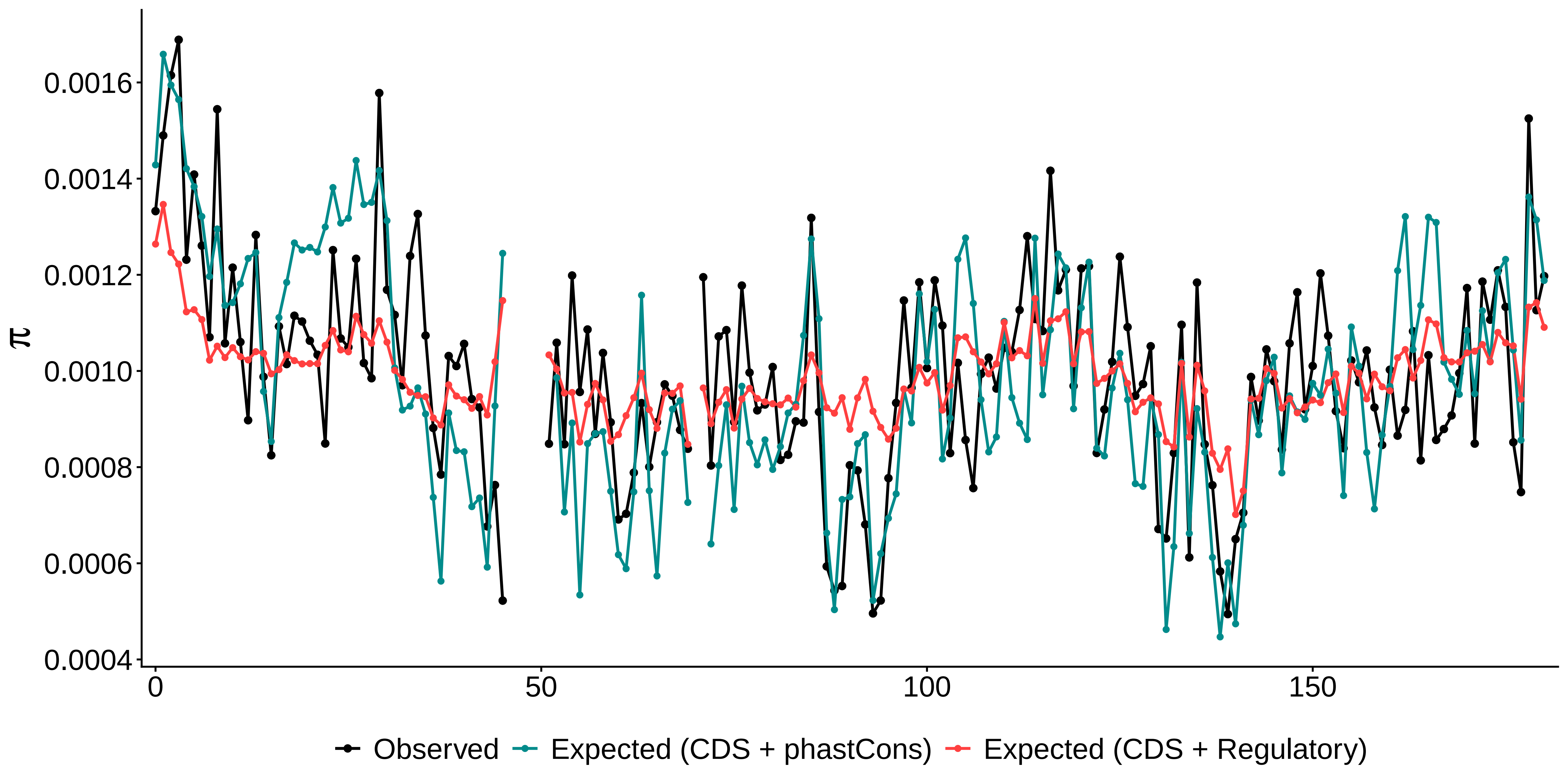

### pi_c6_1Mb.pdf

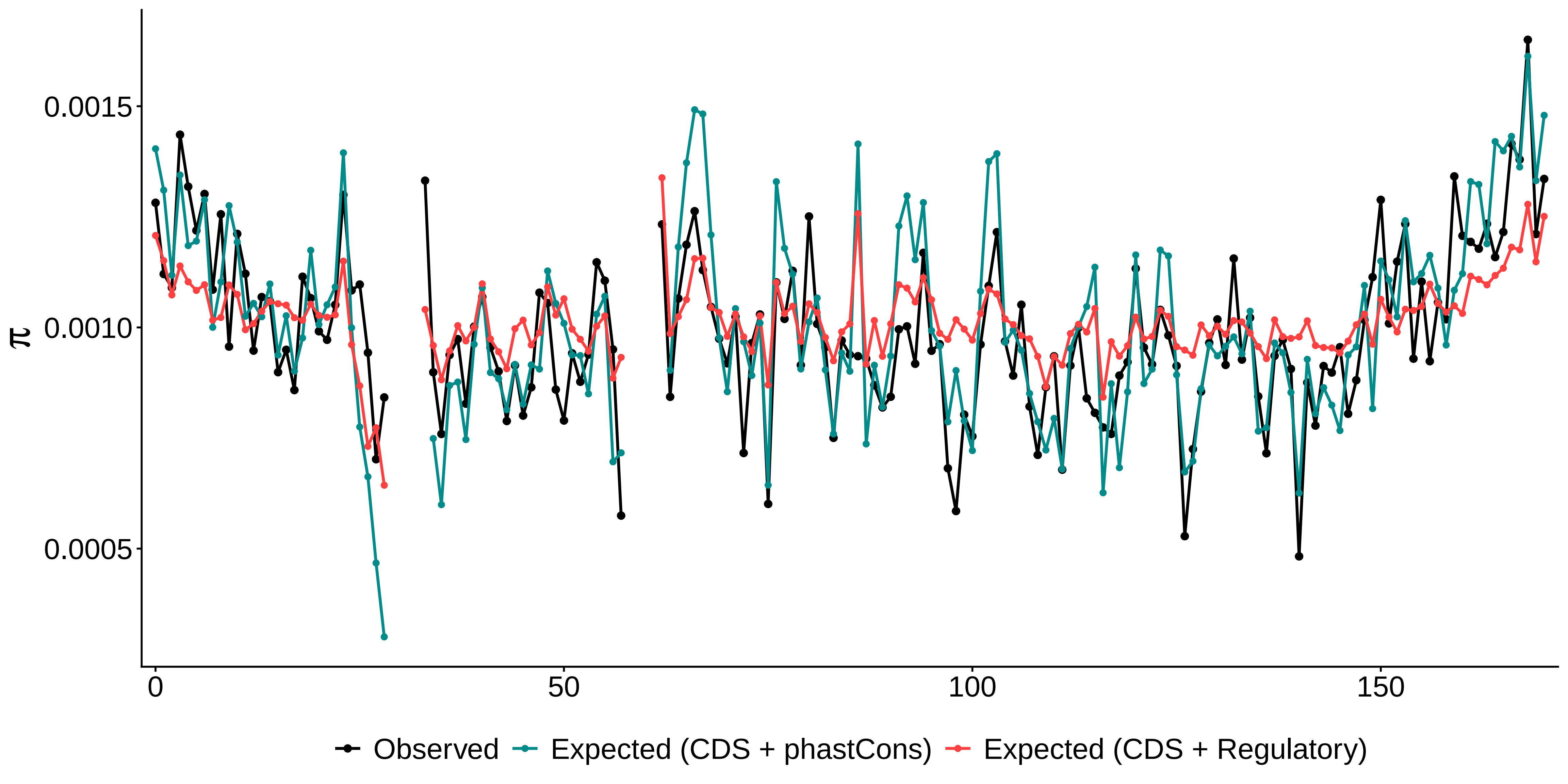

### pi_c7_1Mb.pdf

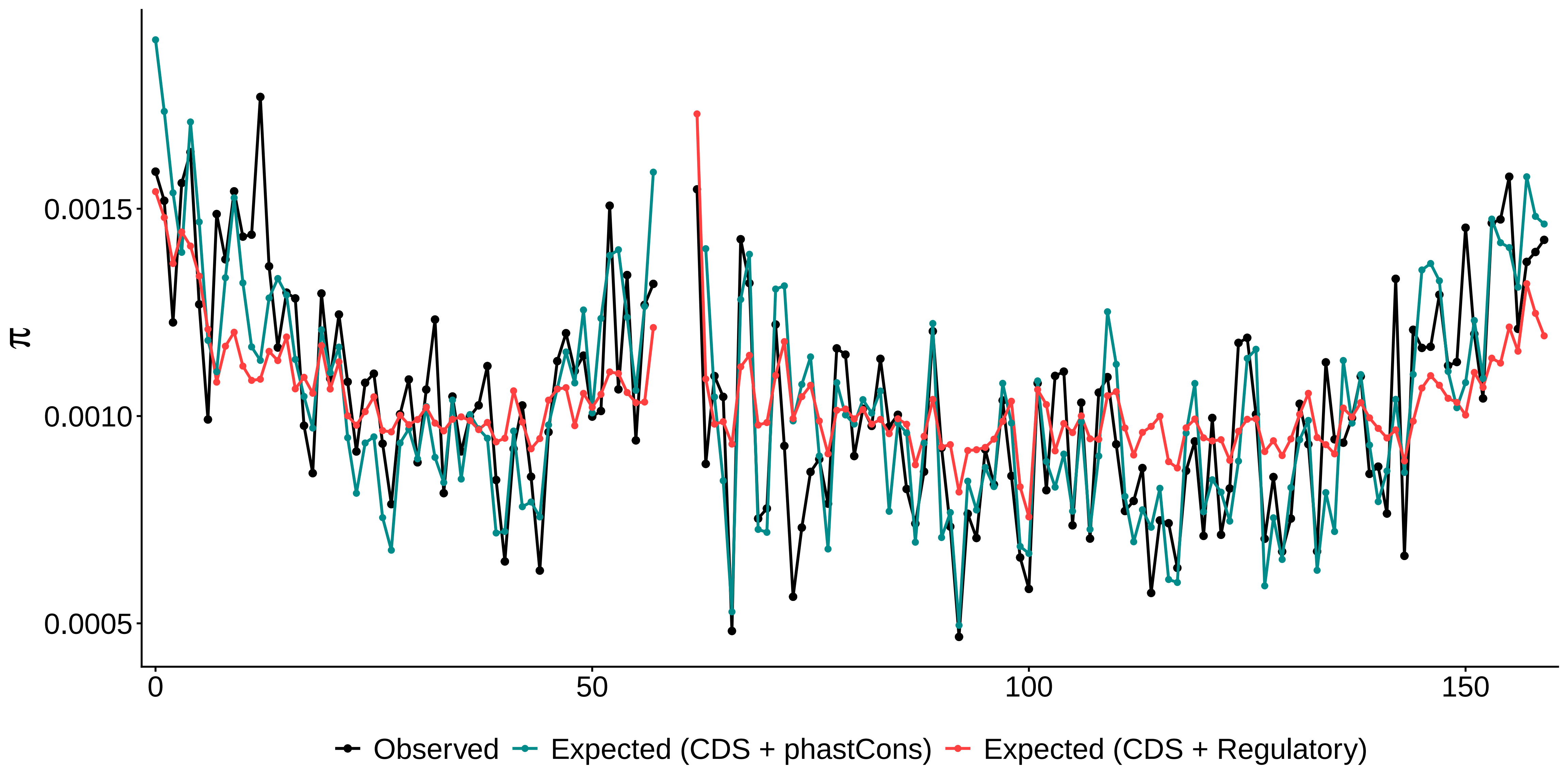

### pi_c8_1Mb.pdf

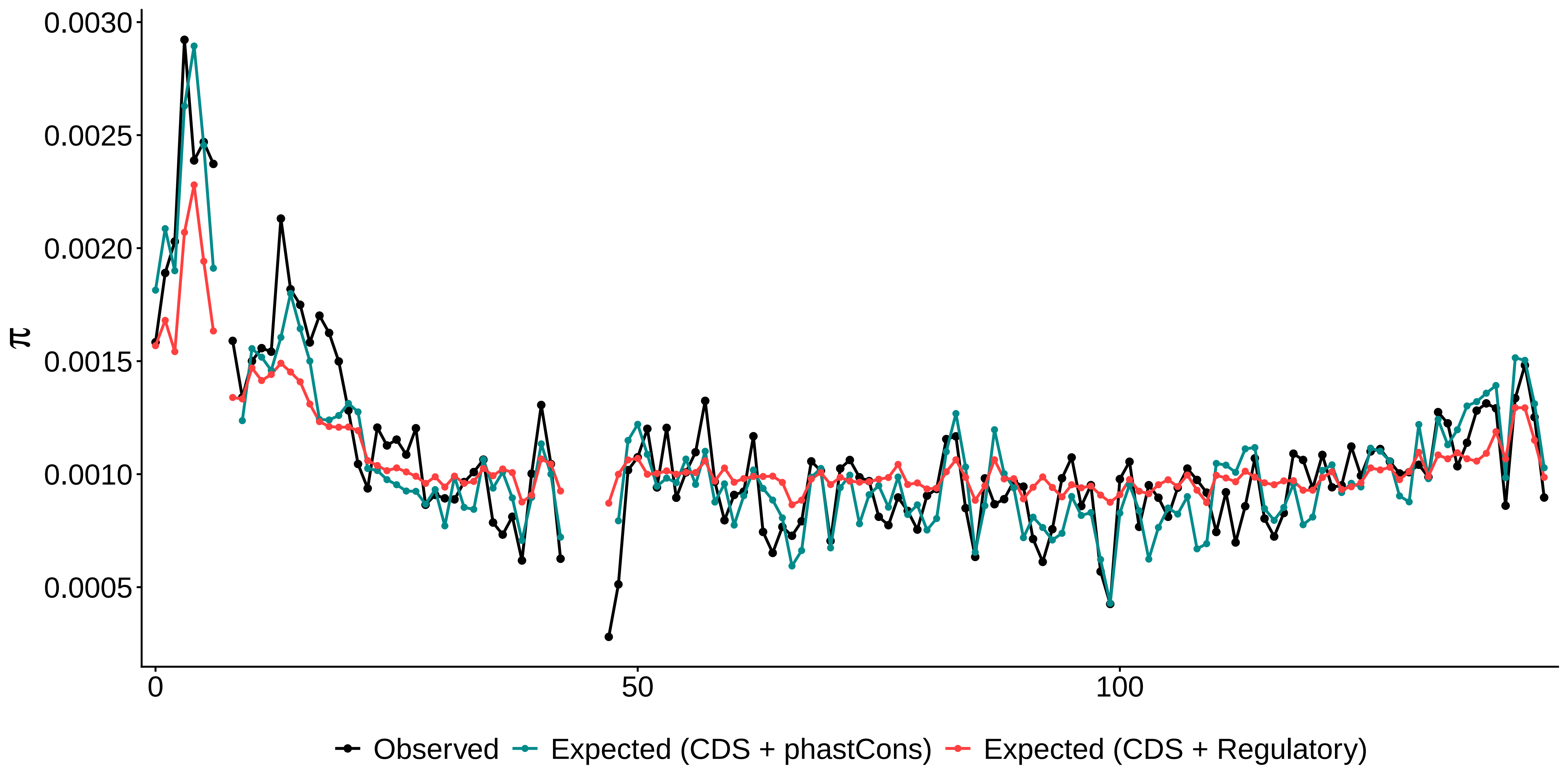

### pi_c9_1Mb.pdf

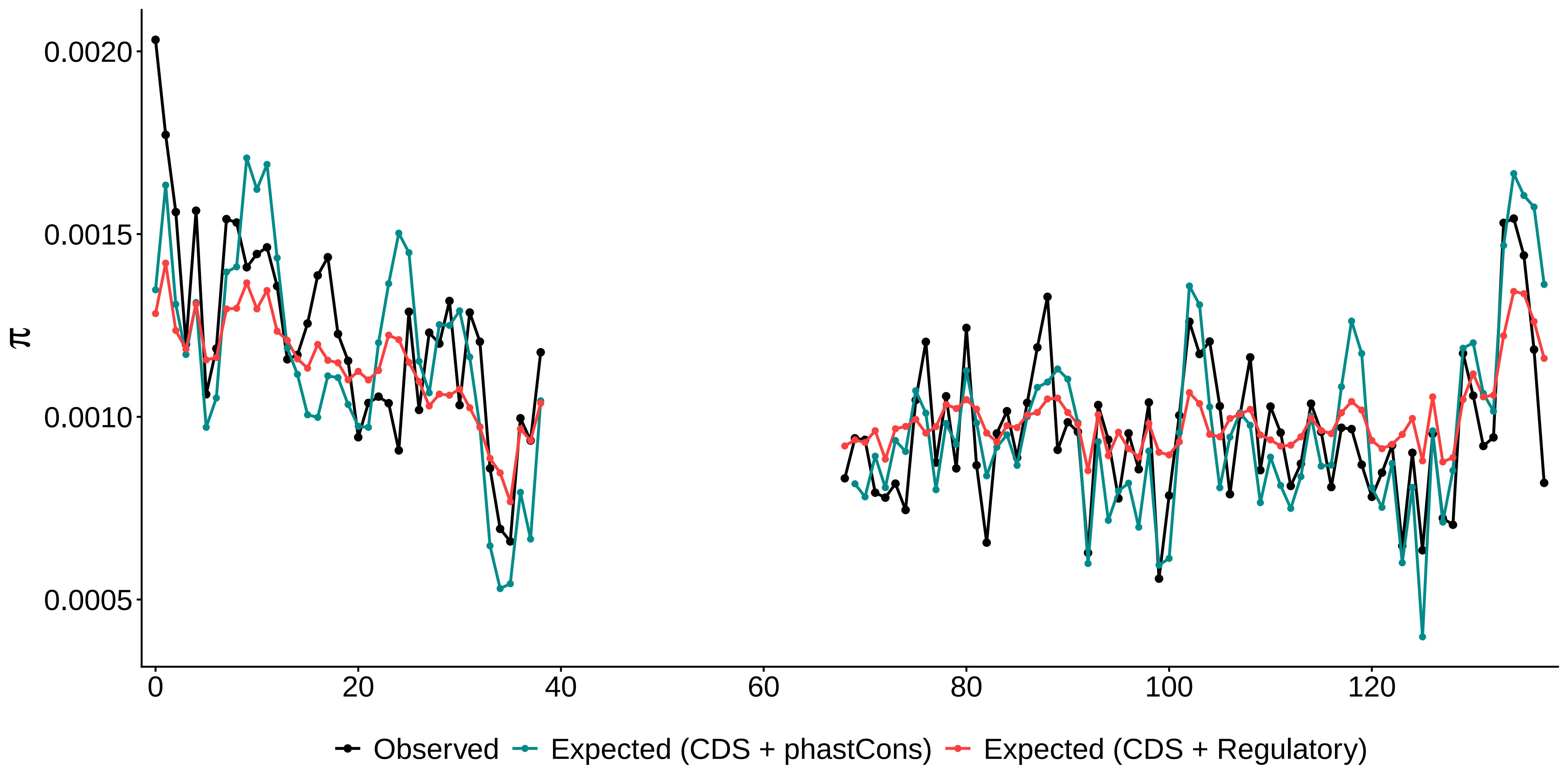

### pi_c10_1Mb.pdf

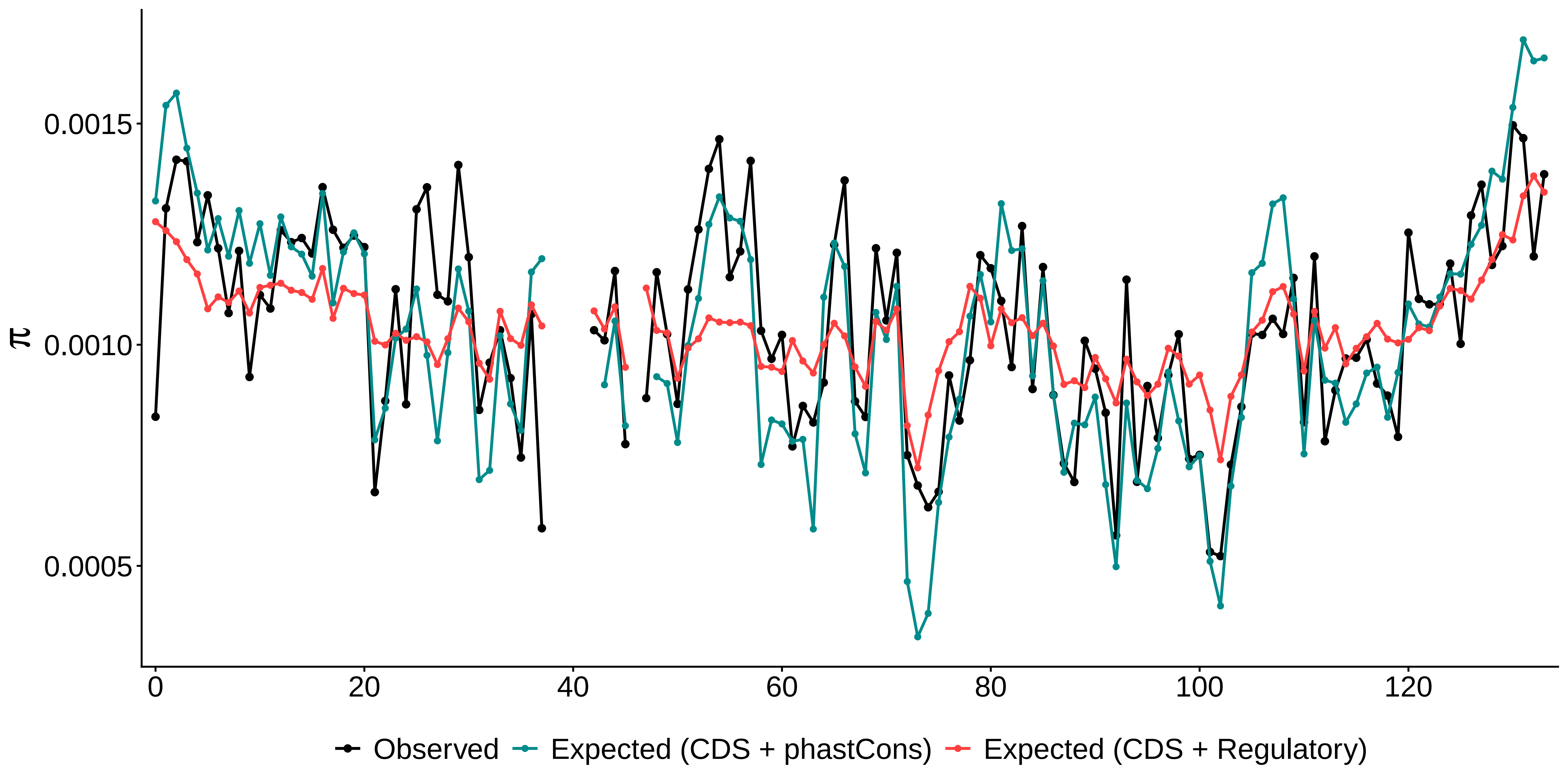

### pi_c11_1Mb.pdf

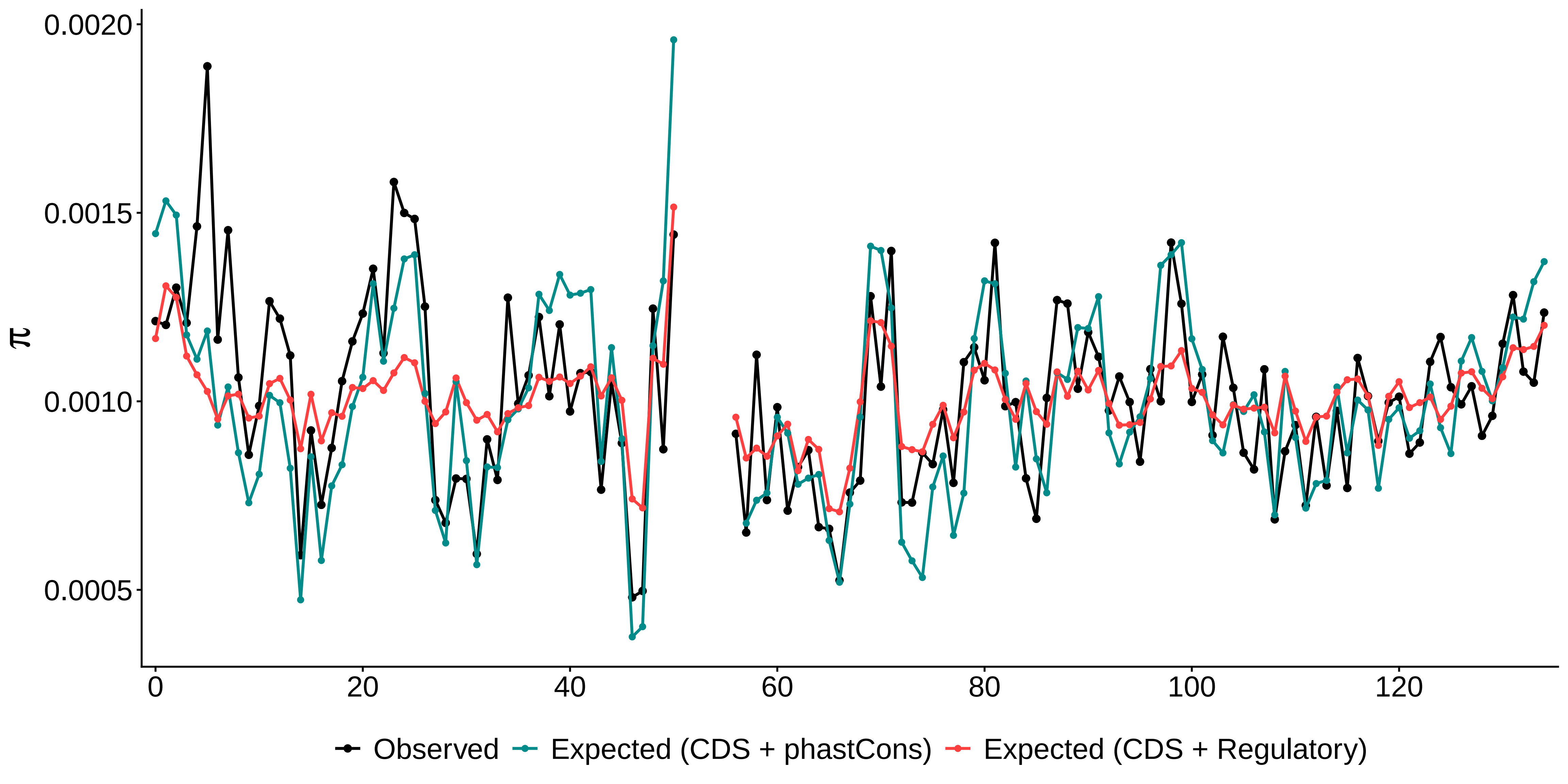

### pi_c12_1Mb.pdf

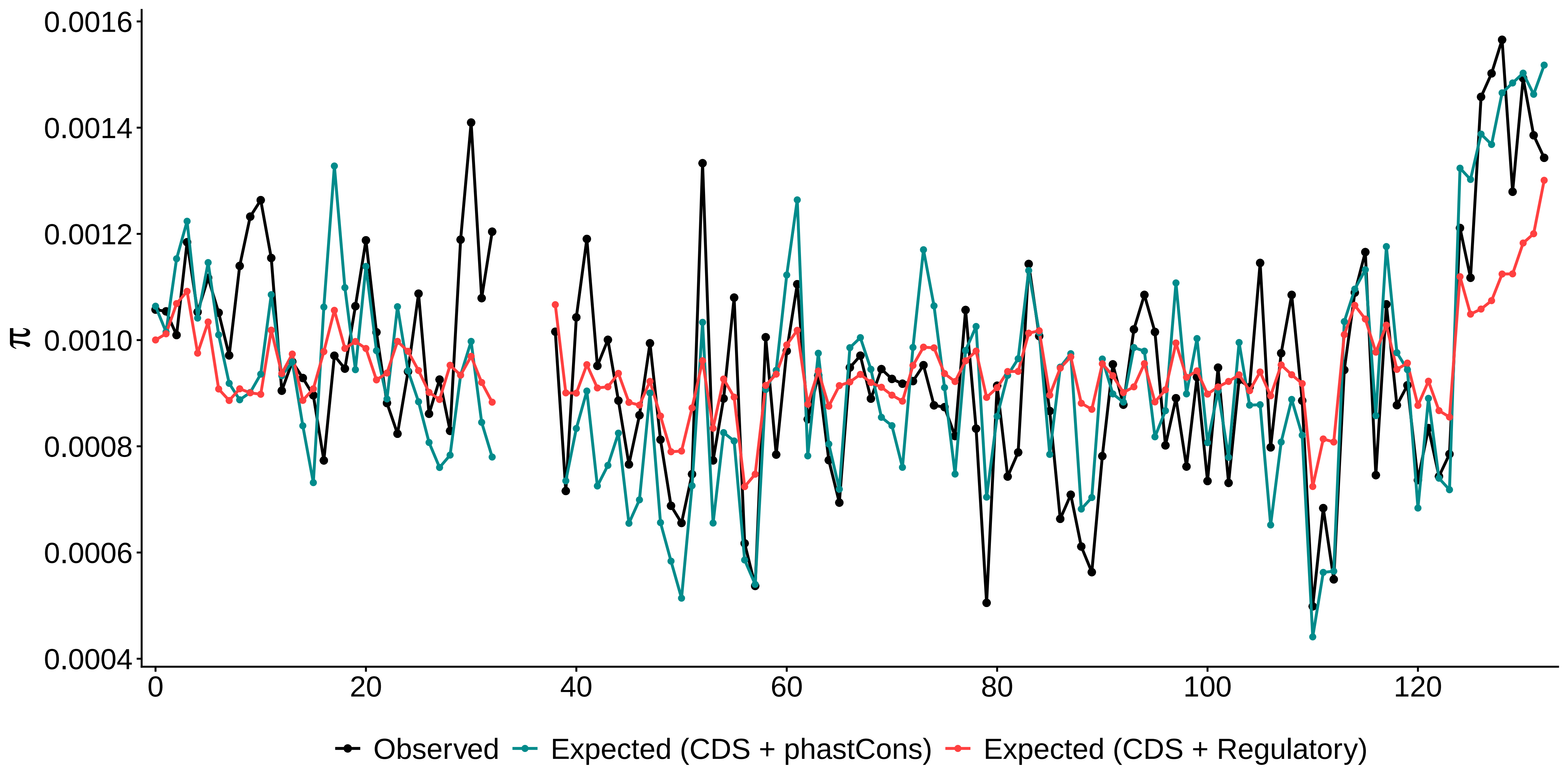

### pi_c13_1Mb.pdf

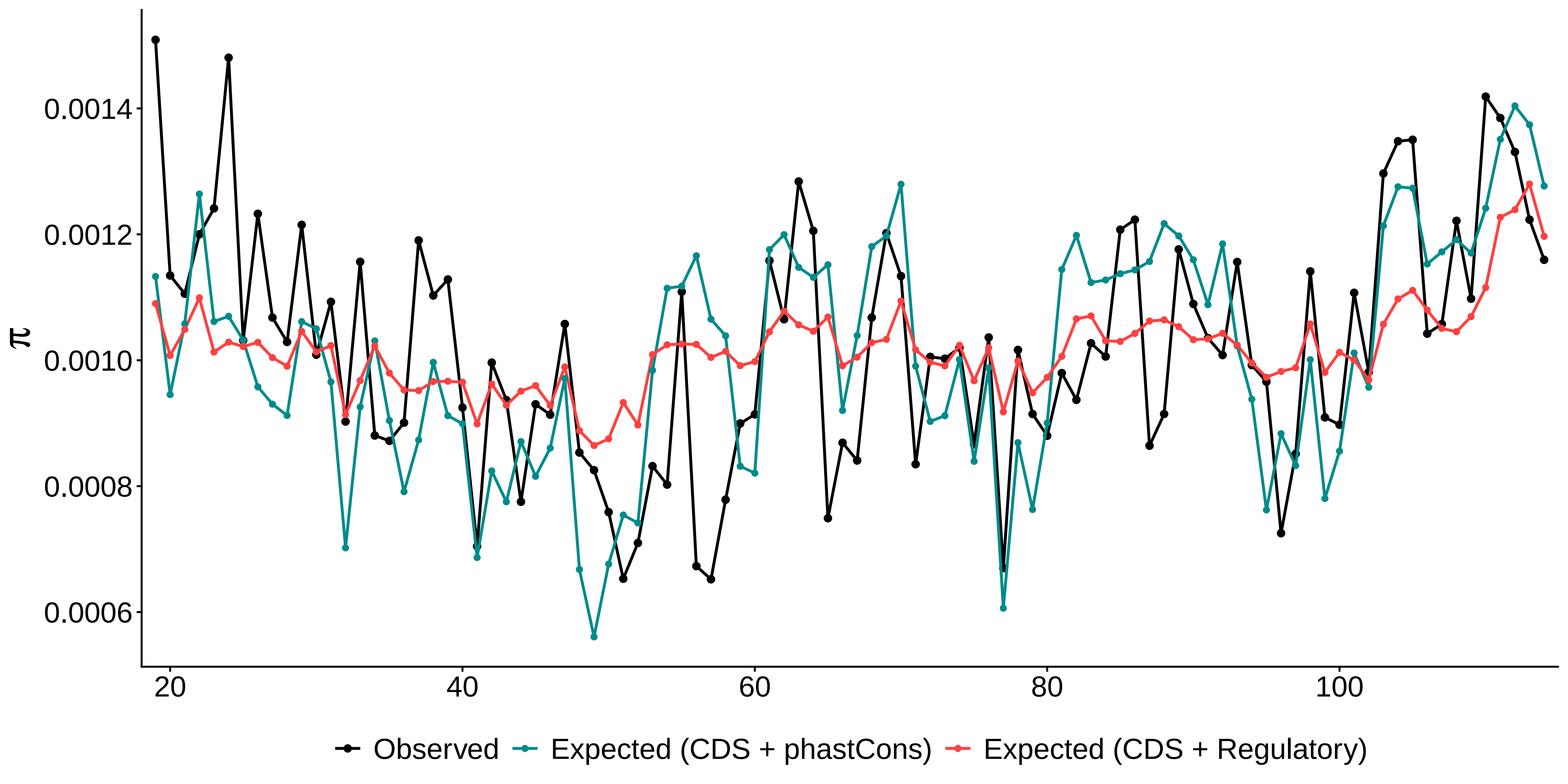

### pi_c14_1Mb.pdf

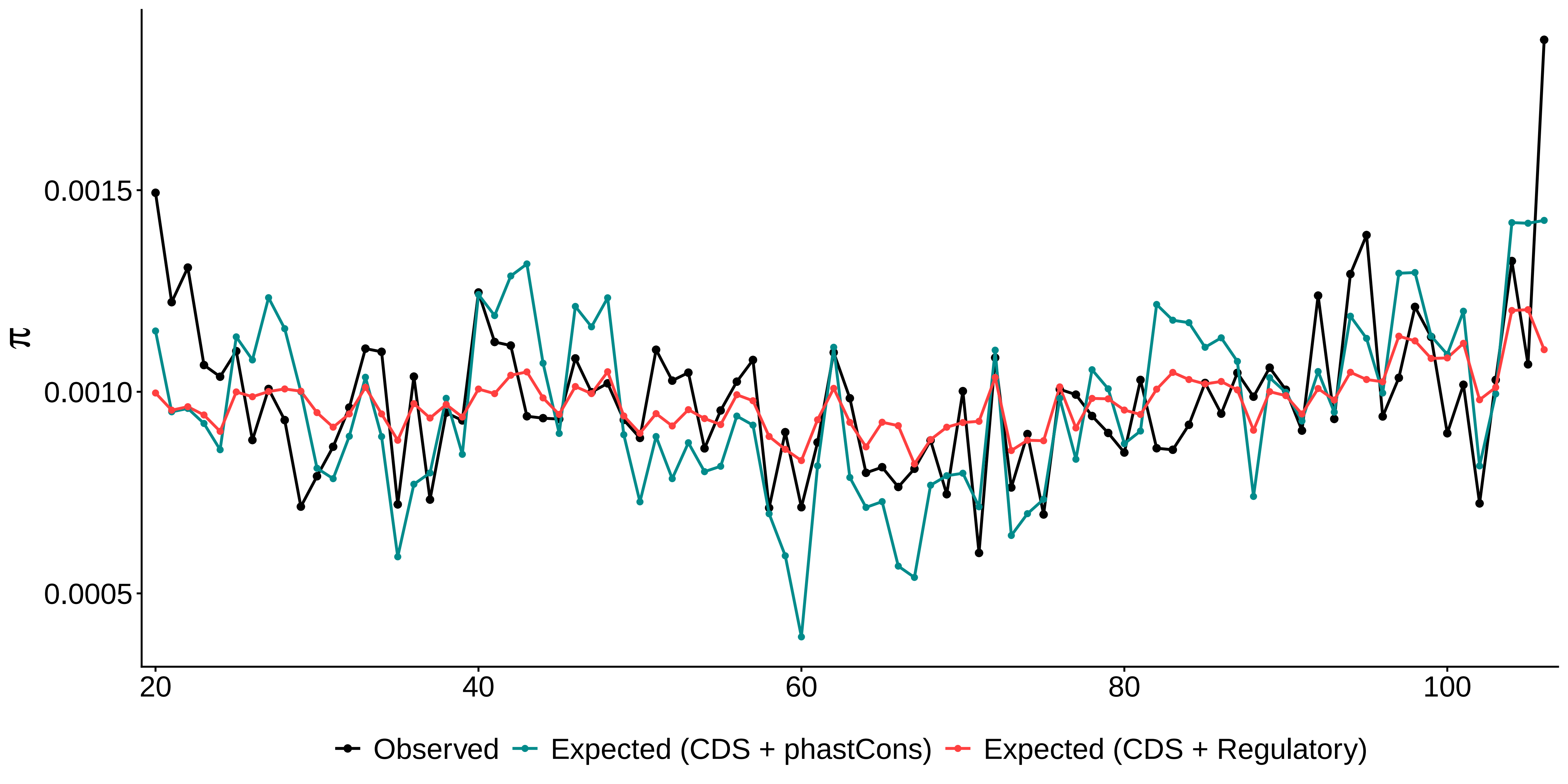

### pi_c15_1Mb.pdf

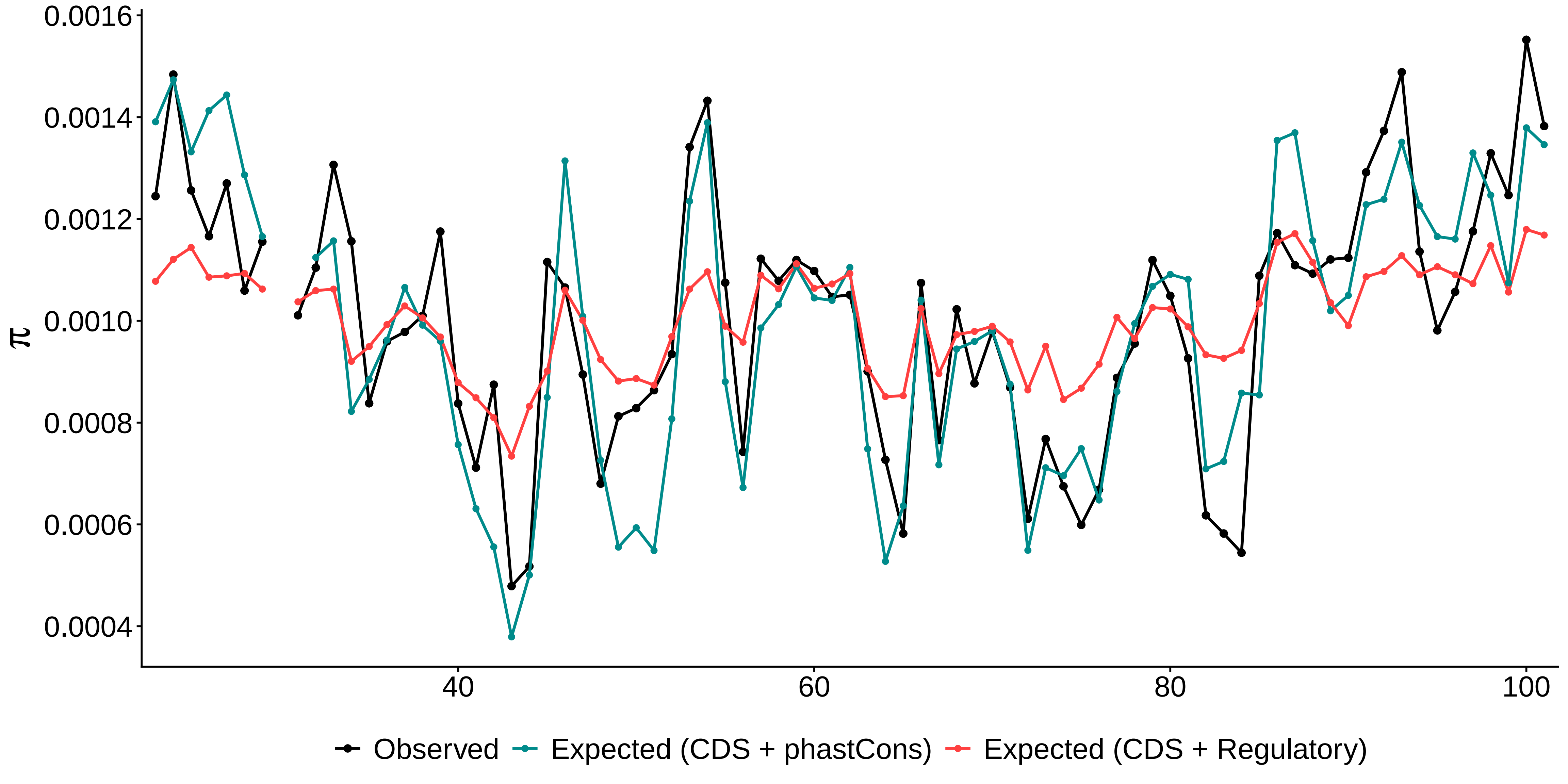

### pi_c16_1Mb.pdf

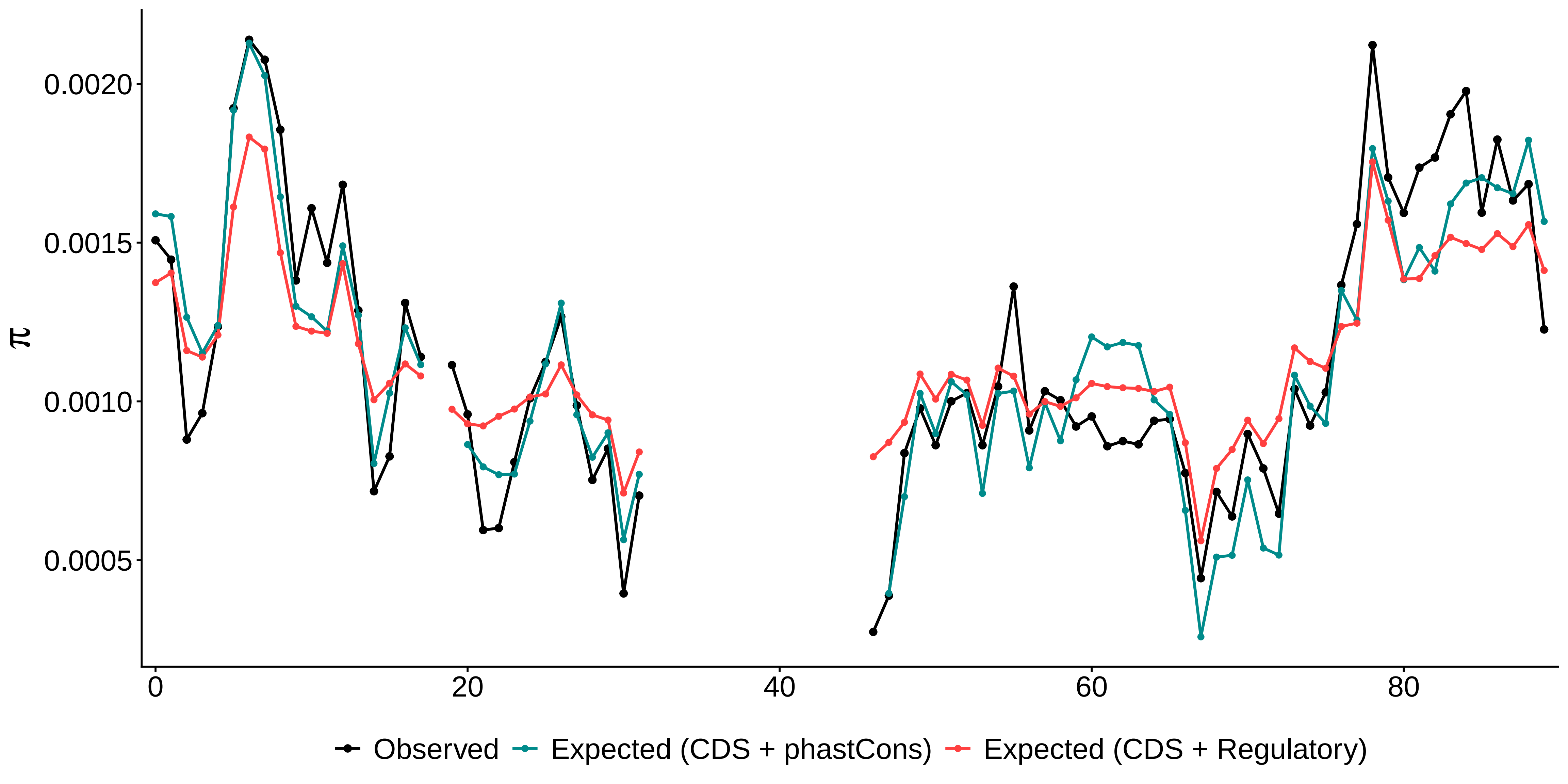

### pi_c17_1Mb.pdf

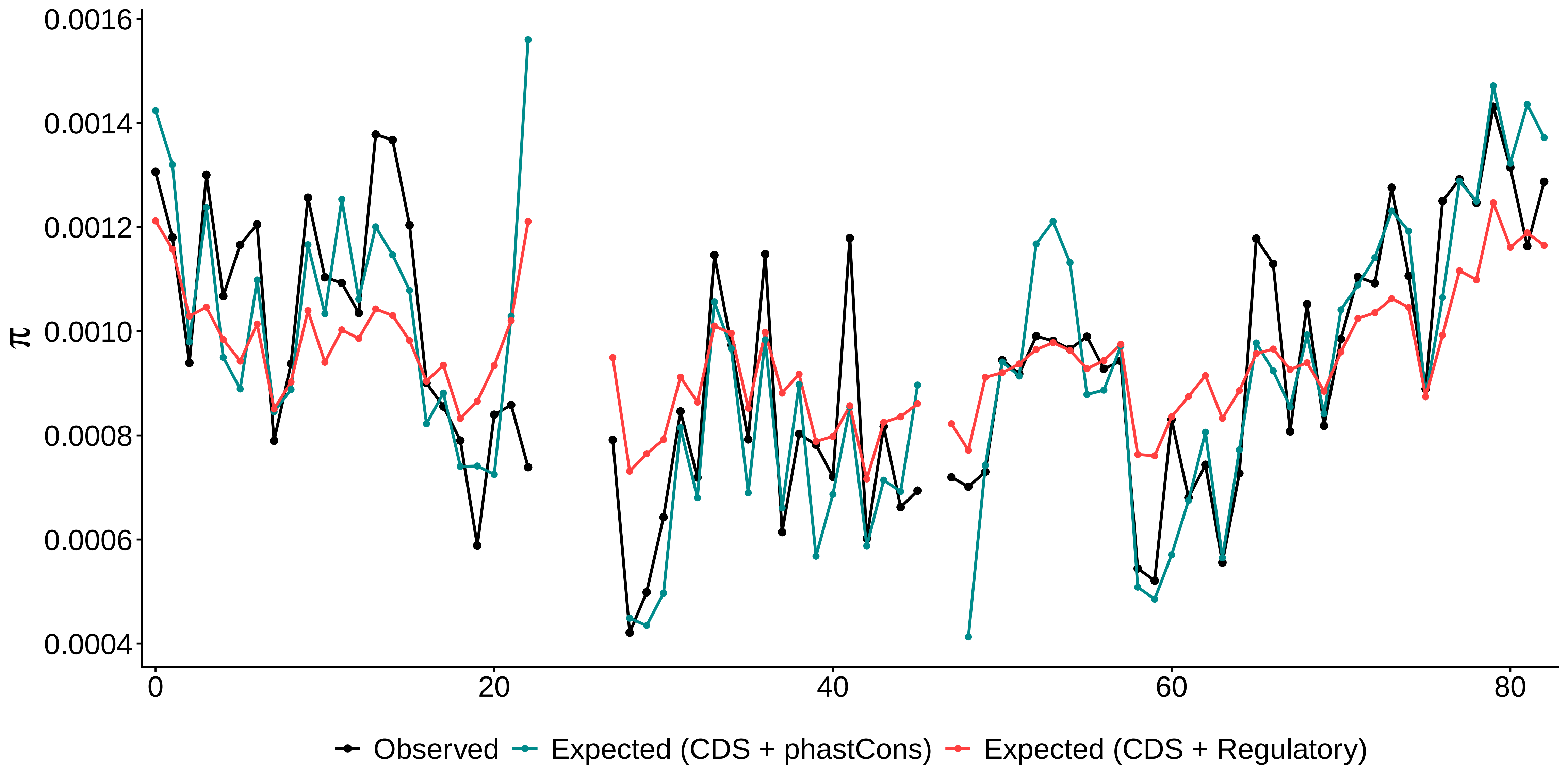

### pi_c18_1Mb.pdf

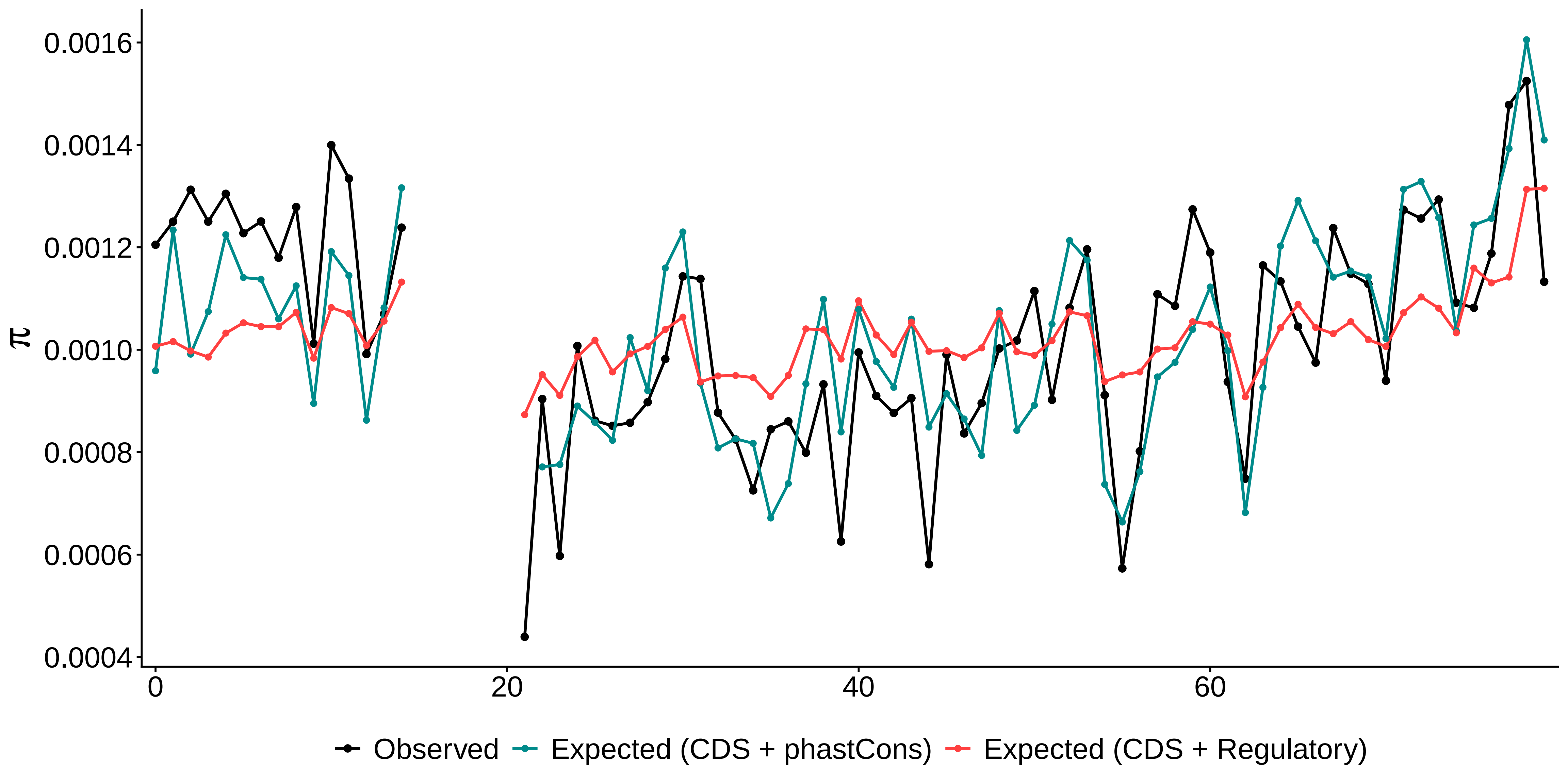

### pi_c19_1Mb.pdf

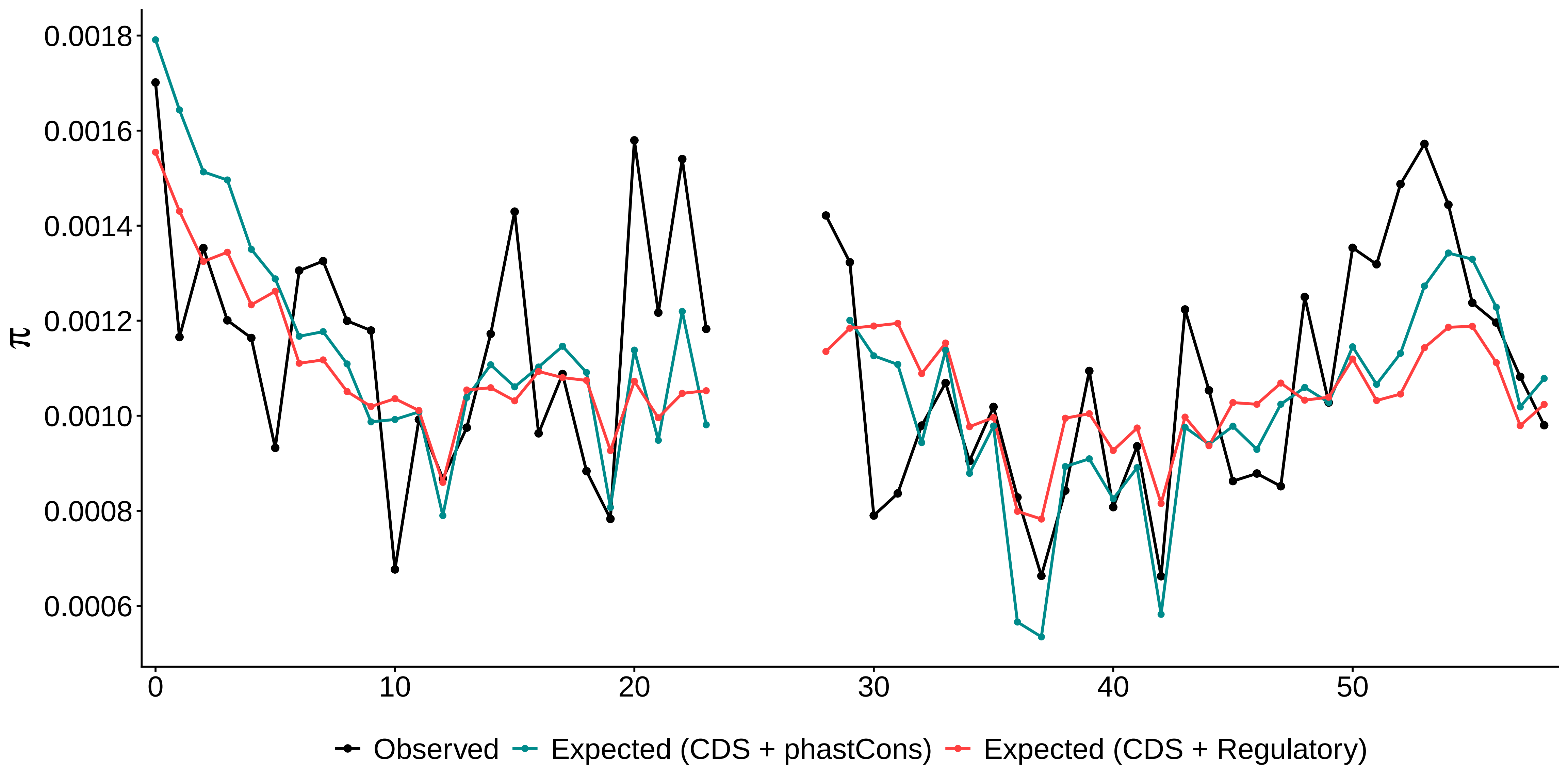

### pi_c20_1Mb.pdf

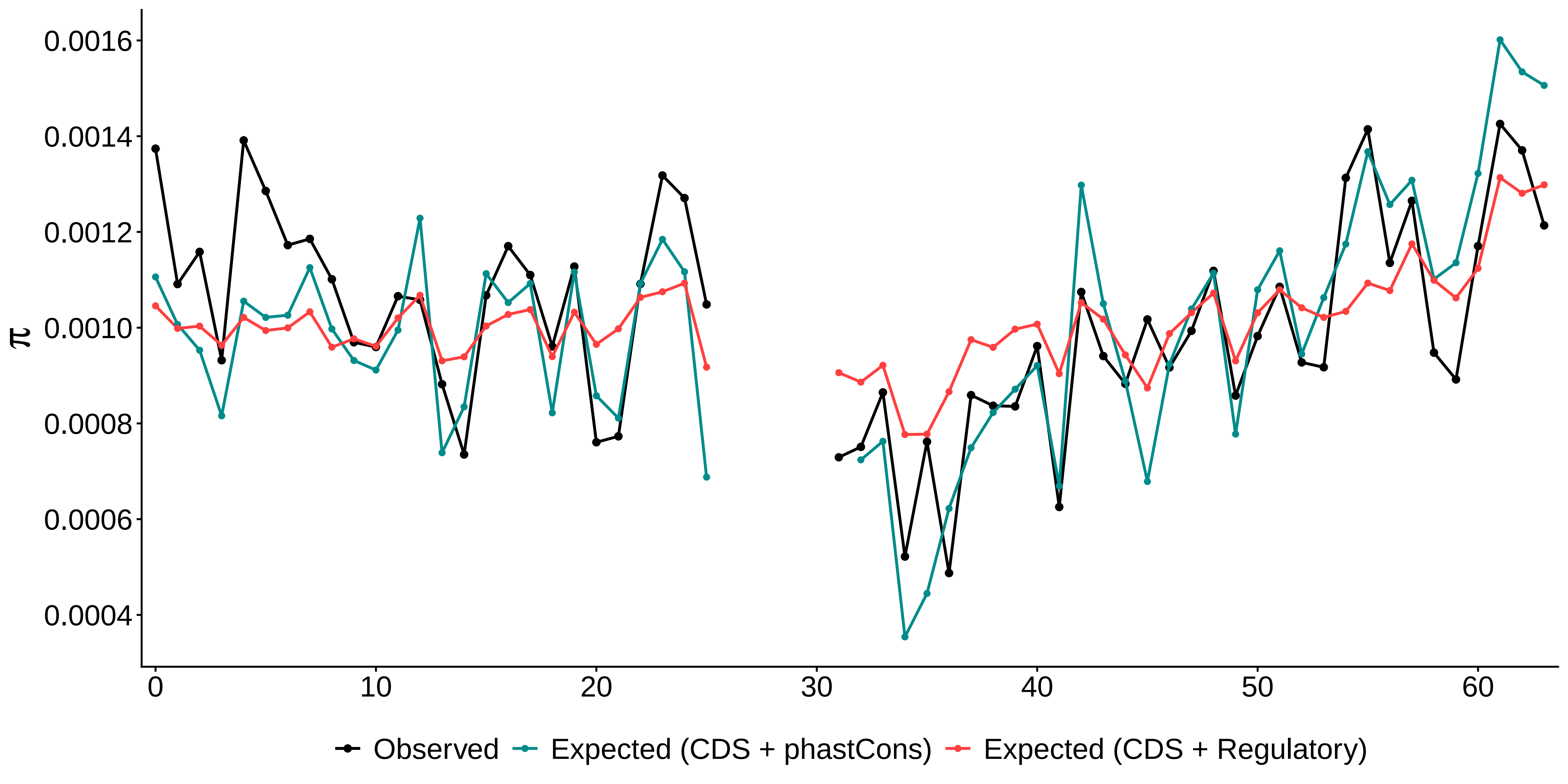

### pi_c21_1Mb.pdf

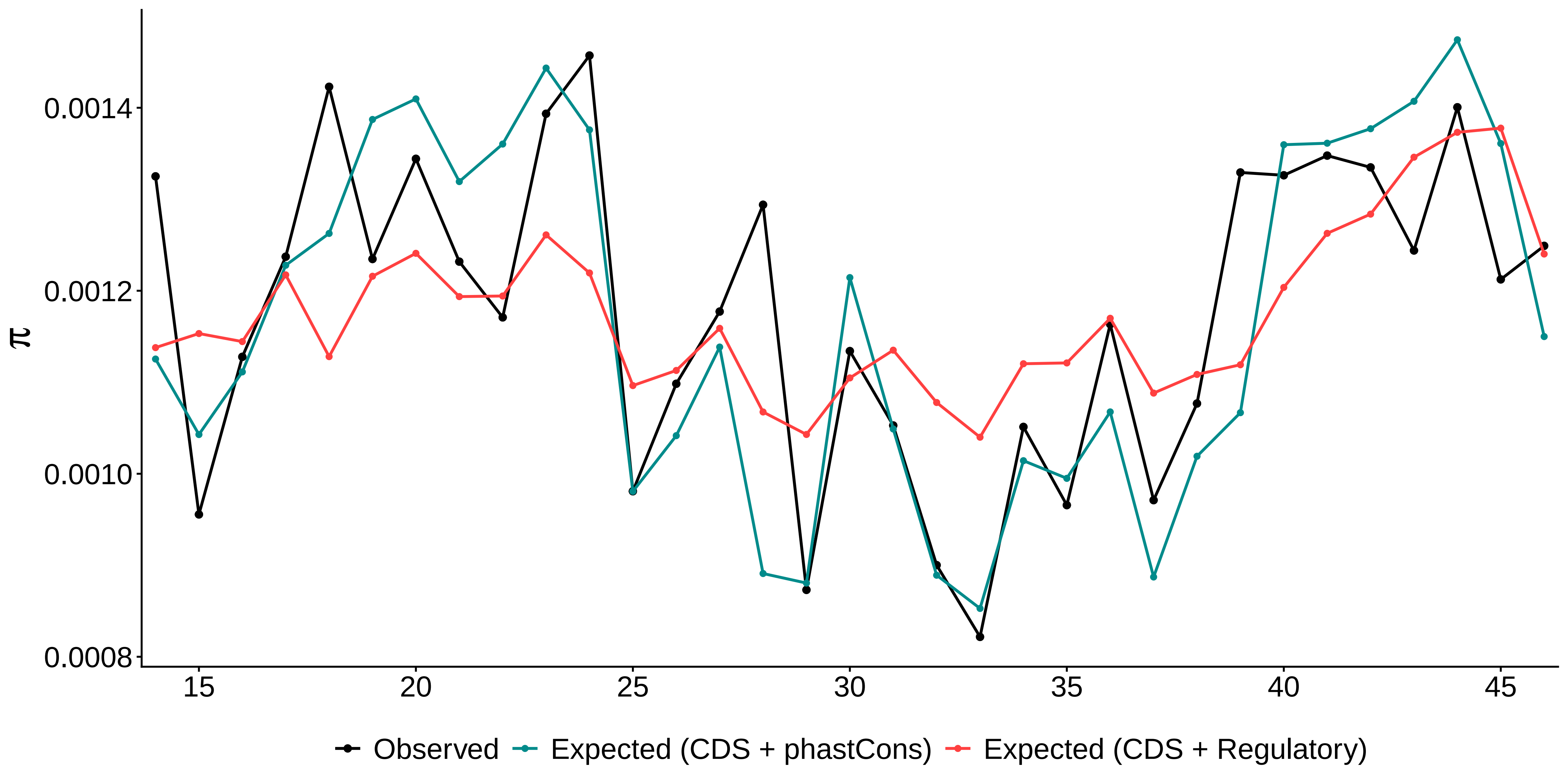

### pi_c22_1Mb.pdf

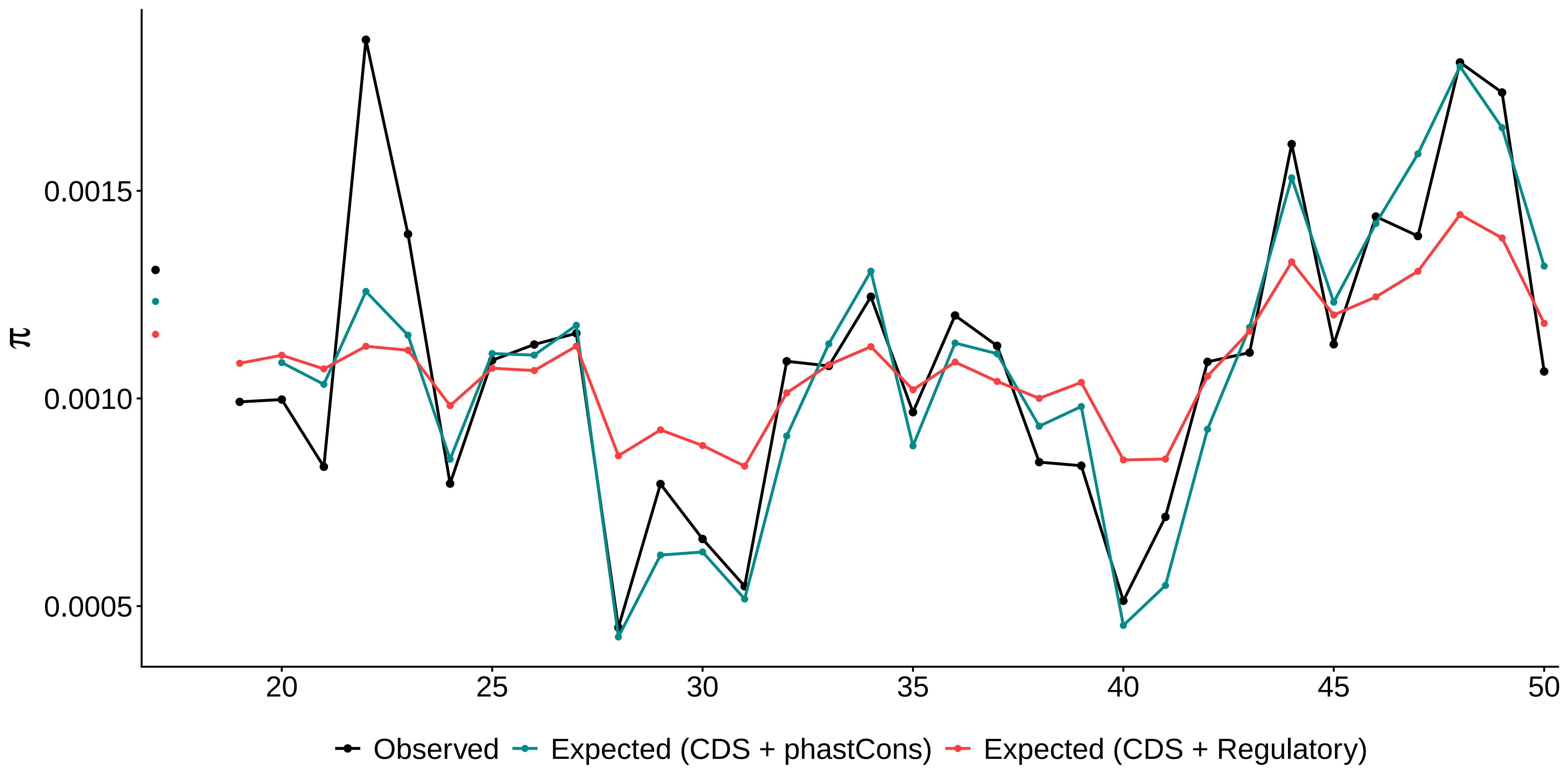
